# Ancient tree-topologies and gene-flow processes among human lineages in Africa

**DOI:** 10.1101/2024.07.15.603519

**Authors:** Gwenna Breton, Per Sjödin, Panagiotis I. Zervakis, Romain Laurent, Alain Froment, Agnès E. Sjöstrand, Barry S. Hewlett, Luis B. Barreiro, George H. Perry, Himla Soodyall, Evelyne Heyer, Carina M. Schlebusch, Mattias Jakobsson, Paul Verdu

## Abstract

The deep history of human evolution in Africa remains intensely debated, with increasingly complex models being proposed. To investigate this, we sequenced and analysed 73 novel high-quality whole genomes from 14 Central and Southern African populations with diverse cultural practices. Using extensive simulations and machine-learning Approximate Bayesian Computation (ABC), we jointly reconstructed their demographic history of divergences and migrations. We find extensive genome-wide diversity within and among populations, including substantial local genetic differentiation not fully explained by geography or cultural practices. These patterns highlight the importance of explicitly considering local genomic diversity when reconstructing human evolutionary history. We find that tree-like population histories with long periods of drift separated by short pulses of unidirectional gene-flow better explain the data than continuous gene-flow. Without invoking archaic admixture, our models accurately fit observed genomic variation and identify multiple episodes of gene-flow coinciding with major ecological and cultural changes in Sub-Saharan Africa.

## Introduction

Unraveling genetic structure and gene flow among human lineages in Africa is crucial to the understanding of the biological evolution and diversity of *Homo sapiens* throughout the continent ^1–3^.

Population geneticists largely agree today that *Homo sapiens* spent a large part of its genetic evolution within Africa only, between its gradual emergence from anatomically archaic forms 600.000-200,000 years ago and the beginning of the Out-of-Africa expansions to the rest of the world around 100,000-50,000 years ago ^2^. For the past 20 years, in various regions of Africa, the detailed demographic and evolutionary history that shaped the genetic diversity of extant populations has been the subject of numerous investigations ^3–18^.

However, how ancient genetic divergences and the dynamics of admixture and/or migration events among ancient human lineages influenced the genetic landscape observed today throughout Africa largely remains to be assessed ^19^. Classically tested demographic models ^1,20–22^, range from a unique ancestral *Homo sapiens* genetic population having recently and rapidly diverged into extant African populations via series of founding events and/or multifurcations (often referred to as tree-like models), to models where extant populations diverged in a remote past and remained isolated over long periods of time until relatively recently (often referred to as within Africa multiregional models). Moreover, each model may encompass possible gene exchanges among lineages via migration or admixture processes. Importantly, the timing, duration, and/or intensity of such gene-flow events may fundamentally change the most likely topology of bifurcating tree-like models compared to results obtained without gene flows, and may also create reticulations among pairs of lineages throughout history ^22^. Finally, these classical albeit highly complex models have recently been enriched with possible introgression events from now extinct, unknown, or unsampled non-*Homo sapiens* or ancient “ghost” *Homo-sapiens* lineages ^3,22–26^. Such latter models became plausible in Africa in analogy to the likely events of introgressions from now extinct hominid species into *Homo sapien*s lineages unveiled outside of Africa by the major advances of paleogenomics ^27,28^.

In this context, several recent investigations tested a variety of the above models with maximum-likelihood approaches relying on whole genome sequences from several extant populations sampled throughout the continent. They often reached highly contrasted results, with different ancient tree-topologies between different numbers of ancient lineages ^2,22,24–26^. Furthermore, they found that reticulations among a limited number of ancient lineages explained the observed genetic patterns without the necessity of archaic introgressions ^22^; or, instead, that admixture with other non-*Homo sapiens* or ancient *Homo sapiens* “ghost” populations most satisfactorily explained the results ^23–26^.

A possible source of such vast differences among obtained results is likely the variety of population sets considered in each study at a continental scale. Indeed, genetic diversity and differentiation among populations is maximal in Africa ^3,29,30^, and previously shown to be due to substantial differences in local demographic histories (e.g. ^4,6,9,11^). Therefore, representing vast regions of the continent by a single population ^22,24^, or merging data from differentiated groups based on shared genetic ancestry ^26^, possibly influenced results obtained across studies and their interpretations.

In addition, and most importantly, all above models are highly nested, i.e. overlapping and genetically largely indistinguishable for vast spaces of model-parameter values. Indeed, depending on values of divergence times, effective population sizes, timing, duration and intensity of migration or admixture events among lineages, tree-like, reticulated, and even multiregional models, with or without ancient admixture, may be hardly identifiable. This essential methodological difficulty is well illustrated by the often relatively similar values of posterior likelihoods of the various competing models, thus indeed proved empirically hard to discriminate with the maximum-likelihood approaches deployed, even more so when competing models differ in the number and specifications of parameters explored with different statistics. These fundamental statistical issues remain despite the use of high-quality whole genomes, an increasing number of populations considered at once, and increasingly powerful methods based on innovative statistics ^22,24,26,31–33^. Finally, the lack of ancient DNA data older than a few thousand years in Africa, strongly hampers the empirical testing of the possible occurrence of ancient *Homo sapiens* “ghost” or non-*Homo sapiens* introgression events in *Homo sapiens* lineages in Africa ^22–25,30^.

In this context, we investigated 74 high coverage (>30x) whole genome sequences from an anthropologically well-characterized sample of hunter-gatherer, herder, and agricultural neighboring populations from Central and Southern Africa (**Figure 1**, **Table 1, Table S1**), merged together with 105 previously published high-quality whole genomes ^27,34–37^. In particular, we investigated genomic diversity patterns and aimed at reconstructing ancient demographic histories among lineages having led to different groups of Khoe-San hunter-gatherer or herder populations from Southern Africa ^6^, and to different groups of both Eastern or Western Congo Basin hunter-gatherer populations (often historically designated “Pygmies” by Europeans) and their agriculturalist (“non-Pygmy”) neighbors today with whom they share complex socio-economic interactions ^4,5,38^. Investigating detailed ancient demography and gene-flow among these three groups of populations, taking explicitly into account local differentiations and possible gene-flow among and within groups, is of particular interest as they have previously been identified to be among the most deeply diverged lineages in our species ^2,3,29,39,40^.

**Figure 1:**
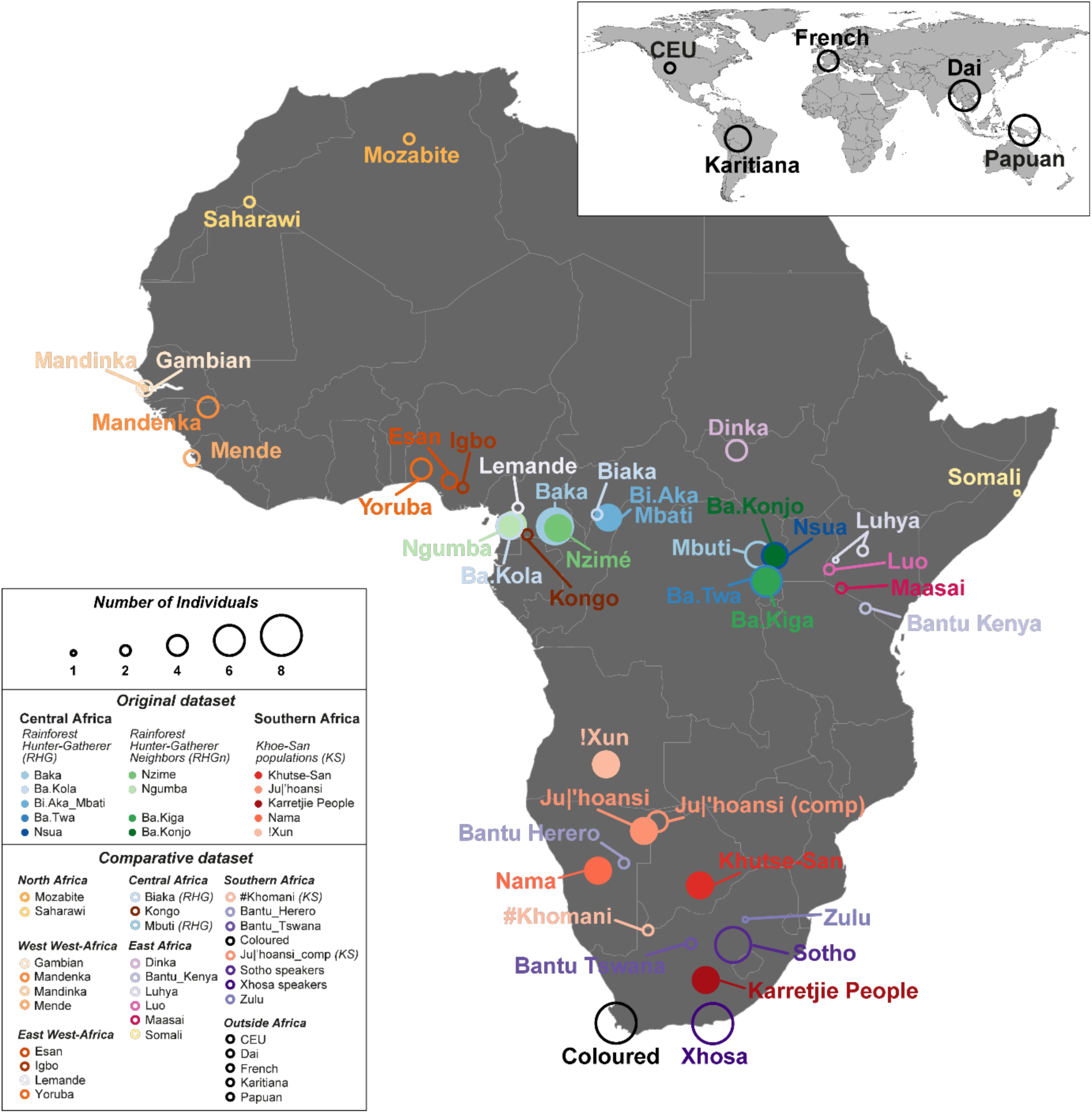
Sampling location and number of whole-genome sequenced individuals. Individuals and populations originally sequenced in this work are indicated with filled circles. Individuals and populations from previously published data merged with the original data set for comparison purposes are indicated with open circles. Circles’ diameters are proportional to the number of individuals as indicated in the top row of the legend in the bottom left corner. Note that “Ju|’hoansi_comp” refers to the genomic dataset for the Ju|’hoansi previously published by the SGDP and HGDP and used for comparison to the original data generated here for another sample of the Ju|’hoansi ethno-linguistic group. Detailed information about population samples is provided in **Table 1**, **Table S1**, and **Material and Methods**.

**Table 1:**
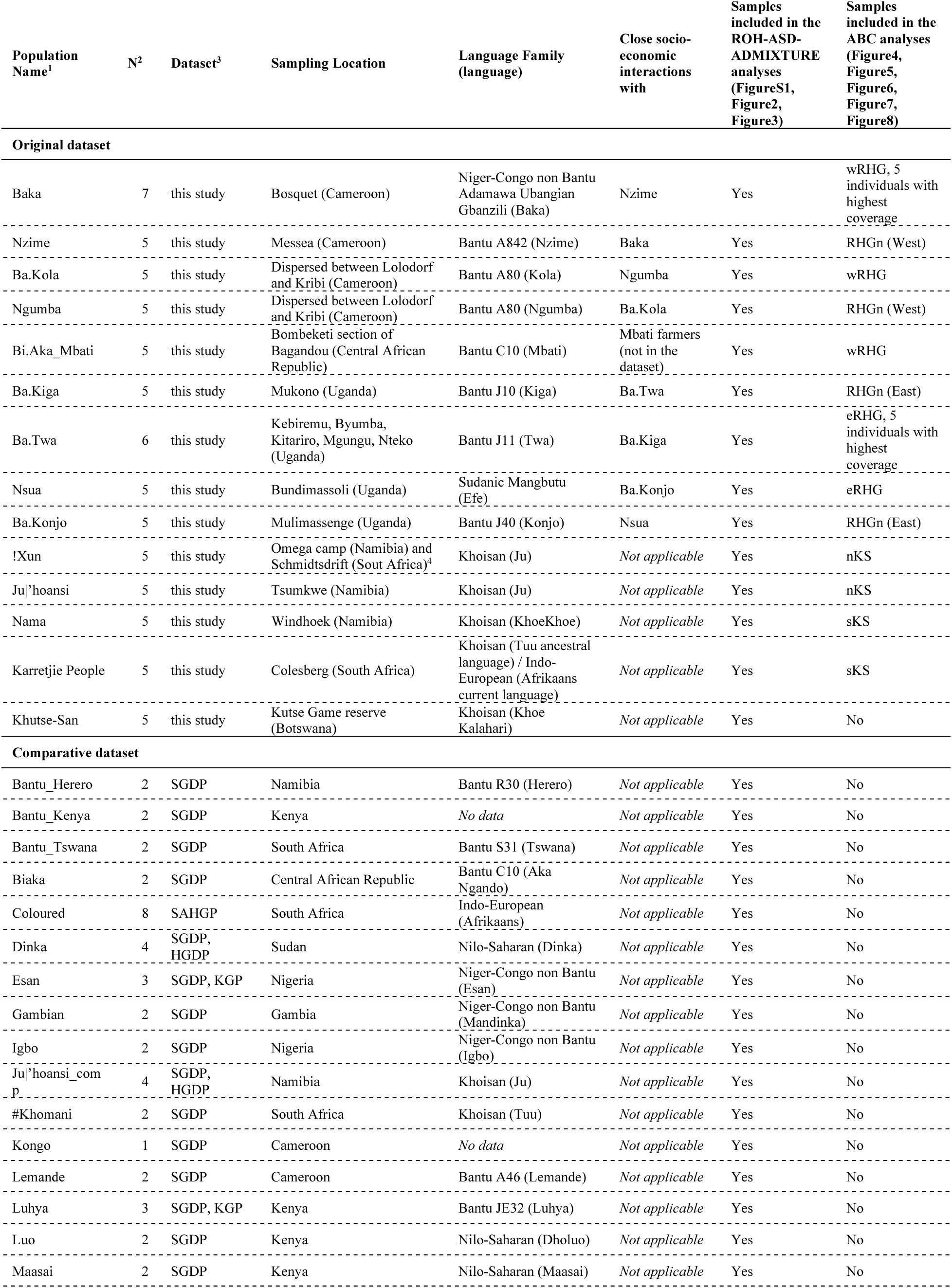
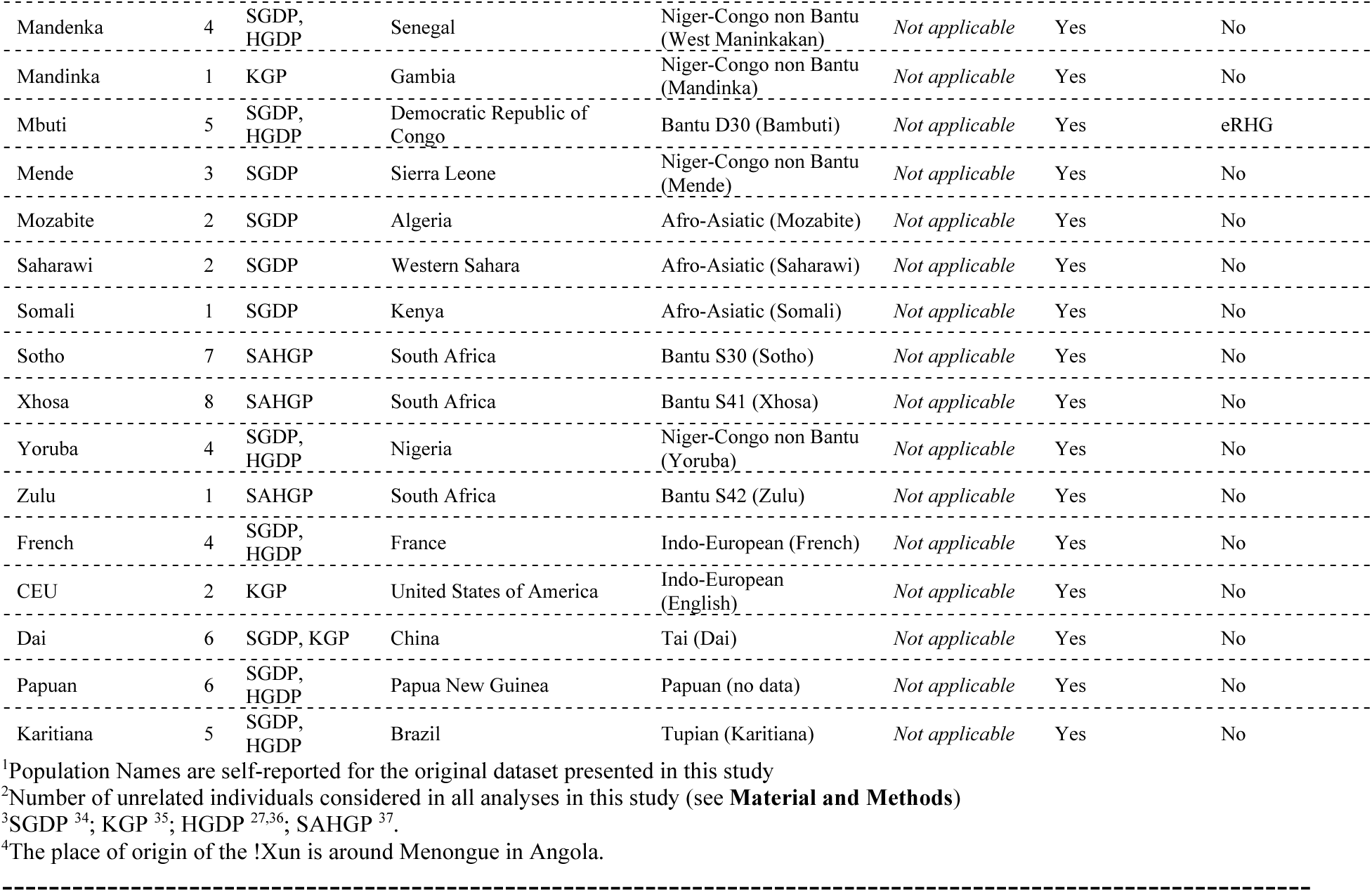
Population table. Geographical location of population samples are indicated in **Figure 1**. Complementary information for genetic-geography and ANOVA analyses are provided in **Table S1**.

We conducted formal statistical choice of competing scenarios and subsequent joint estimation of parameters under the best scenario using machine-learning Approximate Bayesian Computation (ABC) ^41–45^, a methodological approach fundamentally differing from all maximum-likelihood methods previously deployed, and only rarely explored in previous studies reconstructing ancient demography in Africa ^24^. ABC allows us to compare a range of complex demographic scenarios presumed to have led to observed genetic patterns in numerous population samples, and to estimate posterior parameter distributions best mimicking the data under the best scenario (e.g. ^4,18,24^). Briefly, this is achieved based on informative summary-statistics computed on the observed data and on numerous explicit genetic simulations for which scenario-parameters are drawn randomly in large prior distributions set by the user. We thus aimed at jointly inferring competing tree-topologies, divergence times, effective population size changes, and timing, duration and intensity of possible asymmetric gene-flow events, to reconstruct the detailed evolutionary mechanisms underlying the genomic diversity of extant Central and Southern African populations.

## Results

We generated high-coverage whole genome sequences (>30X) in 74 individuals from 14 Central and Southern African populations for whom detailed ethno-anthropological information was gathered in the field jointly with DNA samples (**Figure 1** and **Table 1**). We merged this original dataset with a comparative dataset comprising 105 individuals from 27 African and five non-African populations for whom similar high-quality raw whole genome sequencing data were made available to the community ^27,34–37^. After quality-control, variant-calling, and relatedness filtering procedures conducted on all 179 individuals together, we retained 73 and 104 family unrelated individuals in our original and comparative datasets, respectively, for all subsequent analyses (see **Material and Methods**).

Considering only the 73 Central and Southern African unrelated individuals newly sequenced here, we identify a total of 26,780,319 biallelic SNPs (241,428 multiallelic SNPs), and 2,454,965 simple insertions/deletions (969,025 complex indels), compared to the reference sequence of the human genome GRCh38 for autosomes (chromosomes 1 to 22), out of which 854,114 (3.1893%) and 114,362 (4.6584%), respectively, were not previously reported in dbSNP 156 (**Table S2**). Among the 177 unrelated worldwide individuals including our original dataset, we identify 36,272,545 biallelic SNPs (257,906 multiallelic SNPs), and 3,159,306 simple insertions/deletions (977,542 complex indels), out of which 1,055,245 (2.9092%) and 189,124 (5.9863%), respectively, were not previously reported in dbSNP 156 (**Table S2**).

Overall (**Figure S1A, Table S3**), Southern African Khoe-San populations exhibit the highest mean number of biallelic SNPs as well as numbers of previously unreported SNPs, followed by Central African Rainforest Hunter-Gatherer populations from the Congo Basin, and then by all other African populations in our dataset. These results show that a substantial number of previously unknown variants can still be found when investigating high-quality whole genome sequences from relatively under-studied Sub-Saharan African populations, consistent with previous studies ^22,26,27,34–37,40,46^.

## 1. Large genetic diversities and differentiations in Central and Southern Africa

### 1.A. Heterozygosities and Runs of Homozygosity

We observe significant differences across populations at a local scale in Central and Southern Africa in unbiased heterozygosities ^47^, calculated on both varying and non-varying autosomal sites (**Figure S1A**, **Figure S1B**, and **Table S3**. We find overall more variation in Rainforest Hunter-Gatherer (RHG) populations from the western part of the Congo Basin than in RHG populations from the East (Wilcoxon one-sided test p-value=0.00328 for all RHG populations in our dataset and 1.641×10^-5^ considering only those with original data). Furthermore, note that the Baka and Ba.Kola Western RHG populations exhibited significantly larger genetic diversities than their Nzime and Ngumba RHG neighbors (Wilcoxon one-sided test p-value=0.04637). Finally, observed genetic variation across Northern Khoe-San (KS) populations (!Xun, Ju|’hoansi, Ju|’hoansi_comp) and Southern KS populations (Nama, Karretjie People and #Khomani) was not significant (Wilcoxon two-sided test p-value=0.7726). Note that unbiased estimates of heterozygosities significantly corelate positively (Spearman rho=0.8490, p-value=2.2×10^-16^) with genome-wide numbers of autosomal biallelic SNPs (**Figure S2**).

In addition, the distribution of Runs of Homozygosity (ROH) sizes has been shown to be highly informative about important demographic processes such as levels of inbreeding or endogamy within populations, and reproductive isolation or recent admixture among populations ^18,48–50^. In general (**Supplementary Note**, **Figure S1C**), we find highly varying patterns of ROH depending on ROH length-classes at regional and local scales, within and between groups of Central and Southern African populations.

Altogether, our heterozygosity and ROH results show the vast genetic diversity of Sub-Saharan African populations, as well as substantially diverging genomic patterns at a local geographical scale, in particular among and within groups of Rainforest Hunter-Gatherers and RHG neighbors.

### 1.B. Individual pairwise genetic differentiation

We investigated individual pairwise genomic differentiation across the 177 worldwide unrelated individuals in our dataset, including the 73 unrelated novel Central and Southern African individuals, using Allele-Sharing Dissimilarities (ASD, ^51^). The Neighbor Joining Tree (NJT, ^52,53^) representation of the pairwise ASD matrix shows three main clusters among all African individuals, separated by a longer genomic distance from all non-African individuals (**Figure 2A**, **Figure S3**). One cluster corresponds to all Southern African Khoe-San populations (represented by the pink and red symbols, see **Figure 2B**), largely separated from a cluster corresponding to Rainforest Hunter-Gatherer populations from the Congo Basin (represented by the blue symbols); itself also largely separated from a third cluster grouping all other Western, Eastern, Southern and Central African populations in our data set, the latter group including Rainforest Hunter-Gatherer neighboring populations in green symbols (see, **Figure 2B)**.

**Figure 2:**
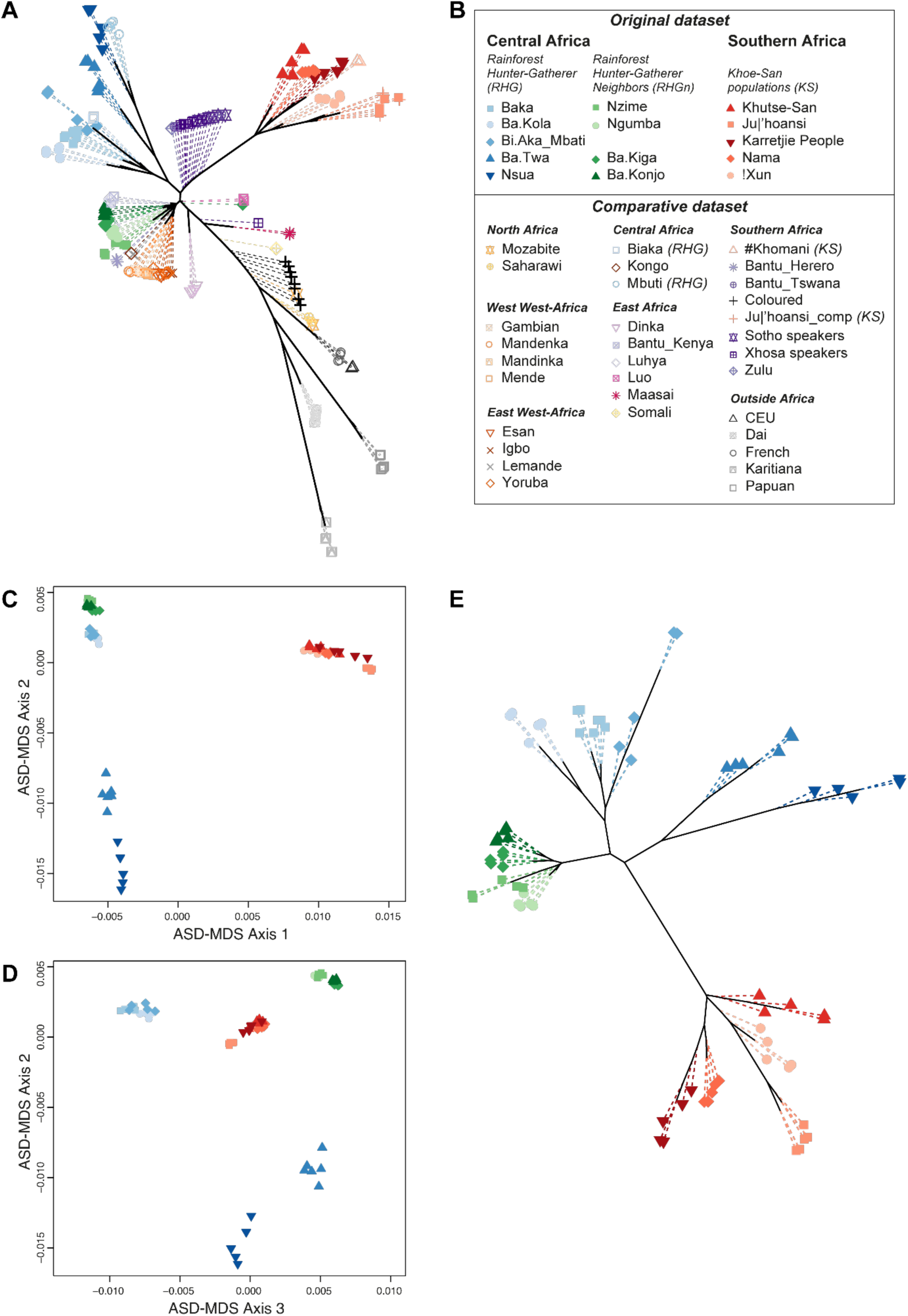
Individual pairwise genetic differentiation patterns. Neighbor-Joining Tree NJT ^52,53^ and Multi-Dimensional scaling representation of genome-wide Allele Sharing Dissimilarities ^51^ among pairs of individuals at worldwide and regional African scales. We considered the 14,182,615 genome-wide autosomal SNPs pruned for low LD (r^2^ threshold 0.1) to calculate ASD between all pairs of individuals. (**A**) NJT computed for all 177 worldwide individuals in the ASD matrix. For easing the visualization of the internal branches of the NJT, all terminal edges are represented in dotted lines each measuring 1/10th of their true size. See **Figure S3** for an MDS version of this ASD matrix. (**B**) Individual symbols and colors identifying the 73 Central and Southern African individuals originally whole-genome sequenced with filled symbols, and the 104 individuals from worldwide populations, including Africa, merged with the original data with open symbols. (**C**) and (**D**) First three axes of ASD-MDS computed on a subset of the full ASD matrix comprising only the individuals sequenced anew, whose symbols are provided in (**B**). (**E**) NJT computed on the same individual subset of the ASD matrix used for (**C**) and (**D**). Terminal edges of this latter tree are represented with dotted lines each measuring 1/30th of their true size.

Note, importantly, that individual pairwise dissimilarities at the genome-wide scale do not trivially reflect geography (**Figure 2**, **Figure S3**, **Table S4**), highlighting the large genetic differentiation found across African populations and the complexity of its distribution at both a local and continental scale. As can be seen in **Figure 2**, from West to East of the Congo Basin throughout Central Africa, RHG populations cluster separately from RHG geographic neighbors with whom they nevertheless share close socio-economic interactions. Central African RHGn are more genetically resembling certain other Western, Eastern, and Southern Africans than their immediate RHG geographical neighbors. Analogously, Southern African Khoe-San populations are also clustering together and much more distant from other geographically neighboring Southern Africans.

ANOVA analyses (**Table S5**, **Table S6**, **Table S7**), show that linguistic families, sub-linguistic families, and traditional food-production strategies (**Table S1**), respectively explained a significant proportion of the variance in average population pairwise ASD at the continental scale throughout Africa (partial *r*^2^ = 15.064%, 26.109%, and 9.021%, respectively, each with ANOVA permutation p <0.001), and throughout Sub-Saharan Africa only (partial *r*^2^ = 14.811%, 22.759%, and 10.010%, respectively, each with ANOVA permutation p <0.001), as reported previously (e.g. ^26^). However, most interestingly, we find more complex ANOVA results at regional scales. Indeed, we find that linguistic families and sub-linguistic families do not significantly explain average population pairwise ASD patterns in Central Africa nor in the Congo Basin, while traditional food-production strategies do so in both regions (**Table S5**, **Table S6**, **Table S7**). Conversely, we find that traditional food-production strategies do not significantly explain population differentiation patterns among Southern African populations, while, here, linguistic and sub-linguistic families do. Finally, and further exemplifying the complexity of genomic differentiation patterns at a local scale in Africa, note that neither linguistic categories nor food-production strategies significantly explain average population pairwise ASD variation among Eastern African populations in our dataset; a result that will need to be challenged in future studies considering additional population samples from this sub-region.

The Multidimensional Scaling (MDS) projection of the subsampled ASD matrix for the 73 novel individuals from Central and Southern Africa, further shows substantial differentiation among groups of individuals within each one of the three clusters identified at the continental scale. Indeed, **Figure 2C**, **Figure 2D** and **Figure 2E** show substantial differentiation between Western and Eastern Central African Rainforest Hunter-Gatherer populations, and even substantial differentiation between Ba.Twa and Nsua RHG very locally in Uganda. Conversely, we find relatively much smaller genetic differentiation among pairs of RHG neighbors from West to East of the Congo Basin. Finally, results also show substantial differentiation across Southern African Khoe-San populations, albeit these populations are relatively closer from one another than the different RHG populations.

NJT and MDS based on the ASD pairwise matrix provide an important view of the major axes of genetic differentiation and variation across samples, but do not easily allow to visually describe higher-order axes of genomic variations. To do so ^18,54,56–60^, we conducted an ADMIXTURE analysis ^54^, on the same entire worldwide dataset for increasing values of K (**Figure 3**, **Figure S4**).

**Figure 3:**
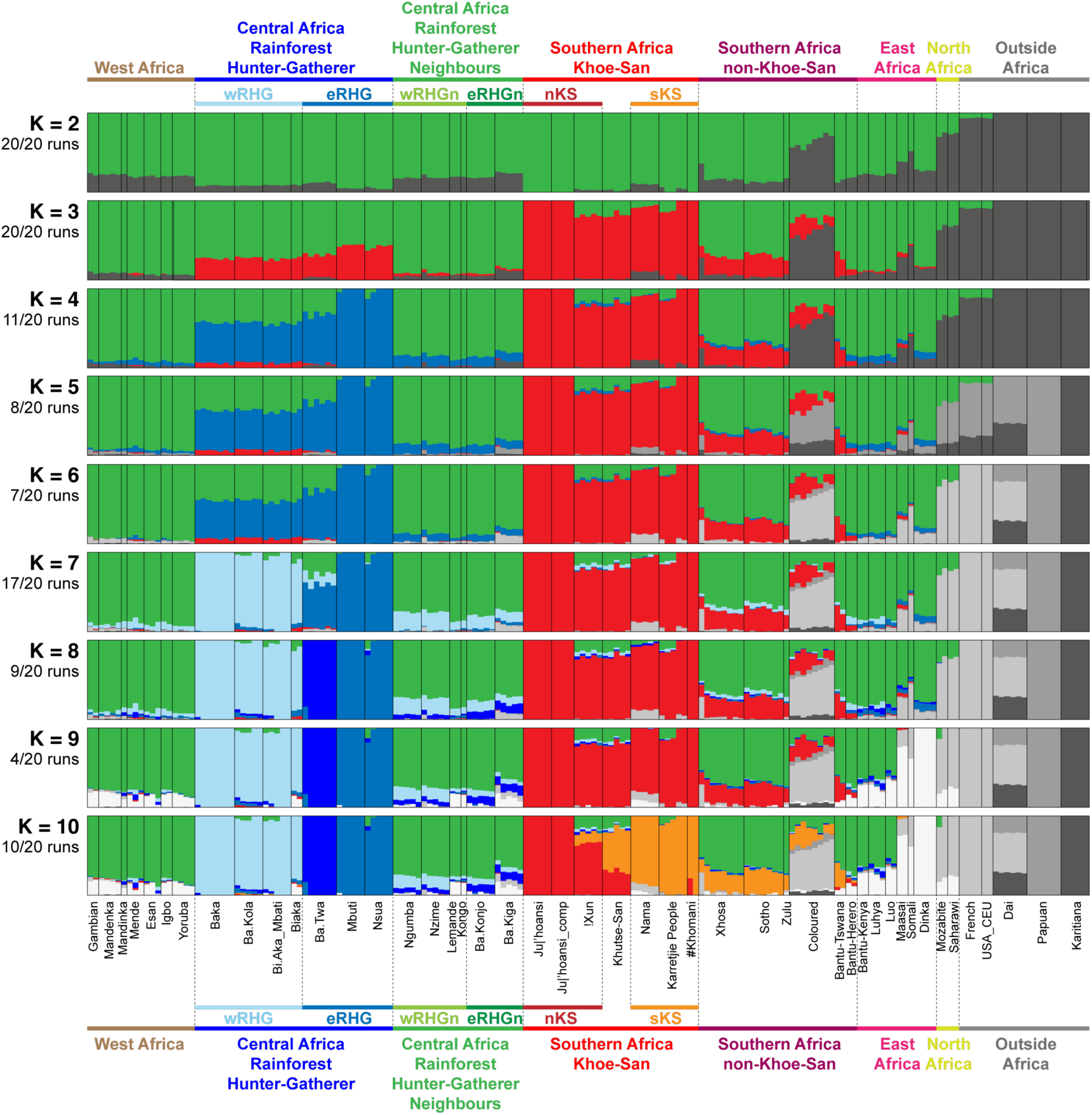
Worldwide interindividual genetic diversity patterns with ADMIXTURE. Each individual is represented by a single vertical line divided in *K* colors each proportional to an individual’s genotype membership proportion assigned by ADMIXTURE ^54^ into the virtual cluster of that color. Two individuals showing the same relative proportions of each color at a given value of *K* are genetically more resembling one another than two individuals with different relative proportions of the same colors. Population and regional geographic groupings are not considered for the calculations, individuals are *a posteriori* grouped by populations and ordered by the user. Each population is separated by a thin vertical black line. Unsupervised clustering conducted with 840,031 genome-wide SNPs pruned for LD (r^2^ threshold 0.1) and minor allele frequency (0.1) using ADMIXTURE for values of *K* ranging from 2 to 10, considering 177 worldwide individuals. For each value of *K* separately, we performed 20 independent runs and used PONG ^55^, to identify groups of resembling results (called “modes”) based on Symmetric Similarity Coefficient measures of individual results. The number of runs among the 20 independent runs belonging to the same mode result (i.e. that have pairwise SSC above 0.998), is indicated below the number of *K* on the left of each barplot separately. Only those runs are averaged per individual to provide the barplot result for each value of *K* separately. For each value of *K*, all alternative modes identified among the 20 runs are provided in **Figure S4**.

At K=2 in **Figure 3**, individual genotype memberships to the green virtual genetic cluster are maximized in individuals from Northern KS populations while those for the dark gray cluster are maximized in non-African individuals, consistent with largest genetic distances in the ASD-NJT in **Figure 2**, with all other individuals in our dataset presenting intermediate, unresolved, genotype membership proportions to either cluster.

At K=3, a novel red cluster is maximized in all Southern KS populations separating them from the rest of African individuals, also consistent with the ASD-NJT. At K=4, the novel blue cluster differentiates Central African RHG populations, and in particular certain Eastern RHG populations who maximize their membership proportions to this cluster.

The new gray and lighter gray clusters respectively at K=5 and K=6 differentiate Oceanian Papuan, South American Karitiana, and Western European French and North American USA-CEU individuals from one another and from all other non-African individuals. These two new clusters do not substantially affect genotype membership proportions in African individuals, except for the Southern African Coloured who show substantial membership to the light gray “European” cluster as well as reduced memberships to this cluster in Nama individuals and one Xhosa individual.

At K=7, the novel light blue cluster differentiates Eastern and Western RHG populations, as observed in **Figure 2**, with Ba.Twa Eastern RHG showing largely unresolved clustering at this value of K with intermediate membership proportions to the light blue, the medium blue, and the green clusters.

At K=8, the novel dark blue cluster is now maximized in Ba.Twa Eastern RHG individuals who exhibited unresolved clustering at previous values of K. Note that this novel dark blue cluster is present only to very limited extents in some other RHG and RHGn populations.

At K=9, the novel white cluster is maximized in Dinka populations, located North-East of Central Africa and the Congo Basin, and substantially represented in other, but not all, East African populations.

Finally, at K=10, the novel orange cluster differentiates mainly among Northern and Southern Khoe-San populations, with the Central Khutse San showing substantial membership proportions to both the red and the orange clusters, respectively maximized in Northern and Southern KS. Note that, at K=10, the green cluster is maximized in all non RHG and non KS Sub-Saharan Africa populations except certain East African populations and the Southern African Coloured population, albeit note that this cluster is not fully resolved at this value of K since no individual exhibits close to 100% “green” genotype membership.

In summary, these ADMIXTURE results recapitulate and further dissect the pairwise genetic differentiation visually observed with ASD-MDS and ASD-NJT in **Figure 2**. Most importantly, note that numerous Central and Southern African individuals retain, from K=2 to 10, intermediate genotype membership proportions in between certain clusters. This should be interpreted primarily as these individuals being at intermediate genetic distance between individuals presenting 100% membership to either cluster, which is consistent with ASD-MDS and ASD-NJT projections (**Figure 2**). In turn, such intermediate distances may be either due to yet-unresolved ADMIXTURE clustering, which may appear at higher values of K, or can be due to admixture having occurred between ancestors respective to each cluster, the latter interpretation being likely but not formally tested by the ADMIXTURE method. The possible occurrence of gene-flow events across pairs of lineages and their influence on tree-topologies are incorporated explicitly in our Approximate Bayesian Computation (ABC) inferences detailed below ^41–45,61,62^.

Altogether, our descriptive results highlight the vast genetic diversity and differentiation of African populations, maximized in certain groups of Central and Southern African populations, as previously reported (e.g. ^17,22,26,27,34,36,37^). Finally, and most importantly, our results highlight in particular the vast genetic diversity and differentiation of populations at a local scale in Africa, including among immediate neighbors, not trivially correlated, at continental, regional and local scales, with geographical distances among populations (**Table S4**), linguistic family classification (**Table S5**), sub-linguistic family classification (**Table S6**), nor traditional mode of subsistence (**Table S7**). In fact, these results anticipate that historical and demographic inferences in Africa may substantially differ when considering different population samples across the continent and even at a local scale, as well as advocate for caution when grouping individuals and populations samples into larger categories.

## 2. Demographic and migration histories in Central and Southern African

### 2.A. Summary of the Approximate Bayesian Computation inference design

We aimed at reconstructing the demographic and migration history that produced extant genomic patterns observed within and among Central and Southern African populations (**Figure S1**, **Figure 2**, **Figure 3**), with machine-learning ABC scenario-choices followed by posterior-parameter estimations. To do so, we considered eight competing possible tree-topologies among five extant Northern and Southern Khoe-San, Western and Eastern Rainforest Hunter-Gatherer, and Rainforest Hunter-Gatherer neighboring populations (**Figure 4**). Furthermore, for each topology, we considered two possible gene-flow processes among recent and ancient genetic lineages: instantaneous asymmetric gene-flow corresponding to instantaneous unidirectional introgression events between pairs of lineages; and recurring asymmetric gene-flow corresponding to unidirectional recurring migrations among pairs of lineages (scenarios “i” and “r” respectively, **Figure 4**). Finally, we considered, for each topology and each gene-flow process, three nested gene-flow intensities for each event among pairs of lineages separately: no to very high gene-flow rates (scenarios “i1” and “r1”); no to moderate rates (scenarios “i2” and “r2”); and no to limited rates (scenarios “i3” and “r3”). This design allowed us to evaluate fairly the classes of migration-rate scenarios that best explained the data, without having numerous a priori unrealistic groups of scenarios (e.g. very strong migration everywhere or no-migration anywhere), directly challenging more intermediate realistic scenarios.

**Figure 4:**
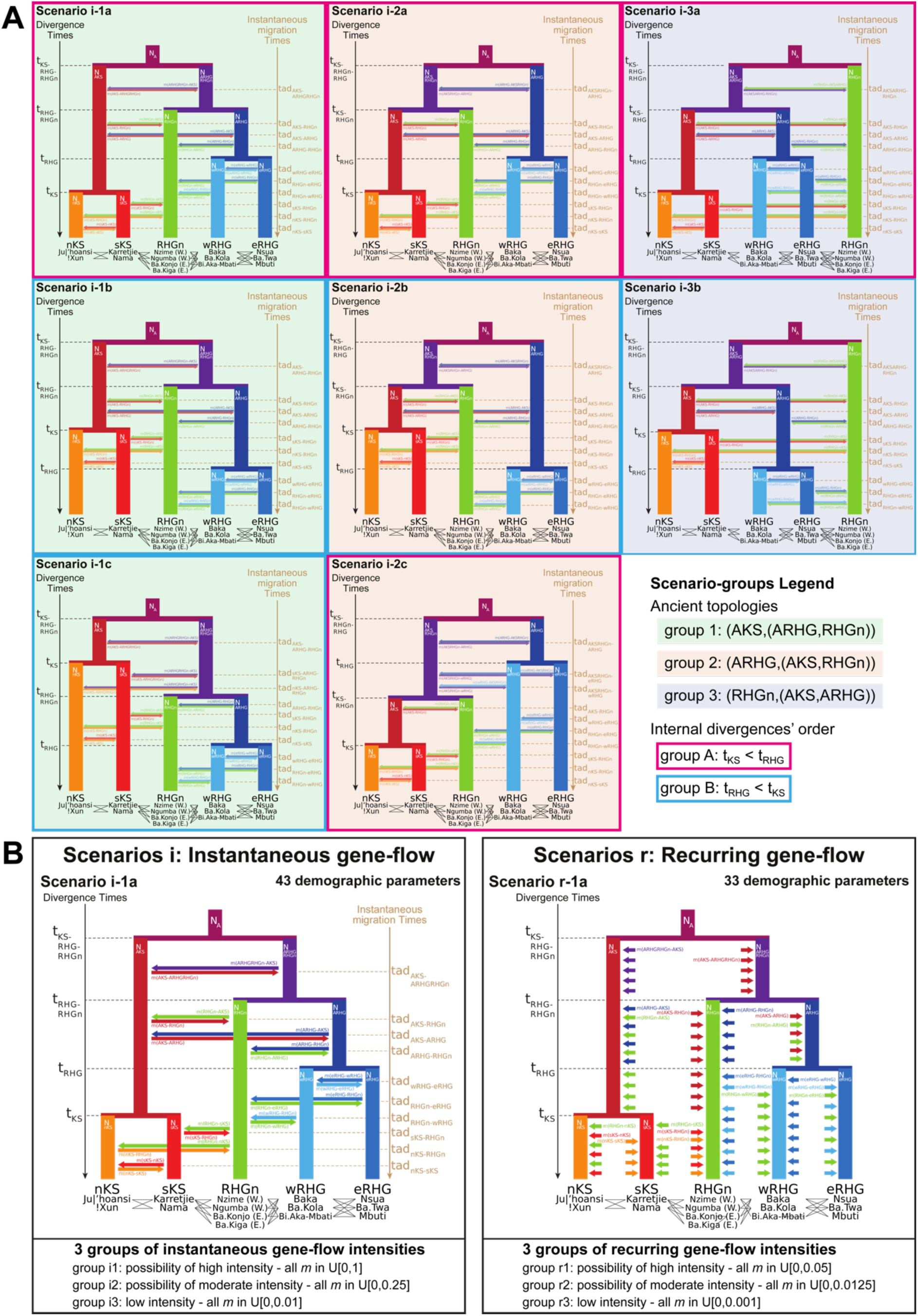
48 competing scenarios for the history of Central and Southern African populations. (**A**) Eight competing topologies for the demographic history of five Central and Southern African extant lineages. The Northern Khoe-San lineage (nKS) is represented by either the Ju|’hoansi or the !Xun population. The Southern Khoe-San lineage (sKS) is represented by either the Karretjie People or the Nama population. The Rainforest Hunter-Gatherer Neighbors lineage (RHGn) is represented either by the Western Congo Basin Nzime or Ngumba populations, or the Eastern Ba.Konjo or Ba.Kiga populations. The Western Rainforest Hunter-Gatherer lineage (wRHG) is represented by either the Baka, the Ba.Kola, or the Bi.Aka_Mbati population. The Eastern Rainforest Hunter-Gatherer lineage (eRHG) is represented by either the Nsua, the Ba.Twa, or the Mbuti population. As indicated in legend in the bottom right corner of the panel, the eight topologies can be grouped according to i) which ancestral lineage (Ancestral RHG or Ancestral KS) diverged first from the two others, or ii) whether the nKS and sKS lineages split earlier or later than the split of the wRHG and eRHG lineages. For each topology, we consider possible changes in lineages’ constant effective population size, *Ne*, after each divergence event, *t*. For each topology, we considered possible gene-flow events among ancestral lineages and among recent lineages, represented as horizontal uni-directional arrows (forward in time). (**B**) Example for a given topology (1a), of the fact that each one of the eight topologies are considered under two alternative competing “instantaneous” (Scenario i-1a) or “recurring” (Scenario r-1a) gene-flow processes. Instantaneous gene-flow processes consider that genes can be exchanged among pairs of ancestral or recent lineages at a single parameterized time, *tad*, with independent parameters of gene-flow intensity, *m*, from lineage “A” to “B” and from lineage “B” to “A”. Recurring gene-flow processes consider instead that gene-flow occur at each generation between pairs of lineages with a constant independent rate, *m*, from lineage “A” to “B” and from lineage “B” to “A”. Note that for all eight topologies and for both instantaneous and recurring gene-flow processes, we parameterized separately the gene-flow from lineage “A” into lineage “B”, and the gene-flow from lineage “B” into “A”, in order to allow for possibly asymmetrical gene-flow events among pairs of lineages. For both instantaneous and recurring gene-flow processes, we considered three nested intensities of gene-flow parameters indicated at the bottom of (**B**). Scenarios i1 or r1 consider that the gene-flow parameters *m* in each topology are independently drawn in Uniform distributions between 0 and 1 or 0 and 0.05, respectively, thus allowing for the possibility of no to high gene-flows from one lineage to the other. Scenarios i2 or r2 consider that the gene-flow parameters *m* in each topology are independently drawn in Uniform distributions between 0 and 0.25 or 0 and 0.0125, respectively, thus allowing only for the possibility of no to intermediate gene-flow from one lineage to the other. Finally, scenarios i3 or r3 consider that the gene-flow parameters *m* in each topology are independently drawn in Uniform distributions between 0 and 0.01 or 0 and 0.001, respectively, thus only allowing for the possibility of no to reduced gene-flow from one lineage to the other. Altogether we thus considered 8×2×3=48 competing scenarios for the demographic and migration history of Central and Southern African populations inferred with machine-learning Approximate Bayesian Computation. We conducted Random-Forest ABC scenario choice procedures among groups of these 48 scenarios in turn for 54 different sets of five observed populations each, represented as grey lines between population names, each population comprising five whole-genome sequenced individuals. Sampled-population names considered in the 54 combinations are given below each tree-leave, and combinations are limited to those pairing one RHGn with the specific eastern or western RHG population with whom they share complex socio-economic interactions, as indicated in **Table 1**. See the detailed description of scenarios and their grouping, their parameters and their respective prior distributions used for ABC inference in **Material and Methods** and **Table S8**.

This led to 8×2×3=48 competing scenarios in total which can be grouped in different ways to address formally two nested major questions:

1. which ancestral lineages to extant populations diverged first from the others and when did they do so, when considering complex gene-flow events among recent and ancient lineages?
2. did gene-flow events occur recurrently or more instantaneously among pairs of lineages during the evolutionary history of Central and Southern African populations? When did these events occur? How intense were they?

Importantly, we aimed at considering the vast genetic diversity and differentiation within and among populations and groups of populations at a local and regional scale (**Figure S1**, **Figure 2**, **Figure 3**, and ^3,29,30^). Therefore, we replicated the same ABC scenario-choice and posterior-parameter estimation procedures considering, in turn, 54 different combinations of five sampled populations of five sampled-individuals using large sets of autosomal genome-wide summary-statistics (**Figure 4**).

We thus conducted a total of 240,000 coalescent simulations under these 48 competing scenarios (5000 simulations per scenario), by drawing parameter values from prior distributions set by the user (**Figure 4**, **Table S8**). For each simulation, we then calculated a vector of 337 summary-statistics (**Table S9**), within and among the five simulated populations; each vector of summary-statistics thus corresponds to a vector of parameter values drawn randomly from prior distributions and used for one simulation. We then employed Random Forest ABC procedures ^45,63^, to identify the best scenario or group of scenarios for each 54 combinations of five sampled “real” populations separately. Finally, under the best scenario, we produced 100,000 simulations, computed 202 summary-statistics for each simulation (**Table S9**), and performed Neural Network ABC posterior parameter joint-estimation ^43,44^, providing posterior-parameter distributions most likely to have produced observed genomic data, for each scenario-parameter.

### 2.B. Are summary-statistics representative of large portions of the genome and are we able to mimic observed data with simulations under the 48 competing scenarios?

We first empirically checked that the observed data for each one of the 54 combinations of five sampled populations used for all ABC inferences hereafter –which considered 100 high-quality 1 Mb-long genomic windows pruned from the full genomes in our working dataset (see **Material and Methods**)– reasonably echoed the statistical descriptions obtained above from our full genomic SNP set. We thus compared standardized individual pairwise ASD matrices for each one of the 54 sets of population-combinations with the corresponding individual subsets of the standardized ASD matrix used for MDS and NJT analyses in **Figure 2**. We found that each 54 sets of population-combinations and associated pruned SNP sets for ABC inferences highly resembled the genomic individual pairwise differentiation patterns reported above using the entire SNP set. Indeed, we found significant (1000 Mantel permutation two-sided p < 0.002), Pearson *r* correlations ranging from 0.945 to 0.974 across the 54 sets of five population combinations (**Table S10**).

Furthermore, we compared entire vectors of summary-statistics computed on the 100 1Mb windows for each 54 combinations of sampled-populations separately, with the same summary-statistics computed on 724 other 1Mb such genomic regions throughout the genome. We found that the 100 windows used for ABC inferences hereafter provided summary-statistics not significantly different from- (Wilcoxon two-sided p-values between 0.6497 and 0.9995 across 54 combinations of five sampled populations), and highly correlated with- (Spearman rho between 0.8566 and 0.99637, all p-values <2.2×10^-16^), those computed on much larger portions of the genome (**Table S11**).

In order to conduct ABC inferences, we first checked whether we were able to simulate data, under the 48 competing scenarios, for which vectors of summary statistics could mimic those obtained separately from the 54 different combinations of five sampled populations each.

We first conducted goodness-of-fit permutation tests, separately for the 54 combinations of five populations, and found that the observed vectors of 337 summary statistics were never significantly different (54 goodness-of-fit permutation p-values > 0.56), from the 240,000 vectors of 337 summary statistics computed from simulations under the 48 competing scenarios. Second, we computed a two-dimensional PCA on the vectors of summary-statistics obtained from simulations on which we projected, separately, the 54 vectors of these summary-statistics computed on observed data, and found that observations each fell well within the space of simulations (**Figure S5**).

Both results show that we were able to simulate data for which summary-statistics were reasonably close to the observed ones, at least in parts of the parameter space used for simulations from the 48 scenarios and for each one of the 54 separate combinations of five sampled populations. We could therefore conclude that the observed patterns of variation could be generated under some of the 48 scenarios, and we proceeded with Random Forest ABC (RF-ABC) scenario-choice and Neural Network ABC (NN-ABC) posterior-parameter estimation procedures.

Note that the prior error in RF-ABC scenario-choice, using simulations as pseudo-observed data and considering 5000 simulations under each one of the 48 scenarios in competition, converged towards a minimum when using 1000 trees for all RF-ABC scenario-choice analyses conducted hereafter (**Figure S6**).

Importantly, the *a priori* discriminatory powers of the RF, estimated as cross-validation procedures based on all simulations used in-turn as pseudo-observed data, were overall good across all analyses described hereafter, despite increased confusion among scenarios based on gene-flow intensity within groups of instantaneous or recurring processes (**Figure S7, Figure S8, Figure S9, Figure S10**). This was expected as classes of gene-flow intensities distinguishing among scenarios with, respectively, no-to-low, no-to-moderate, or no-to-high gene-flow intensities, were designed to have nested parameter priors (**Figure 4**, **Table S8**). Note that this confusion among gene-flow intensities due to nestedness was more acute for recurring gene-flow scenarios than for instantaneous gene-flow scenarios (**Figure S9**).

Finally, we evaluated separately for each analysis how each one of the 337 summary-statistics (**Table S9**), contributed *a priori* to the RF-ABC scenario-choice in the entire space of simulations, by considering each simulation “out-of-bag” as pseudo-observed data and without considering observed data. Most interestingly, we found that individual-pairwise ASD-MDS summary-statistics describing how individuals are genetically differing from one another within and among populations and population-pairwise F_ST_, in particular, were always among the 50 most informative summary-statistics for RF-ABC scenario-choice *a priori*. Alternatively, note that we found that all statistics summarizing the 1D unfolded standardized Site Frequency Spectrum considered here were never among the 50 most informative statistics *a priori* using the Gini index of classification, for any of the RF-ABC scenario-choice analyses (**Figure S11**, **Figure S12**, **Figure S13**, **Figure S14**, **Figure S15**, **Figure S16**, **Figure S17**, **Figure S18**, **Figure S19**, **Figure S20**).

### 2.C. Which historic scenario best explains genomic patterns in extant African populations?

#### 2.C.1. Instantaneous or recurring gene-flows of which intensity among African lineages?

**Figure 5A** shows that gene-flow processes among recent or ancient genetic lineages most likely occurred during very limited “instantaneous” periods of time, rather than recurrently, throughout the entire evolutionary history of Central and Southern African populations. Indeed, for each one of the 54 different combinations of five sampled populations, the group of 24 scenarios considering instantaneous gene-flow processes provides vectors of summary statistics systematically closest to the observed ones, whichever the tree-topology or intensity of the gene-flow considered. Note that we find that posterior probabilities of choosing the correct group of scenarios for each 54 sets of population combinations range from 0.97828 to 0.99510, compared to a prior probability of 1/2 in this particular test (**Table S12**).

**Figure 5:**
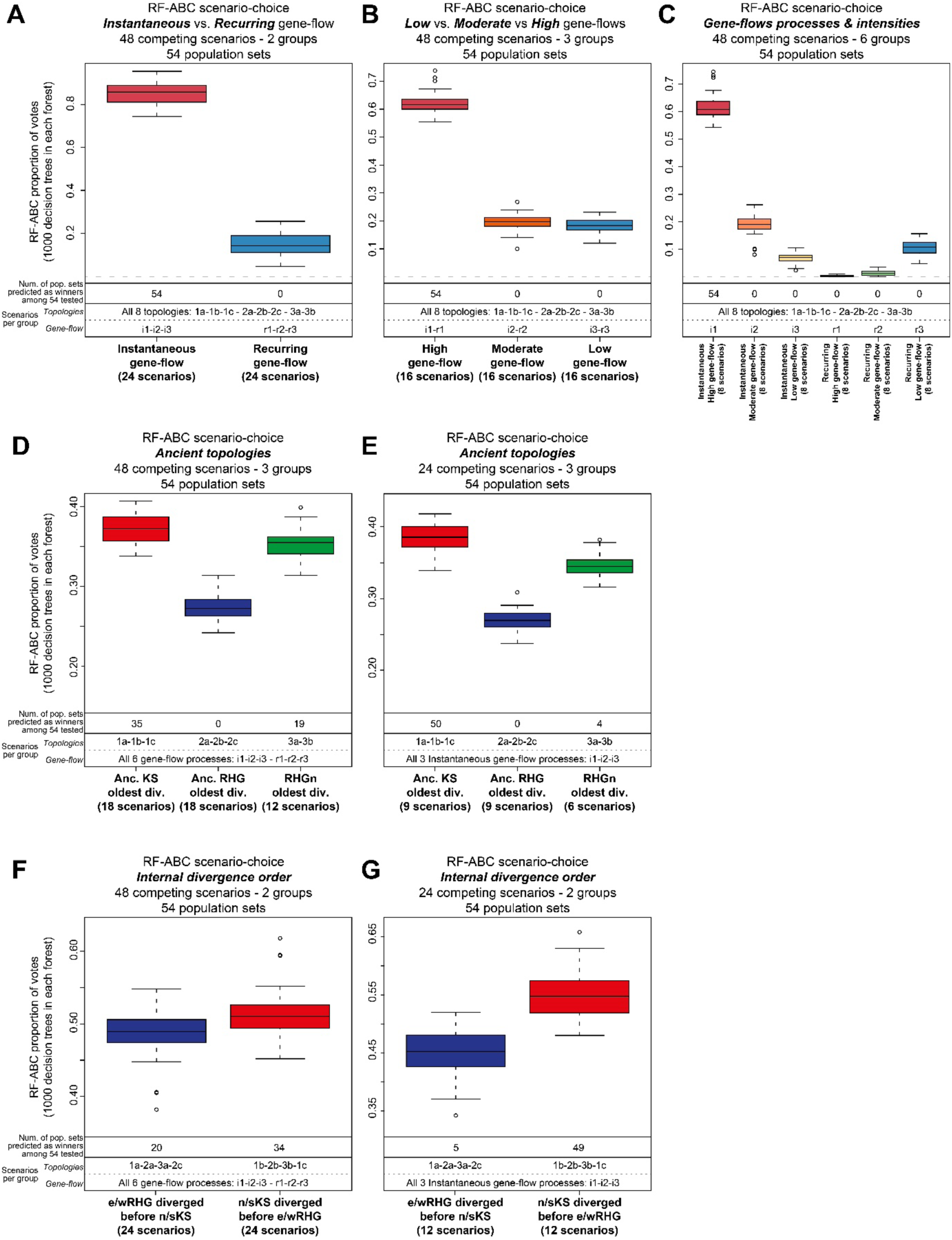
Random-Forest ABC scenario-choices among 48 competing scenarios. Random Forest Approximate Bayesian Computation scenario-choice results ^45,63^, are conducted, for each analysis in each panel, separately for 54 different combinations of five Central and Southern African sampled populations each. Posterior proportions of votes obtained for each 54 sampled-populations combinations, and for RF-ABC analysis in each panel respectively, are provided as box-plots indicating the median in between the first and third quartile of the box-limits, whiskers extending to data points no more than 1.5 times the interquartile range of the distribution, and empty circles for all more extreme points beyond this limit, if any. In each RF-ABC analysis in each panel, proportion of votes (indicated in the y-axis) are obtained using 1,000 decision trees in the random forest for each group of competing scenarios (indicated in the x-axis table and labels), using 5,000 simulations per scenario and 337 summary-statistics (see **Material and Methods** and **Table S9** for details). Detailed competing-scenarios grouped in each group are indicated in the two bottom lines of the x-axis table, with topology and gene-flow processes codes provided in **Figure 4** and explained in detail in **Material and Methods**. The top-line of the x-axis table indicates, for each group of competing scenarios in each analysis, the number of times the corresponding group of scenarios was predicted as the best one among the 54 different combinations of five sampled-populations. All figure results can be found in table format in **Table S12**. (**A**) Scenario-choice results for the two groups of instantaneous or recurring gene-flow processes (24 scenarios and 120,000 simulations in each two competing groups), all gene-flow intensities and all topologies “being equal”. Posterior probabilities of choosing the correct group of scenarios ^64^, for each 54 sets of population combinations ranged from 0.97828 to 0.99510, compared to a prior probability of 1/2 in this particular test. (**B**) Scenario-choice results for the three groups of intensities for gene-flow processes (16 scenarios and 80,000 simulations in each three competing groups), all topologies and both instantaneous and recurring gene-flow processes “being equal”. Posterior probabilities of choosing the correct group of scenarios for each 54 sets of population combinations ranged from 0.66318 to 0.87138, compared to a prior probability of 1/3 in this particular test. (**C**) Scenario-choice results for the six groups of gene-flow processes and intensities separately (8 scenarios and 40,000 simulations in each six competing groups), all topologies “being equal”. Posterior probabilities of choosing the correct group of scenarios for each 54 sets of population combinations ranged from 0.67765 to 0.83957, compared to a prior probability of 1/6 in this particular test. (**D**) Scenario-choice results for the three groups of scenario topologies differing in the order of ancient lineages divergence (differing number of scenarios in each three competing groups are evened by randomly sampling the same number of simulations, equal to 60,000, in each group, see **Material and Methods**), all gene-flow processes and intensities “being equal”. Posterior probabilities of choosing the correct group of scenarios for each 54 sets of population combinations ranged from 0.39862 to 0.48967, compared to a prior probability of 1/3 in this particular test. (**E**) Scenario-choice results for the same test as (**D**) restricted to the 24 competing scenarios considering instantaneous gene-flow processes only (differing number of scenarios in each three competing groups are evened by randomly sampling the same number of simulations, equal to 30,000, in each group, see **Material and Methods**), all gene-flow intensities “being equal”. Posterior probabilities of choosing the correct group of scenarios for each 54 sets of population combinations ranged from 0.43333 to 0.51857, compared to a prior probability of 1/3 in this particular test. (**F**) Scenario-choice results for the two groups of scenario topologies differing in the relative order of divergences between Northern and Southern KS, and Western and Eastern RHG lineages, respectively (24 competing scenarios and 120,000 simulations in each two groups), all gene-flow processes and intensities “being equal”. Posterior probabilities of choosing the correct group of scenarios for each 54 sets of population combinations ranged from 0.62115 to 0.72015, compared to a prior probability of 1/2 in this particular test. (**G**) Scenario-choice results for the same test as (**F**) restricted to the 24 competing scenarios considering instantaneous gene-flow processes only (12 competing scenarios and 60,000 simulations in each two groups), all gene-flow intensities “being equal”. Posterior probabilities of choosing the correct group of scenarios for each 54 sets of population combinations ranged from 0.62033 to 0.71883, compared to a prior probability of 1/2 in this particular test.

Furthermore, we find that we imperatively need these gene-flow processes to be potentially high in order to best explain the observed data for each one of the 54 combinations of population samples, whichever the tree-topology and gene-flow process (**Figure 5B**); whichever the tree-topology but considering gene-flow processes separately (**Figure 5C**); or even considering 24 competing scenarios under instantaneous gene-flow processes only, whichever the tree-topology (**Figure S8**). Note that, for these three particular tests, we find that posterior probabilities of choosing the correct group of scenarios ^64^, for each 54 sets of population combinations range, respectively, from 0.66318 to 0.87138 (**Figure 5B, Table S12**), from 0.67765 to 0.83957 (**Figure 5C, Table S12**), and from 0.63935 to 0.87000 (**Figure S8**), compared to a prior probability of 1/3, 1/6, and 1/3 in these three particular tests, respectively. Finally, note that when considering the 48 competing-scenarios separately, a very challenging task *a priori* given the number of competing scenarios and the high level of nestedness among scenarios, the best scenarios predicted by the RF-ABC procedure is, among the 54 population-combination tests conducted, systematically found for scenarios considering an instantaneous gene-flow process allowing for possibly highly intense gene-flow rates **(Figure** S9, **Figure** S10, TableS12).

#### 2.C.2. Ancient tree-topology among African lineages?

**Figure 5D** shows that, overall, the RF-ABC scenario-choice procedures vote in favor of an ancient tree-topology where the ancestral Khoe-San lineage diverged first followed by the divergence between the ancient Rainforest Hunter-Gatherer lineage and that of their neighbors (Scenarios tree-topologies 1a-1b-1c, **Figure 4**), in a majority (35/54) of the 54 combinations of observed populations tested here, whichever the gene-flow process and intensity. Note that we find that posterior probabilities of choosing the correct group of scenarios for each 54 sets of population combinations range from 0.39862 to 0.48967, compared to a prior probability of 1/3 in this particular test (**Figure 5, Table S12**).

Furthermore, we find that this majority of votes is much larger (50/54), when considering instantaneous gene-flow processes only, whichever their intensities (**Figure 5E**). Note that we find that posterior probabilities of choosing the correct group of scenarios for each 54 sets of population combinations range from 0.43333 to 0.51857, compared to a prior probability of 1/3 in this particular test (**Figure 5, Table S12**).

Finally, note also that the tree-topologies where the RHGn lineage diverge first from the lineage ancestral to that of the RHG and the KS (Scenarios tree-topologies 3a-3b, **Figure 4**), are predicted to be the best ones in a minority of the tests (19 out of 54 combinations of sampled populations, **Figure 5D**), whichever the gene-flow process and/or intensity. Furthermore, they do so in an even smaller minority (4/54) for instantaneous gene-flow processes only, whichever the intensities (**Figure 5E**). Interestingly, we found that tree-topologies where the ancient lineage for RHG populations diverges first from the two others (Scenarios tree-topologies 2a-2b-2c, **Figure 4**), is never favored in our analyses, whichever the gene-flow process and/or intensity.

#### 2.C.3. Recent tree-topology among African lineages?

Finally, **Figure 5** shows that Northern and Southern Khoe-San extant lineages likely diverged from one-another before the divergence between Eastern and Western Rainforest Hunter-Gatherer lineages (Scenarios tree-topologies 1b-1c-2b-3b, **Figure 4**), whichever the ancient tree-topology and gene-flow process and intensity (**Figure 5F**), and whichever the ancient tree-topology for instantaneous gene-flow processes only, all classes of gene-flow intensities “being equal” (**Figure 5G**). Interestingly, the “b” group of tree-topologies is found to best-describe observed summary-statistics for 34 out of 54 combinations of sampled populations and this majority of RF-ABC votes is even larger (49/54) when considering only the group of 24 competing scenarios for which gene-flows are instantaneous rather than recurring. Note that, for these two particular tests, we find that posterior probabilities of choosing the correct group of scenarios for each 54 sets of population combinations range, respectively, from 0.62115 to 0.72015 (**Figure 5F, Table S12**), and from 0.62033 to 0.71883 (**Figure 5G, Table S12**), compared to a prior probability of 1/2 in both these two particular tests.

#### 2.C.4. Conclusion: intersecting scenario-choice results for the history of Africa

Altogether, when intersecting RF-ABC scenario-choice inferences in groups of gene-flow processes and tree-topologies (**Figure 5**), with inferences considering all competing-scenarios separately (**Figure S9, Figure S10**), and among 54 combinations of five sampled populations, we find that Scenario i1-1b (**Figure 4**), systematically best-describes the observed data in all conformations of scenario-choice tests for 25 combinations of populations (**Table S12, Table S13**), while the second scenario best explaining the data (Scenario i1-3b) does so consistently across all tests for only four combinations of population out of 54.

Therefore, our results point to a scenario for the history of Central and Southern African populations where the ancestral KS lineage diverged first, followed by the divergence between the ancestral RHG and the RHGn lineages, with a divergence between the extant Northern and Southern KS lineages occurring independently before that of the extant Eastern and Western RHG lineages. Most importantly, the best scenario necessarily involves gene-flow events among all pairs of lineages throughout history which occurred during relatively short “instantaneous” periods of time, rather than recurrently over longer periods of time. Furthermore, these instantaneous gene-flow events must each have been possibly of high intensity, rather than all limited to moderate or low intensities, and possibly asymmetrical between pairs of lineages, to best explain the observed data. Finally, our results also highlight that considering different combinations of extant populations separately in the analyses may provide contrasted results where alternative evolutionary scenarios may best explain observations.

For conservativeness and to further consider the genetic diversity of extant Central and Southern African populations at a local or regional scale, we will henceforth conduct all Neural Network ABC posterior-parameter estimations separately for the 25 combinations of sampled populations systematically providing the Scenario i1-1b as the most-likely one among the 48 competing scenarios. **Table S13** shows no specific statistical or anthropological pattern among the specific sampled-populations in the 25 combinations providing consistent results across RF-ABC analyses. Indeed, the least represented sampled-population, the Ngumba RHGn, appears in 20% of the combinations (four RHGn sampled populations are considered in turn); the most represented sampled-population, the Ju|’hoansi nKS, appears in 76% of the combinations (two nKS sampled populations are considered in turn); and all 14 sampled-populations considered in each RF-ABC scenario-choice analysis are present in the 25 combinations.

### 2.D. Which scenario-parameters best explain genomic patterns in extant African populations?

#### 2.D.1. Divergence times

Divergence times are most often well estimated by our Neural Network ABC posterior parameter inferences (**Figure 6** and **Table S14**), with relatively narrow 90% Credibility Intervals (CI), posterior distributions very often largely departing from the priors, and low cross-validation posterior parameter estimation errors in the vicinity of the observed data (**Table S16**). Interestingly, we find relatively consistent posterior estimates across the 25 different combinations of five populations for, separately, the Eastern and Western RHG divergence (*t*_RHG_), between 355 generations before present (gbp) (mode point estimate, 90%CI=[203-1,220]) and 1,518 gbp (90%CI=[805-3,016], **Table S14**). We find divergences between the Northern and Southern KS (*t*_KS_), between 2.1 and 8.0 times older than the corresponding *t*_RHG_ for the 25 combinations of sampled-populations. These correspond in absolute to *t*_KS_ between 1,882 gbp (mode point estimate, 90%CI=[1,114-3,270]) and 3,823 gbp (90%CI=[2,336-5,669], **Table S14**). Combining (concatenating) the posterior distributions for these two parameters (**Figure 6A**, **Figure 6B**, **Table S17**), respectively, provides, synthetically, an absolute modal point-estimate divergence time between eRHG and wRHG of 751 gbp (90%CI =[370-2,222]), and one between nKS and sKS, largely more ancient, having occurred some 2,857 gbp (90%CI=[1,624-4,662]).

**Figure 6:**
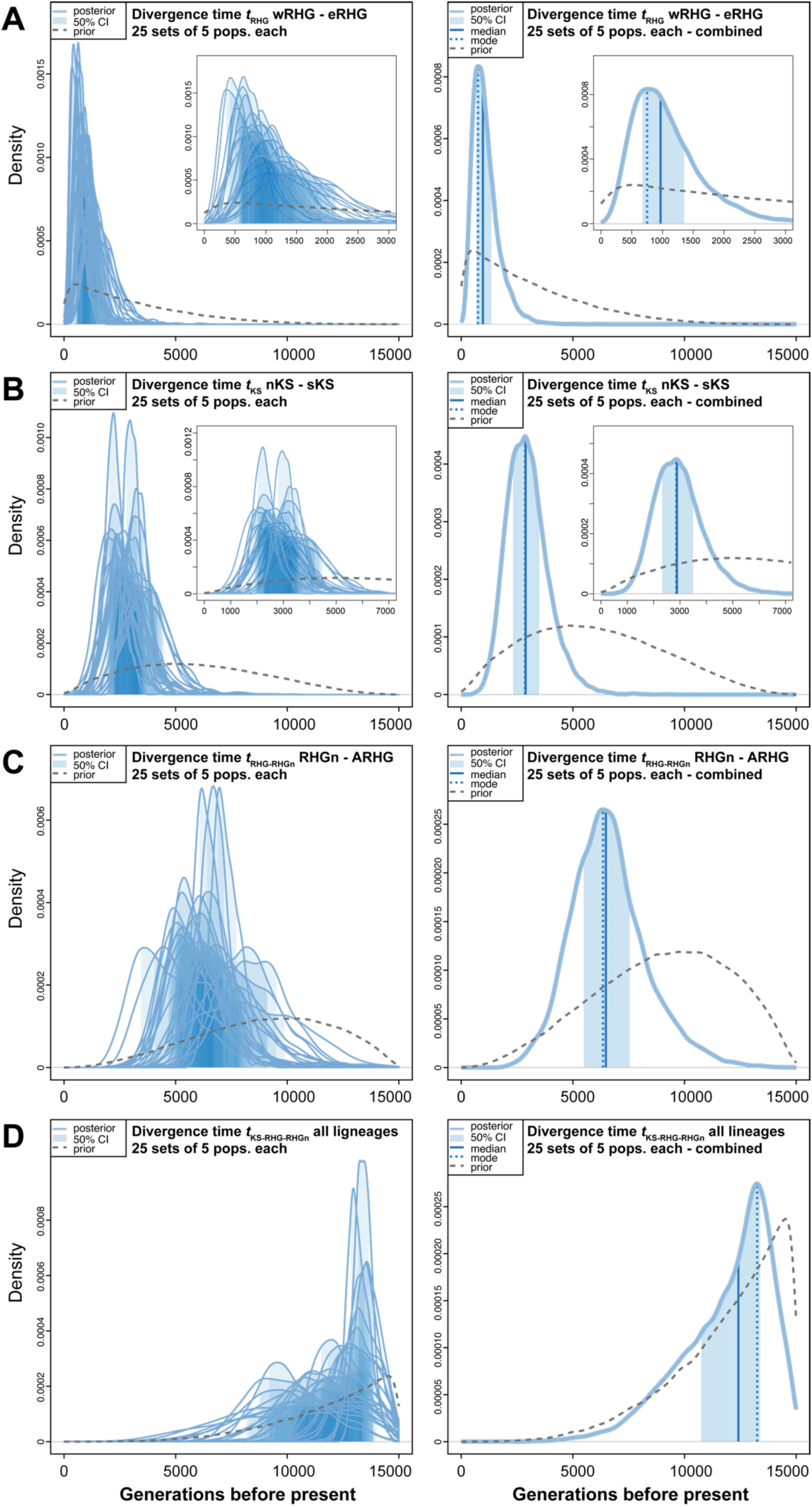
Neural Network ABC posterior parameter distributions of divergence times. Neural Network Approximate Bayesian Computation posterior parameter joint estimations ^43,44^, of topological divergence-times (in generations before present) for 25 sets of five Central and Southern African populations for which the best scenario identified by RF-ABC was Scenario i1-1b (**Figure 4**). NN ABC posterior parameter estimation procedures were conducted using 100,000 simulations under Scenario i1-1b, each simulation corresponding to a single vector of parameter values drawn randomly from prior distributions provided in **Table S8**. We considered 43 neurons in the hidden layer of the NN and a tolerance level of 0.01, corresponding to the 1,000 simulations providing summary-statistics closest to the observed one. NN posterior estimates are based on the logit transformation of parameter values using an Epanechnikov kernel between the corresponding parameter’s prior bounds (see **Material and Methods** and **Table S8**). Posterior parameter densities are represented with solid blue lines. 50% Credibility Intervals are represented as the light blue area under the density. The median and mode values are represented as a solid and dotted blue vertical line, respectively. Parameter prior distributions are represented as dotted gray lines. For all panels, the left plots represent the NN-ABC posterior parameter distributions for each 25 sets of five Central and Southern African populations best explained under Scenario i1-1b, separately (**Table S13** and **Table S14**). For all panels, the right plots represent a single parameter posterior distribution obtained from combining (concatenating) the 25 posterior distributions together. (**A**) Results for parameter *t*_RHG_ corresponding to the split time between the Western and Eastern Rainforest Hunter-Gatherer (RHG) lineages (**Figure 4**). (**B**) Results for parameter *t*_KS_ corresponding to the split time between Northern Khoe-San (nKS) and Southern Khoe-San (sKS) lineages (**Figure 4**). (**C**) Results for parameter *t*_RHG-RHGn_ corresponding to the split time between the Rainforest Hunter-Gatherer neighboring population lineage (RHGn) and the lineage ancestral to Western and Eastern RHG (**Figure 4**). (**D**) Results for parameter *t*_KS-RHG-RHGn_ corresponding to the split time between the lineage ancestral to KS populations and the lineage ancestral to all RHG and RHGn lineages; this event thus corresponding to the split time of all lineages in the history of Central and Southern African populations (**Figure 4**). Results for all left panels are summarized in **Table S17** and cross-validation posterior parameter estimation errors in the vicinity of the observed data in **Table S16**.

In a more remote past, the divergence time between the lineage ancestral to all RHG populations and the RHG neighbors’ lineage is also most often well estimated, based on CI-width and departure from prior distributions, and cross-validation posterior parameter estimation errors in the vicinity of the observed data, for each 25 combinations of populations respectively, albeit results are more variable across sets of population combinations than for more recent divergence times (**Figure 6C**, **Table S14, Table S16**). The ancient RHG and RHGn lineages’ divergences (*t*_RHG-RHGn_) are between 4.5 and 22.3 times older than the corresponding t_RHG_ for the 25 combinations of sampled-populations. This translates in absolute between 3,595 gbp (mode point estimate, 90%CI=[2,571-7,368]) and 8,199 gbp (90%CI =[6,083-10,682], **Table S14**). Synthetically combining results together provides an absolute divergence time between ancestors of extant RHGn and extant RHG of 6,347 gbp (90%CI=[4,098-9,734], **Figure 6C** and **Table S17**).

Finally, we also find relatively variable posterior estimates for the most ancient divergence time in our tree-topology, between the ancestral lineages to all extant KS lineages and the ancestral lineages to all RHG and the RHG neighbors; with, again, posterior-parameter distributions across 25 combinations of sampled populations very often satisfactorily estimated, with relatively reduced 90%CI and substantial departure from the priors, and relatively low cross-validation posterior parameter estimation errors in the vicinity of the observed data (**Figure 6D**, **Table S14, Table S16**). We find the original most ancient divergence (*t*_KS-RHG-RHGn_) in our tree-topology between 8.6 and 33.6 times older than the corresponding most recent divergence *t*_RHG_ for the 25 combinations of sampled-populations. This translates in absolute to a modal ancestral divergence among all investigated Central and Southern African lineages that occurred some 13,254 gbp (90%CI=[8,044-14,397]), when all results are combined (concatenated) together synthetically (**Figure 6D** and **Table S17**).

#### 2.D.2. Effective population sizes

Effective population sizes (Ne) for all recent Northern and Southern KS, Western and Eastern RHG, and RHG neighbors’ lineages are often reasonably well estimated with relatively reduced 90%CI, substantial departure from their priors, and low cross-validation posterior parameter estimation errors in the vicinity of the observed data (**Table S16**), for each 25 sets of sampled population combinations (**Figure S21**, **Table S14**). Combining posterior distributions among the 25 separate tests, we find Ne posterior estimates ranging from modal point-estimates of 10,702 diploid effective individuals (90%CI=[5,769-67,086]) in the eRHG to 17,083 (90%CI=[7,606-71,133]) in the wRHG extant lineages (**Figure S21**, **Table S17**). Importantly, we were unable to satisfactorily estimate, for almost all 25 combinations of sampled populations, the three effective population sizes for, respectively, the ancient KS lineage, the ancient RHG lineage, and the lineage ancestral to all RHG and RHG neighbors’ extant lineages (**Table S17**, **Figure S21**, **Table S14, Table S16**).

Conversely, our posterior estimates of the effective size of the lineage most ancestral to all our populations was satisfactorily estimated, with relatively narrow 90%CI, substantial departure from the prior distributions, and among the lowest cross-validation posterior parameter estimation errors in the vicinity of the observed data (**Table S16**), for almost all 25 sets of population combinations. Concatenating posterior distributions, despite a noteworthy variation across the 25 tests (**Figure S21**, **Table S14**), we find an ancestral effective size for the lineage ancestral to all our Central and Southern African extant populations of 17,005 diploid effective individuals (90%CI=[9,890-69,775]), the second largest posterior estimate across all recent and ancient lineages in our analyses (**Table S17**). Note that we considered large priors (Uniform[10-100,000] diploid individuals) for constant effective population sizes in all lineages with possible changes at each divergence time, for simplicity. Therefore, the posterior estimates here found may be different in future procedures considering more complex effective demographic regimes likely to have occurred in certain lineages, such as possible bottlenecks and/or population expansions (e.g. ^8,40,65^).

#### 2.D.3. Instantaneous gene-flow times

In between each lineage divergence times, we estimated the time of occurrence of instantaneous and potentially asymmetric gene-flow exchanges across pairs of lineages (**Figure 4**), for each 25 combinations of sampled populations separately. For four out of the six recent gene-flow times between pairs of extant lineages (tad_nKS-sKS_, tad_wRHG-eRHG_, tad_wRHG-RHGn_, tad_eRHG-RHGn_, **Figure 4**, **Table S8**), we find relatively consistent posterior estimates across the 25 tests, almost all with relatively reduced 90%CI, substantial departure from the priors, and low cross-validation posterior parameter estimation errors in the vicinity of the observed data (**Figure 7A – Figure 7D**, **Table S14, Table S16**, **Table S17**). Note that, while we also obtain similarly satisfactory posterior distributions among the 25 tests for gene-flows among Northern KS and RHGn lineages (tad_nKS-RHGn_), and among Southern KS and RHGn lineages (tad_sKS-RHGn_), respectively, we found much larger variance across the 25 different population sets (**Figure 7E – Figure 7F**, **Table S14**).

**Figure 7:**
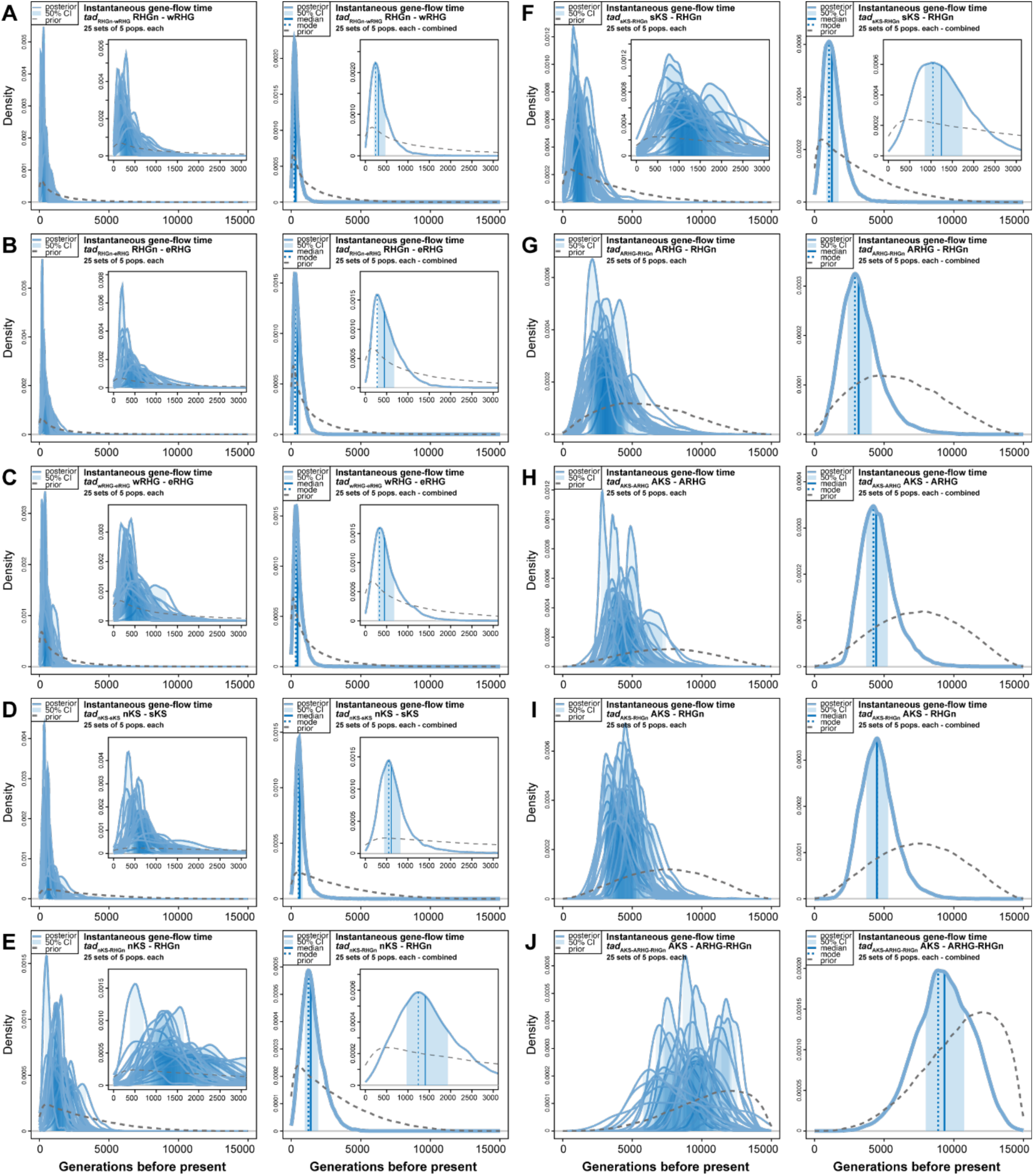
ABC posterior parameter distribution of gene-flow times. Neural Network Approximate Bayesian Computation ^43,44^, posterior parameter joint estimations of gene-flow instantaneous times *tad* (in generations before present) for 25 sets of five Central and Southern African populations for which the best scenario identified by RF-ABC was Scenario i1-1b (Figure 4, **Table S13**). Methodological details are provided in Figure 6 caption and detailed in **Material and Methods**.

Interestingly, we also obtain satisfactory posterior estimates of gene-flow timing among ancient lineages for almost all 25 sets of population combinations (tad_ARHG-RHGn_, tad_AKS-RHGn_, tad_AKS-ARHG_, tad_AKS-ARHG-RHGn_, **Figure 4**, **Table S8**). Nevertheless, note that posterior estimates are increasingly variable from one set of sampled populations to the other as the estimated parameter is further back in time in the tree-topology (**Figure 7G – Figure 7J**, **Table S17**).

#### 2.D.4. Gene-flow intensities

We obtain overall unsatisfactory posterior estimates of the 20 separate unidirectional gene-flow rates for the 10 separate instantaneous gene-flow events, for a majority of the 25 sets of population combinations (**Figure S22, Table S14**), with relatively large 90%CI, reduced posterior distributions’ departure from priors, and high cross-validation posterior parameter estimation errors in the vicinity of the observed data (**Table S16**). Considering only the several satisfactory posterior estimates for each gene-flow parameter separately, we notice their very large variability across the sets of population combinations for each parameter, making it unreasonable to combine posterior distributions to obtain a synthetic value (**Table S17**). The strong parameter-estimation limitation encountered here is extensively discussed in the **Discussion** section below.

#### 2.D.5. A synthesis of the demographic history of Central and Southern African populations

We propose, in **Figure 8**, a schematic synthesis of all the above inference results for the demographic history of 25 out of 54 combinations of five Central and Southern African populations, for which all our RF-ABC scenario-choice procedures provided systematically consistent results. **Figure 8** synthesizes mode estimates of all reliably inferred parameters obtained from the combined (concatenated) posterior distributions for the 25 combinations of sampled populations provided in **Table S17** and presented in **Figure 6**, **Figure 7**, **Figure S21**, and **Figure S22**. Analogous schematic representations for each 25 combinations of sampled populations separately corresponding to results in **Table S14** are presented *in extenso* in **Figure S23** to **Figure S47.**

**Figure 8:**
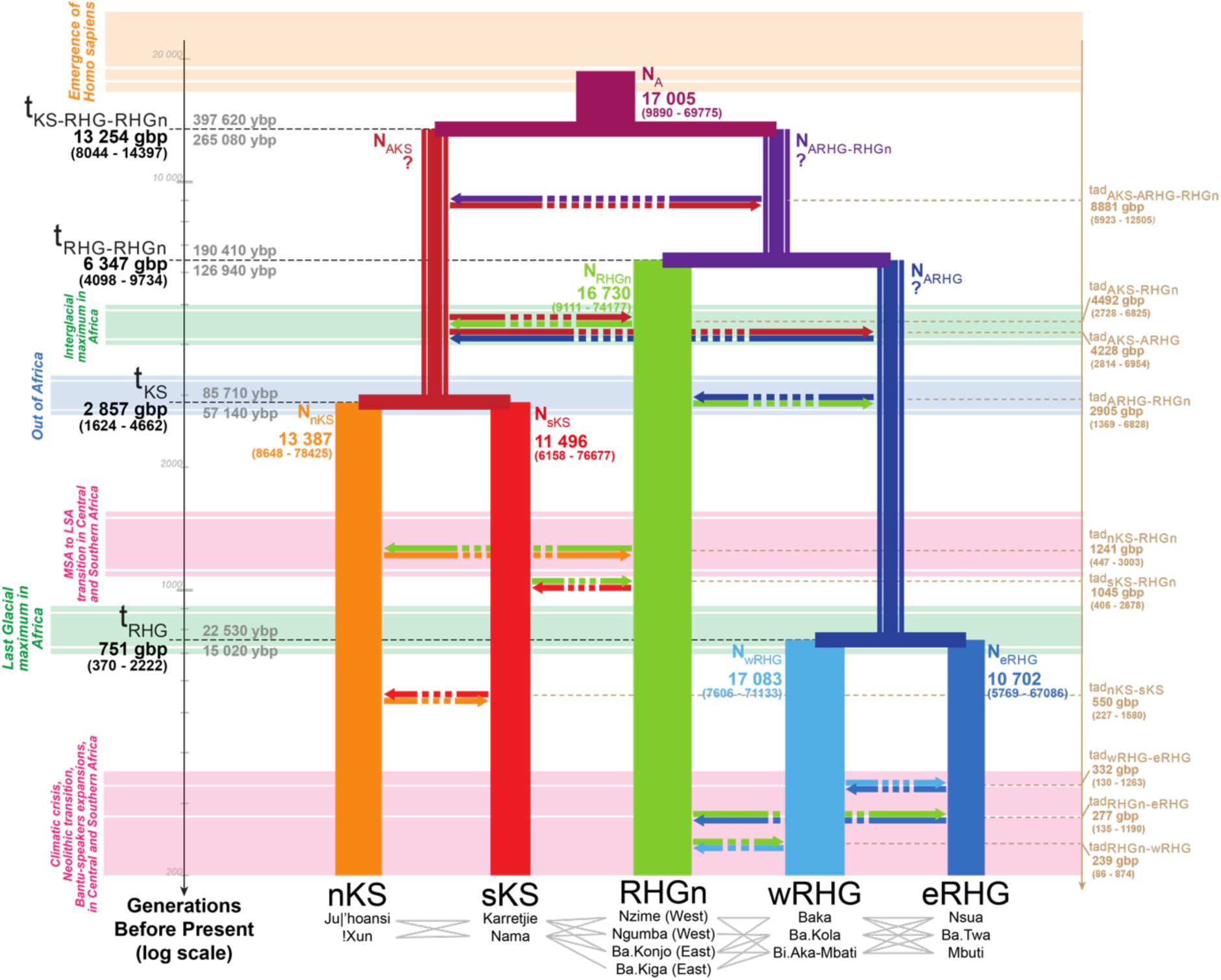
Schematic inferred demographic and migration history of Central and Southern African populations. Schematic representation of Scenario i1-1b (Figure 4), and Neural Network ABC posterior parameter mode estimates summarizing results obtained separately for 25 sets of five Central and Southern African populations (**Table S12**, **Table S13, Figure S23 to Figure S47**), represented by the gray lines in between population names. Gene-flow arrows are indicated forward in time. For the time of each divergence and gene-flow event, mode point estimates are provided in generations before present (gbp) in bold, and 90% Credibility Intervals are provided between parentheses (**Table S17**). We provide two estimates of the divergence times estimates in years before present (ybp), one (upper) corresponding to 30 years per generation and the other (lower) to 20 years per generation ^66,67^. Mode point estimates of effective population sizes *Ne* are provided in numbers of diploid effective individuals and width of lineages are proportional to the estimated *Ne* (**Table S17**, **Figure S21**). NN-ABC posterior distributions for the effective population sizes of the ancestral Khoe-San lineage (AKS), for the ancestral RHG lineage (ARHG), and for the lineage ancestral to RHGn and ARHG, were all three poorly distinguished from their respective prior distributions and with high cross-validation posterior parameter estimation errors (**Figure S21**, **Table S16, Table S17**), as indicated by the question marks. NN-ABC posterior distributions for instantaneous asymmetric gene-flow rates were overall poorly departing from their priors and with high cross-validation posterior parameter estimation errors (**Figure S22**, **Table S14, Table S16**), as indicated by the doted arrows. All posterior distributions are shown graphically in Figure 6, Figure 7**, Figure S21, Figure S22**, and detailed in **Table S14** and **Table S17**, with cross-validation posterior errors for each parameter in **Table S16**. An analogous synthetic schematic for the original results obtained for each one of the 25 combinations of five sampled populations is presented separately in **Figure S23** to **Figure S47**.

Posterior checks show that 25 separate pseudo-observed simulations computed under the mode point-estimates for this best synthetic demographic scenario, as well as 25 separate pseudo-observed simulations each computed under the best demographic scenario obtained respectively for each 25 observed combination of populations (**Table S16**), resembles those from the 25 observed combinations of five sampled populations in the first two axes of a PCA computed on the 202 summary-statistics used for all NN-ABC inferences (**Figure S48, Figure S49**). Furthermore, **Figure S48** shows that, as expected, more variance in summary-statistics is observed among the 25 observed combinations of sampled populations than among the 25 pseudo-observed simulations under the proposed synthetic model. This shows that, while the synthetic demographic history proposed here results from artificially combining the diversity of demographic histories inferred separately for each combination of sampled-populations, this synthesis provides reasonably similar genomic patterns as those observed in real data; keeping nevertheless in mind that the relatively large Credibility Intervals reported under this best synthetic scenario also reflect the relatively large variation of histories across specific combinations of sampled populations under this scenario (**Table S14** and **Figure S23** to **Figure S47**).

We conducted NN-ABC posterior-parameter estimations as for the observed datasets on the 25 pseudo-observed simulations. We find substantially more variance in combined posterior-parameter estimates for most parameters that were reasonably accurately estimated across the 25 combinations of five Central and Southern African sampled-populations, compared to the variance of combined (concatenated) posterior-parameter distributions obtained for 25 pseudo-observed simulations under the proposed synthetic demographic scenario (**Figure S50**, **Figure S51**, **Figure S52**, **Figure S53**, and **Table S18**). Indeed, **Table S18** shows that 15 out of the 20 demographic parameters posterior combined distributions had a significantly larger variance with observed datasets than with pseudo-observed simulations; and an additional two parameters exhibit the same pattern but only marginally significant.

In particular (**Table S18**), the divergence times obtained with observed data are between 22% (t_RHG_; Fligner test p-value=5.46×10^-146^) and 154% (t_KS-RHG-RHGn_; Fligner test p-value=0), more variable than those obtained with pseudo-observed simulations. Furthermore, the admixture times obtained with observed data are between 16% (tad_AKS-RHGn_; Fligner test p-value=4.67×10^-25^) and 211% (tad_eRHG-RHGn_; Fligner test p-value=0), more variable than those obtained with pseudo-observed simulations, excluding the two parameters with the marginally significant patterns.

Altogether, these results demonstrate empirically that the large variation in divergence and instantaneous gene-flow time posterior estimates obtained across combinations of sampled populations largely depend on the specific samples considered when investigating highly differentiated Sub-Saharan populations, rather than on variation in inference performances. Our results thus explicitly advocate for caution when summarizing results obtained across population sets and further likely explain the, sometimes, apparent discrepancies that arise across studies considering different sampling designs prior to conducting demographic evolutionary reconstructions (e.g. ^22,26^).

## Discussion

We aimed at inferring jointly the tree-topology, instantaneous or recurring gene-flow processes, and asymmetric gene-flow intensities across pairs of recent and ancient lineages that most likely produced genome-wide genetic patterns in 54 different combinations of five Central and Southern African populations. Therefore, we explicitly considered the large genetic differentiation observed across African populations at regional and local scales, and the possible confounding effects of complex gene-flow processes on tree-topologies’ predictions among 48 complex scenarios in formal statistical competition.

First, our results altogether show that, in fact, considering different sets of populations at a regional and local scale in Africa highlights different aspects of the demographic histories of African populations, sometimes substantially divergent. This demonstrates that apparently discrepant results obtained across previous studies may not necessarily be completely reconciled, but rather interpreted as separate illustrations of the large diversity of demographic histories experienced by our species in Africa since its emergence in a remote past. This further strongly advocates for being careful when merging individual samples among genetically differentiated populations in *a priori* categories based on geography, subsistence-strategy, linguistics, and/or shared genetic ancestry criteria. Instead, we recommend developing novel approaches, probably at a large computational cost as experienced here, to explicitly take into account African genetic diversity and genetic structure at the continental, regional, and/or local scales, and *a posteriori* interpreting the diversity of results obtained separately.

### Short periods of gene-flow rather than recurring migrations in the ancient history of Africa

As pointed out here and in numerous previous theoretical or empirical studies (e.g. ^22–24,26,31,68–72^), whether ancient gene-flow processes occurred recurrently or more instantaneously across lineages has fundamental consequences on the biological and cultural evolution of our species. Indeed, recurrent gene-flow processes allow for continuous allelic exchanges across populations-thus never reproductively isolated throughout entire periods of evolution-, which may strongly influence the relative influence of drift and selection across populations and the notion of ancient tree-like evolution in human populations in Africa. We found that instantaneous gene-flow processes systematically vastly outperformed recurring migration processes, whichever the tree-topology for ancient and recent Central and Southern African lineages divergences, and for all combinations of five observed populations. Therefore, our results unambiguously favor an evolutionary history of African lineages ancestral to a variety of Central and Southern African populations where *Homo sapiens* populations experienced long periods of isolation and drift, followed by short periods of possibly asymmetric gene-flow, which may in turn have induced some reticulations among lineages ^22,73^. Notably, this scenario further allows for differential selection and adaptation processes across lineages (e.g. ^6,10,23^), and for ancient admixture-related selection processes within Africa, analogous to previous findings of such processes having occurred more recently throughout the continent ^7,9,74,75^).

Nevertheless, it is certain that *Homo sapiens* evolutionary history of ancient and recent gene-flow in Africa has involved more complex processes among pairs of lineages than the ones here identified. Previous studies considered arguably more realistic models to explain their data, with either a conjunction of recurring symmetric migrations among pairs of lineages with possible instantaneous introgressions among certain pre-specified recent and/or ancient lineages ^22,24^, or only pre-specified instantaneous introgressions without recurring gene-flows ^23,26^, including specific events involving the contributions from *Homo sapiens* or non*-Homo sapiens* unsampled lineages. In this context, our results highlight that complex instantaneous introgression scenarios should probably be preferred as a starting-point from where one could then increase complexity, for instance by considering more than a single instantaneous gene-flow event among pairs of ancient lineages.

Moreover, our results advocate for the possibility of at least some such introgressions to be relatively intense, as scenarios considering low or moderate gene-flows, whether instantaneous or recurring, are systematically poorly mimetic of observed genomic patterns. Echoing this result, a previous study identified high levels of instantaneous introgressions across African lineages as well ^26^; keeping in mind that the authors considered very different maximum-likelihood approaches based on different statistics and scenario specifications, and different population samples.

Nevertheless, our approach failed to provide satisfactory posterior estimates of gene-flow intensities, a serious limitation which we discuss extensively in the methodological limits and perspectives section below. Importantly, previous studies investigating conjunctions of recurring and more instantaneous migrations overall estimated that symmetric recurring gene-flows were weak, whether recent or ancient (e.g. ^22^). Although our results are difficult to compare with previous studies due to strong differences in model specifications, statistics used and methodological approaches as well as population samples, this latter previous result may be reflected in ours that disfavor recurring migration models as being poorly explicative of our data.

### Inferred demographic history in Central and Southern Africa

Extensive previous work agrees that divergences among lineages ancestral respectively to extant Khoe-San, Rainforest Hunter-Gatherers, and Rainforest Hunter-Gatherer neighbors were among the most ancient in *Homo sapiens* evolution (e.g. reviewed in ^2,3^). However, the relative order of their divergences as well as their timing has been a matter of extensive debate ^25,26,40^. This is due in part to split-time inferences conducted on tree-like topologies considering gene-flow or no gene-flow, using ancient DNA data or not, and to differing methods, statistics, and population samples.

#### Most ancient divergences in Africa

Here, we formally tested, with Random-Forest ABC scenario-choice, which ancient tree-topology best explained extant genomic patterns, whichever the gene-flow processes and intensities across pairs of recent and ancient lineages, for 54 different combinations of five different Khoe-San, Rainforest Hunter-Gatherer, and Rainforest Hunter-Gatherer neighboring populations. It is the first time to our knowledge that such formal comparison of numerous competing-scenarios is conducted systematically for a variety of sets of population samples at a regional and local scale, rather than comparing maximum-likelihood values from vastly differing models, each obtained separately using differing population samples ^22,25,26^. Importantly, note that ^24^ also explored highly complex demographic scenarios in Africa with ABC, focusing on varying unidirectional introgression processes among certain lineages for a fixed tree-shape among, in particular, three extant African populations at the continental scale.

Our results show that whichever the gene-flow processes considered, the lineage ancestral to extant KS populations diverged first from the lineage ancestral to RHG and RHGn, for almost all combinations of sampled populations. Furthermore, we estimated with Neural Network ABC posterior-parameter inferences that this original divergence likely occurred in a remote past ∼265,000-398,000 years ago, followed by the divergence between ancestral RHG and RHGn lineages ∼127,000-190,000 years ago, considering conservatively 20 or 30 years per generation ^66,67^ and with significant variation depending on combinations of sampled populations considered. Recent paleo-genomic data from ancient southern African individuals also demonstrate deep population structure and long-term isolation in southern Africa, and show that present-day Khoe-San populations retain only part of this ancient structure following later gene-flow ^76^. Importantly, the ABC framework employed here explicitly models gene-flow processes when inferring past population structure, allowing deep divergence signals to be evaluated in the presence of migration.

Both our estimates fall within the upper bound of previously obtained results considering a variety of similar population samples, genetic data, inference methods, and statistics, albeit scenarios may not have always been specified in analogous ways ^4–6,8,25,26,40,77^. Note however, that considering different combinations of populations at the regional and local scales allowed us to identify a small minority (4/54) of such combinations for which an alternative scenario (Scenario i1-3b), where RHGn lineages diverged first from a lineage ancestral to all RHG and KS extant populations, somewhat analogous to those proposed in ^26^, would better explain the data consistently across our analyses, again whichever the gene-flow processes and range of intensities considered. This latter result, in addition to the variation in posterior estimates observed across combinations of populations for the proposed synthetic scenario, may explain apparently discrepant results across some previous studies ^22,24–26^, which would then be due to differing population samples used for demographic inferences and/or to merging samples from differentiated populations with shared genetic ancestries, thus advocating for further accounting for the vast genetic diversity among African populations even at a local scale in future studies ^72^.

Note that, in ^22^, the fundamental divergence between the Nama Khoe-San sample and other Sub-Saharan populations was dated to ∼110,000-135,000 years ago under the two best-fitting models, thus strongly discrepant with our findings. However, these important results are very difficult to compare with ours, since the authors did not consider any Central African Rainforest Hunter-Gatherers nor their neighbors in their inferences, and since they investigated highly complex models specified completely differently from those here envisioned. In particular, Ragsdale and colleagues included in their models possible very ancient genetic structures, long before *Homo sapiens* emergence, a feature that is unspecified in our scenarios which considered simply a single ancestral population in which all extant lineages ultimately coalesce. Nevertheless, note that substructure and reticulation within the ancestral population is not *per se* incompatible with our scenarios. In fact, it may be compatible with our posterior estimates of a large effective population ancestral to all extant populations here investigated, among the largest of inferred ancient and recent Central and Southern African effective population sizes. Therefore, it will be reasonable in future work to complexify the scenarios here proposed to evaluate whether very ancient substructures and reticulations within our ancestral population, prior to the original divergence between Southern and Central African populations, may improve the fit to the observed genomic data, as proposed by ^22^.

#### More recent divergences in Central and Southern Africa

More recently during the evolutionary history of Sub-Saharan Africa, we found that the divergence among Northern and Southern Khoe-San populations largely pre-dated the divergence of Western and Eastern Congo Basin Rainforest Hunter-Gatherers, a question rarely addressed to our knowledge. First, we found that KS divergence dated sometime between 57,000 and 85,000 years ago, thus substantially more recently than estimates previously proposed ^2^. Beyond vast differences in models, methods, and statistics used to provide either inference, note that our results for this divergence time were substantially variable across pairs of sampled populations used in each analysis, some specific sets of populations providing posterior estimates consistent with previous results.

Interestingly, our synthesized estimates for the Northern and Southern Khoe-San population divergence were relatively synchronic to the genetic onset of the Out-of-Africa (e.g. ^2^). Population genetics inferences only provide possible mechanisms to explain observed genetic patterns and are, in essence, not addressing the possible causes underlying the inferred mechanisms. In this context, we may hypothesize that the global climatic shifts inducing massive ecological changes that have occurred in Africa at that time (e.g. ^78^), sometimes proposed to have triggered ancient *Homo sapiens* movements Out-of-Africa, may also have triggered, independently, the genetic isolation among ancestral Khoe-San populations. Nevertheless, where the ancestors of extant Khoe-San populations lived at that time remains unknown and is nevertheless needed to further elaborate possible scenarios for the causes of the genetic divergence here inferred.

We found that Rainforest Hunter-Gatherer populations across the Congo Basin diverged long after this divergence, roughly between 15,000 and 23,000 years ago, relatively consistently across pairs of sampled populations used for inferences; estimates highly consistent with previous studies ^8,77,79^, despite major differences in gene-flow specifications across RHG groups between studies. Interestingly, this divergence time is relatively synchronic with absolute estimates for the Last-Glacial Maximum in Sub-Saharan Africa (e.g. ^80^). The fragmentation of the rainforest massif during this period in the Congo Basin may have induced isolation between Eastern and Western RHG extant populations, as plausibly previously proposed ^77^. However, similarly as above for the Northern and Southern Khoe-San populations divergence, where the ancestors of extant Eastern and Western Rainforest Hunter-Gatherers lived remains unknown, which prevents us from formally testing this hypothesis ^81^.

Altogether, these results show that Central African Rainforest Hunter-Gatherer and Southern African Khoe-San populations have had, respectively, extensive time for selection processes, including adaptive introgression processes, to have influenced independently both groups of populations as well as populations within each group separately (e.g. ^6,7,9,10^).

#### Ancient and recent instantaneous gene-flow times in Africa

We obtained reasonably well estimated instantaneous asymmetric gene-flow times among almost all pairs of ancient and recent lineages, albeit the variation of estimates across sets of Central and Southern African sampled populations increased substantially with most ancient times. In this context, we deem it hard to confidently try to interpret the most ancient event of instantaneous gene-flow between the two most ancestral lineages in our tree-topology. However, other instantaneous gene-flow time estimates throughout the topology were more consistently estimated overall, and showed relative synchronicity in some cases, which has never been reported before to our knowledge, even if they were specified independently in our models and drawn from large distributions *a priori*.

Interestingly, we found strong indications for almost synchronic events of introgressions having occurred during the Last Interglacial Maximum in Africa ^73^, between ∼85,000 and ∼135,000 years ago (when considering conservatively 20 or 30 years per generation ^66,67^), with coherent, even if relatively large, credibility intervals across combinations of sampled populations. They involved gene-flow between lineages ancestral to Khoe-San populations and ancestors of Rainforest Hunter-Gatherer neighbors on the one hand and, on the other hand, between lineages ancestral to Khoe-San populations and the lineage ancestral to all Rainforest Hunter-Gatherers. An increase in material-based culture diversification and innovation, possibly linked to climatic and environmental changes locally, has previously been observed during this period of the Middle Stone Age in diverse regions of continental Africa; prompting a long-standing debate as to its causes if human populations were subdivided and isolated biologically and culturally at the time ^82–85^.

In this context, our results instead may suggest that population movements at that time among previously isolated populations may itself have triggered the observed increased cultural diversification locally, even if obvious signs of pan-African cultural spread at the time are difficult to assess ^84^. In turn, it would interestingly echo the known effect of within-population genetic diversity increase induced by genetic admixture between previously isolated populations (e.g. ^18,31,68,86^).

Then, note that we estimated that the instantaneous gene-flow event between the ancestral Rainforest Hunter-Gatherers lineage and that of their extant neighbors seemingly occurred synchronically to the genetic Out-of-Africa (^78^; see above). This would imply that possible climatic and ecological shifts at that time may not have only induced population divergences and displacement, but may also have triggered population gene-flow.

Relatively more recently, around 30,000 years ago, we found two loosely synchronic gene-flow events between ancestors to extant Central African Rainforest Hunter-Gatherer neighbors’ lineages and, separately, Northern and Southern Khoe-San lineages. This corresponds to the end of the Interglacial Maximum and a period of major cultural changes and innovations during the complex transition from Middle Stone Age to Late Stone Age in Central and Southern Africa ^82,85,87–89^. Nevertheless, connecting the two lines of genetic and archaeological evidence to conclude for increased population movements at the time and their possible causes should be considered with caution. Indeed, in addition to genetic-dating credibility-intervals being inherently much larger than archaeological dating, this period remains highly debated in paleoanthropology mainly due to the scarcity and complexity of the material-based culture records, and that of climatic and ecological changes locally, across vast regions going from the Congo Basin to the Cape of Good Hope ^82,85,87–89^.

Finally, we found several signals for instantaneous gene-flow events having occurred among Central African lineages and among Southern African lineages, between 6,000 and 15,000 years ago, during the onset of the Holocene in that region, shortly before or during the beginning of the last Post Glacial Maximum climatic crisis in Western Central Africa ^90^, the emergence and spread of agricultural techniques ^91^, and the demic expansion of now-Bantu-speaking populations from West Central Africa into the rest of Central and Southern Africa ^9,17,92^. These results are consistent with previous investigations that demonstrated the determining influence of Rainforest Hunter-Gatherer neighboring populations’ migrations through the Congo Basin in shaping complex socio-culturally determined admixture patterns ^4,5,8,77^, including admixture-related natural selection processes ^9,10,79,93^. Our estimates for introgression events are in the upper bound of previous estimates for the onset of the so-called “Bantu expansion” throughout Central and Southern Africa. Since we do not try to determine the geographical areas of occupation of ancestral lineages in this work, we may hypothesize here that major climatic and ecological changes that have occurred at that time may have triggered increased population mobility and gene-flow events between previously isolated populations, rather than consider that the Bantu-expansions themselves were the cause for the gene-flow events here identified.

Finally, we did not find signals of more recent introgression events from Bantu-speaking agriculturalists populations into Northern or Southern Khoe-San populations, in particular among the !Xun, albeit such events have been identified in several previous studies (see ^2^). This is likely due to the fact that we considered only a limited number of individual samples from each population with low apparent levels of recent admixture, and therefore may lack power to detect these very recent events with our data and approach.

### Conceptual, methodological, and empirical limitations and perspectives for inferring ancient histories from observed genomic data

All previous attempts at reconstructing the human evolutionary histories which led to genomic patterns observed today across African populations have faced major conceptual, methodological, and empirical challenges. Conceptually, large amounts of more-or-less nested scenarios can be envisioned *a priori* to explain extant genetic diversity, based on previous results from paleo-anthropology, population genetics, and paleogenomics. These scenarios may range from tree-like models without gene-flow events among ancient or recent lineages to complex networks of weakly differentiated populations exchanging migrants over large periods of time, with or without the contribution of ancient *Homo* or non-*Homo* now extinct or unsampled lineages. Systematically exploring all possible models is often methodologically out of reach due to differing fundamental scenario-specifications or, when scenarios are specified and parameterized in analogous ways, due to scenarios’ nestedness and un-identifiability in certain parts of their parameter spaces (e.g. ^94^). Empirically, formal scenario comparisons are first hampered by necessarily limited amounts of genomic data representative, at continental, regional, and local scales, of the known diversity and differentiation of human populations in Africa; in addition to yet limited amounts of ancient DNA data throughout the continent. Finally, empirical limitations also emerge from the use of different statistics to explore genomic diversity patterns, which possibly each capture different facets of human evolutionary histories, thus providing discrepant results and interpretations only in appearances. Machine-learning ABC procedures provide advantages, in particular concerning the formal exploration of competing scenarios’ fit to observed data across numerous highly complex sometimes nested scenarios and using numerous summary-statistics ^43,45,63,94^. However, all the above challenges are also largely faced in ABC and, therefore, in this study.

While most divergence and gene-flow times, as well as some effective sizes, were inferred satisfactorily in our results, the lack of satisfactory posterior estimates for all 20 gene-flow rates parameters consistent among the 25 sets of population combinations, as well as that for some ancient effective sizes, likely stems from different limitations. First, considering only five individual genomes per population is inherently limiting when trying to estimate gene-flow rates from inter-population summary-statistics in ABC ^95^. Furthermore, our inferences rely on summary-statistics computed over only a portion of the human genome and therefore do not integrate information from the entire available genomic data. In future studies, increasing sample sizes and adding statistics based on coalescent times, inter- and intra-individual distribution of admixture fractions and local-ancestries, for larger parts of the genome ^31,32,68,95^, will likely improve the posterior estimation of these parameters using machine-learning ABC approaches ^24^. Note, that simulating larger parts of the genome and computing these complex statistics require significantly improving computation time in ABC frameworks comprising hundreds of thousands of simulations ^96,97^. Furthermore, in all cases, considering multiple sets of different and substantially genetically diverse populations as the ones here considered from Sub-Saharan Africa may inevitably lead to large variation across gene-flow rates’ posterior estimates depending on the specific population-combinations.

Second, for simplicity and as a starting point, we considered a single instantaneous time for the two separate unidirectional gene-flow events, for each gene-flow event between pairs of lineages separately. Therefore, while our RF-ABC scenario-choice results strongly support instantaneous asymmetric gene-flow events rather than recurring asymmetric gene-flows to best explain extant genomic patterns, it is plausible that scenarios where each unidirectional gene-flow event may occur at a different time ^24^, will more realistically explain the observed data. If this is the case, it is possible that the choice of a unique time for two separate gene-flow events rendered the corresponding gene-flow rates harder to identify with our joint NN-ABC posterior parameter estimation procedure. Furthermore, we explored two extreme gene-flow processes, establishing an open competition between only instantaneous gene-flow and only recurring gene-flow processes. While we found that only instantaneous gene-flow processes outperformed only recurring gene-flow processes, whichever the range of intensities for each event and for all combinations of sampled populations, we provide here a possible starting point for future more complex evolutionary scenarios to be tested. For instance, it will be of natural interest to consider, next, more than a single “pulse” of gene-flow between any two ancient or recent lineage.

Third, to reduce the number of free parameters, we considered constant effective population sizes in the terminal branches of our topologies, albeit extensive previous work have inferred diverse recent demographic dynamics across Sub-Saharan African populations (e.g. ^17,65,98–100)^. Importantly, how these recent demographic changes may influence divergence times posterior estimates is not trivial in models as complex as the ones here investigated. For instance, previous studies considered ABC frameworks and a variety of autosomal and uniparental genetic data ^4,5,9,77,101^. They investigated divergence times among sampled lineages assuming possible gene-flow processes and models with very different population-size changes. Overall, while informing a variety of bottlenecks, constant, or increases in effective population sizes, divergence times remained largely consistent across studies that considered the same sets of sampled populations, and consistent with our genome-wide results for comparable sets of sampled populations. Therefore, it will be crucial in future studies to further complexify effective population sizes changes in the proposed topologies to empirically evaluate how they affect or not divergence times estimates; and further disentangle the complex evolutionary and migration history of African populations using increasingly realistic scenarios.

Fourth, the neural networks used here may have been unable to satisfactorily identify the 20 gene-flow parameters among the 43 jointly-estimated parameters, due to their lack of complexity when considering a unique layer of hidden neurons. For instance, and as a first step, considering Multilayer-Perceptrons in the future, *i.e.* more complex feed-forward neural networks with additional layers of hidden neurons, may help posterior estimations of these parameters (e.g. ^102^), at the known cost of non-trivial parameterization and optimization of the neural networks themselves (e.g. ^97,103,104^).

Altogether, joint posterior estimation of numerous gene-flow rates parameters and the timing of their occurrence under highly complex demographic scenarios remains one of the most challenging tasks in population genetics. It will undoubtedly benefit from future analytical theoretical developments ^105,106^, and the improvement of machine-learning-based inference procedures, whether for ABC or other analysis frameworks (e.g. ^104,107–109)^.

### Archaic admixture with *Homo sapiens* or non-*Homo sapiens* unsampled lineages?

Our results show that explicitly considering archaic admixture from unsampled populations is not strictly necessary to explain parts of the observed genomic diversity of extant Central and Southern African populations when considering jointly the 337 relatively classical population genetics summary-statistics used here for demographic inferences, consistently with a previous study ^22^, and conversely to others ^3,25,26^. We did not explore possible contributions from unsampled lineages, whether from non-*Homo sapiens* or from ancient “ghost” human populations, and therefore cannot formally evaluate such possibilities.

Our set-up formally comparing competing-scenarios rather than comparing posterior likelihoods of highly complex yet vastly differing models, provides a reasonable framework for future complexification of scenarios comprising possible contributions from ancient or ghost unsampled populations; where the burden of proof lies on showing that scenarios with archaic admixture fit specific parts of the data significantly better than scenarios without such archaic admixture (incorporating the increased complexity and more parameters). Such endeavors will also benefit from the explicit use of additional novel summary-statistics (^22,26,32^; see also above).

In any case, the complexification of scenario-specifications to account for possible past “archaic” or “ancient” introgressions will not fundamentally solve the issue of the current lack of reliable ancient genomic data older than a few hundreds or thousands of years from Sub-Saharan Africa ^3,110,111^. Indeed, analogously to archaic admixture signals that were identified outside Africa only when ancient DNA data were made available for Neanderthals and Denisovans (e.g. ^27,28^), we imperatively need to overcome this lack of empirical ancient DNA data in Africa to formally test whether, or not, ancient human or non-human now extinct lineages have significantly contributed to shaping extant African diversity.

## Material and Methods

### Population samples

#### Central and Southern Africa dataset

We investigated high-coverage whole genomes newly generated for 74 individual samples (73 after family relatedness filtering, see below), from 14 Central and Southern African populations (**Figure 1**, **Table 1, Table S1**). Based on extensive ethno-anthropological data collected from semi-directed interviews with donors and their respective communities, we grouped *a posteriori* the 73 individual samples from 14 populations in three larger categories.

The Baka, Ba.Kola, Bi.Aka_Mbati, Ba.Twa, and Nsua individuals were categorized as Rainforest Hunter-Gatherers (RHG), based on several criteria including self-identification and relationships with other-than-self, socio-economic practices and ecology, mobility behavior, and musical practices ^4,38^. Note that the Bi.Aka_Mbati individuals are Aka RHG speaking Mbati (Bantu C10) and from a different Aka community than the Bi.Aka from the HGDP-CEPH and SGDP databases who speak Aka and Ngando Bantu C10 languages, each language corresponding to the specific group of neighboring populations with whom each group shares privileged complex socio-economic relationships ^4,38^.

The Nzime, Ngumba, Ba.Kiga, and Ba.Konjo individuals were categorized as Rainforest Hunter-Gatherer neighbors (RHGn), based on the same set of categorization-criteria as RHG. Indeed, RHG and RHGn populations in Central Africa are known to identify themselves separately and to live in different ways in the Central African rainforest, whilst often sharing languages locally as well as complex socio-economic relationships and interactions that may include intermarriages ^5^. Historically, RHG populations have been designated as “Pygmies” by European colonists, an exogenous term from ancient Greek that is sometimes used derogatorily by the RHGn.

Finally, the Nama, Ju|’hoansi, Karretjie People, !Xun, and Khutse_San individuals were categorized as Khoe-San populations (KS) based primarily on self-identification and relationships with other-than-self locally, lifestyle, and languages ^112^. Note that research was approved by the South African San Council. Individuals from these populations were whole-genome sequenced anew with improved methods (see below), compared to previous work ^40^. The Botswana and Namibia genomes were previously published in ^40^, where the collection procedures, national ethical approvals, sequencing approvals, and publication permissions are described in detail. In the present study, we generated new sequencing data from the DNA libraries in order to minimize potential technical biases between previously published genomes and the newly sequenced Central African samples.

Note that the three groups can be further subdivided based on geography into Western and Eastern RHG and RHGn, and into Northern, Central, and Southern KS (**Figure 1**, **Table 1**).

#### Comparative dataset

The 74 genomes were analyzed together with 105 (104 after family relatedness filtering, see below) high-coverage genomes from 32 worldwide populations (**Figure 1**, **Table 1, Table S1**). 64 samples from the Simon’s Genome Diversity Project (SGDP) ^34^; 9 HGDP samples ^27^; 7 samples from the 1000 Genomes project (KGP) ^35^; one Karitiana sample ^36^; 24 samples from the South African Human Genome Project (SAHGP) ^37^. Selection criteria were: Illumina paired-end reads, high-coverage (>30X), access to raw data and no (known) genetic substructure within population.

Note that, although samples from these datasets have not been collected by us for the purpose of this study, based on previous knowledge and publications, the Biaka and Mbuti HGDP samples can be categorized as RHG and the #Khomani and Ju|’hoansi_comp samples as KS (**Figure 1**, **Table 1**).

### Data generation

#### DNA extraction

DNA was extracted from saliva with the DNA Genotek OG-250 kit for the Baka, Nzime, Bi.Aka_Mbati, Nsua, and Ba.Konjo samples. DNA was extracted from buffy coats with DNeasy Blood&Tissue spin-column Qiagen^TM^ kits for the Ba.Kola, Ngumba and Ba.Kiga samples. Both extraction methods were used for the Ba.Twa samples. For the Nama, Ju|’hoansi, Karretjie People, !Xun, and Khutse_San samples, DNA was extracted from EDTA-blood using the salting-out method ^113^.

#### Library preparation and sequencing

Library preparation and sequencing was performed by the SciLifeLab SNP&SEQ Technology platform in Uppsala, Sweden. Libraries were prepared with TruSeq DNA preparation kits, and paired-end sequencing (150 bp read length), was performed on Illumina HiSeqX machines with v2 sequencing chemistry, to a coverage of 30X or more. For the Southern African samples, we included paired-end data (100 bp read length) sequenced on Illumina HiSeq2000 machines obtained previously for the same libraries ^40^.

#### Sequencing data quality-control processing and relatedness filtering

The processing pipeline used in the present study is adapted from the Genome Analyses Toolkit (GATK) “Germline short variant discovery (SNPs + Indels)” Best Practices workflow ^114–116^. It is described and compared to the original GATK Best Practices workflow in ^117^. **Figure S54.** gives an overview of the pipelines used for the processing; all template codes with accompanying detailed explanations for all processing steps are provided below and in the corresponding GitHub repository (https://github.com/Gwennid/africa-wgs-descriptive).

Reads were mapped to a decoy version of the human reference genome, GRCh38 (1000 Genomes Project version), with BWA-MEM ^118^ from BWAKIT v0.7.12 using a strategy appropriate for ALT contigs. The input was either FASTQ or BAM files that were reverted to unmapped BAM or to FASTQ prior to mapping. Mapped reads were selected with SAMtools version 1.1 and Picard version 1.126 (https://broadinstitute.github.io/picard/).

Note that, as mentioned in ^119^, the GRCh38 is a “mosaic haploid representation of the human genome”. The dominant sequence contribution (70%) is from a donor of “likely African-European admixed ancestry”. Therefore, this assembly of the reference genome is not equally genetically related to all extant individuals within and among human populations. It is important to keep in mind, thus, that while numbers of variants can be compared across studies referring to the same genome assembly, in any given study, some populations may have artificially high number of variants as compared to the reference.

Duplicate reads were marked (at the lane level) with Picard version 1.126 MarkDuplicates. Realignment around indels was performed (at the lane level) with GATK version 3.5.0 RealignerTargetCreator and IndelRealigner. We then performed a “triple mask base quality score recalibration (BQSR)”, where the sample’s variation is used together with dbSNP, as described in ^40,117^. This step includes merging the reads from a given sample with SAMtools version 1.1; calling variants with GATK version 3.5.0 HaplotypeCaller with “--genotyping_mode DISCOVERY”; and (in a parallel track) standard BQSR with GATK version 3.5.0 BaseRecalibrator and PrintReads, and dbSNP version 144. Following triple mask BQSR (performed at the lane level), the recalibrated reads from a given sample are merged with SAMtools version 1.1, sorted and indexed with Picard version 1.126 SortSam. Duplicate marking and realignment around indels were performed again, at the sample level.

Variant calling was performed independently for the autosomes and the X chromosome, due to the difference in ploidy. SNP and short indels were first called with GATK version 3.7 HaplotypeCaller in each genome, with the “--emitRefConfidence BP_RESOLUTION” option. Multi-sample GVCFs were then generated with CombineGVCF, and variants were finally jointly called with GATK GenotypeGVCFs. Genotypes were emitted for each site (option “allSites”).

Variants were filtered with GATK version 3.7 Variant Quality Score Recalibration (VQSR) using recommended resources from the GATK bundle: HapMap (version 3.3), 1000 Genomes Project Illumina Omni 2.5M SNP array, 1000 Genomes Project phase 1 high confidence SNPs, and dbSNP version 151 for the SNPs; and gold standard indels ^120^ and dbSNP version 151 for the indels. For the X chromosome, autosomal variants were included for training the VQSR model (VariantRecalibrator step).

We controlled the dataset for related individuals with KING ^121^ “--kinship” and plink version 1.90b4.9 ^122^ “--genome --ppc-gap 100”. The dataset was pruned for linkage disequilibrium (LD) before relatedness estimation with plink, using plink “--indep-pairwise” with sliding windows of 50 SNPs, shifting by five SNPs, and a r^2^ threshold of 0.5. Both methods identified two pairs of first-degree relatives. We excluded the sample with greatest missingness from each pair using GATK version 3.7 SelectVariants to produce the family unrelated working datasets used henceforth.

The callset was then further refined; a “FAIL” filter status was set on sites that are ambiguous in the reference genome (base “N”) or had greater than 10% missingness, identified with VCFtools version 0.1.13 “--missing-site” ^123^. Moreover, for the autosomes, variants heterozygous in all samples were marked as failed (Hardy Weinberg Equilibrium-HWE-filter, identified with VCFtools version 0.1.13 “--hardy”).

Coverage was computed with QualiMap version 2.2 ^124^ including or not duplicates (“-sd” option for the latter). The average coverage (without duplicates) across individuals within populations is provided in **Table S3**.

### Descriptive analyses

#### Variant counts

We obtained per-sample and aggregated metrics of variant counts (**Figure S1**), with the tool *CollectVariantCallingMetrics* of the software package Picard v2.10.3 (https://broadinstitute.github.io/picard/), applied to the full 177 worldwide individuals’ dataset after VQSR, family relatedness, HWE and 10% site missingness filtering. We conducted variant counts procedures for chromosomes 1 to 22. We used dbSNP 156 as a reference; we downloaded “GCF_000001405.40.gz” and modified the contig names to match contig names in the VCFs. Separately, we applied the same variant counts pipeline as above to the Central and Southern African 73 individuals’ original dataset extracted from the full dataset with BCFtools version 1.17 (https://github.com/samtools/bcftools), using view with the option “-S”. All scripts are provided in the corresponding GitHub repository (https://github.com/Gwennid/africa-wgs-descriptive).

#### Genome-wide Heterozygosity

We calculated observed and expected heterozygosities, for the autosomes and for the X-chromosome separately using custom Python, Bash, and R scripts (full pipeline available at GitHub https://github.com/Gwennid/africa-wgs-descriptive), for the 177 worldwide individuals after variant-counts filtering described above. For autosomes, we considered the main contigs (“chr1” to “chr22”). For the X chromosome, we excluded the Pseudo-Autosomal Regions with coordinates from GRCh38 (PAR1: 10,000 to 2,781,480, PAR2: 155,701,382 to 156,030,896).

For each population with more than one individual and for each autosome and X-chromosome separately, we counted the number of variable and non-variable sites and excluded multi-allelic sites, indels, and sites with missing genotypes in at least one individual in the population for simplicity and conservativeness. For each autosome and each individual, we then counted separately the numbers of observed homozygous and heterozygous sites of each configuration compared to the reference (0/0; 0/1; 1/0; 1/1). Based on these counts, for each population and each chromosome separately, we first calculated observed heterozygosities simply as the average proportion of heterozygous individuals per variable sites with no-missing data in the population sample. We also computed unbiased expected multi-locus heterozygosities for all variable loci with no missing genotypes in the population as in equations (2) and (3) in ^47^. We averaged this value across all sites with no missing information in the population sample including both variable and non-variable sites, and finally corrected it for haploid population sample sizes (**Figure S1**). We compared unbiased heterozygosities by chromosomes across groups of populations using Wilcoxon signed-rank tests. All scripts are provided in the corresponding GitHub repository (https://github.com/Gwennid/africa-wgs-descriptive).

#### Runs of homozygosity

We identified runs of homozygosity (ROH) using the *homozyg* tool from PLINK version 1.90b4.9 ^122^. We selected autosomal biallelic SNPs from the variant-counts pipeline described above with GATK version 3.7 SelectVariants with options “-selectType SNP-restrictAllelesTo BIALLELIC-excludeFiltered”. We converted the VCF to TPED and then binary plink fileset with VCFtools version 0.1.13 “-plink-tped” and plink. For the 177 individuals worldwide, we considered only ROHs measuring more than 200 Kb (--*homozyg-kb* 200), containing at least 200 SNPs (*--homozyg-snp* 200), containing at least one variable site per 20 Kb on average (*-- homozyg-density* 20), and with possible gaps of up to 50 Kb (*--homozyg-gap* 50). Default values were considered for all other parameters: *--homozyg-window-het* 1 *--homozyg-window-snp* 50 *--homozyg-window-threshold* 0.05. Finally, the total mean ROH lengths were calculated for each population separately using the “.hom” output-file for each of four length categories: 0.2 to 0.5 Mb, 0.5 to 1 Mb, 1 to 2 Mb, and 2 to 4 Mb (**Figure S1**). Corresponding pipelines are provided in the corresponding GitHub repository (https://github.com/Gwennid/africa-wgs-descriptive).

#### Individual pairwise genome-wide genetic differentiation

We explored genome-wide genetic differentiation between pairs of individuals with Neighbor-Joining Tree (NJT) ^52,53^, and Multi-Dimensional Scaling (MDS) approaches based on the pairwise matrix of Allele-Sharing Dissimilarities (ASD) ^51^ computed using 14,182,615 genome-wide autosomal SNPs pruned for low LD, with PLINK version 1.90b4.9 ^122^ *indep-pairwise* function (*--indep-pairwise 50 5 0.1*). We considered at first all 177 unrelated individuals in our dataset to calculate the ASD matrix, and then subset this ASD matrix for the 73 Central and Southern African unrelated individuals original to this study before computing NJT and MDS analyses anew (**Figure 2, Figure S3**).

We computed the ASD matrix considering, for each pair of individuals, only those 14,182,615 SNPs without missing data, using the *asd* software (v1.1.0a; https://github.com/szpiech/asd; Szpiech, 2020). We computed the unrooted NJT using the *bionj* function of the R package *ape* and setting the branch-length option to “true”. We computed the MDS using the *cmdscale* function in R.

We compared ASD values with populations’ geographic locations by computing the standardized average ASD across all pairs of individuals from two different populations, for all pairs of African populations separately (**Figure 1**, **Table 1, Table S1**), and compared these values with the logarithm of geographic distances among pairs of populations (setting 0 km distances to “1”), using Spearman *ρ* and Pearson *r* correlations for different continental, sub-continental, and regional sets of populations. We tested these correlations using 1000 Mantel permutation tests (**Table S4**), with the *partial.mantel.test* function of the *ncf* package in R. We computed the shortest geographic distance between two populations’ GPS coordinates on an Earth-like ellipsoid (with an earth radius of 6378.137 km by default in WGS84), using the *distm* function with *distGeo* method from the *geosphere* package in R.

We conducted permutational ANOVA analyses to test, separately, whether linguistic families, sub-linguistic families, or traditional food-producing strategies (**Table S1**), significantly explain average population-pairwise ASD patterns, using the *adonis2* function from the *vegan* package in R with 1000 permutations for each analysis conducted on subsets of the above ASD matrix (**Figure 2, Figure S3**) at varying geographical scales (**Table S5**, **Table S6**, **Table S7**).

The clustering software ADMIXTURE ^54^ allows researchers to further explore inter-individual genome-wide levels of dissimilarity and resemblance. Indeed, while it is tedious and often cognitively difficult to explore multiple combinations of dimensions of genetic variation using an MDS or a NJT approach, ADMIXTURE instead allows exploring K higher such dimensions at once ^54,56–59^. However, note that the visual representation of relative “distances” across pairs of individuals is lost in the classical bar-plot representation of ADMIXTURE, hence showing the complementarity of this descriptive analysis to the above MDS and NJT.

We considered here only 840,031 genome-wide autosomal SNPs pruned for LD (r^2^ threshold 0.1) and with Minimum Allele Frequency above 0.1, for the 177 worldwide unrelated individuals, using PLINK *indep-pairwise* and *maf* functions. We computed unsupervised ADMIXTURE version 1.3.0 clustering with values of K ranging from 2 to 10, considering 20 independent runs for each value of K separately. We then calculated ADMIXTURE results symmetric-similarity-coefficient SSC for each value of K separately in order to find the groups of runs providing highly similar results (SSC>99.8%). Individual’s genotype membership proportions to each K cluster were then averaged, per individual, across such highly resembling runs, and then plotted. SSC calculations, averaging results across similar runs, and producing barplots were conducted using the software PONG with the “greedy” algorithm ^55^. Corresponding pipelines are provided in the corresponding GitHub repository (https://github.com/Gwennid/africa-wgs-descriptive). For each value of K separately, results representing the majority of the 20 independent runs are presented in **Figure 3**, and all other results are presented in supplementary **Figure S4**.

### Machine-Learning Approximate Bayesian Computation scenario-choice and posterior parameter estimation

We reconstructed the complex demographic history of Central and Southern African populations using machine-learning Approximate Bayesian Computation (ABC) ^41–45^. In principle, in ABC, researchers first simulate numerous genetic datasets under competing scenarios by drawing randomly a vector of parameter values for each simulation in distributions set a priori by the user. For each simulation separately, we then calculate a vector of summary statistics, thus corresponding to a vector of parameter values used for the simulation. The same set of summary statistics is computed on the observed data. ABC scenario-choice then allows the researcher to identify which one of the competing scenarios produces the simulations for which summary-statistics are closest to the observed ones. Under this best scenario, ABC posterior-parameter inference procedures allow the researcher to estimate the posterior distribution of parameter values most likely underlying the observed genetic patterns.

### 48 competing scenarios for the demographic history of Central and Southern African populations

We designed 48 competing demographic-history scenarios possibly underlying the genetic patterns observed in five Central and Southern African populations of five individuals each (**Table 1**, **Figure 4**); one Eastern Rainforest Hunter-Gatherer population (eRHG: Nsua, Ba.Twa, or Mbuti), one Western Rainforest Hunter-Gatherer population (wRHG: Baka, Ba.Kola, or Bi.Aka_Mbati), one Eastern or Western RHG neighboring population (eRHGn: Ba.Kiga or Ba.Konjo; wRHGn: Nzime or Ngumba), one Northern Khoe-San population (nKS: Ju|’hoansi or !Xun), and one Southern Khoe-San population (sKS: Karretjie People or Nama). Note that we did not include the Khutse_San population in ABC inferences for simplicity, as this population is located at intermediate geographic distances between the nKS and sKS groups of populations.

The 48 competing scenarios differed in their ancient tree-topologies and relative timing of the more recent divergence events, combined with different (duration and intensities) possibly asymmetric gene-flow processes among ancient and recent lineages (**Figure 4**). This design, while complex, allowed us, for the first time to our knowledge, to consider explicitly the expected confounding effects of gene-flow processes on otherwise different tree-topologies, which may also possibly induce certain reticulations among ancient lineages, while jointly estimating divergence times, gene-flow events’ timing and intensities, and effective population size changes over time.

Importantly, we relied on extensive previous findings having formally demonstrated the common origin of Congo Basin RHG populations ^4,77^, and that of KS populations ^6,40^, respectively, and thus did not consider all possible tree-topologies for five tree-leaves. Moreover, we considered only a single RHGn sample in each combination, in turn from the East or the West of Central Africa, in order to simplify already highly complex scenarios, as we were not interested here in the detailed demographic history of RHGn which has been extensively studied previously (e.g. ^9,17^). This simplification was deemed reasonable as RHGn populations throughout the Congo Basin and into Southern Africa have previously been shown to be strongly more genetically resembling one another, compared to RHG and KS neighboring populations, or compared to much more genetically dissimilar RHG and KS populations, respectively (e.g. ^4,5,8,9,17,79,93^); a result that we also obtained here (**Figure 4**).

#### Eight competing topologies

We investigated eight competing topologies starting with a single ancestral population and resulting in five different sampled populations in the present (**Figure 4**). The eight topologies differ in which lineage ancestral to modern KS, RHG, or RHGn diverged first from the two others, and in the relative order of divergence events internal to KS (nKS-sKS divergence) and internal to RHG (wRHG-eRHG divergence), in order to consider the known variable demographic histories of Sub-Saharan populations at a regional scale.

Note that divergence and introgression times (all *t* and *tad*, see **Figure 4**, **Table S8**), were each randomly drawn in uniform prior distributions between 10 and 15,000 generations, thus effectively setting an upper limit for the most ancient divergence times among lineages at roughly 450,000 years ago considering an upper-bound of 30 years for human generation duration ^125^. This was vastly anterior to the current estimates for the genetic or morphological emergence of *Homo sapiens* ^126,127^, which allowed us to estimate *a posteriori* the most ancient divergences among our populations, without constraining our assumptions based on previous results obtained with different methods, data, and models. Furthermore, these prior-distribution boundaries allowed for gene-flow events among ancient lineages (see next section), to influence, potentially, the timing of the earliest divergence events among human lineages previously estimated ^2,22,26^. Note that we retained for simulations only those vectors of randomly-drawn parameter-values that satisfied the chronological order of lineage divergences set for each eight topologies, respectively (**Figure 4**, **Table S8**)

In each eight topologies, we incorporated the possibility for changes in effective population sizes, *N* (**Figure 4**), during history along each lineage separately by defining constant diploid effective population sizes parameters separately for each tree-branch, each drawn randomly in U[10-100,000] (**Figure 4**, **Table S8**).

#### Asymmetric instantaneous or recurring gene-flows and their intensities

Migration of individuals between populations is ubiquitous in human history (for an overview in Africa, see e.g. ^2,3^). Here we aimed at disentangling the nature of migration processes that may have occurred across pairs of lineages throughout the history of Central and Southern African populations.

In particular, we first aimed at determining whether gene-flow events during history occurred relatively instantaneously, leading to reticulations in tree-topologies, or, conversely, occurred recurrently over longer periods of time leading to un- or weakly-differentiated lineages throughout history ^1,21,22^. To do so, for each eight topologies described above (**Figure 4**), we simulated gene-flow either as single-generation gene-flow pulses across pairs of lineages, or as constant recurring gene-flow across pairs of lineages in-between each lineage-divergence event. This design resulted in 16 competing scenarios encompassing eight different possible topologies, contrasting instantaneous gene-flow events with recurring gene-flows. Note that, for simplicity, we did not consider possible gene-flows between KS and RHG recent lineages, as genetic signatures of such recent migrations have never been identified in previous genetic studies nor in historical records to our knowledge.

Importantly, we aimed at determining whether gene-flow occurred symmetrically, or not, across pairs of lineages, in particular in the past. Indeed, previous studies already identified recent asymmetric admixture processes between RHG and RHGn ^4,5,8,77^, and such possible asymmetries have not been explored among ancient lineages in previous studies ^22,26^, although they may be influencing ancient tree-topologies or reticulations across lineages and the subsequent past evolution of extant populations.

To do so, for each gene-flow event in each 16 competing scenarios (**Figure 4**), we parameterized separately the introgression of lineage A into lineage B, from that of lineage B into lineage A, by drawing randomly the corresponding parameter values independently in the same prior distribution (**Table S8**). ABC posterior-parameter estimations would thus reveal possible asymmetries in gene-flows across pairs of lineages for each event separately, if identifiable from genomic data.

Finally, for each one of the 16 competing combinations of topologies and gene-flow processes, we considered three classes of gene-flow intensities by setting different boundaries of gene-flow parameters’ uniform prior-distributions (**Table S8**): no to low possible gene-flow (U[0, 0.001] for recurring processes, and U[0, 0.01] for instantaneous processes), no to moderate gene-flow (U[0,0.0125] for recurring processes, and U[0, 0.25] for instantaneous processes), or no to intense gene-flow events (U[0,0.05] for recurring processes and U[0,1] for instantaneous processes). Note that, for each three classes of gene-flow intensities, each gene-flow parameter from one lineage to another at each point or period in time was drawn independently in the above prior-distributions’ boundaries, and thus may differ across events within a scenario, and across scenarios (**Figure 4**, **Table S8**).

Altogether, this design resulted in 16×3 = 48 competing scenarios for the complex demographic history of Central and Southern African populations. Importantly, note that for any given class of gene-flow process or intensity, the eight topologies are highly nested in certain parts of the space of parameter values ^94^. In particular, despite the fact that we did not consider competing scenarios with explicitly trifurcating ancient tree-topologies, scenarios in which the parameter values for the oldest and second oldest divergence times are similar are expected to provide results highly resembling those obtained with ancient trifurcation scenarios.

All scenario-parameters are represented schematically in **Figure 4**, and their prior distributions and constraints are indicated in **Table S8**.

### Simulating genomic datasets

We performed simulations under the coalescent using *fastsimcoal2* ^128,129^, for each one of the 48 scenarios described above, separately. For each simulation, we generated a vector of parameter values randomly drawn in prior-distributions and satisfying the topological constraints as described above and in **Table S8**, with custom-made Python and Bash scripts available in the GitHub repository for this article (https://github.com/Gwennid/africa-wgs-abc). The genetic mutation model is based on the model deployed in ^97^. We simulated 100 independent loci (or “chromosomes”) with the same structure. Each such “chromosome” corresponds to a linkage block, with the following properties: the type of marker was “DNA”, the length of the loci 1 Mb, the recombination rate was 1×10^−8^ per base pair, the mutation rate 1.25 × 10^−8^ per base pair, and there was no transition bias (transition rate of 0.33).

We performed 5,000 such simulations for each 48 competing scenarios separately in order to conduct Random Forest ABC scenario-choice, hence producing 240,000 separate simulated dataset each corresponding to a single vector of parameter values randomly drawn in prior distributions (**Table S8**). Then we performed an additional 95,000 separate simulations under the best scenario obtained with RF-ABC for each 54 separate combinations of observed population samples (see below). We thus reached 100,000 simulations under each best scenario identified for each 54 observed datasets respectively, to be used for Neural Network ABC posterior parameter inference.

### Building observed genome-wide data-sets for 54 sets of five Central and Southern African populations

We considered 54 separate combinations of four eRHG, wRHG, nKS, sKS, and one eRHGn or wRHGn populations, each with five unrelated individuals (**Table 1**). We prepared a call-set of high-quality regions by applying the 1000 Genomes phase 3 accessibility mask and filtering out indels on the variant callset comprising all individuals. This resulted in a “fragmented” genome, with fragments of high-quality sites and fragments that are filtered out. We then agglutinated high-quality fragments spanning 1 Mb in total agglutinated-length. We calculated the mapped bp distance between the first and the last fragment of each 1Mb window and then randomly chose 100 windows spanning less than 1.2 mapped Mb and located on chromosomes 1 to 10, among the 824 such windows identified on chromosomes 1 to 22. We extracted these 100 independent autosomal loci of 1 Mb each for the 54 sets of 25 individuals.

In order to test whether the 100 windows used in all ABC inferences were resembling larger portions of the autosomal genome, we calculated the summary statistics used for ABC inferences (see next section) for each one of the 54 combinations of five sampled-populations each on the 100 windows and separately on the 724 other such windows identified genome-wide. Some of the summary statistics, such as the total number of biallelic SNPs in the sample, were scaled by the number of windows, to allow comparisons between the 100 and the 724 windows. We then performed Wilcoxon’s two-sided rank test and Spearman’s correlation test using the *wilcox.test* and *cor.test* functions in R.

Finally, we checked that these pruned datasets provided individual pairwise genomic differentiation patterns resembling those obtained with the entire SNP set, by computing standardized ASD matrices for each one of the 54 combinations of five populations separately. We compared these ASD matrices with the same individual subsets of the entire ASD matrix computed as above and used for MDS and NJT analyses in **Figure 2**, using Spearman *ρ* and Pearson *r* correlations tested using 1000 Mantel permutation tests (**Table S10**), with the *partial.mantel.test* function of the *ncf* package in R.

### Calculating 337 summary statistics

For each simulated genetic dataset, we computed 337 summary statistics. 41 summary statistics were computed within each five populations separately (hence 205 statistics for five populations), 132 summary statistics were computed across the five populations, for each one of the 54 combinations of five populations separately. In brief, all summary statistics were calculated with plink v1.90b4.9 ^122^, R v3.6.1 ^130^, Python v2.7.15, bash, awk, and scripts developed by ^97^, available at https://gitlab.inria.fr/ml_genetics/public/demoseq/-/tree/master, and using the software *asd* (https://github.com/szpiech/asd). The list of calculated summary statistics is provided in **Table S9**. The same statistics were computed for each one of the 54 observed data-sets separately, using the same computational tools and pipeline. Custom-made Python and Bash scripts for computations of all summary-statistics for each simulation and for the observed data are available in the GitHub repository for this article in the “fsc-simulations/code” and in the “observed-sum-stats” folders (https://github.com/Gwennid/africa-wgs-abc).

We used the complete set of 337 statistics to perform Random-Forest ABC scenario-choice, as this method is relatively fast and is unaffected by correlations among statistics ^45^. We then considered a subset of 202 statistics among the 337 for Neural Network ABC posterior-parameter estimation, in order to diminish computation time (**Table S9**).

Concerning the Site Frequency Spectrum statistics, note that we inferred the ancestral state using an alignment of three non-human apes (gorilla – reference genome gorGor5, chimpanzee – reference genome panTro6 - and orangutan – reference genome ponAbe3). We considered that the ancestral state of a site could be reliably inferred if the site had the same allele (A, C, T or G) in the three apes. Only these sites were used to calculate the proportion of homozygous ancestral positions per population and the unfolded standardized SFS.

### Prior-checking simulations’ fit to the observed data

Before conducting ABC scenario-choice and posterior parameter estimation inferences, we checked that summary-statistics calculated on the observed data fell within the range of values of summary-statistics obtained from our simulations. First, we visually verified that each observed vector of summary-statistics computed separately from 54 combinations of five populations each, clustered with the 240,000 vectors of summary-statistics obtained from simulations under the 48 competing scenarios, using two-dimensional PCA calculated with the *prcomp* function R (**Figure S5**). Second, we used the *gfit* function in R to perform goodness-of-fit 100-permutations tests between the 240,000 vectors of summary-statistics obtained from simulations under the 48 competing scenarios and, in turn, each vector of summary-statistics obtained from the 54 observed datasets respectively.

Results both showed that our simulation design could successfully mimic summary-statistics observed in our five-population sample sets, for each 54 combinations of five populations separately. Hence, ABC inferences could be confidently conducted a priori based on the simulations and observed data here considered.

### Random Forest ABC grouped scenario-choice

We conducted series of Random Forest ABC scenario-choice procedures for different groups of scenarios ^45,63^, elaborated specifically to address our different questions of interest regarding gene-flow processes, their intensities, and topological features of the history of Central and Southern African populations. RF-ABC scenario-choice has proven to be performing efficiently and satisfactorily with a significantly lower number of simulations compared to any other ABC scenario-choice procedure. Moreover, RF-ABC scenario choice is insensitive to correlations among summary-statistics ^45,63^.

Each RF-ABC scenario-choice procedure presented below was conducted using the *predict.abcrf* and the *abcrf* functions with the “group” option of the *abcrf* package in R ^45^, with 1000 decision trees to train the algorithm after checking that error rates were appropriately low with the *err.abcrf* function (**Figure S6**), separately for each 54 combinations of five sampled populations. Posterior probabilities that the selected scenario, for each analysis separately, is the true scenario, conditional on the classification error a priori is conducted with the abcrf function as explicated in Algorithm 3 in Section 2.4 of ^64^.

We conducted “out-of-bag” error cross-validation procedures ^64^, considering in turn each one of the simulations as pseudo-observed data and all remaining simulations to train the algorithm, for each RF-ABC analysis separately (**Figure S7, Figure S8, Figure S9, Figure S10**). While not necessarily predicting the outcome of the scenario-choice for the observed data, these out-of-bag procedures provide us with a sense of the discriminatory power, *a priori*, of RF-ABC for our set of competing-scenarios, as well as empirical levels of nestedness among scenarios or groups of scenarios ^94,95^. Finally, we estimated the relative contribution of each one of the 337 summary-statistics to each one of the *a priori* RF-ABC scenario-choice procedure, i.e. the relative importance of each statistics to discriminate among competing scenarios for the entire simulation space using out-of-bag cross-validations on all simulations only without observed data, using the “n.var” option in the *abcrf* plotting function. For simplicity and readability, we provide results for the 50 most important such summary-statistics for each RF-ABC scenario-choice procedure performed in **Figure S11** to **Figure S20**.

#### Instantaneous or recurring gene-flows in the history of Central and Southern African populations?

The 5,000 simulations performed under each one of the 48 competing scenarios were first gathered into two groups of 24 scenarios each (**Figure 4**), corresponding to either instantaneous asymmetric gene-flow processes or to recurring constant asymmetric gene-flow between each pair of lineages, respectively. Both groups thus contained all simulations from the eight competing topologies and all three sets of possible gene-flow intensities (low, moderate, or high, see above), and only differed in the process of gene-flow itself. For each 54 combinations of five sampled populations separately, we performed such RF-ABC scenario-choice to determine the best group of scenarios (**Figure 5A, Table S12**).

#### Low, moderate, or high gene-flow intensities?

For each 54 combinations of five sampled populations separately, we performed RF-ABC scenario-choice in order to determine which class of intensities of gene-flows best explained the data, everything else (topology and gene-flow process) being-equal. We thus considered three groups of scenarios in our RF-ABC scenario-choice, each corresponding to “low”, “moderate”, or possibly “high” intensities, and each grouping the 16 scenarios corresponding to eight topologies and instantaneous or recurring gene-flow processes indiscriminately (**Figure 5B, Table S12**).

Then, we conducted, for each 54 population combinations separately, the same RF-ABC scenario-choice procedure to disentangle groups of gene-flow intensities “all topologies being equal”, considering here only the 24 scenarios from the best group among instantaneous or recurring gene-flow processes obtained from the above scenario-choice procedure (**Figure S8, Table S12**).

Finally, we performed, for each 54 combinations of five populations separately, RF-ABC scenario choice for the 48 competing scenarios grouped in six different groups (encompassing eight scenarios each), combining instantaneous or recurring processes with the three classes of intensities respectively (**Figure 5C, Table S12**), thus “intersecting” the two corresponding group analyses (see above).

#### Which ancestral Central or Southern African lineage diverged first?

The eight topologies considered in the 48 competing scenarios can also be grouped according to their ancient topologies, by considering which ancestral lineage diverged first from all others at the oldest divergence event in the tree (**Figure 4**). The ancient lineage which separated first from the two others can either be the lineage ancestral to Northern and Southern Khoe-San populations (AKS, scenario topologies 1a, 1b, 1c in **Figure 4**), the lineage ancestral to Eastern and Western Rainforest Hunter-Gatherer populations (ARHG, scenario topologies 2a, 2b, 2c in **Figure 4**), or the lineage leading to Rainforest Hunter-Gatherer Neighboring populations (RHGn, scenario topologies 3a, 3b in **Figure 4**).

For each 54 combinations of five populations separately, we conducted RF-ABC scenario-choice across these three groups of scenarios, randomly drawing 2/3 of the 5,000 simulations per scenario for the scenarios 1a, 1b, and 1c, and for the scenarios 2a, 2b, and 2c, respectively, and kept all simulations for the scenarios 3a and 3b, in order to even the total number of simulations in competition among the three groups of scenarios. We thus performed RF-ABC scenario-choice across the three groups of topologies “all gene-flow processes and intensities being equal” (**Figure 5D, Table S12**).

Then we restricted the RF-ABC scenario-choice of the three-competing groups of topologies only for the 24 scenarios from the best instantaneous or recurring gene-flow processes obtained above, all gene-flow intensities being equal (**Figure 5E, Table S12**).

#### Northern and Southern Khoe-San populations diverged before or after the divergence between Eastern and Western RHG?

The eight topologies considered in the 48 competing scenarios can alternatively be grouped according to the relative order of divergence-time between the divergence event among Northern and Southern KS populations, and that among Eastern and Western RHG populations; an important question to understand the relative duration of separate evolution of each group of populations.

To address this specific question we thus conducted, for the 54 combinations of five populations separately, RF-ABC scenario-choice procedures by grouping the 48 competing scenarios in two separate groups of 24 scenarios each. One group corresponded to KS populations diverging from one-another before the RHG divergence (scenario topologies 1b, 2b, 3b, and 1c in **Figure 4**), and the other group where RHG diverged from one another before the KS divergence (scenario topologies 1a, 2a, 3a, and 2c in **Figure 4**), instantaneous or recurring gene-flow and all “low”, “moderate”, or “high” gene-flow intensities being equal among the two groups (**Figure 5F, Table S12**). Last, we conducted the same RF-ABC scenario-choice procedure between two groups of four topologies each, but restricted to the 24 best scenarios among the instantaneous or recurring gene-flow processes as determined from above procedures (**Figure 5G, Table S12**).

Finally, we conducted RF-ABC scenario choice procedures considering, respectively, all 48 competing scenarios separately (**Figure S9, Table S12**), and all 24 best scenarios among the instantaneous or recurring gene-flow processes as determined above (**Figure S10, Table S12**).

### Neural Network ABC posterior-parameter estimation

We intersected all the RF-ABC scenario-choice results considering the different groups of scenarios or all scenarios independently, as described above, in order to determine, which single scenario was explaining data most accurately in the majority of the 54 combinations of five Central and Southern African populations. We found that the Scenario i1-1b conservatively produced simulations whose observed summary-statistics most resembled those obtained in 25 out of the 54 possible population combinations here tested (see **Results**, the detailed list of 25 population combinations is provided in **Table S13**). The second most often best-scenario, Scenario i1-3b, only succeeded for four population combinations among the 54 tested.

Based on this *a posteriori* Scenario i1-1b, we performed an additional 95,000 simulations considering the same prior distributions and constraints among parameters, in order to obtain 100,000 simulations for further ABC posterior parameter inferences, separately for each one of the 25 population combinations for which it was identified confidently as the winner. As it remains difficult to estimate jointly the posterior distribution of all parameters of the complex scenarios here explored with RF-ABC ^62^, we instead conducted Neural Network ABC joint posterior parameter inferences using the *abc* package in R ^43,44^, following best-practices previously bench-marked for highly complex demographic scenarios ^95,97^.

There are no evident criteria to choose *a priori* the best tolerance level and numbers of neurons in the neural network’s hidden layer for parameterizing the NN-ABC posterior-parameter estimation procedure ^97,104^. As the total number of parameters in the best Scenario i1-1b was large (43, **Figure 4** and **Table S8**), and as the number of summary-statistics considered was also large (202, **Table S9**), we chose to conduct the 25 NN-ABC posterior-parameter inferences for the 25 combinations of sampled populations considering, in-turn, 7, 14, 21, 28, 35, 42, or 43 neurons in the hidden layer (43 being the assumed number of dimensions equal to the number of parameters for this scenario ^43,97^), and a tolerance level of 0.01, thus considering the 1,000 simulations closest to each observed dataset out of the 100,000 performed, respectively. We found *a posteriori* that, considering 43 neurons in the hidden layer and a maximum allowable number of weights MaxNWts = 12100 provided overall posterior parameter distributions departing from their priors, and in particular for divergence-times and gene-flow-times parameters which we were highly interested in, and therefore decided to provide all posterior parameter distributions’ results using this parameterization (tolerance = 0.01, number of hidden neurons = 43, MaxNWts = 12100) for the training of each one of the neural networks.

We thus performed 25 separate NN-ABC joint parameter posterior inferences, using the “neuralnet” method option in the function *abc* of the *abc* package in R ^44^, with logit-transformed (“logit” transformation option), an “epanechnikov” kernel, parameter values within parameter-priors’ boundaries, a tolerance level of 0.01, 43 neurons in the hidden layer, and maximum allowable number of weights MaxNWts = 12100. Adjusted posterior parameter distributions obtained with this method were then plotted for each parameter separately using a gaussian kernel truncated at the boundaries of the parameter prior-distributions. We then estimated the mode, median, mean, and 50% and 90% Credibility Intervals of these distributions *a posteriori*. We provide all posterior-parameter distributions together with their priors in figure-format (**Figure 6**, **Figure 7**, **Figure S21, Figure S22**), and in table-format (**Table S17**, **Table S14**).

### Neural Network ABC posterior-parameter estimation cross-validation errors

For each one of the 25 combinations of sampled population separately, we randomly chose 50 simulations (5%) out of the 1000 simulations closest to each observed dataset as determined above. We then conducted NN-ABC joint posterior parameter inference with the same hyper-parameters as above and considering, for each chosen simulation considered as pseudo-observed, the other 99,999 simulations in the reference table, for each one of these 50 pseudo-observed simulations and for each 25 combinations of sampled populations separately (1250 separate NN-ABC joint posterior parameter inferences in total). We then calculated, for each 43 parameter and each 25 populations’ combination, the cross-validation posterior parameter estimation error as 1. the mean-squared error of the median NN-ABC posterior estimate (*θ̂*_*i*_) scaled by the true-parameter’s (*θ*_*i*_) variance (**Table S16**), 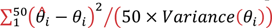 ^44^, and 2. simple average estimation bias of the median NN-ABC posterior estimate 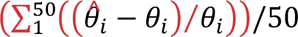 (**Table S16**).

### Neural Network ABC posterior-parameter estimation posterior checks

We performed 25 separate simulations of Scenario i1-1b each using the same mode point-estimate parameter-values obtained combining (concatenating) the results obtained separately for the 25 different observed combinations of five Central and Southern African populations each and presented in results (**Figure 6**, **Figure 8**, **Figure S21**, **Figure S22**, **Table S17** and synthesized schematically in **Figure 8**). We also performed 25 separate simulations of the same scenario using the mode point-estimate parameter-values obtained separately for each 25 combinations of five populations and provided in **Table S14**.

First, we aimed to determine *a posteriori* whether our synthetic scenario that best explains genomic patterns in 25 different observed combinations of five Central and Southern African populations each, provided summary-statistics reasonably resembling that of the 25 observed datasets. We also aimed to determine a posteriori whether each best scenario inferred for each 25 combinations of five sampled populations provided summary-statistics reasonably resembling those of the synthetic scenario and those from the observed sets of summary-statistics. Therefore, we performed PCA using the *prcomp* function in R, computed on the 202 summary-statistics used for our NN-ABC posterior-parameter inferences (**Table S9**), calculated on each one of the 100,000 simulations used as a reference table for NN-ABC posterior-parameter inferences, on the 25 pseudo-observed simulations under the best synthetic scenario, on the 25 pseudo-observed simulations under each best scenario for each one of the 25 combinations of five sampled populations separately (using the mode point estimates in **Table S14**), and the 25 observed datasets similarly as above (**Figure S48**). We normalized (min-max) each summary-statistics calculated separately for each one of the 25 combinations of five sampled population, together with the corresponding statistics computed on the 25 pseudo-observed simulations performed under the synthetic scenario (**Table S17**). For each summary-statistics and each 25 observed combinations of population, we then calculated the difference between the normalized observed value and the average of the normalized value among the 25 pseudo-observed simulations. We finally averaged the absolute value of the obtained difference across the 25 observed combination of populations and provide the 50 most differentiated and 10 least differentiated summary-statistics in **Figure S49**.

Second, beyond the NN-ABC posterior parameter inferred CI and cross-validation error results here obtained, we aimed to empirically estimate whether the variance of posterior parameter estimates obtained among the 25 different combinations of five Central and Southern African populations originated from variation in performances across NN-ABC inferences rather than from different demographic evolutionary histories of the different population combinations considered.

To do so, we conducted, similarly as above for the 25 observed combinations of five populations, NN-ABC joint posterior-parameter inferences for each one of the 25 pseudo-observed simulated datasets under our proposed synthetic scenario (**Figure 6**, **Figure 7**, **Figure S21**, **Figure S22**, **Table S17** and synthesized schematically in **Figure 8** and **Figure S23** to **Figure S47**). We plotted obtained posterior-parameter distributions for each pseudo-observed simulations separately as well as for combined (concatenated) results similarly as for the observed datasets (**Figure S50**, **Figure S51**, **Figure S52**, **Figure S53**). For each parameter separately, we then performed Fligner and F tests of homogeneity of variance, using, respectively, the *fligner.test* and *var.test* functions in R between combined (concatenated) posterior-parameter distributions obtained with 25 pseudo-observed simulations and those obtained previously with 25 observed combinations of five populations (**Table S18**).

## Acknowledgments

We are grateful to all subjects who participated in this research. The computations were performed at the Swedish National Infrastructure for Computing (SNIC-UPPMAX). Sequencing was performed by the SNP&SEQ Technology Platform in Uppsala. We thank the platform Paléogénomique et Génétique Moléculaire (P2GM) of the MNHN at the Musée de l’Homme in Paris for helping to prepare, in part, the Central African samples for sequencing. We thank the Working Group of Indigenous Minorities in Southern Africa (WIMSA) and the South African San Council for their support and facilitating fieldwork. The project was approved by the South African San Council. We thank Rémi Tournebize, four other anonymous reviewers, and the editors for their help improving this work.

## Authors’ contributions

GB – Designed the study, conducted data curation and quality controls, conducted statistical descriptions, conducted simulations and ABC inferences, interpreted the results, wrote the original draft.

PS – Designed the study, conducted statistical descriptions, interpreted the results, participated in writing and editing.

PLZ – Conducted data curation and quality controls, conducted statistical descriptions, participated in editing.

RL –Conducted simulations and ABC inferences, participated in writing and editing.

AF – Conducted fieldwork and provided samples, participated in editing.

AES – Conducted fieldwork and provided samples, participated in editing.

BSH – Conducted fieldwork and provided samples, participated in editing.

LBB – Conducted fieldwork and provided samples, participated in editing.

GHP – Conducted fieldwork and provided samples, participated in editing.

HS – Conducted fieldwork and provided samples, participated in editing.

EH – Acquired funding, interpreted the results, participated in editing.

CMS – Supervised the project, designed the study, acquired funding, conducted fieldwork and provided samples, interpreted the results, participated in writing and editing.

MJ – Supervised the project, designed the study, acquired funding, interpreted the results, participated in writing and editing.

PV – Supervised the project, designed the study, acquired funding, conducted fieldwork and provided samples, conducted statistical descriptions, conducted simulations and ABC inferences, interpreted the results, wrote the original draft.

## GitHub repositories for material availability

https://github.com/Gwennid/africa-wgs-descriptive https://github.com/Gwennid/africa-wgs-abc

## Data availability statement

All raw whole-genome sequence FASTQ data original to this study are made available on the European Genome-Phenome Archive (https://ega-archive.org/), provided that requests conform to ethical requirements and informed consents provided by donors, as detailed in the corresponding Data Access Policies associated with each data repository. Data access requests are evaluated by the associated Data Access Committees. EGA accession numbers: EGAD50000001559 and EGAD50000001560.

## Ethical statement

DNA samples from individuals were collected with the subjects’ informed consent following the Declaration of Helsinki guidelines and approved by the following Ethics committees: Institut de Recherche pour le Développement (IRD, France); Cameroon Ministry of Scientific Research and Technology; Central African Republic Ministère de l’Enseignement Supérieur et de la Recherche Scientifique; France Ministère de l’Enseignement Supérieur, de la Recherche et de l’Innovation (Comité de Protection des Personnes DC-2016-2740); French Commission Nationale Informatique et Liberté CNIL (declaration n°1972648); Uganda National Council for Science and Technology; Institutional Review Board of the University of Chicago Biological Sciences Division and the Pritzker School of Medicine University of Chicago Hospitals (protocol number 16986A); Committee for the Protection of Human Subjects of the Trustees of Dartmouth College and the Dartmouth-Hitchcock Medical Center (protocol number 22410); Human Research Ethics Committee (Medical) at the University of the Witwatersrand, Johannesburg (Protocol Numbers: M980553, M050902, M090576, M10270, M180654); Working Group of Indigenous Minorities in Southern Africa (WIMSA); South African San Council (SASC); and the Swedish Ethical Review Authority (Dnr 2019-05174).

## Supplementary Material

## Supplementary Note

### Additional description of results on ROH patterns

In general (**Figure S1C**), individuals from Eastern RHG populations (Nsua, Mbuti, Ba.Twa), exhibit the longest total proportion of their autosomal genome in ROH across Sub-Saharan African populations, for all ROH length-classes (Class 0.2-0.5 median total length: 71.28 Mbp; Class 0.5-1: 35.97 Mbp; Class 1-2: 15.79 Mbp; Class 2-4: 9.72 Mbp), with the exception of #Khomani individuals from Southern Africa, and that of Coloured individuals for short ROH only (Class 0.2-0.5 median total length= 98.15 Mbp and 108.18 Mbp, respectively). Interestingly, we find that Western RHG populations (Ba.Kola, Baka, Biaka, Bi.Aka_Mbati) have significantly less total length of ROH of short and medium classes than Eastern RHG (Class 0.2-0.5: Wilcoxon one-sided test p-value= 4.8×10^-8^; Class 0.5-1: p-value=0.0055; Class 1-2: p-value=0.1284). Furthermore, we find significantly less short ROH (Class 0.2-0.5: Wilcoxon one-sided test p-value=1.4×10^-5^), and significantly more long ROH (Class 1-2: p-value=2.4×10^-7^), in Western RHG than in all RHG neighboring populations across the Congo Basin except for the Ba.Kiga RHGn from Uganda. Finally, we find that all Northern and Southern Khoe-San populations have relatively similar total lengths of short ROH compared to that of RHG neighbors (Class 0.2-0.5 Wilcoxon one-sided p-value=0.01675), with the notable exception of the !Xun, who have among the smallest proportion of their genomes in short ROH, worldwide (Class 0.2-0.5 median total length=42.59 Mb). Nevertheless, we find significantly more ROH of longer class in KS populations than in RHG neighbors (Class 1-2: Wilcoxon one-sided p-value=3.7×10^-5^), a relative pattern similar to what was observed for Western RHG. Finally, note that total ROH lengths significantly correlated negatively (Spearman rho=-0.5912, p-value=5.243×10^-5^) with unbiased estimates of heterozygosities (**Figure S2**).

## Supplementary Figures and Tables

**Figure S1:**
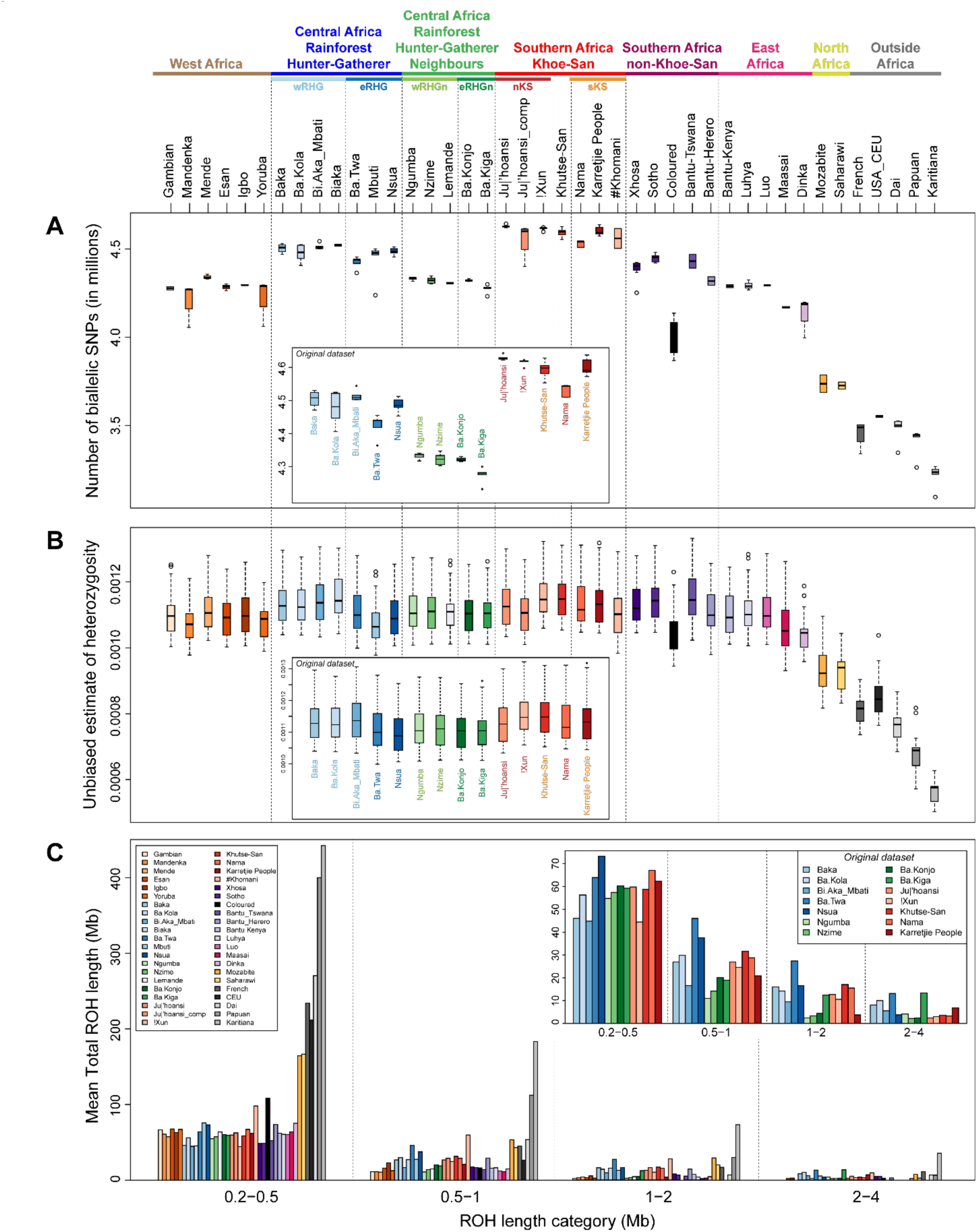
Whole autosomal genome bi-allelic SNPs counts, unbiased heterozygosities, and distributions of Runs of Homozygosity by bins of length, in 37 African and 5 non-African populations. (**A**) Numbers of bi-allelic SNPs across individuals within populations of more than one individual, compared to the reference sequence of the human genome GCRh38 and to previously reported variants in dbSNP 156. Detailed variant-counts are provided in **Table S2 and Table S3**. Note that we find a large variability in the mean number of biallelic SNPs across populations. In particular, we find substantially more biallelic SNPs within African populations and more variation of the mean number of SNPs across African populations (from mean=3,726,018; SD=27,793 across 2 individuals in the Saharawi from Western Sahara, to mean=4,628,957; SD=8,139 across 5 individuals in the Ju|’hoansi from Namibia), than within and among non-African populations (from mean=3,214,086; SD=69,241 across 5 individuals in the Karitiana from Brazil, to mean=3,551,792; SD=7,819 across 2 individuals from the USA CEU population). (**B**) Unbiased estimates of multi-locus heterozygosities ^47^, averaged across all variable (bi-allelic SNPs only) and non-variable autosomal sites with no-missing genotype within each population of more than one individual, corrected then for haploid population sample sizes. (**C**) Mean total ROH lengths in four bins of length categories for each population of more than one individual, separately. For each panel, we present a box zooming specifically on the results obtained for the original dataset of 73 unrelated individuals whole-genome sequenced from Central and Southern Africa. See **Material and Methods** for filtering and calculation details and the software packages used. For (**A**) and (**B**), box-plots indicate the population median in between the first and third quartile of the box-limits, whiskers extending to data points no more than 1.5 times the interquartile range of the distribution, and empty circles for all more extreme points beyond this limit, if any. Population geographical location, categorization, and descriptions are provided in **Figure 1, Table 1, and Table S1**.

**Figure S2:**
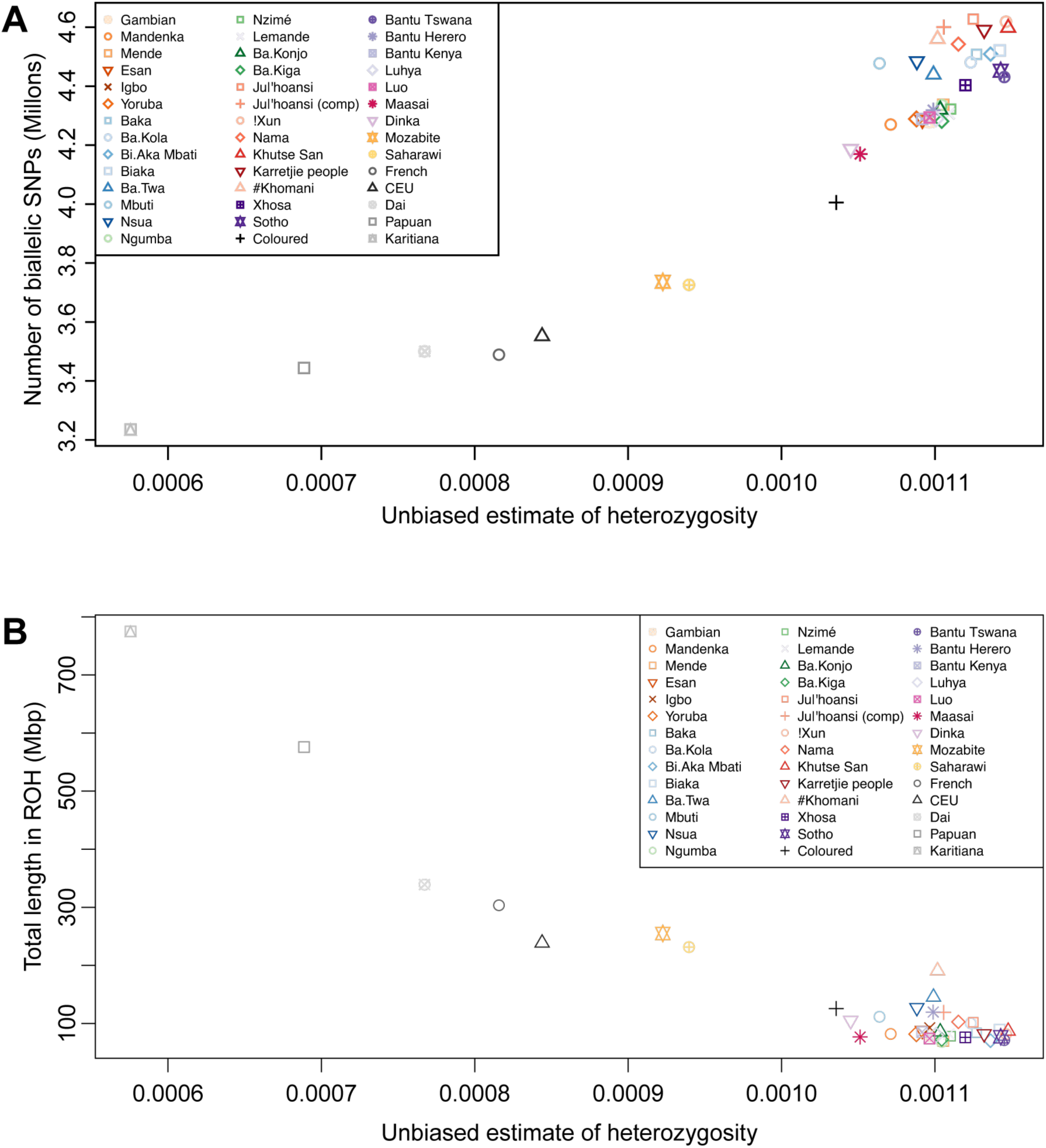
Correlations between genome-wide autosomal number of biallelic SNPs, unbiased estimates of heterozygosities and total ROH lengths at the world-wide scale. Median values across individuals within populations reported in **Figure S1** panels A-C are used here in panel **A**) Unbiased estimates of heterozygosity vs. total number of autosomal biallelic SNPs, and in panel **B**) Unbiased estimates of heterozygosity vs. total autosomal length found in ROH. As expected, we found a significant positive correlation in Panel **A** (Spearman rho=0.9490, p-value<2.2×10^-16^), and a significant negative correlation in Panel **B** (Spearman rho=-0.59128, p-value=5.243×10^-^_5_).

**Figure S3:**
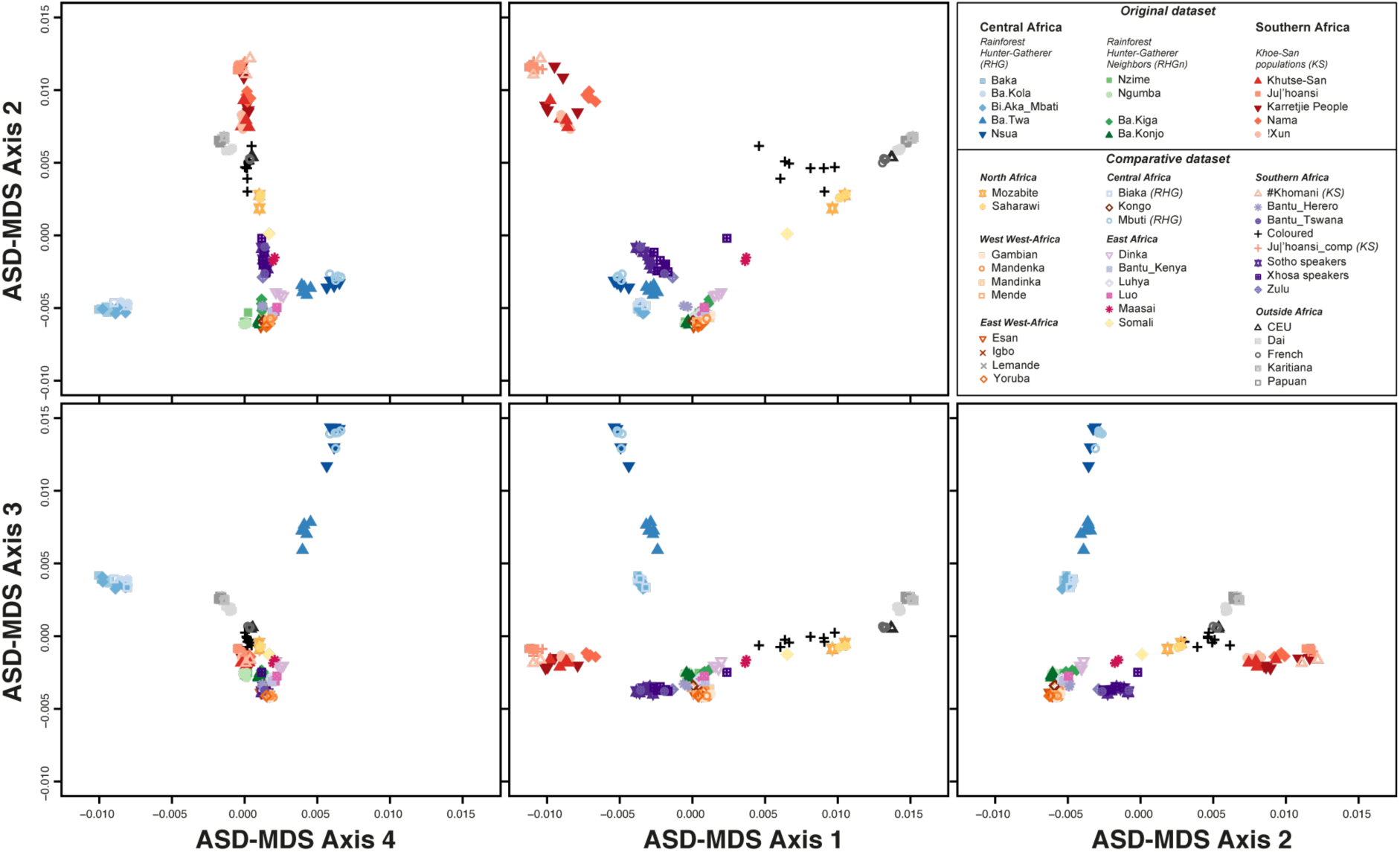
Individual pairwise genetic differentiation patterns at the world-wide scale. Multi-Dimensional scaling representation of genome-wide Allele Sharing Dissimilarities ^51^ among pairs of individuals at worldwide scale. We considered the 14,182,615 genome-wide autosomal SNPs pruned for low LD (r^2^ threshold 0.1) to calculate ASD between all pairs of individuals. This figure complements the NJ representation provided in **Figure 2** panel A.

**Figure S4:**
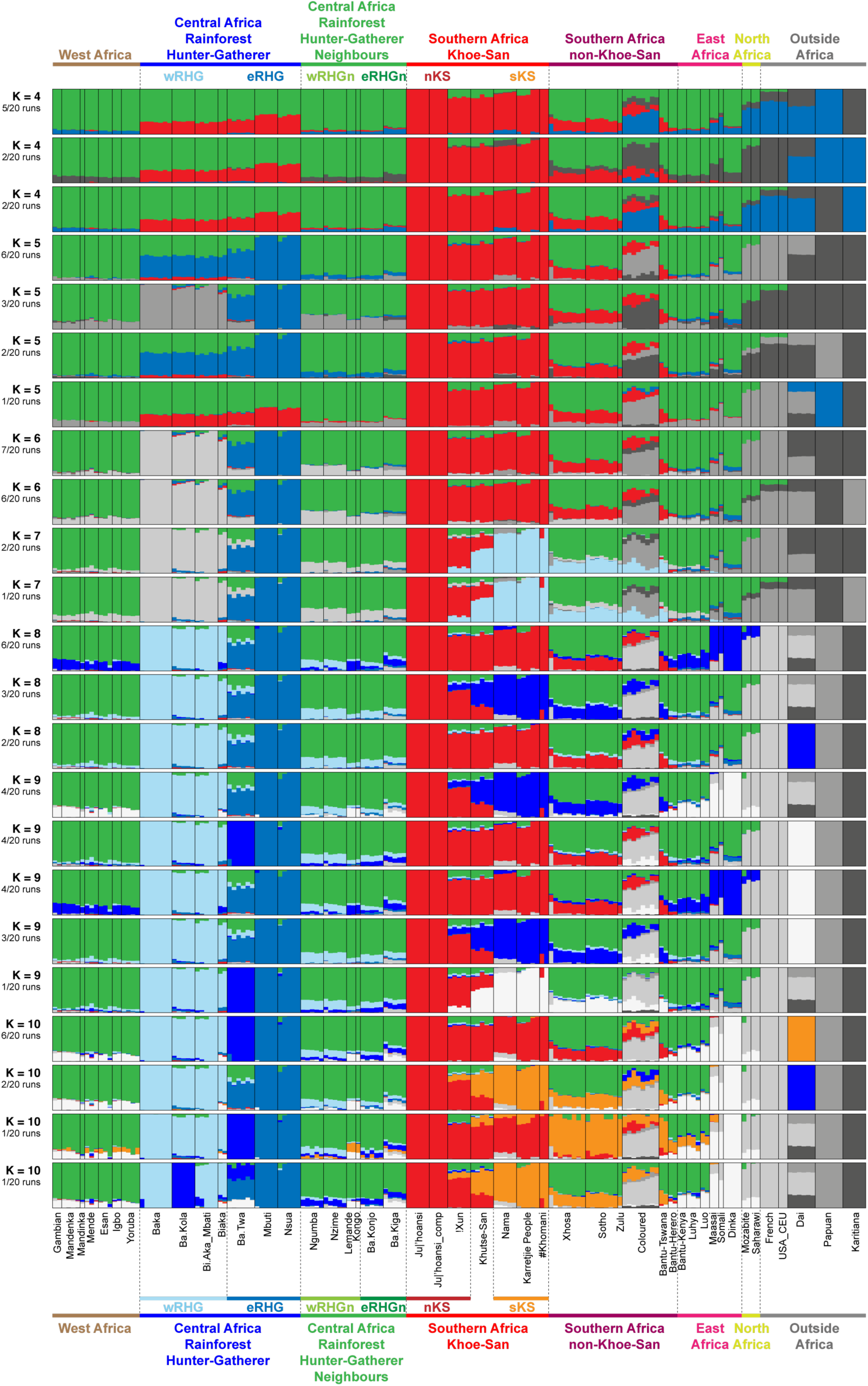
Alternative ADMIXTURE modal results for K=4 to 10. Modal results each containing the highest number of runs among the 20 independent runs performed for each value of K are provided in main **Figure 3**. All alternative modal results are provided here. See **Material and Methods** for details.

**Figure S5:**
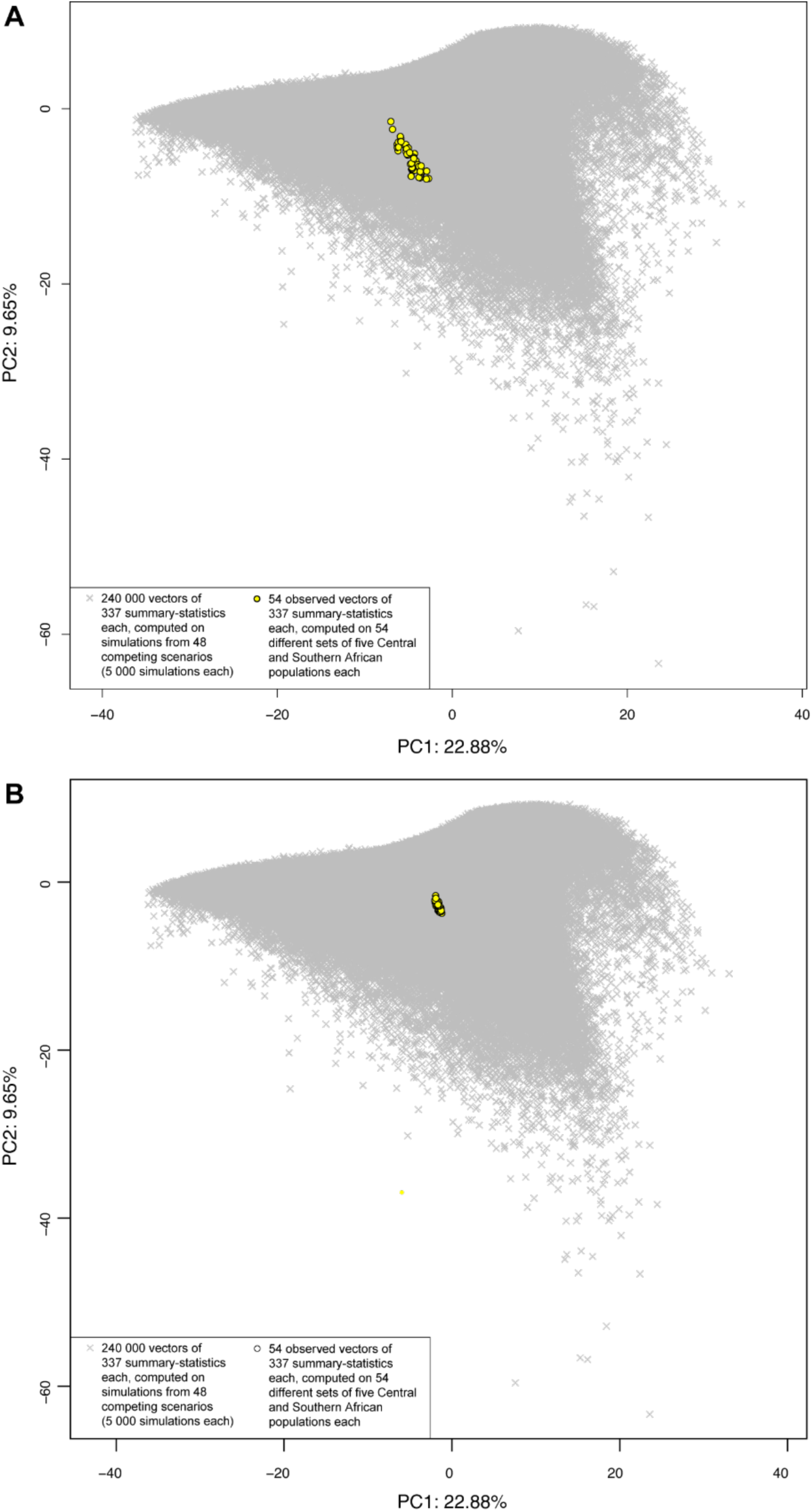
PCA for ABC prior-checking of simulations fit to the observed data. We calculated 337 summary-statistics for each one of the 240 000 simulated data sets, 5000 under each one of the 48 competing scenarios (**Figure 4** and **Material and Methods**), which we projected on the first two axes of a principal component analysis. Each vector (corresponding thus to a single simulation) is represented by a gray cross. We separately computed the same 337 summary-statistics, separately for each one of the 54 sets of five observed Central and Southern African populations included in our analyses. A) Each observed vector is then projected, in turn, on the PCA obtained from simulations only. Each such observed vector is represented as a single yellow dot. B) Alternatively, a PCA is performed on all simulated and observed vectors of summary statistics together. Each observed vector is also represented as a single yellow dot in panel B. We can see that, in both cases, the 54 observed sets of populations each fall well within the space of simulated dataset, thus allowing us a priori to conduct machine-learning ABC scenario choice and posterior parameter estimation procedures.

**Figure S6:**
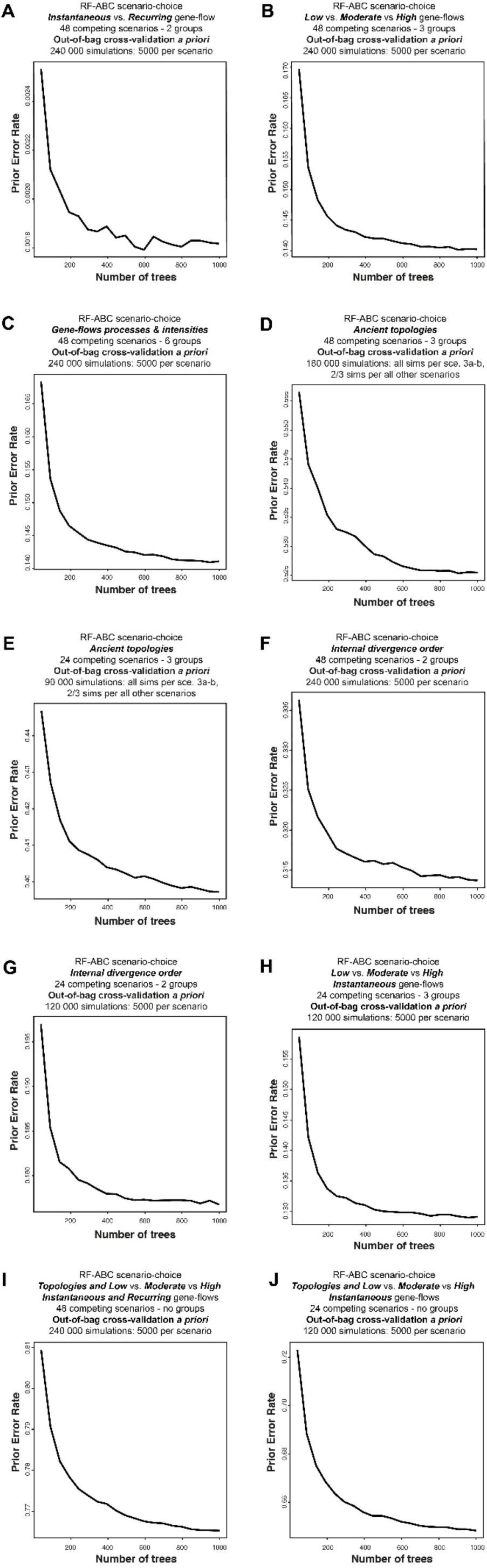
Cross-validation prior error of Random-Forest ABC scenario-choice procedures as a function of the number of trees in the forest. For panels A to G, out-of-bag cross-validation prior error results for each RF-ABC scenario choice analysis are shown in **Figure S7 panels A to G** respectively. For panel H, the corresponding out-of-bag cross-validation prior error results are shown in **Figure S8 panel B.** For panel I, the corresponding out-of-bag cross-validation prior error results are shown in **Figure S9 panel B.** For panel J, the corresponding out-of-bag cross-validation prior error results are shown in **Figure S10 panel B.**

**Figure S7:**
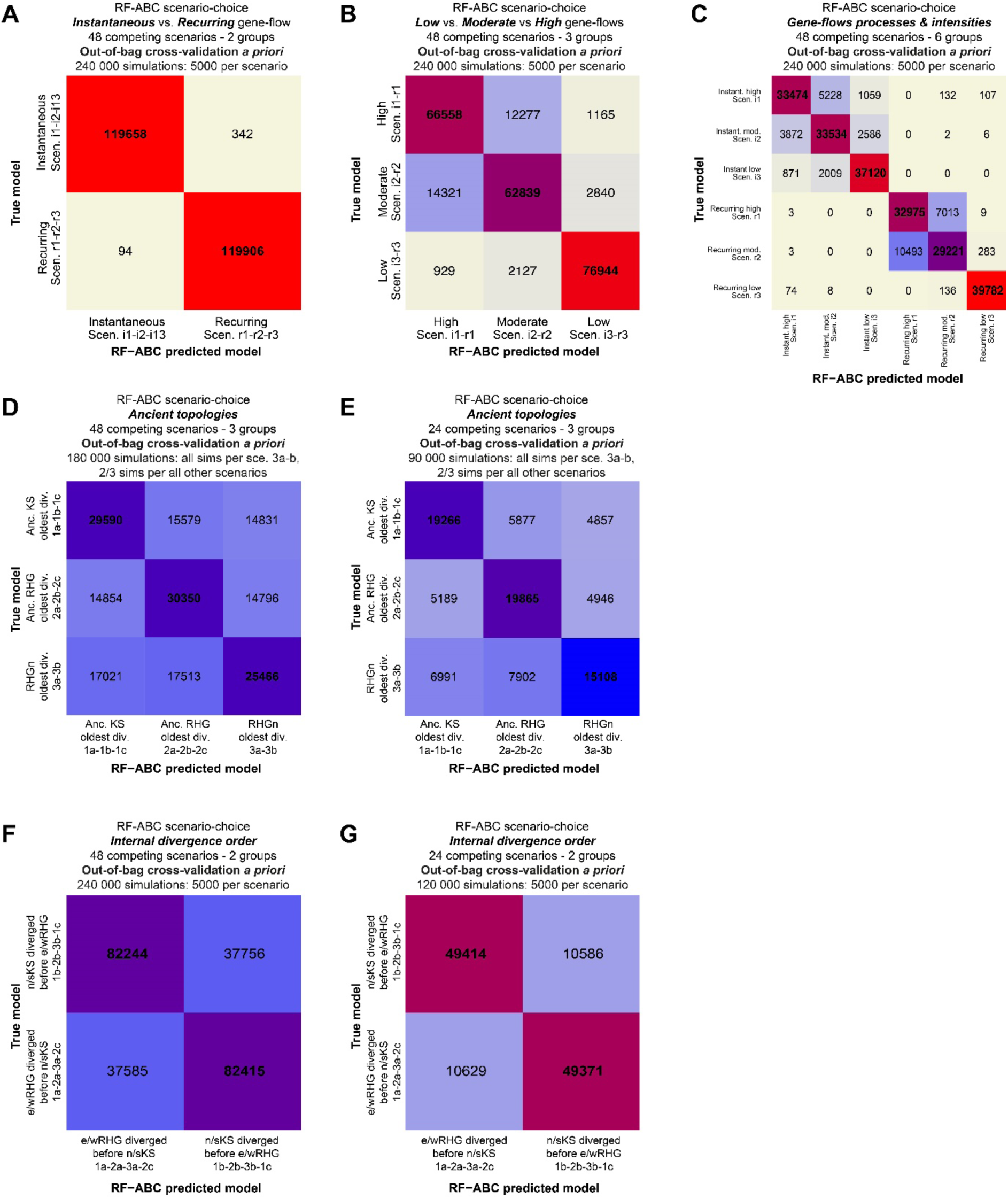
Out-of-bag cross-validation errors for Random-Forest ABC scenario-choices among 48 competing scenarios. All RF-ABC cross-validation analyses are conducted *a priori*, without using any observed data, by considering, in-turn, each simulation as observed data while all remaining simulations are used to train the Random Forest. Each panel provides the *a priori* cross-validation errors for the corresponding RF-ABC scenario-choice analysis detailed in each panel in **Figure 5**. While these cross-validation errors do not predict the specific outcome of RF-ABC prediction for the observed data, they provide levels of scenarios and groups of scenarios nestedness *a priori* for the entire space of parameters used for simulations. (**A**) shows that intermediate and recurring gene-flow processes are, all topologies and gene-flow intensities “being equal”, clearly differentiable by RF-ABC and little nested. (**B**) and (**C**) show that different classes of gene-flow intensities are also relatively clearly identifiable, albeit more nested, as expected by design since the “high gene-flow” class comprises parameter values of the moderate and the low intensity classes of gene-flows (**Material and Methods**, **Table S8**). (**D**) and (**E**) show that ancient tree topologies are relatively well distinguishable *a priori* all gene-flow processes “being equal”, despite some amounts of errors due to the expected nestedness in the space of parameter values where ancient divergence times are resembling and thus undistinguishable. (**F**) and (**G**) show that the chronological order between the divergence-time of Northern and Southern KS lineages, and that of Western and Eastern RHG lineages, are also relatively well distinguishable a priori with RF-ABC scenario-choice, albeit with an increased cross-validation error expected due to high levels of nestedness in the spaces of parameter-values where both separate events may occur at similar time.

**Figure S8:**
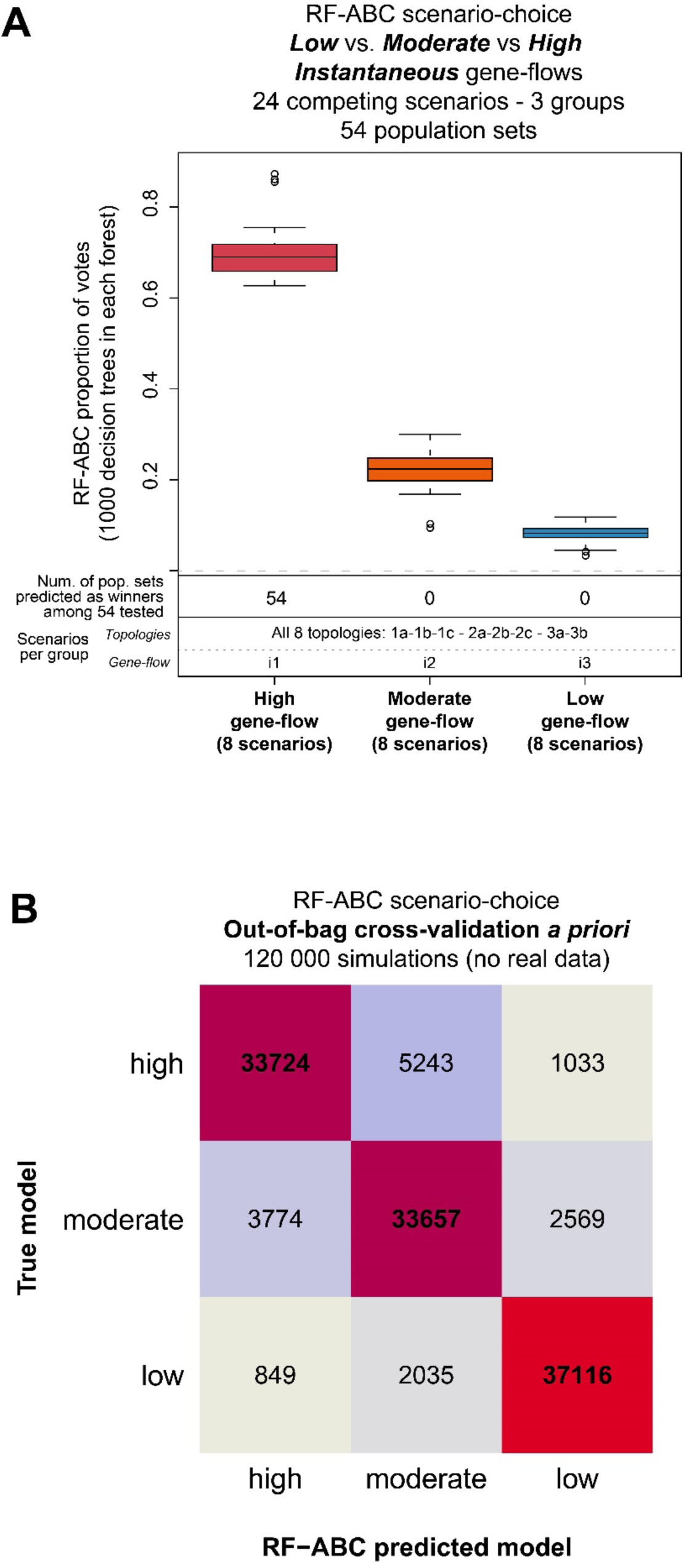
RF-ABC scenario choice among 24 competing scenarios with instantaneous asymmetric gene-flow processes, for three groups of gene-flow intensities, all topologies being equal. (**A**) corresponds to the scenario-choice results for the same test as in **Figure 5C**, restricted to the 24 competing scenarios considering instantaneous gene-flow processes only, all scenario topologies “being equal”. Description of the x and y axis legends are provided in **Figure 5**. Posterior probabilities of choosing the correct scenarios for each 54 sets of population combinations ranged from 0.63935 to 0.87000, compared to a prior probability of 1/3 in this particular test. (**B**) corresponds to the RF-ABC cross-validation prior error for the scenario-choice procedure conducted in (**A**). Errors were obtained without considering observed data and using, instead, all simulations in turn as pseudo-observed data and the remaining simulations to train the RF.

**Figure S9:**
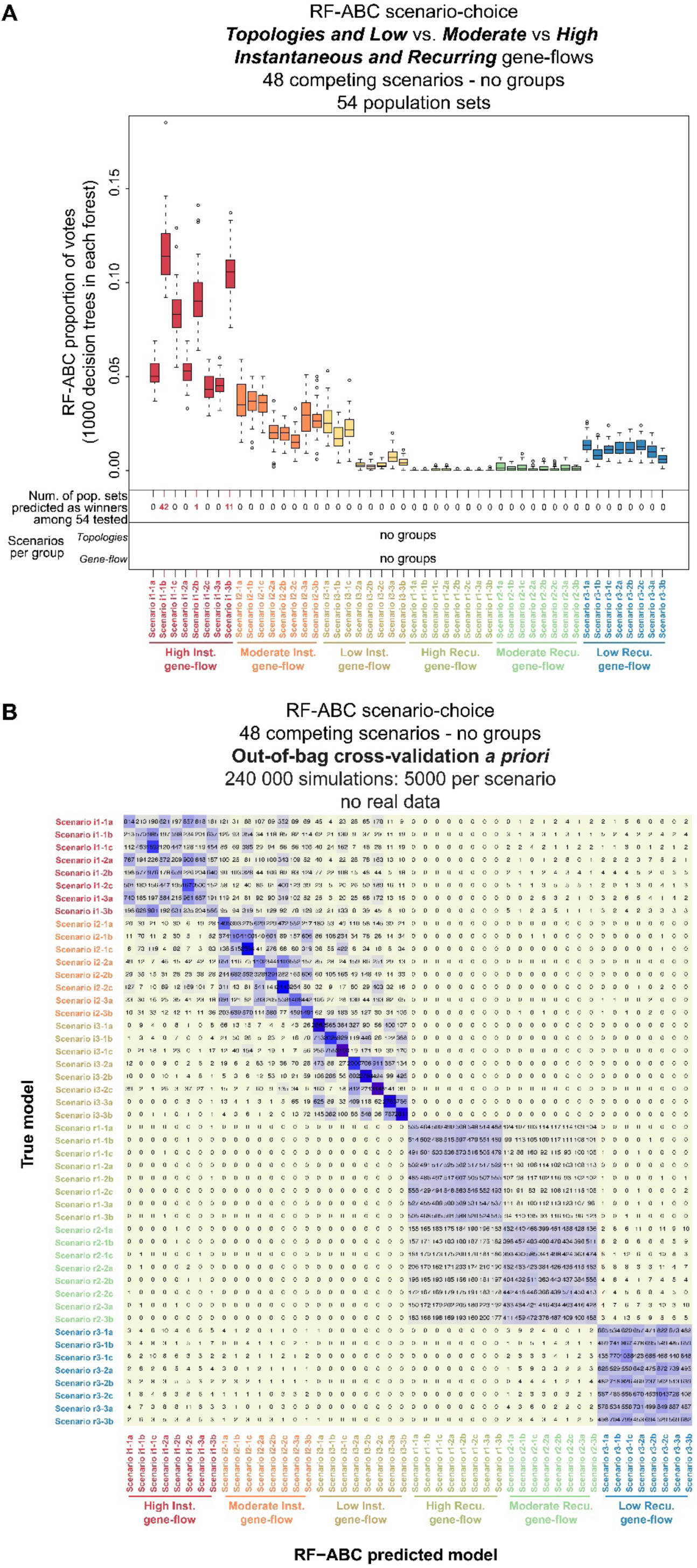
RF-ABC scenario choice among 48 competing scenarios without groups. (**A**) corresponds to the RF-ABC scenario-choice results obtained when considering all 48 scenarios (5,000 simulations per scenario), as separate competitors without grouping them, for each 54 combinations of five sampled-populations separately. Description of the x and y axis legends are provided in **Figure 5**. Posterior probabilities of choosing the correct scenarios for each 54 sets of population combinations ranged from 0.20553 to 0.28588, compared to a prior probability of 1/48 in this particular test. (**B**) corresponds to the RF-ABC cross-validation prior error for the scenario-choice procedure conducted in (**A**). Errors were obtained without considering observed data and using, instead, all simulations in turn as pseudo-observed data and the remaining simulations to train the RF. Note that, in (**B**), whichever the intensity of the gene-flow, when considering only recurring gene-flow processes, topologies are harder to distinguish from one another a priori. Conversely, such difficulty to distinguish a priori among topologies remains overall limited for instantaneous gene-flow processes, and errors increase only when considering the possibility of intense instantaneous gene-flows. However, as mentioned throughout the article, the capacity to distinguish a priori among scenarios in the entire space of summary-statistics values does not predict the power to predict a best scenario in the specific space occupied by observed data ^45^.

**Figure S10:**
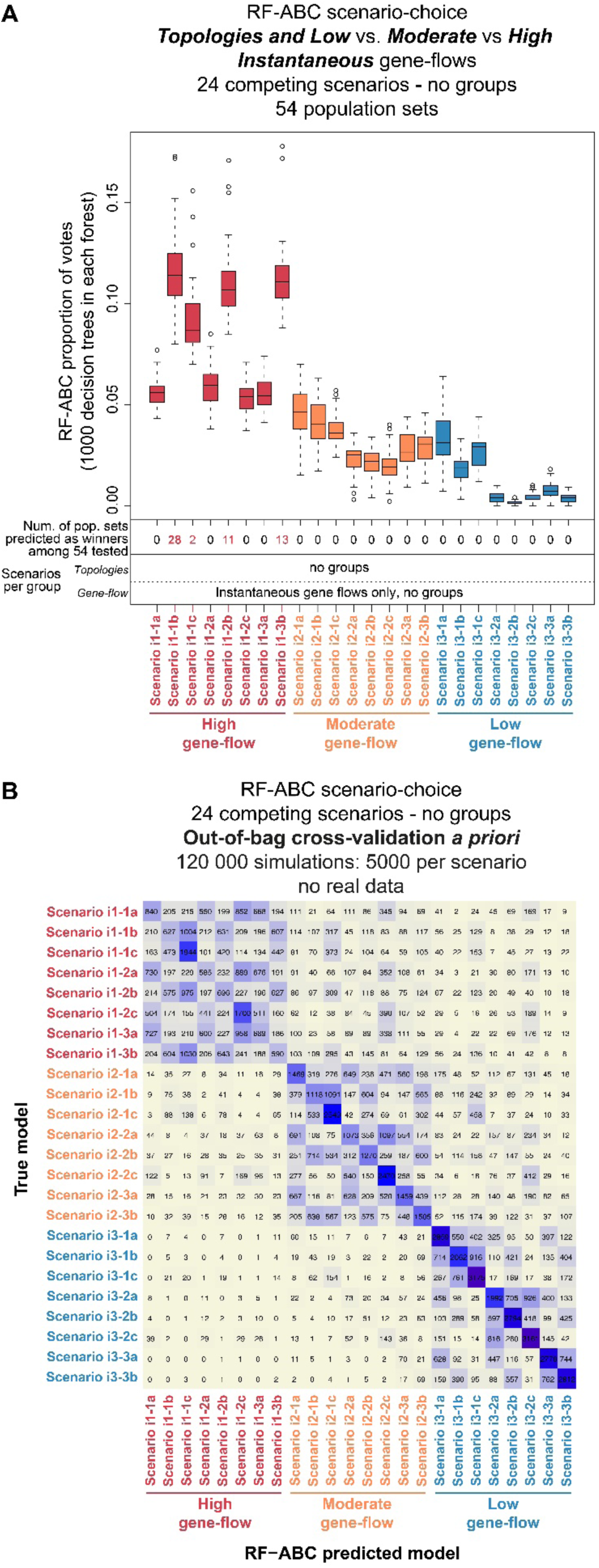
RF-ABC scenario choice among 24 competing scenarios without groups. (**A**) corresponds to the RF-ABC scenario-choice results obtained when considering all 24 instantaneous gene-flow scenarios (5,000 simulations per scenario), as separate competitors without grouping them, for each 54 combinations of five sampled-populations separately. Description of the x and y axis legends are provided in **Figure 5**. Posterior probabilities of choosing the correct scenarios for each 54 sets of population combinations ranged from 0.22380 to 0.30273, compared to a prior probability of 1/24 in this particular test. (**B**) corresponds to the RF-ABC cross-validation prior error for the scenario-choice procedure conducted in (**A**). Errors were obtained without considering observed data and using, instead, all simulations in turn as pseudo-observed data and the remaining simulations to train the RF. Note that prediction and cross-validation error results resemble those obtained for these 24 specific scenarios obtained when considering instead all the 48 scenarios in competition (**Figure S9**).

**Figure S11:**
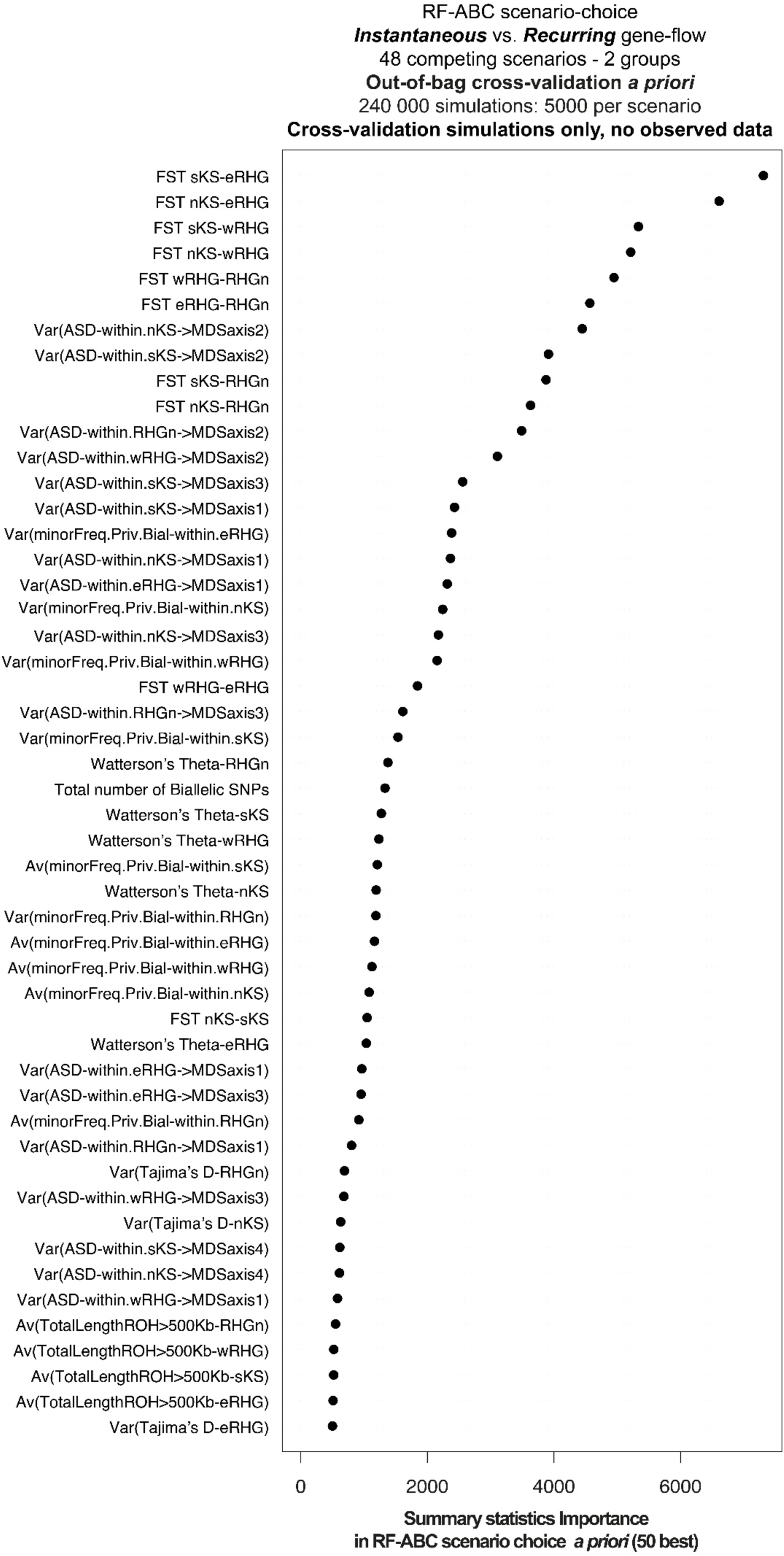
50 best summary statistics relative contribution to the Random Forest ABC scenario-choice *a priori* based on simulations only without observed data – Instantaneous vs Recurring gene-flow processes all gene-flow intensities and all topologies “being equal”. Out-of-bag cross-validation prior error results for this RF-ABC scenario choice analysis are shown in **Figure S7 panel A.**

**Figure S12:**
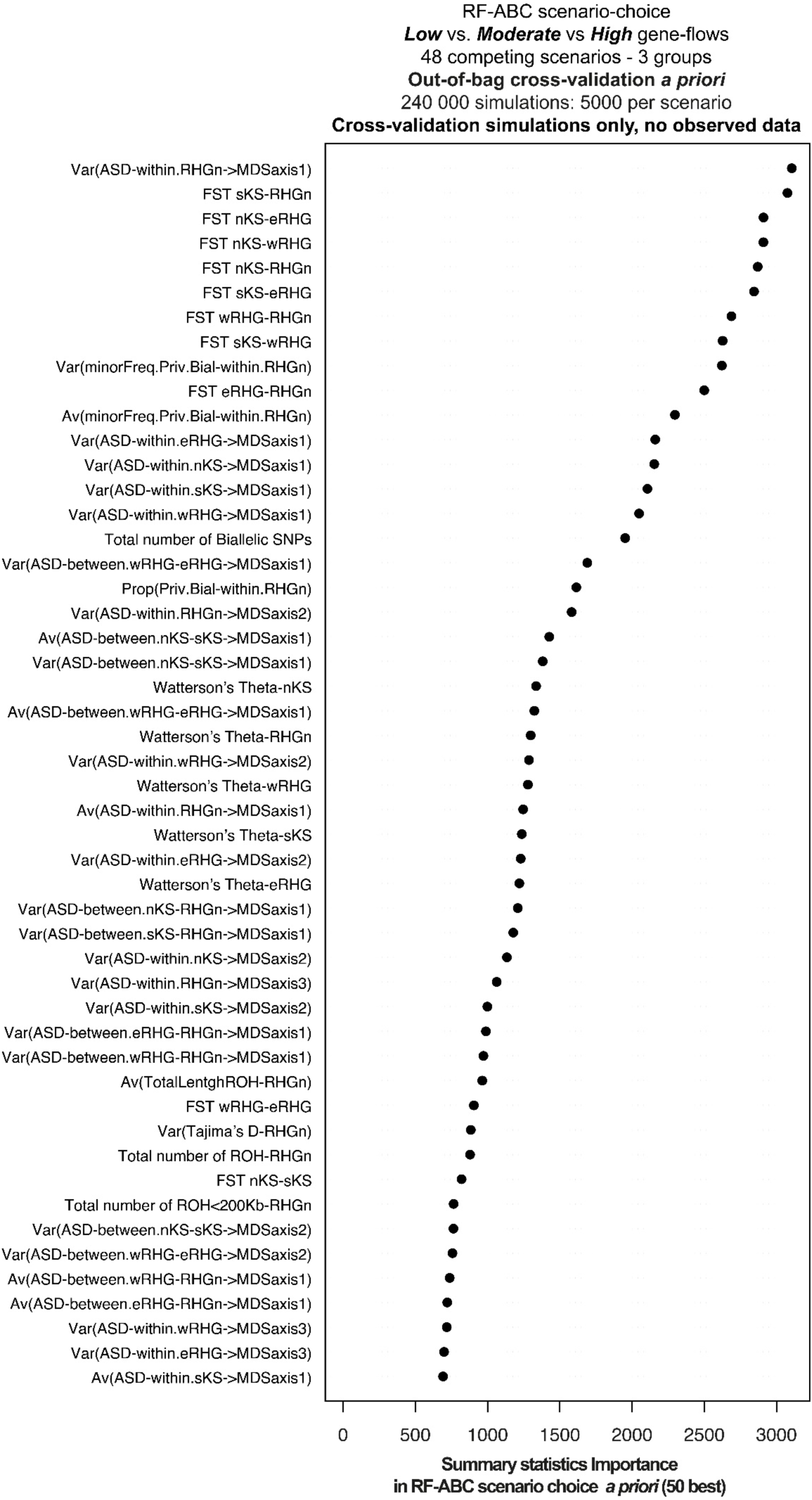
50 best summary statistics relative contribution to the Random Forest ABC scenario-choice *a priori* based on simulations only without observed data – Low vs Moderate vs High gene-flow intensities, all gene-flow processes (instantaneous and recurring) and all topologies “being equal”. Out-of-bag cross-validation prior error results for this RF-ABC scenario choice analysis are shown in **Figure S7 panel B.**

**Figure S13:**
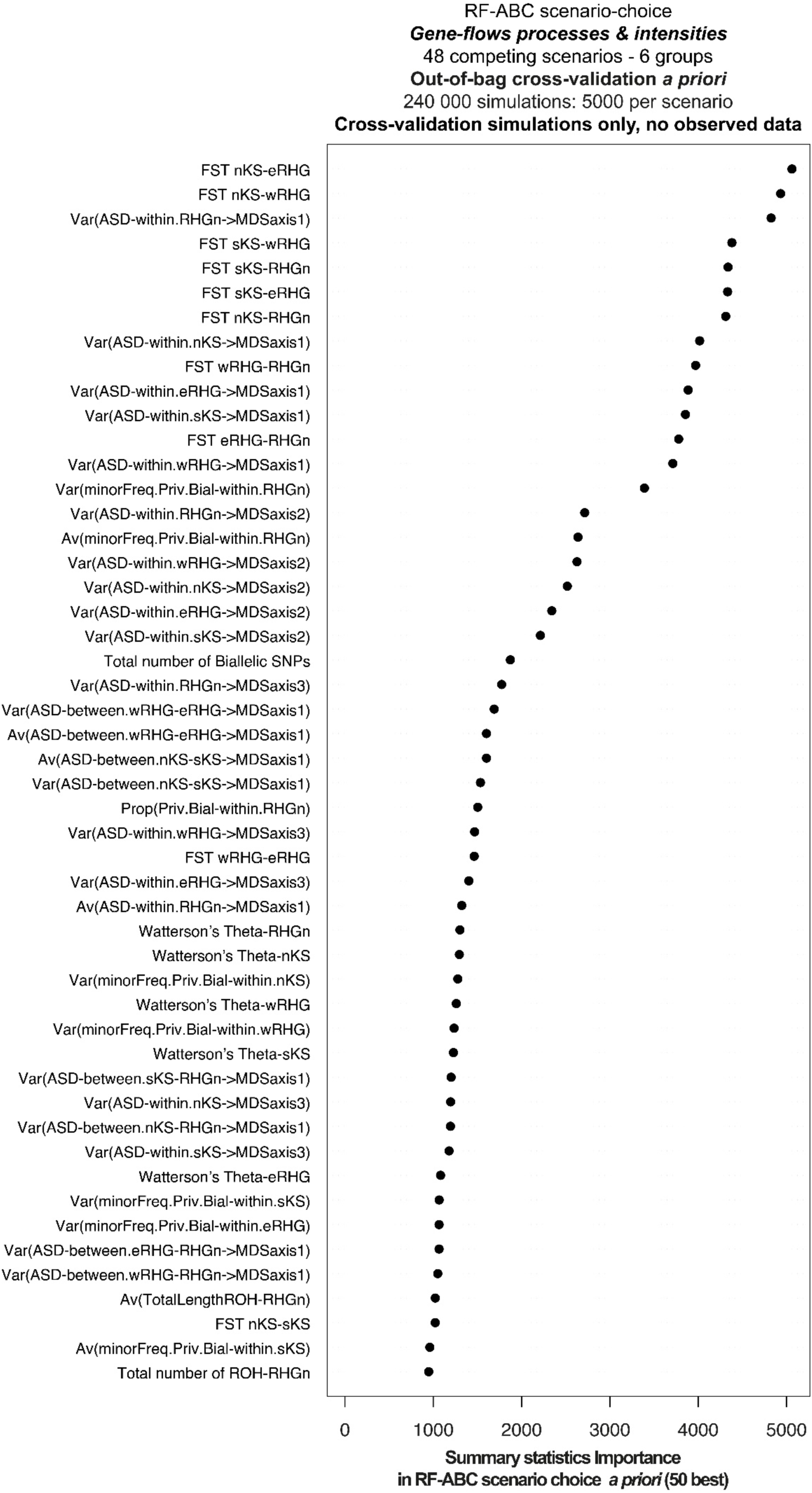
50 best summary statistics relative contribution to the Random Forest ABC scenario-choice *a priori* based on simulations only without observed data – Instantaneous vs Recurring & Low vs Moderate vs High gene-flow intensities, all topologies “being equal”. Out-of-bag cross-validation prior error results for this RF-ABC scenario choice analysis are shown in **Figure S7 panel C.**

**Figure S14:**
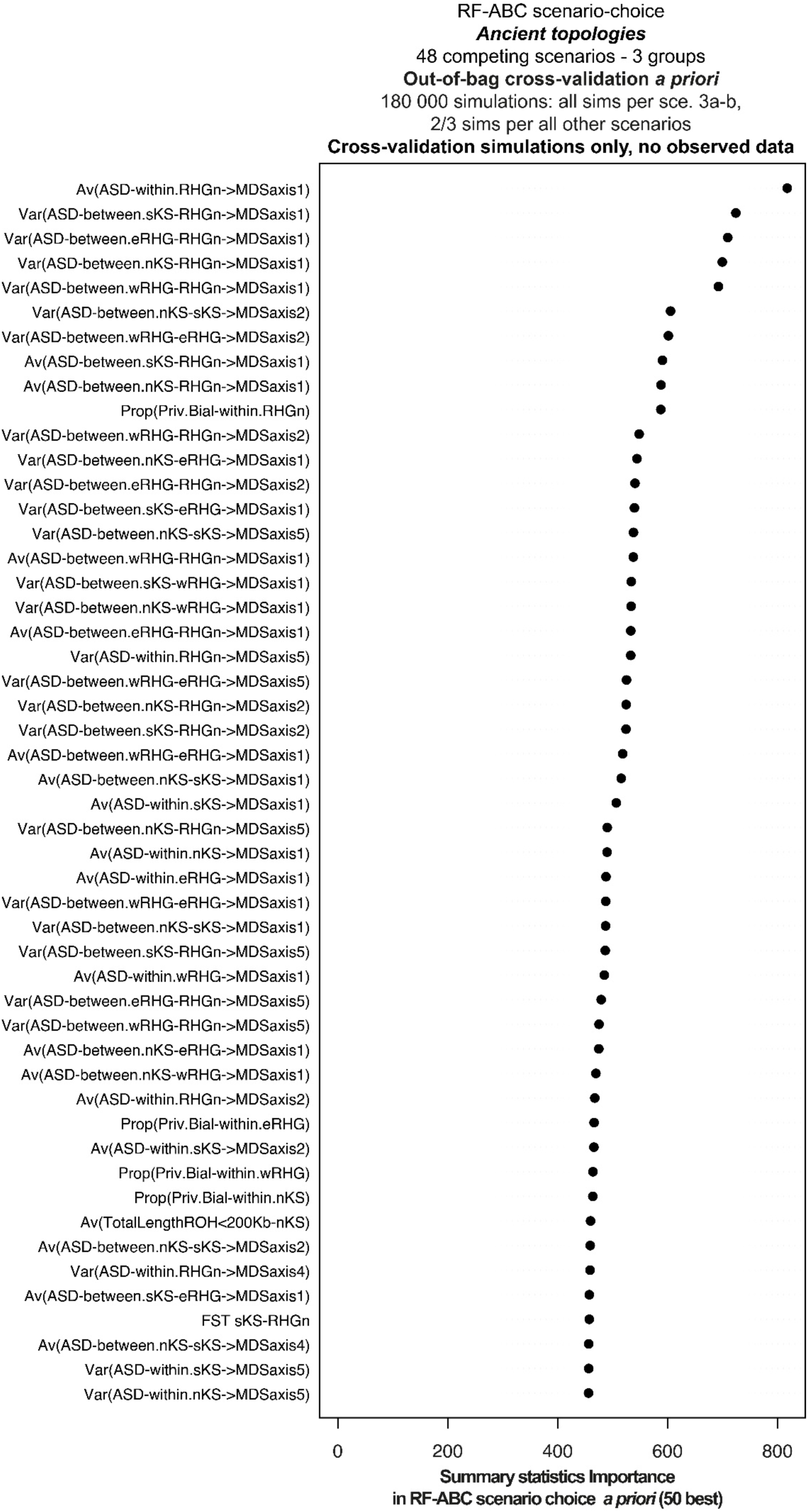
50 best summary statistics relative contribution to the Random Forest ABC scenario-choice *a priori* based on simulations only without observed data – Ancient topologies, all gene-flow processes (instantaneous and recurring) and intensities (Low, moderate, and high), and all recent topologies “being equal”. Out-of-bag cross-validation prior error results for this RF-ABC scenario choice analysis are shown in **Figure S7 panel D.**

**Figure S15:**
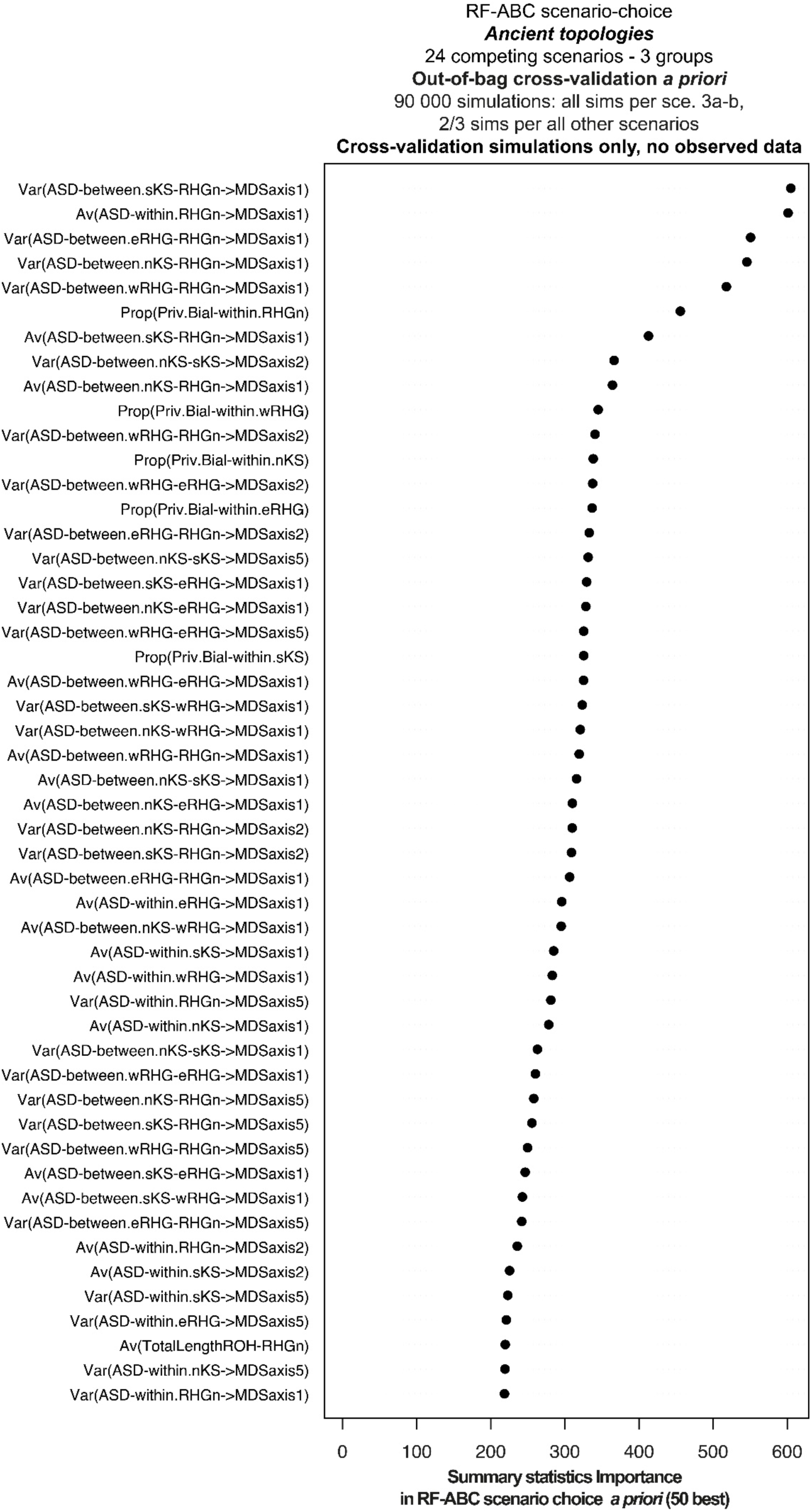
50 best summary statistics relative contribution to the Random Forest ABC scenario-choice *a priori* based on simulations only without observed data – Ancient topologies for instantaneous gene-flow processes only, all intensities (Low, moderate, and high), and all recent topologies “being equal”. Out-of-bag cross-validation prior error results for this RF-ABC scenario choice analysis are shown in **Figure S7 panel E.**

**Figure S16:**
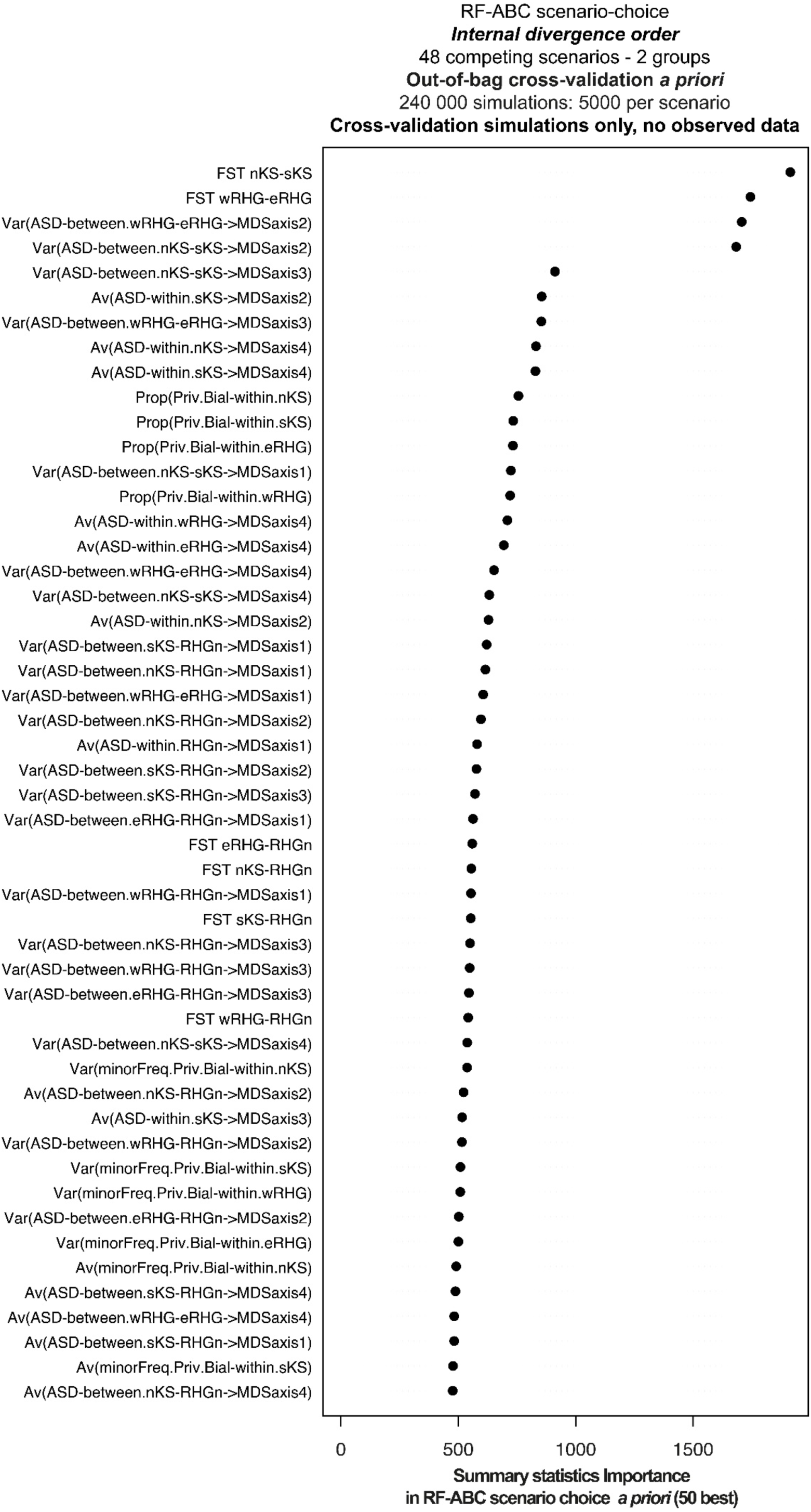
50 best summary statistics relative contribution to the Random Forest ABC scenario-choice *a priori* based on simulations only without observed data – Internal divergence order, all gene-flow processes (instantaneous and recurring) and intensities (Low, moderate, and high), and all ancient topologies “being equal”. Out-of-bag cross-validation prior error results for this RF-ABC scenario choice analysis are shown in **Figure S7 panel F.**

**Figure S17:**
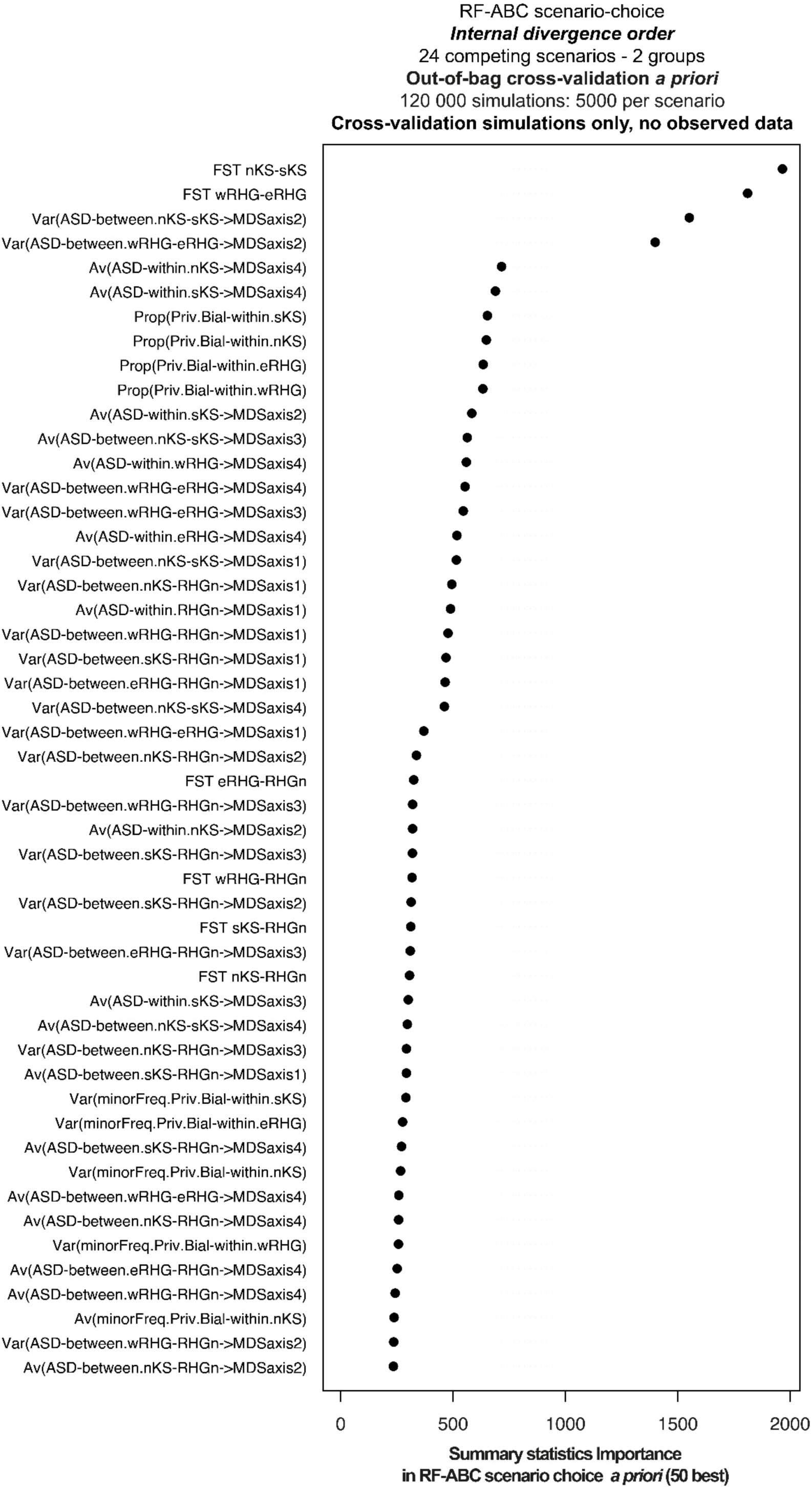
50 best summary statistics relative contribution to the Random Forest ABC scenario-choice *a priori* based on simulations only without observed data – Internal divergence order for instantaneous gene-flow processes only, all intensities (Low, moderate, and high), and all ancient topologies “being equal”. Out-of-bag cross-validation prior error results for this RF-ABC scenario choice analysis are shown in **Figure S7 panel G.**

**Figure S18:**
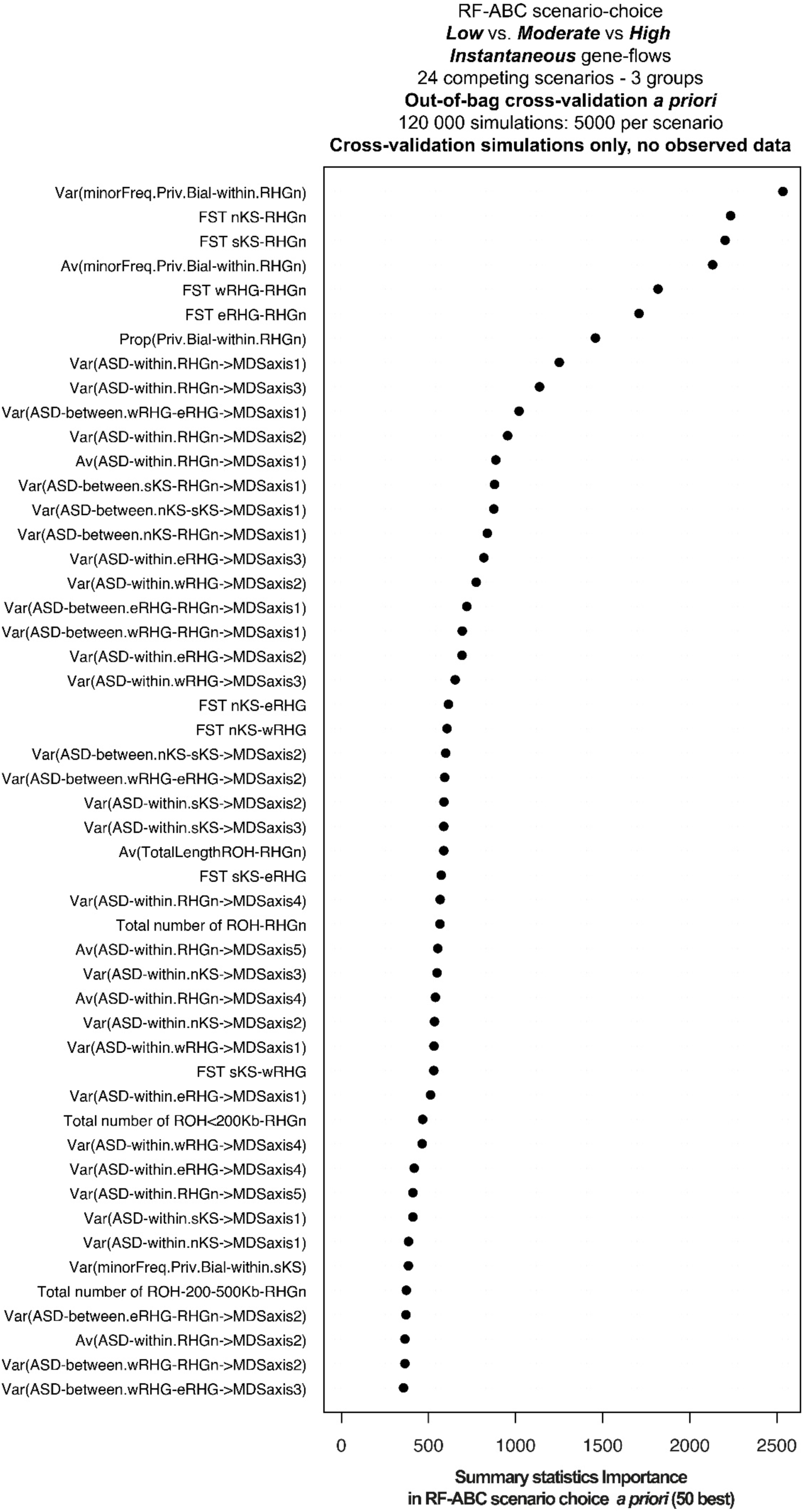
50 best summary statistics relative contribution to the Random Forest ABC scenario-choice *a priori* based on simulations only without observed data – Low vs Moderate vs High gene-flow intensities for instantaneous gene-flow processes only, all topologies “being equal”. Out-of-bag cross-validation prior error results for this RF-ABC scenario choice analysis are shown in **Figure S8 panel B.**

**Figure S19:**
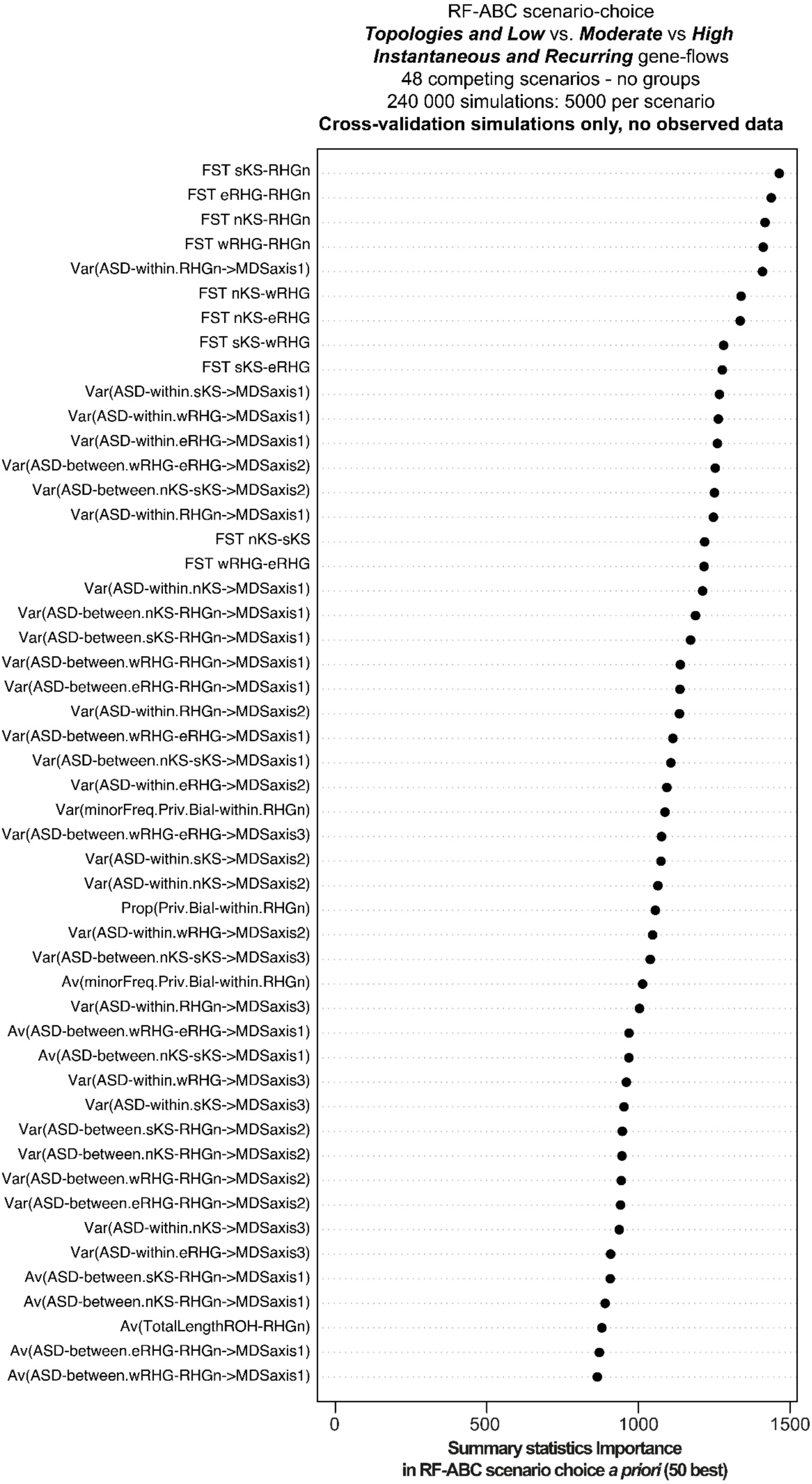
50 best summary statistics relative contribution to the Random Forest ABC scenario-choice *a priori* based on simulations only without observed data – All models in competition. Out-of-bag cross-validation prior error results for this RF-ABC scenario choice analysis are shown in **Figure S9 panel B.**

**Figure S20:**
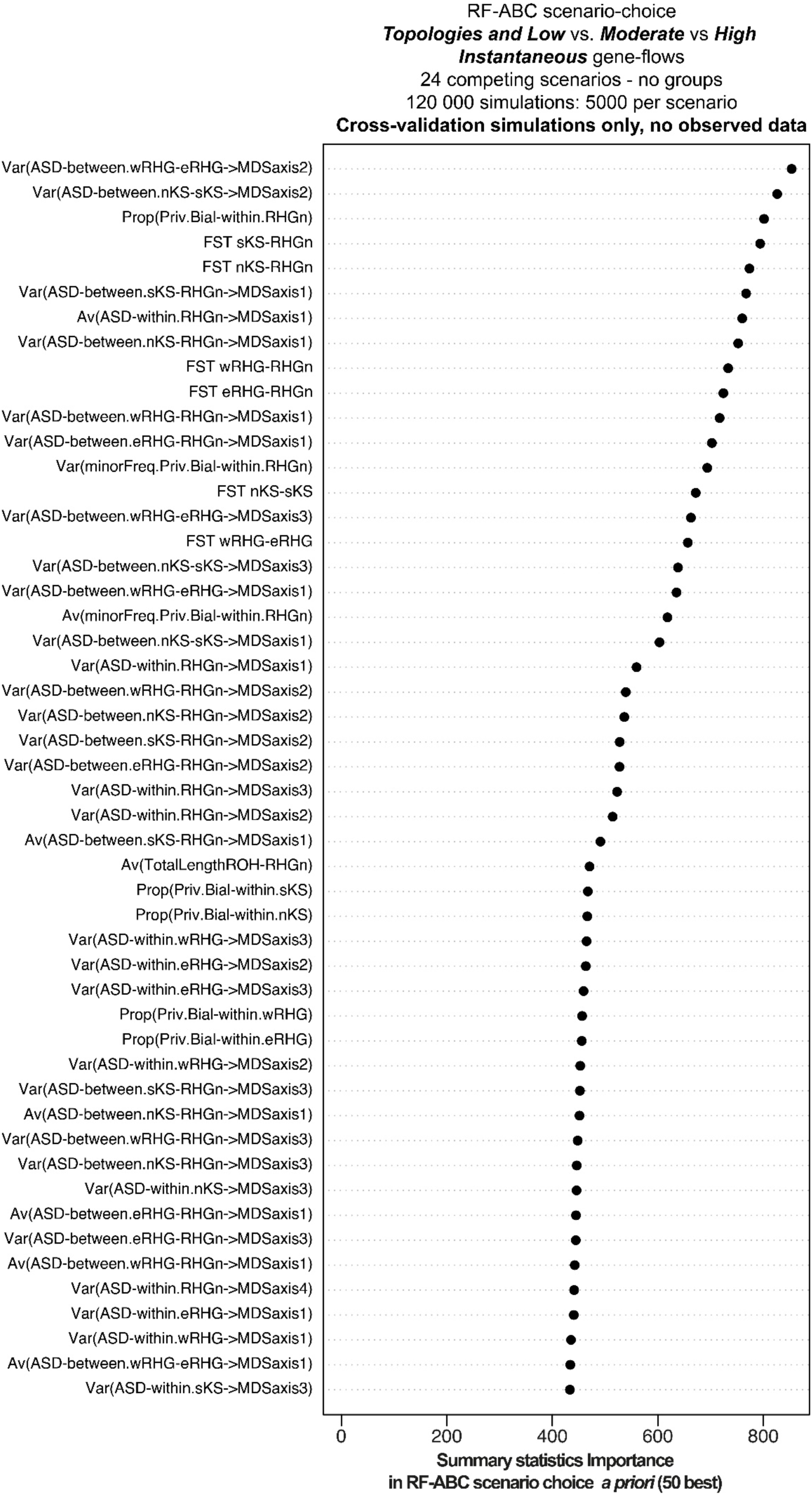
50 best summary statistics relative contribution to the Random Forest ABC scenario-choice *a priori* based on simulations only without observed data – All models in competition for instantaneous gene-flow processes only. Out-of-bag cross-validation prior error results for this RF-ABC scenario choice analysis are shown in **Figure S10 panel B.**

**Figure S21:**
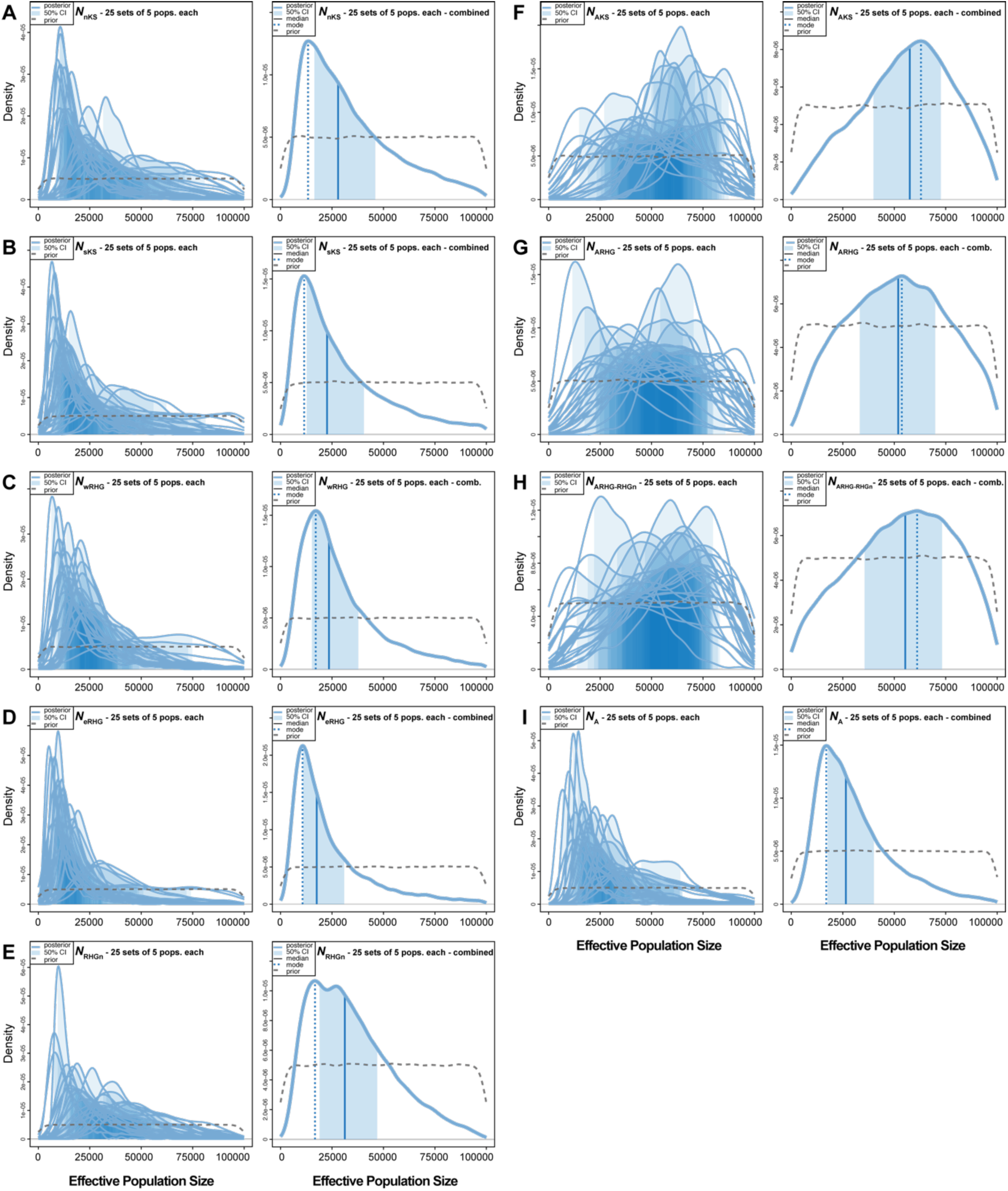
ABC posterior distribution of effective population size parameters. Neural Network Approximate Bayesian Computation ^43,44^, posterior parameter joint estimations of Effective population sizes *Ne* (in generations before present) for 25 sets of five Central and Southern African populations for which the best scenario identified by RF-ABC was Scenario i1-1b (**Figure 4**, **Table S13**). NN ABC posterior parameter estimation procedures were conducted using 100,000 simulations under Scenario i1-1b, each simulation corresponding to a single vector of parameter values drawn randomly from prior distributions provided in **Table S8**. We considered 43 neurons in the hidden layer of the NN and a tolerance level of 0.01, corresponding to the 1,000 simulations providing summary-statistics closest to the observed ones, for each 25 separate analyses. NN posterior estimates are based on the logit transformation of parameter values using an Epanechnikov kernel between the corresponding parameter’s prior bounds (see **Material and Methods** and **Table S8**). Posterior parameter densities are represented with solid blue lines. 50% Credibility Intervals are represented as the light blue area under the density. The median and mode values are represented as a solid and dotted blue vertical line, respectively. Parameter prior distributions are represented as dotted grey lines. For all panels, the left plots represent the NN-ABC posterior parameter distributions for each 25 sets of five Central and Southern African populations under Scenario i1-1b, separately (**Table S13** and **Table S14**). For all panels, the right plots represent a single parameter posterior distribution obtained from combining (concatenating) the 25 posterior distributions together. See **Figure 4** and **Table S8** for all parameters’ descriptions in each panel. Results are also provided in **Table S17**.

**Figure S22:**
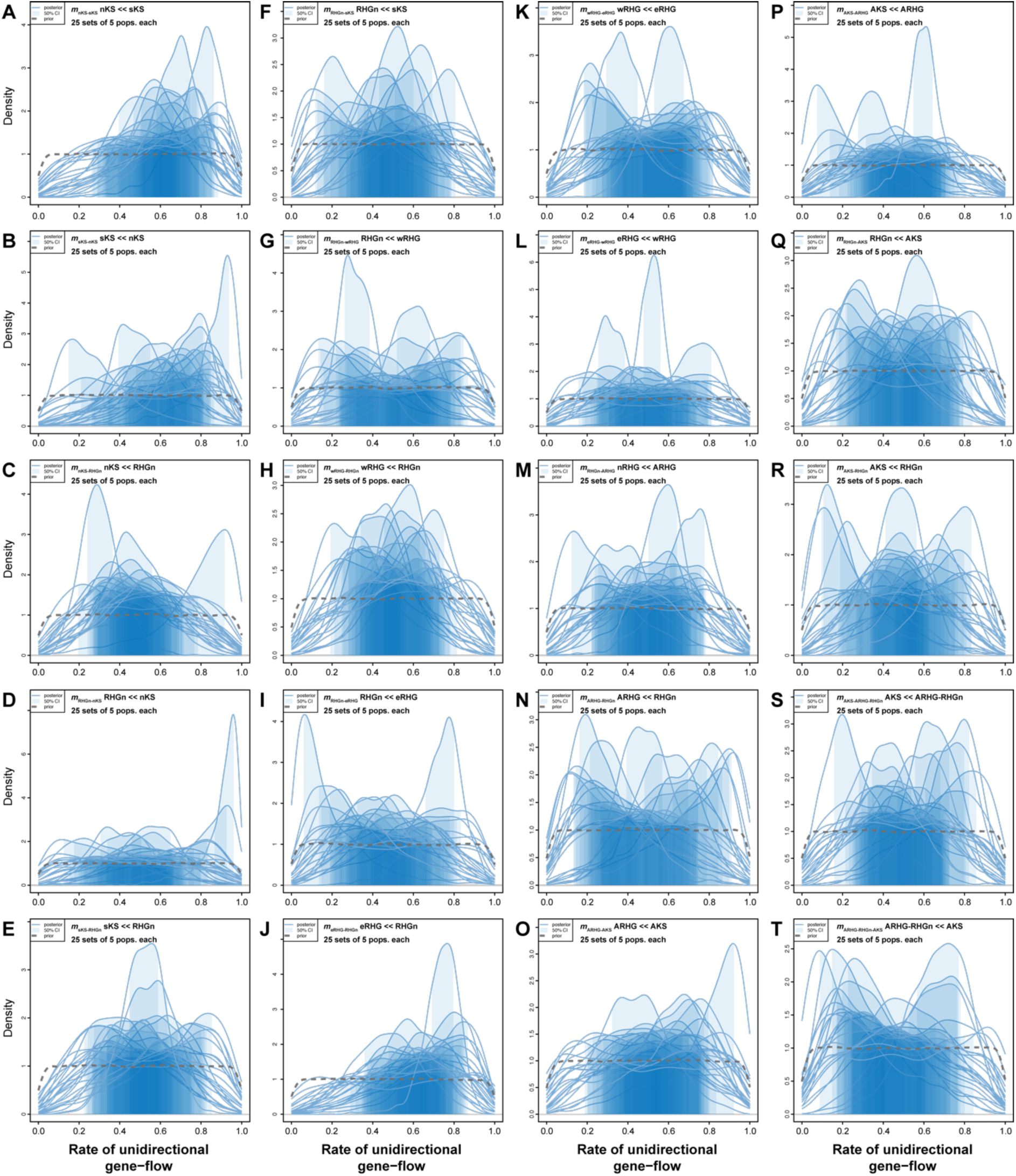
ABC posterior distribution of potentially asymmetric instantaneous gene-flow intensity parameters. Neural Network Approximate Bayesian Computation ^43,44^, posterior parameter joint estimations of gene-flow intensity parameters *m* (in generations before present) for 25 sets of five Central and Southern African populations for which the best scenario identified by RF-ABC was Scenario i1-1b (**Figure 4**, **Table S13**). NN ABC posterior parameter estimation procedures were conducted using 100,000 simulations under Scenario i1-1b, each simulation corresponding to a single vector of parameter values drawn randomly from prior distributions provided in **Table S8**. We considered 43 neurons in the hidden layer of the NN and a tolerance level of 0.01, corresponding to the 1,000 simulations providing summary-statistics closest to the observed ones, for each 25 separate analyses. NN posterior estimates are based on the logit transformation of parameter values using an Epanechnikov kernel between the corresponding parameter’s prior bounds (see **Material and Methods** and **Table S8**). Posterior parameter densities are represented with solid blue lines. 50% Credibility Intervals are represented as the light blue area under the density. Parameter prior distributions are represented as dotted grey lines. For all panels, the plots represent the NN-ABC posterior parameter distributions for each 25 sets of five Central and Southern African populations under Scenario i1-1b, separately (**Table S13** and **Table S14**). See **Figure 4** and **Table S8** for all parameters’ descriptions in each panel. Note that, overall, parameters are relatively little departing from their priors for numerous sets of population combinations among the 25. Also, posterior parameter distributions that are substantially departing from their priors are often highly differing from one another for each parameter. Therefore, we considered these parameters as unsatisfactorily estimated in our analyses and discuss this limitation in the **Discussion** section of the main text.

**Figure S23:**
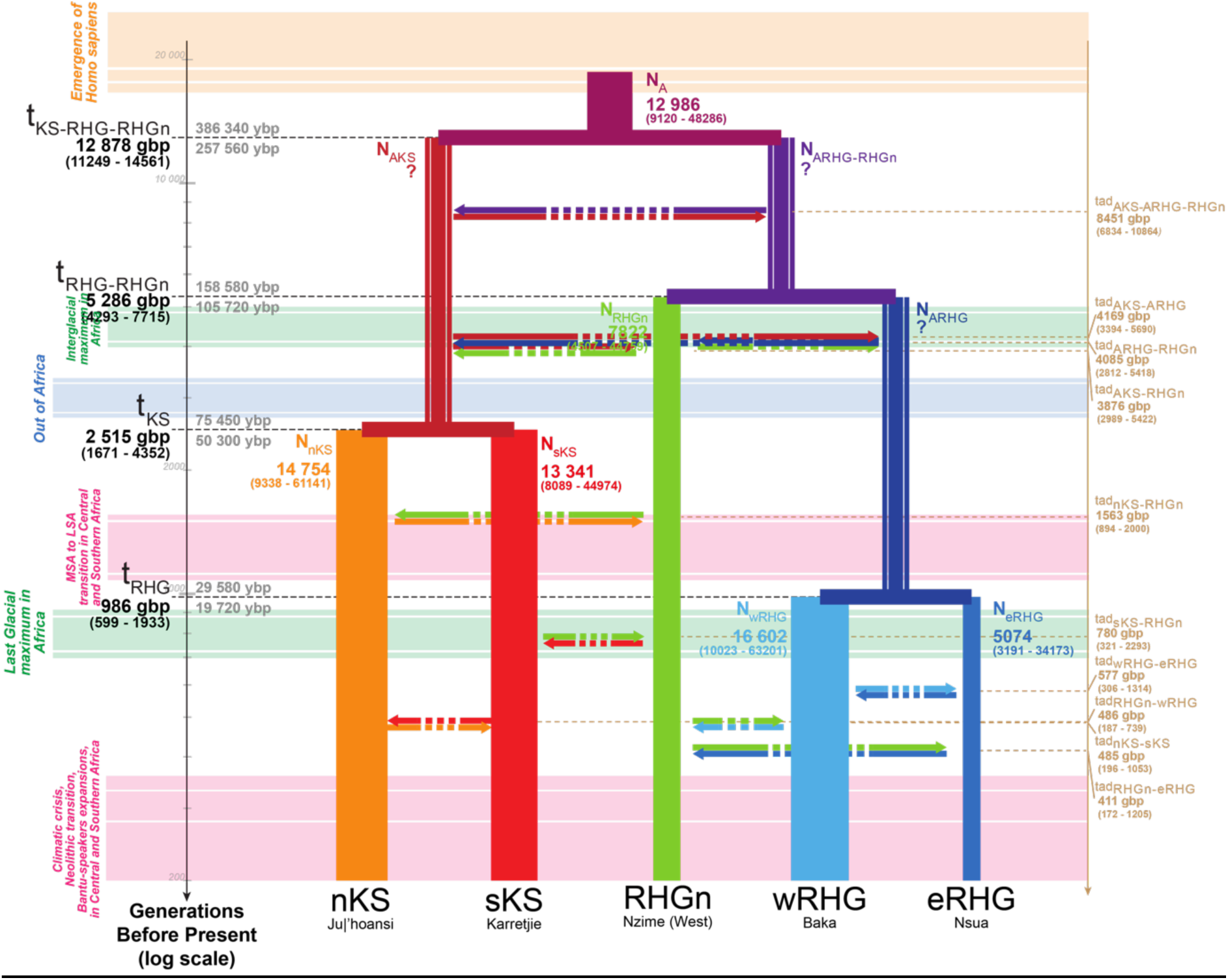
Schematic inferred demographic and migration history of Central and Southern African combination of populations n°1. Schematic representation of Scenario i1-1b (**Figure 4**), and Neural Network ABC posterior parameter mode estimates summarizing results obtained for the first combination of sampled populations (**Table S13, Table S14**). Gene-flow arrows are indicated forward in time. For the time of each divergence and gene-flow event, mode point estimates are provided in generations before present (gbp) in bold, and 90% Credibility Intervals are provided between parentheses (**Table S14**). We provide two estimates of the divergence times estimates in years before present (ybp), one (upper) corresponding to 30 years per generation and the other (lower) to 20 years per generation ^66,67^. Mode point estimates of effective population sizes *Ne* are provided in numbers of diploid effective individuals and width of lineages are proportional to the estimated *Ne* (**Table S14**, **Figure S21**). NN-ABC posterior distributions for the effective population sizes of the ancestral Khoe-San lineage (AKS), for the ancestral RHG lineage (ARHG), and for the lineage ancestral to RHGn and ARHG, were all three poorly distinguished from their respective prior distributions and with high cross-validation posterior parameter estimation errors (**Figure S21**, **Table S16, Table S17**), as indicated by the question marks. NN-ABC posterior distributions for instantaneous asymmetric gene-flow rates were overall poorly departing from their priors and with high cross-validation posterior parameter estimation errors (**Figure S22**, **Table S14, Table S16**), as indicated by the doted arrows. All posterior distributions are shown graphically in **Figure 6**, **Figure 7, Figure S21, Figure S22**, and detailed in **Table S14**, with cross-validation posterior errors for each parameter in **Table S16**.

**Figure S24:**
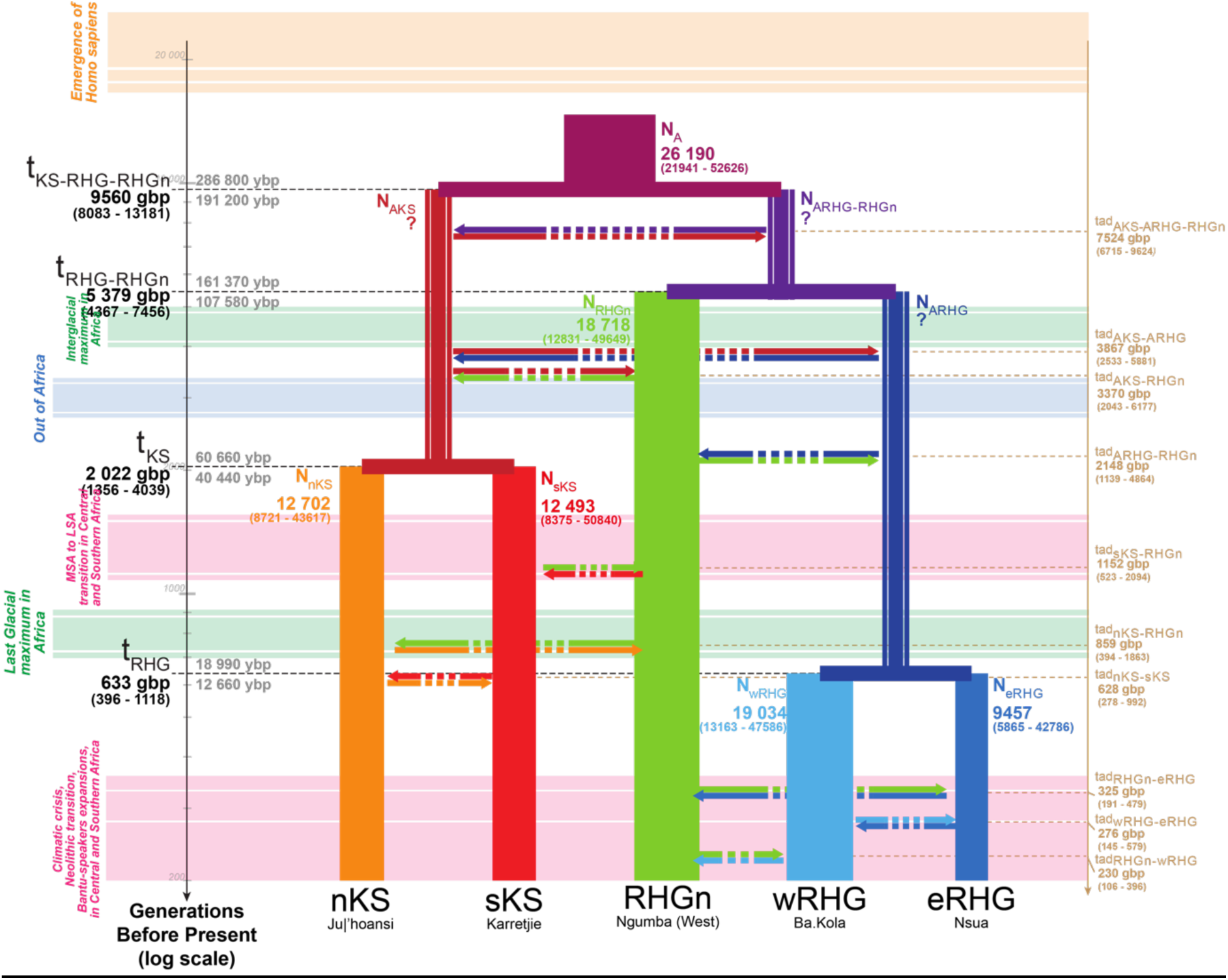
Schematic inferred demographic and migration history of Central and Southern African combination of populations n°2. Schematic representation of Scenario i1-1b (**Figure 4**), and Neural Network ABC posterior parameter mode estimates summarizing results obtained for the second combination of sampled populations (**Table S13, Table S14**). Gene-flow arrows are indicated forward in time. For the time of each divergence and gene-flow event, mode point estimates are provided in generations before present (gbp) in bold, and 90% Credibility Intervals are provided between parentheses (**Table S14**). We provide two estimates of the divergence times estimates in years before present (ybp), one (upper) corresponding to 30 years per generation and the other (lower) to 20 years per generation ^66,67^. Mode point estimates of effective population sizes *Ne* are provided in numbers of diploid effective individuals and width of lineages are proportional to the estimated *Ne* (**Table S14**, **Figure S21**). NN-ABC posterior distributions for the effective population sizes of the ancestral Khoe-San lineage (AKS), for the ancestral RHG lineage (ARHG), and for the lineage ancestral to RHGn and ARHG, were all three poorly distinguished from their respective prior distributions and with high cross-validation posterior parameter estimation errors (**Figure S21**, **Table S16, Table S17**), as indicated by the question marks. NN-ABC posterior distributions for instantaneous asymmetric gene-flow rates were overall poorly departing from their priors and with high cross-validation posterior parameter estimation errors (**Figure S22**, **Table S14, Table S16**), as indicated by the doted arrows. All posterior distributions are shown graphically in **Figure 6**, **Figure 7, Figure S21, Figure S22**, and detailed in **Table S14**, with cross-validation posterior errors for each parameter in **Table S16**.

**Figure S25:**
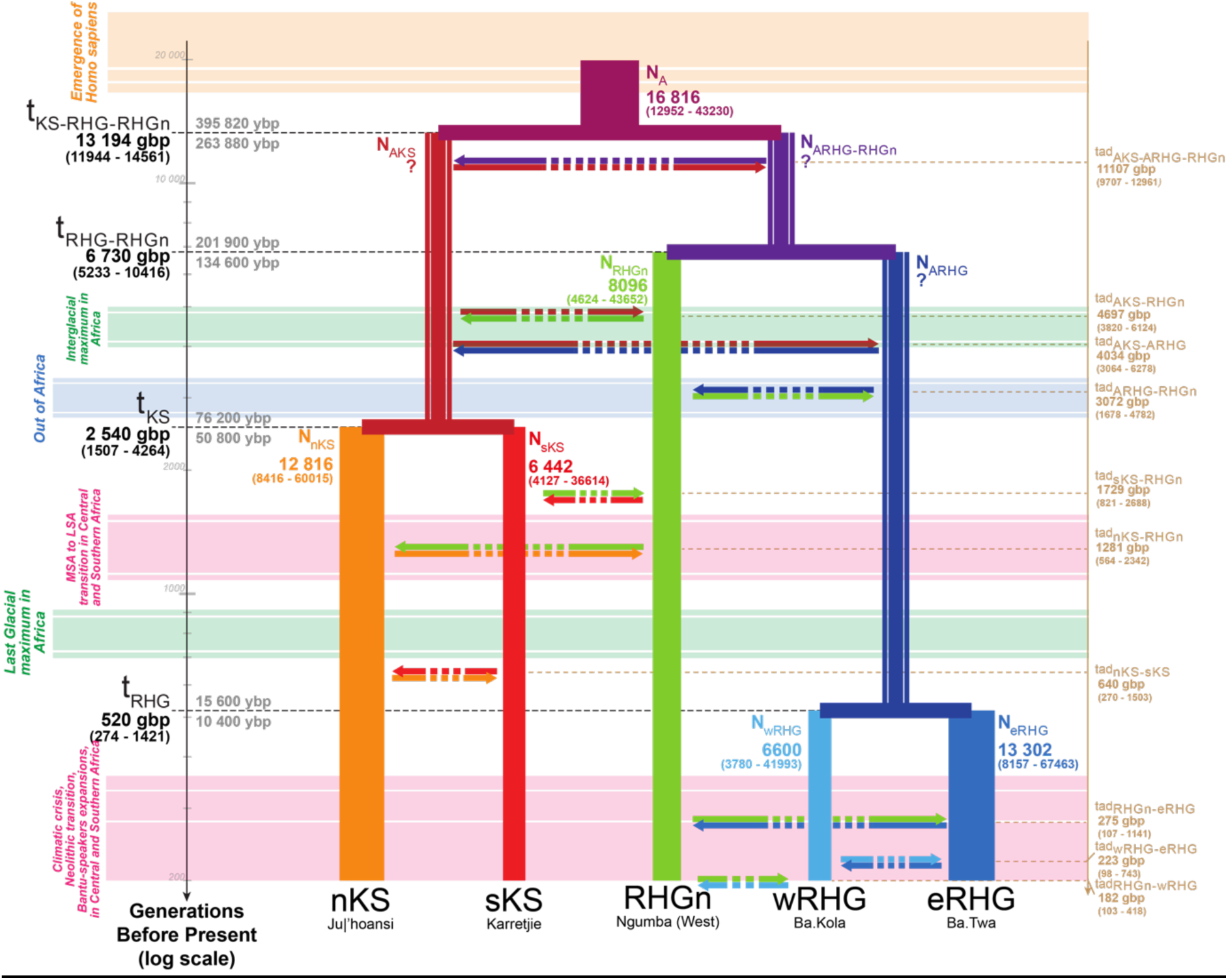
Schematic inferred demographic and migration history of Central and Southern African combination of populations n°3. Schematic representation of Scenario i1-1b (**Figure 4**), and Neural Network ABC posterior parameter mode estimates summarizing results obtained for the third combination of sampled populations (**Table S13, Table S14**). Gene-flow arrows are indicated forward in time. For the time of each divergence and gene-flow event, mode point estimates are provided in generations before present (gbp) in bold, and 90% Credibility Intervals are provided between parentheses (**Table S14**). We provide two estimates of the divergence times estimates in years before present (ybp), one (upper) corresponding to 30 years per generation and the other (lower) to 20 years per generation ^66,67^. Mode point estimates of effective population sizes *Ne* are provided in numbers of diploid effective individuals and width of lineages are proportional to the estimated *Ne* (**Table S14**, **Figure S21**). NN-ABC posterior distributions for the effective population sizes of the ancestral Khoe-San lineage (AKS), for the ancestral RHG lineage (ARHG), and for the lineage ancestral to RHGn and ARHG, were all three poorly distinguished from their respective prior distributions and with high cross-validation posterior parameter estimation errors (**Figure S21**, **Table S16, Table S17**), as indicated by the question marks. NN-ABC posterior distributions for instantaneous asymmetric gene-flow rates were overall poorly departing from their priors and with high cross-validation posterior parameter estimation errors (**Figure S22**, **Table S14, Table S16**), as indicated by the doted arrows. All posterior distributions are shown graphically in **Figure 6**, **Figure 7, Figure S21, Figure S22**, and detailed in **Table S14**, with cross-validation posterior errors for each parameter in **Table S16**.

**Figure S26:**
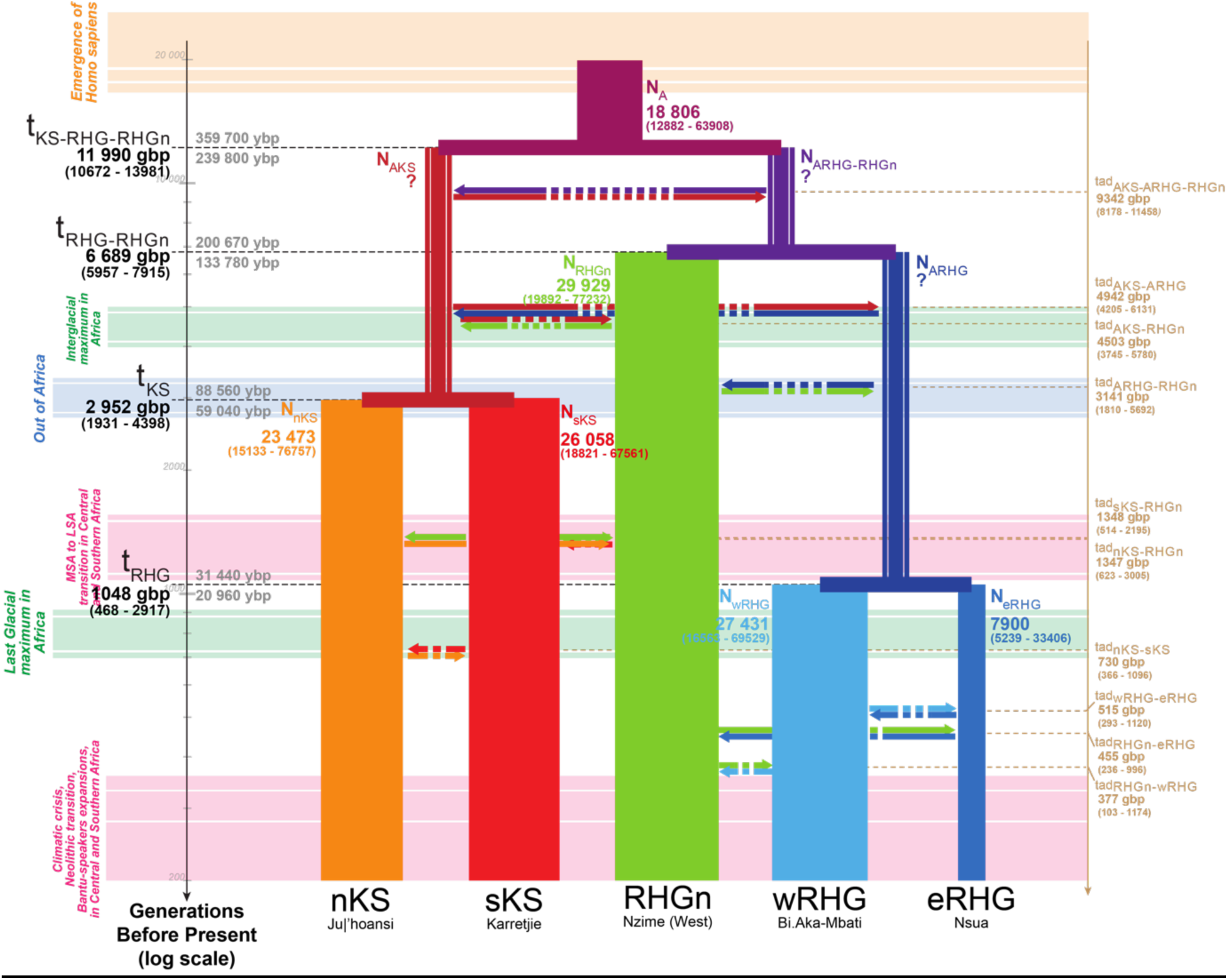
Schematic inferred demographic and migration history of Central and Southern African combination of populations n°4. Schematic representation of Scenario i1-1b (**Figure 4**), and Neural Network ABC posterior parameter mode estimates summarizing results obtained for the fourth combination of sampled populations (**Table S13, Table S14**). Gene-flow arrows are indicated forward in time. For the time of each divergence and gene-flow event, mode point estimates are provided in generations before present (gbp) in bold, and 90% Credibility Intervals are provided between parentheses (**Table S14**). We provide two estimates of the divergence times estimates in years before present (ybp), one (upper) corresponding to 30 years per generation and the other (lower) to 20 years per generation ^66,67^. Mode point estimates of effective population sizes *Ne* are provided in numbers of diploid effective individuals and width of lineages are proportional to the estimated *Ne* (**Table S14**, **Figure S21**). NN-ABC posterior distributions for the effective population sizes of the ancestral Khoe-San lineage (AKS), for the ancestral RHG lineage (ARHG), and for the lineage ancestral to RHGn and ARHG, were all three poorly distinguished from their respective prior distributions and with high cross-validation posterior parameter estimation errors (**Figure S21**, **Table S16, Table S17**), as indicated by the question marks. NN-ABC posterior distributions for instantaneous asymmetric gene-flow rates were overall poorly departing from their priors and with high cross-validation posterior parameter estimation errors (**Figure S22**, **Table S14, Table S16**), as indicated by the doted arrows. All posterior distributions are shown graphically in **Figure 6**, **Figure 7, Figure S21, Figure S22**, and detailed in **Table S14**, with cross-validation posterior errors for each parameter in **Table S16**.

**Figure S27:**
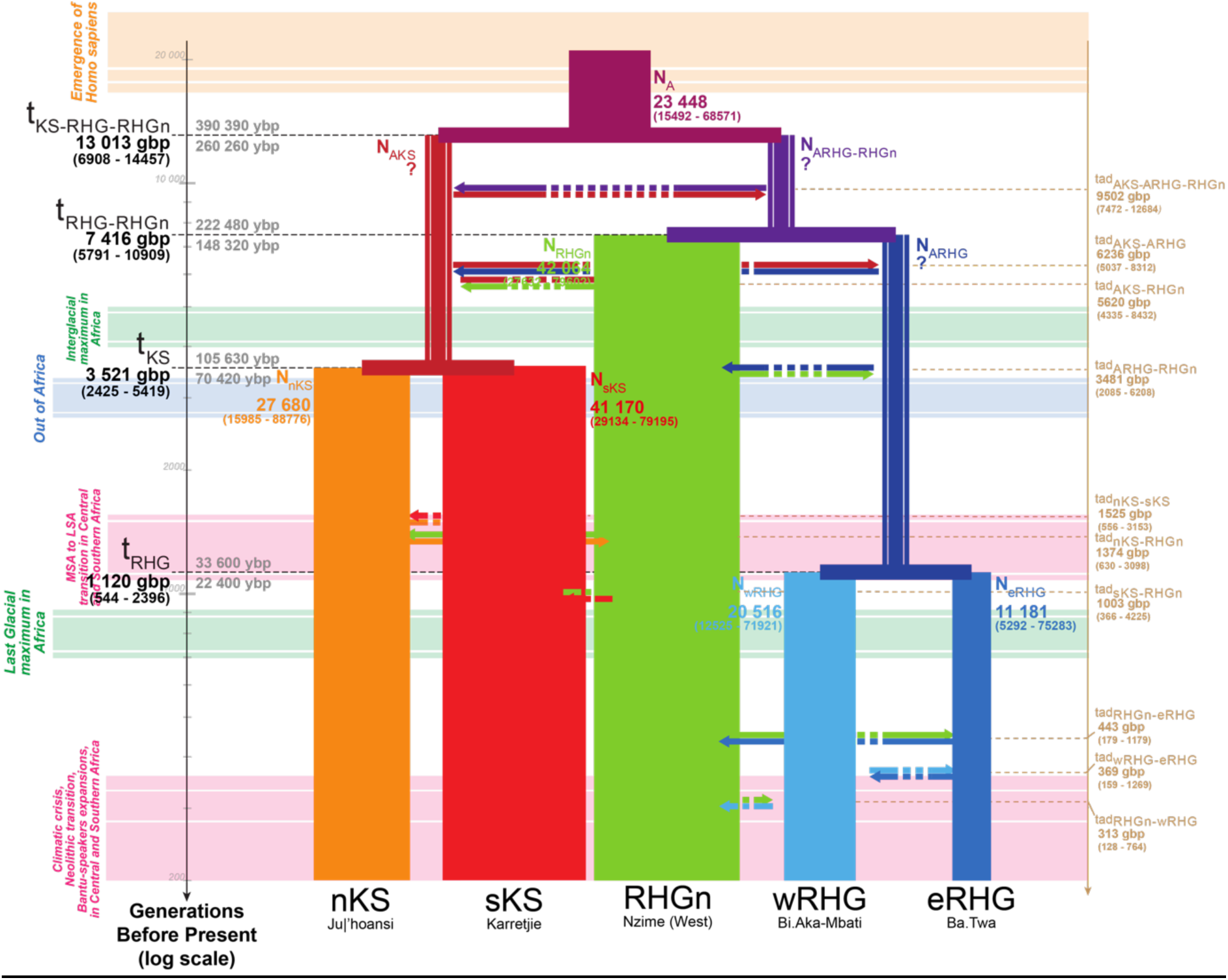
Schematic inferred demographic and migration history of Central and Southern African combination of populations n°5. Schematic representation of Scenario i1-1b (**Figure 4**), and Neural Network ABC posterior parameter mode estimates summarizing results obtained for the fifth combination of sampled populations (**Table S13, Table S14**). Gene-flow arrows are indicated forward in time. For the time of each divergence and gene-flow event, mode point estimates are provided in generations before present (gbp) in bold, and 90% Credibility Intervals are provided between parentheses (**Table S14**). We provide two estimates of the divergence times estimates in years before present (ybp), one (upper) corresponding to 30 years per generation and the other (lower) to 20 years per generation ^66,67^. Mode point estimates of effective population sizes *Ne* are provided in numbers of diploid effective individuals and width of lineages are proportional to the estimated *Ne* (**Table S14**, **Figure S21**). NN-ABC posterior distributions for the effective population sizes of the ancestral Khoe-San lineage (AKS), for the ancestral RHG lineage (ARHG), and for the lineage ancestral to RHGn and ARHG, were all three poorly distinguished from their respective prior distributions and with high cross-validation posterior parameter estimation errors (**Figure S21**, **Table S16, Table S17**), as indicated by the question marks. NN-ABC posterior distributions for instantaneous asymmetric gene-flow rates were overall poorly departing from their priors and with high cross-validation posterior parameter estimation errors (**Figure S22**, **Table S14, Table S16**), as indicated by the doted arrows. All posterior distributions are shown graphically in **Figure 6**, **Figure 7, Figure S21, Figure S22**, and detailed in **Table S14**, with cross-validation posterior errors for each parameter in **Table S16**.

**Figure S28:**
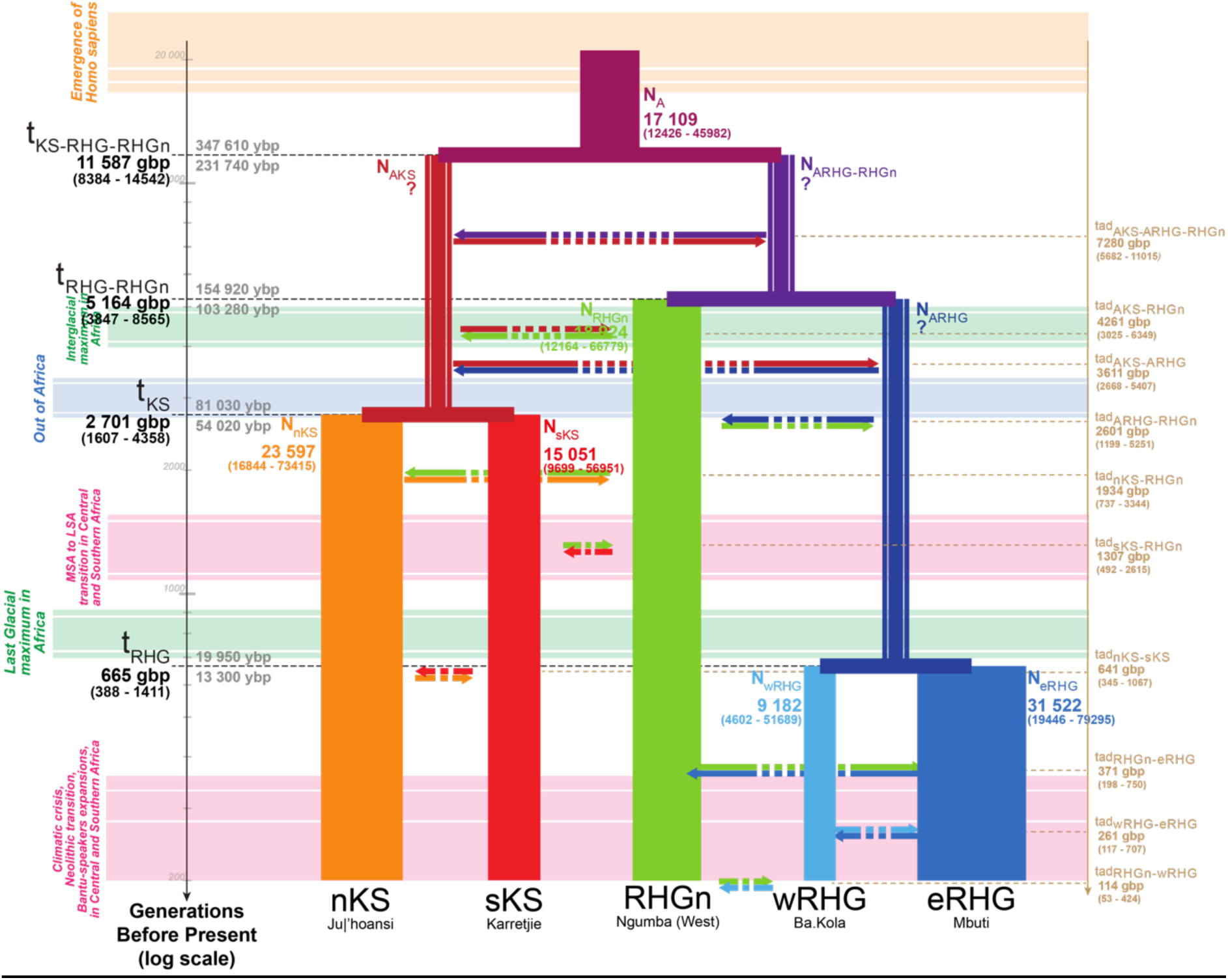
Schematic inferred demographic and migration history of Central and Southern African combination of populations n°6. Schematic representation of Scenario i1-1b (**Figure 4**), and Neural Network ABC posterior parameter mode estimates summarizing results obtained for the sixth combination of sampled populations (**Table S13, Table S14**). Gene-flow arrows are indicated forward in time. For the time of each divergence and gene-flow event, mode point estimates are provided in generations before present (gbp) in bold, and 90% Credibility Intervals are provided between parentheses (**Table S14**). We provide two estimates of the divergence times estimates in years before present (ybp), one (upper) corresponding to 30 years per generation and the other (lower) to 20 years per generation ^66,67^. Mode point estimates of effective population sizes *Ne* are provided in numbers of diploid effective individuals and width of lineages are proportional to the estimated *Ne* (**Table S14**, **Figure S21**). NN-ABC posterior distributions for the effective population sizes of the ancestral Khoe-San lineage (AKS), for the ancestral RHG lineage (ARHG), and for the lineage ancestral to RHGn and ARHG, were all three poorly distinguished from their respective prior distributions and with high cross-validation posterior parameter estimation errors (**Figure S21**, **Table S16, Table S17**), as indicated by the question marks. NN-ABC posterior distributions for instantaneous asymmetric gene-flow rates were overall poorly departing from their priors and with high cross-validation posterior parameter estimation errors (**Figure S22**, **Table S14, Table S16**), as indicated by the doted arrows. All posterior distributions are shown graphically in **Figure 6**, **Figure 7, Figure S21, Figure S22**, and detailed in **Table S14**, with cross-validation posterior errors for each parameter in **Table S16**.

**Figure S29:**
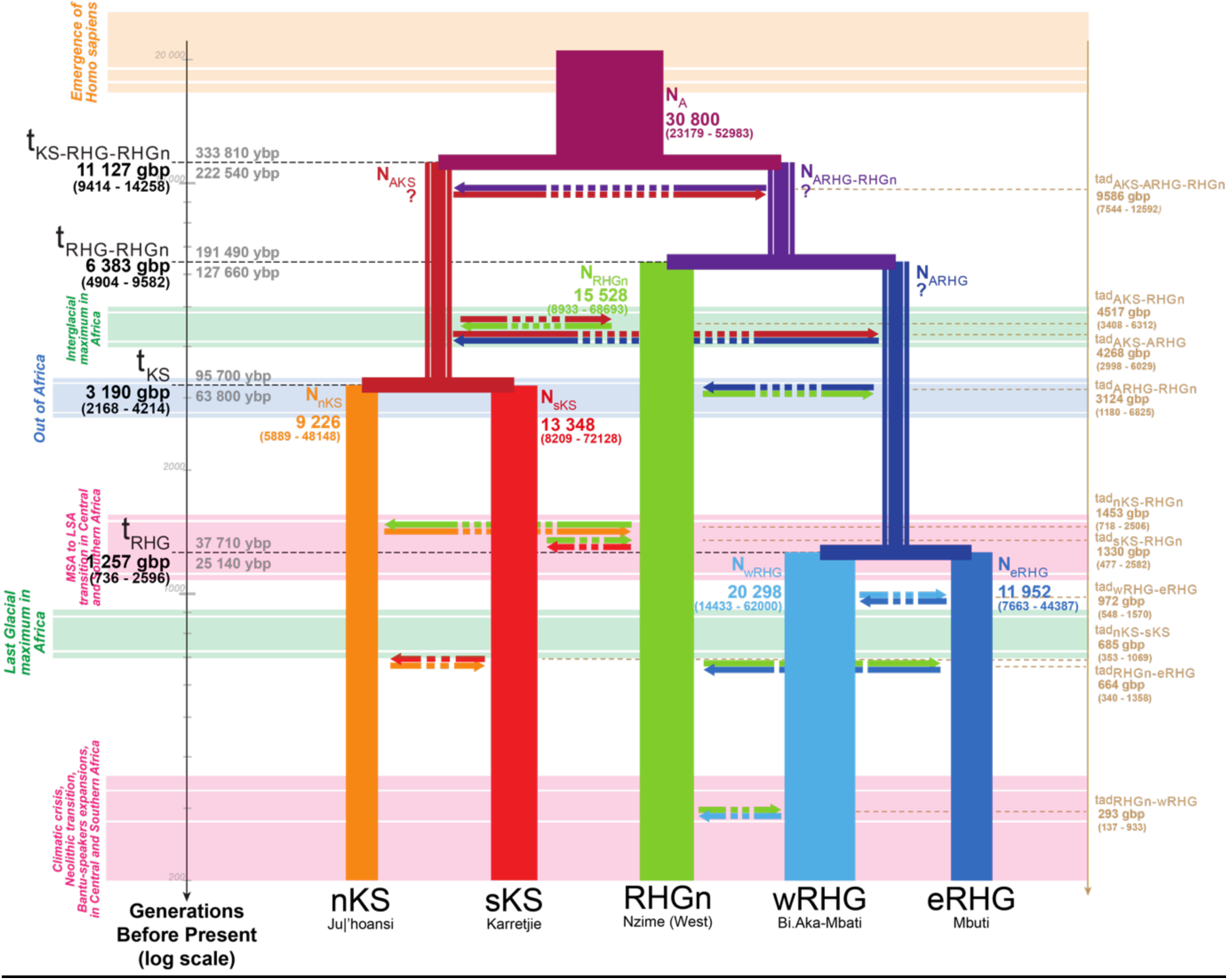
Schematic inferred demographic and migration history of Central and Southern African combination of populations n°7. Schematic representation of Scenario i1-1b (**Figure 4**), and Neural Network ABC posterior parameter mode estimates summarizing results obtained for the seventh combination of sampled populations (**Table S13, Table S14**). Gene-flow arrows are indicated forward in time. For the time of each divergence and gene-flow event, mode point estimates are provided in generations before present (gbp) in bold, and 90% Credibility Intervals are provided between parentheses (**Table S14**). We provide two estimates of the divergence times estimates in years before present (ybp), one (upper) corresponding to 30 years per generation and the other (lower) to 20 years per generation ^66,67^. Mode point estimates of effective population sizes *Ne* are provided in numbers of diploid effective individuals and width of lineages are proportional to the estimated *Ne* (**Table S14**, **Figure S21**). NN-ABC posterior distributions for the effective population sizes of the ancestral Khoe-San lineage (AKS), for the ancestral RHG lineage (ARHG), and for the lineage ancestral to RHGn and ARHG, were all three poorly distinguished from their respective prior distributions and with high cross-validation posterior parameter estimation errors (**Figure S21**, **Table S16, Table S17**), as indicated by the question marks. NN-ABC posterior distributions for instantaneous asymmetric gene-flow rates were overall poorly departing from their priors and with high cross-validation posterior parameter estimation errors (**Figure S22**, **Table S14, Table S16**), as indicated by the doted arrows. All posterior distributions are shown graphically in **Figure 6**, **Figure 7, Figure S21, Figure S22**, and detailed in **Table S14**, with cross-validation posterior errors for each parameter in **Table S16**.

**Figure S30:**
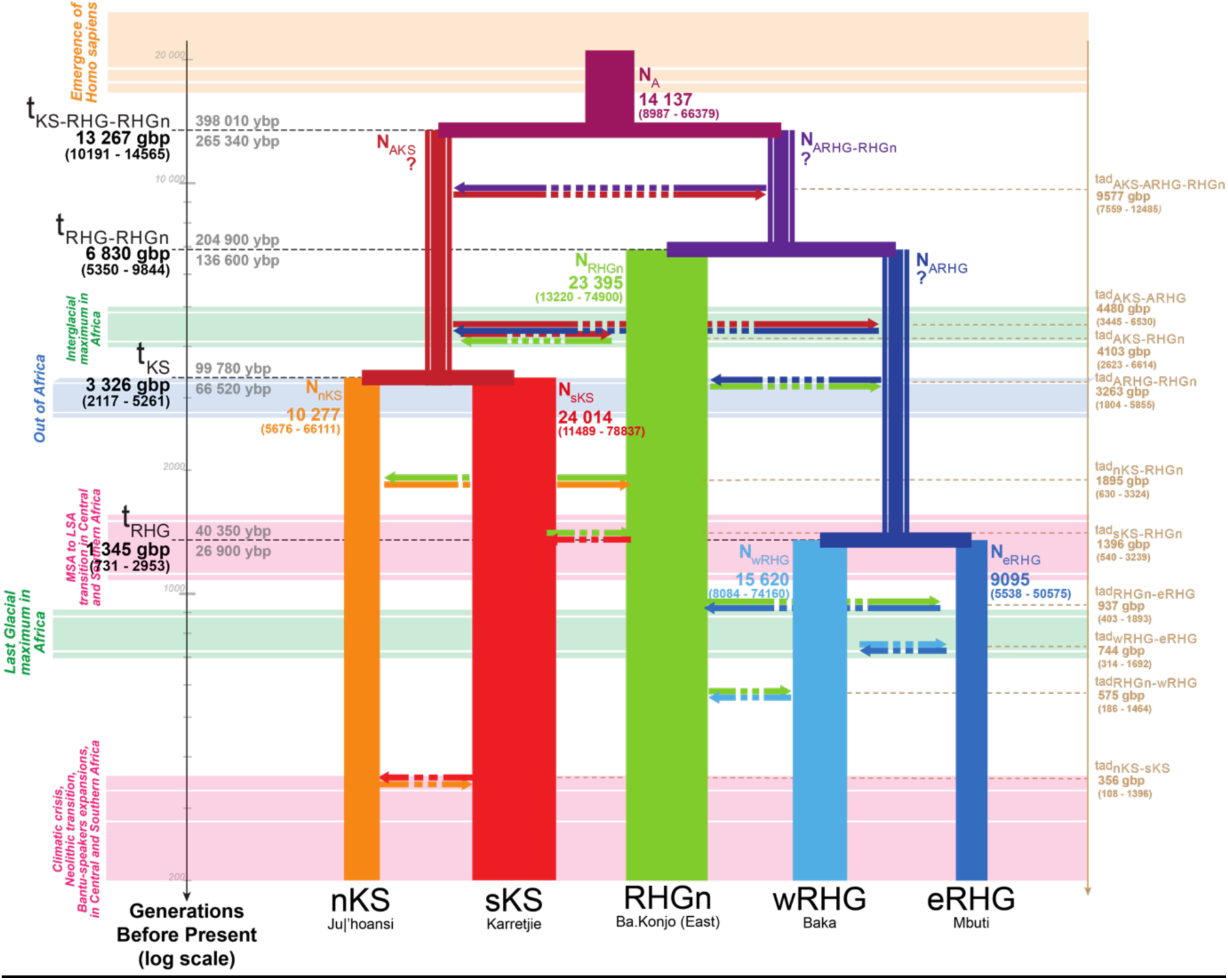
Schematic inferred demographic and migration history of Central and Southern African combination of populations n°8. Schematic representation of Scenario i1-1b (**Figure 4**), and Neural Network ABC posterior parameter mode estimates summarizing results obtained for the eigth combination of sampled populations (**Table S13, Table S14**). Gene-flow arrows are indicated forward in time. For the time of each divergence and gene-flow event, mode point estimates are provided in generations before present (gbp) in bold, and 90% Credibility Intervals are provided between parentheses (**Table S14**). We provide two estimates of the divergence times estimates in years before present (ybp), one (upper) corresponding to 30 years per generation and the other (lower) to 20 years per generation ^66,67^. Mode point estimates of effective population sizes *Ne* are provided in numbers of diploid effective individuals and width of lineages are proportional to the estimated *Ne* (**Table S14**, **Figure S21**). NN-ABC posterior distributions for the effective population sizes of the ancestral Khoe-San lineage (AKS), for the ancestral RHG lineage (ARHG), and for the lineage ancestral to RHGn and ARHG, were all three poorly distinguished from their respective prior distributions and with high cross-validation posterior parameter estimation errors (**Figure S21**, **Table S16, Table S17**), as indicated by the question marks. NN-ABC posterior distributions for instantaneous asymmetric gene-flow rates were overall poorly departing from their priors and with high cross-validation posterior parameter estimation errors (**Figure S22**, **Table S14, Table S16**), as indicated by the doted arrows. All posterior distributions are shown graphically in **Figure 6**, **Figure 7, Figure S21, Figure S22**, and detailed in **Table S14**, with cross-validation posterior errors for each parameter in **Table S16**.

**Figure S31:**
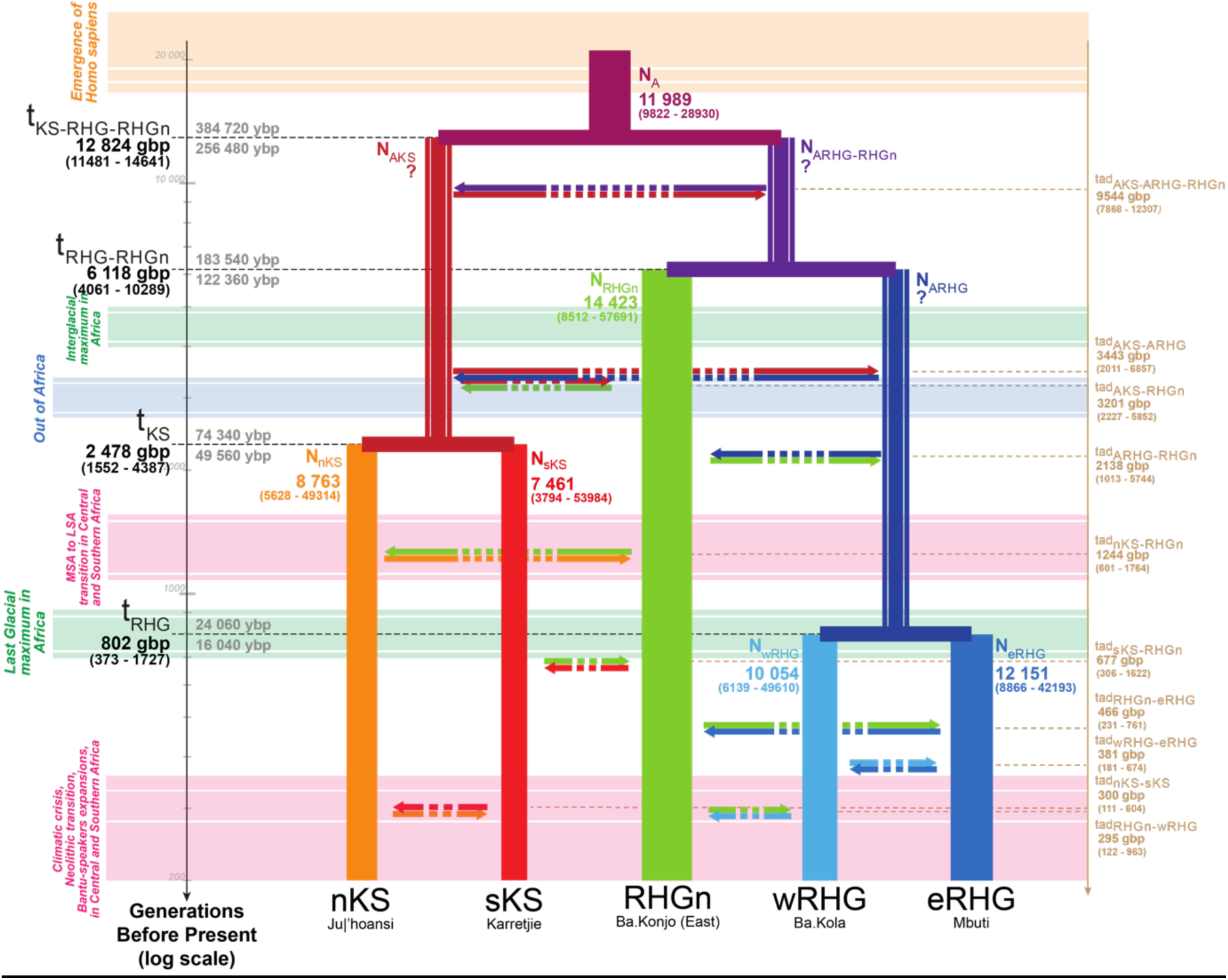
Schematic inferred demographic and migration history of Central and Southern African combination of populations n°9. Schematic representation of Scenario i1-1b (**Figure 4**), and Neural Network ABC posterior parameter mode estimates summarizing results obtained for the ninth combination of sampled populations (**Table S13, Table S14**). Gene-flow arrows are indicated forward in time. For the time of each divergence and gene-flow event, mode point estimates are provided in generations before present (gbp) in bold, and 90% Credibility Intervals are provided between parentheses (**Table S14**). We provide two estimates of the divergence times estimates in years before present (ybp), one (upper) corresponding to 30 years per generation and the other (lower) to 20 years per generation ^66,67^. Mode point estimates of effective population sizes *Ne* are provided in numbers of diploid effective individuals and width of lineages are proportional to the estimated *Ne* (**Table S14**, **Figure S21**). NN-ABC posterior distributions for the effective population sizes of the ancestral Khoe-San lineage (AKS), for the ancestral RHG lineage (ARHG), and for the lineage ancestral to RHGn and ARHG, were all three poorly distinguished from their respective prior distributions and with high cross-validation posterior parameter estimation errors (**Figure S21**, **Table S16, Table S17**), as indicated by the question marks. NN-ABC posterior distributions for instantaneous asymmetric gene-flow rates were overall poorly departing from their priors and with high cross-validation posterior parameter estimation errors (**Figure S22**, **Table S14, Table S16**), as indicated by the doted arrows. All posterior distributions are shown graphically in **Figure 6**, **Figure 7, Figure S21, Figure S22**, and detailed in **Table S14**, with cross-validation posterior errors for each parameter in **Table S16**.

**Figure S32:**
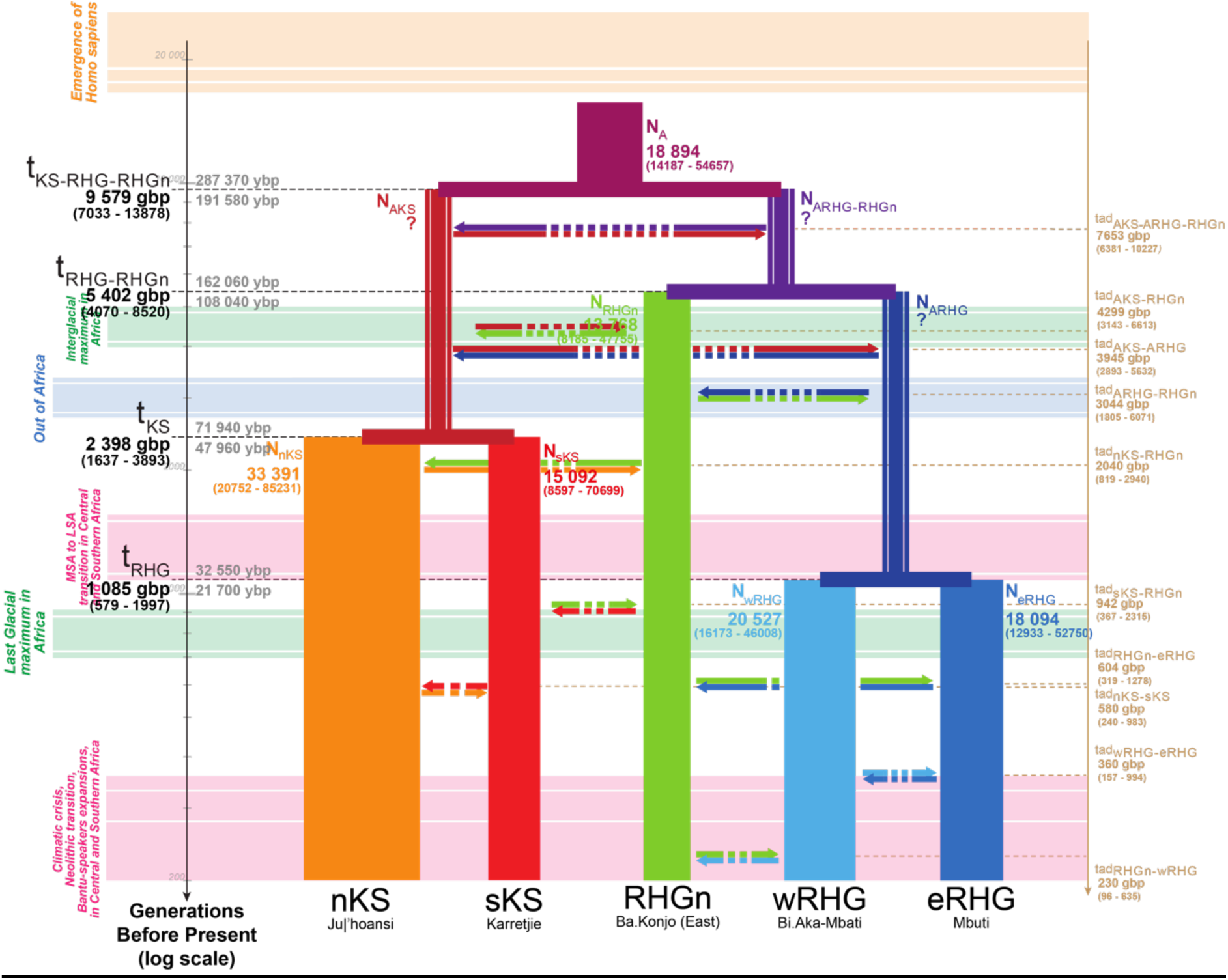
Schematic inferred demographic and migration history of Central and Southern African combination of populations n°10. Schematic representation of Scenario i1-1b (**Figure 4**), and Neural Network ABC posterior parameter mode estimates summarizing results obtained for the 10^th^ combination of sampled populations (**Table S13, Table S14**). Gene-flow arrows are indicated forward in time. For the time of each divergence and gene-flow event, mode point estimates are provided in generations before present (gbp) in bold, and 90% Credibility Intervals are provided between parentheses (**Table S14**). We provide two estimates of the divergence times estimates in years before present (ybp), one (upper) corresponding to 30 years per generation and the other (lower) to 20 years per generation ^66,67^. Mode point estimates of effective population sizes *Ne* are provided in numbers of diploid effective individuals and width of lineages are proportional to the estimated *Ne* (**Table S14**, **Figure S21**). NN-ABC posterior distributions for the effective population sizes of the ancestral Khoe-San lineage (AKS), for the ancestral RHG lineage (ARHG), and for the lineage ancestral to RHGn and ARHG, were all three poorly distinguished from their respective prior distributions and with high cross-validation posterior parameter estimation errors (**Figure S21**, **Table S16, Table S17**), as indicated by the question marks. NN-ABC posterior distributions for instantaneous asymmetric gene-flow rates were overall poorly departing from their priors and with high cross-validation posterior parameter estimation errors (**Figure S22**, **Table S14, Table S16**), as indicated by the doted arrows. All posterior distributions are shown graphically in **Figure 6**, **Figure 7, Figure S21, Figure S22**, and detailed in **Table S14**, with cross-validation posterior errors for each parameter in **Table S16**.

**Figure S33:**
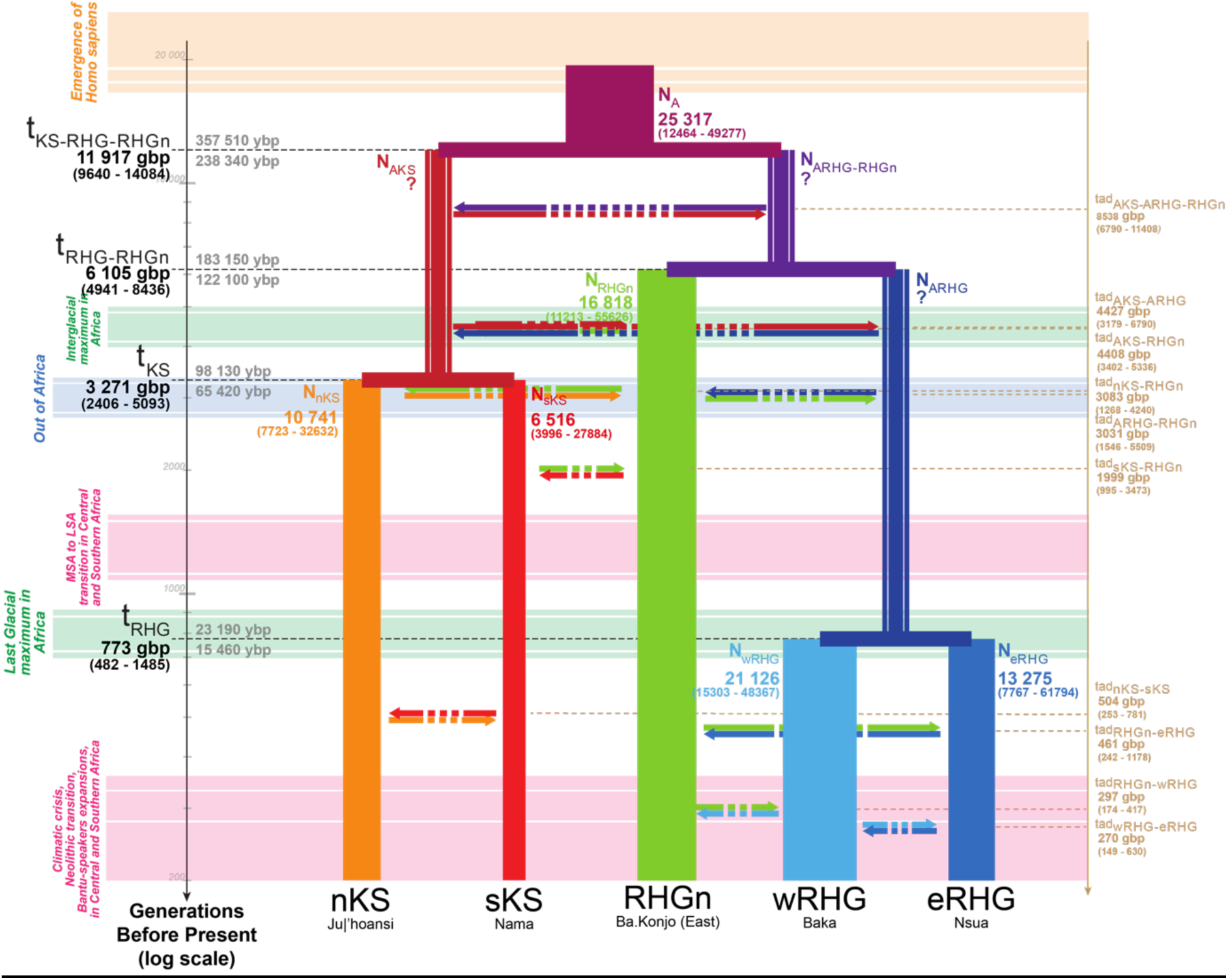
Schematic inferred demographic and migration history of Central and Southern African combination of populations n°11. Schematic representation of Scenario i1-1b (**Figure 4**), and Neural Network ABC posterior parameter mode estimates summarizing results obtained for the 11^th^ combination of sampled populations (**Table S13, Table S14**). Gene-flow arrows are indicated forward in time. For the time of each divergence and gene-flow event, mode point estimates are provided in generations before present (gbp) in bold, and 90% Credibility Intervals are provided between parentheses (**Table S14**). We provide two estimates of the divergence times estimates in years before present (ybp), one (upper) corresponding to 30 years per generation and the other (lower) to 20 years per generation ^66,67^. Mode point estimates of effective population sizes *Ne* are provided in numbers of diploid effective individuals and width of lineages are proportional to the estimated *Ne* (**Table S14**, **Figure S21**). NN-ABC posterior distributions for the effective population sizes of the ancestral Khoe-San lineage (AKS), for the ancestral RHG lineage (ARHG), and for the lineage ancestral to RHGn and ARHG, were all three poorly distinguished from their respective prior distributions and with high cross-validation posterior parameter estimation errors (**Figure S21**, **Table S16, Table S17**), as indicated by the question marks. NN-ABC posterior distributions for instantaneous asymmetric gene-flow rates were overall poorly departing from their priors and with high cross-validation posterior parameter estimation errors (**Figure S22**, **Table S14, Table S16**), as indicated by the doted arrows. All posterior distributions are shown graphically in **Figure 6**, **Figure 7, Figure S21, Figure S22**, and detailed in **Table S14**, with cross-validation posterior errors for each parameter in **Table S16**.

**Figure S34:**
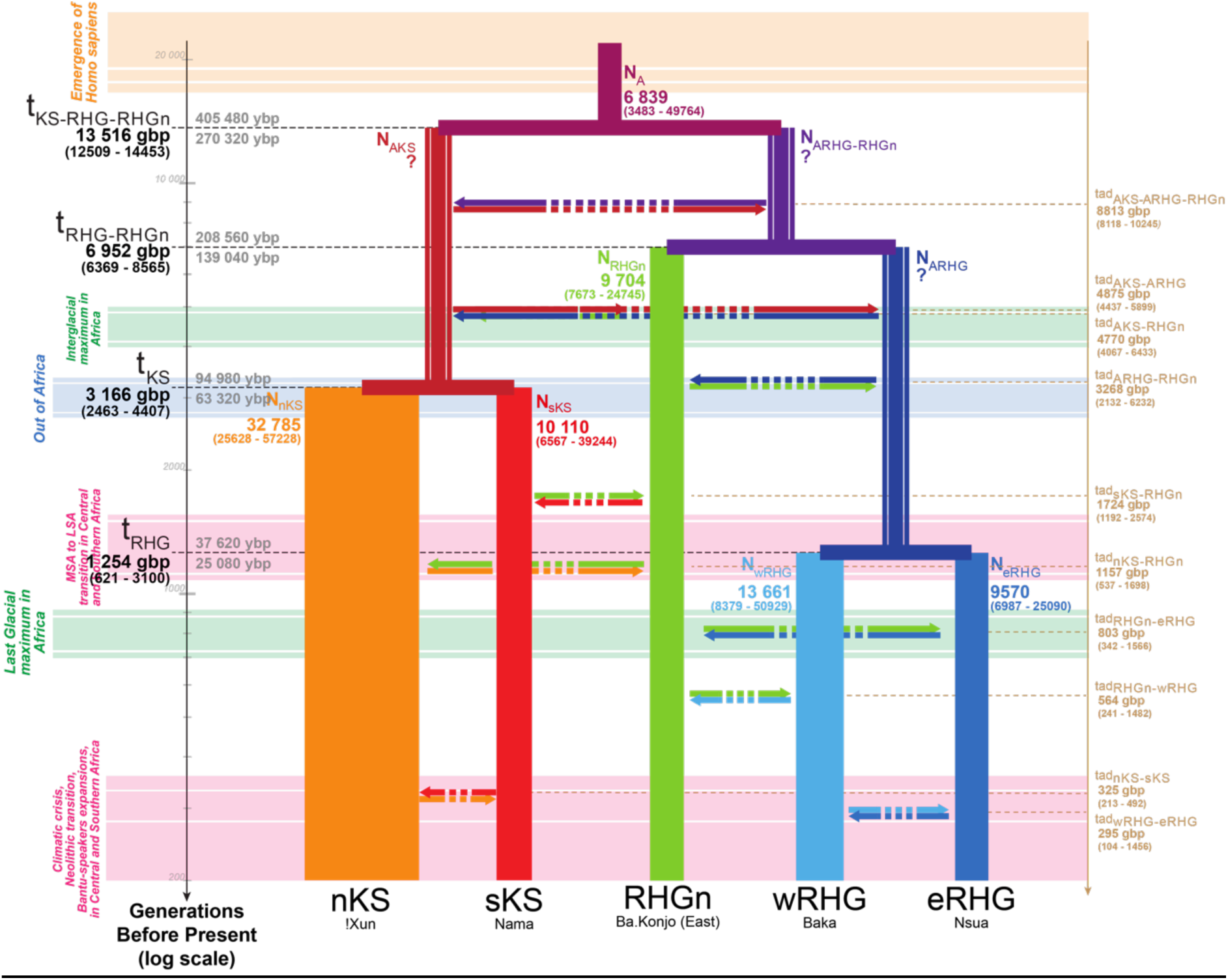
Schematic inferred demographic and migration history of Central and Southern African combination of populations n°12. Schematic representation of Scenario i1-1b (**Figure 4**), and Neural Network ABC posterior parameter mode estimates summarizing results obtained for the 12^th^ combination of sampled populations (**Table S13, Table S14**). Gene-flow arrows are indicated forward in time. For the time of each divergence and gene-flow event, mode point estimates are provided in generations before present (gbp) in bold, and 90% Credibility Intervals are provided between parentheses (**Table S14**). We provide two estimates of the divergence times estimates in years before present (ybp), one (upper) corresponding to 30 years per generation and the other (lower) to 20 years per generation ^66,67^. Mode point estimates of effective population sizes *Ne* are provided in numbers of diploid effective individuals and width of lineages are proportional to the estimated *Ne* (**Table S14**, **Figure S21**). NN-ABC posterior distributions for the effective population sizes of the ancestral Khoe-San lineage (AKS), for the ancestral RHG lineage (ARHG), and for the lineage ancestral to RHGn and ARHG, were all three poorly distinguished from their respective prior distributions and with high cross-validation posterior parameter estimation errors (**Figure S21**, **Table S16, Table S17**), as indicated by the question marks. NN-ABC posterior distributions for instantaneous asymmetric gene-flow rates were overall poorly departing from their priors and with high cross-validation posterior parameter estimation errors (**Figure S22**, **Table S14, Table S16**), as indicated by the doted arrows. All posterior distributions are shown graphically in **Figure 6**, **Figure 7, Figure S21, Figure S22**, and detailed in **Table S14**, with cross-validation posterior errors for each parameter in **Table S16**.

**Figure S35:**
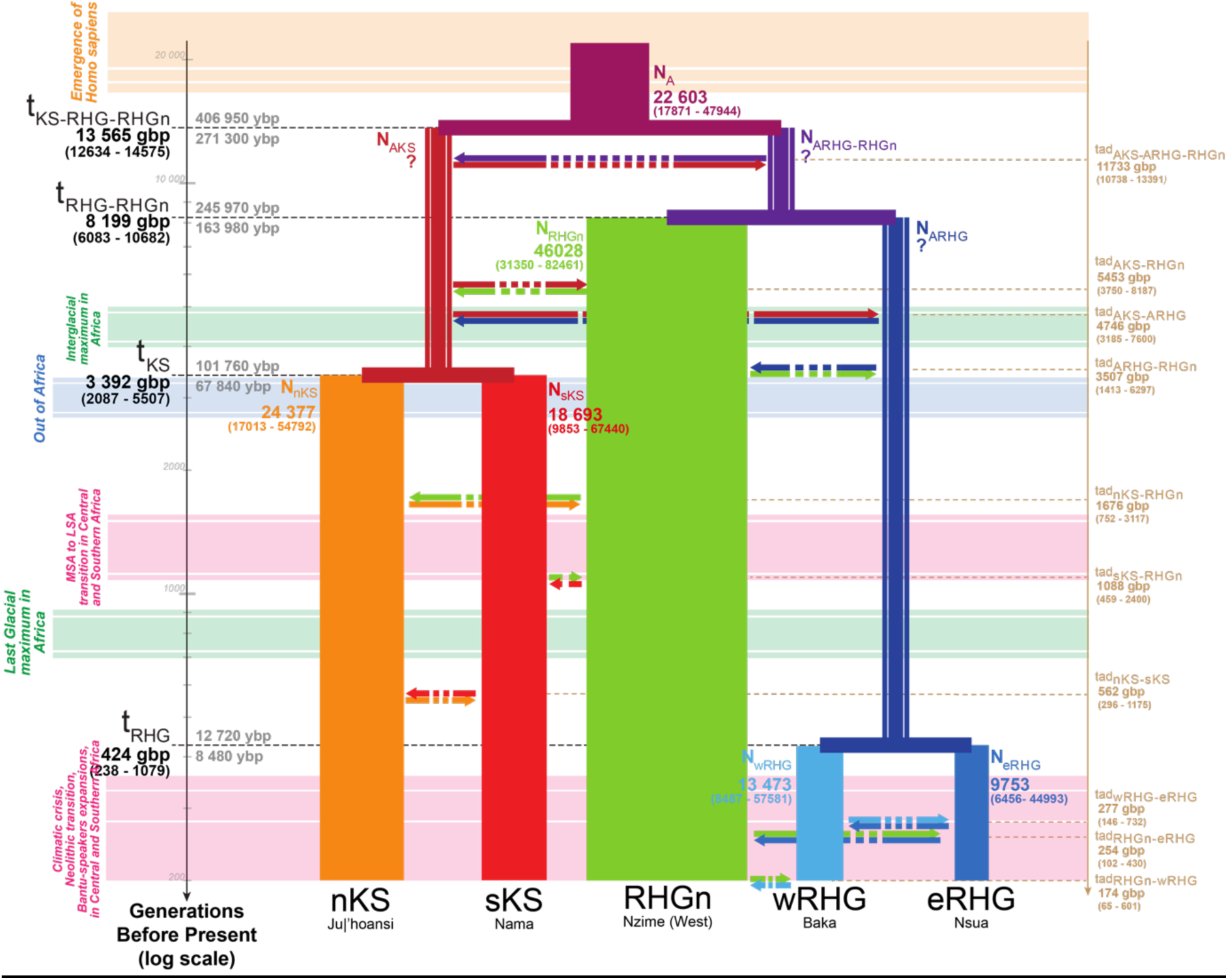
Schematic inferred demographic and migration history of Central and Southern African combination of populations n°13. Schematic representation of Scenario i1-1b (**Figure 4**), and Neural Network ABC posterior parameter mode estimates summarizing results obtained for the 13^th^ combination of sampled populations (**Table S13, Table S14**). Gene-flow arrows are indicated forward in time. For the time of each divergence and gene-flow event, mode point estimates are provided in generations before present (gbp) in bold, and 90% Credibility Intervals are provided between parentheses (**Table S14**). We provide two estimates of the divergence times estimates in years before present (ybp), one (upper) corresponding to 30 years per generation and the other (lower) to 20 years per generation ^66,67^. Mode point estimates of effective population sizes *Ne* are provided in numbers of diploid effective individuals and width of lineages are proportional to the estimated *Ne* (**Table S14**, **Figure S21**). NN-ABC posterior distributions for the effective population sizes of the ancestral Khoe-San lineage (AKS), for the ancestral RHG lineage (ARHG), and for the lineage ancestral to RHGn and ARHG, were all three poorly distinguished from their respective prior distributions and with high cross-validation posterior parameter estimation errors (**Figure S21**, **Table S16, Table S17**), as indicated by the question marks. NN-ABC posterior distributions for instantaneous asymmetric gene-flow rates were overall poorly departing from their priors and with high cross-validation posterior parameter estimation errors (**Figure S22**, **Table S14, Table S16**), as indicated by the doted arrows. All posterior distributions are shown graphically in **Figure 6**, **Figure 7, Figure S21, Figure S22**, and detailed in **Table S14**, with cross-validation posterior errors for each parameter in **Table S16**.

**Figure S36:**
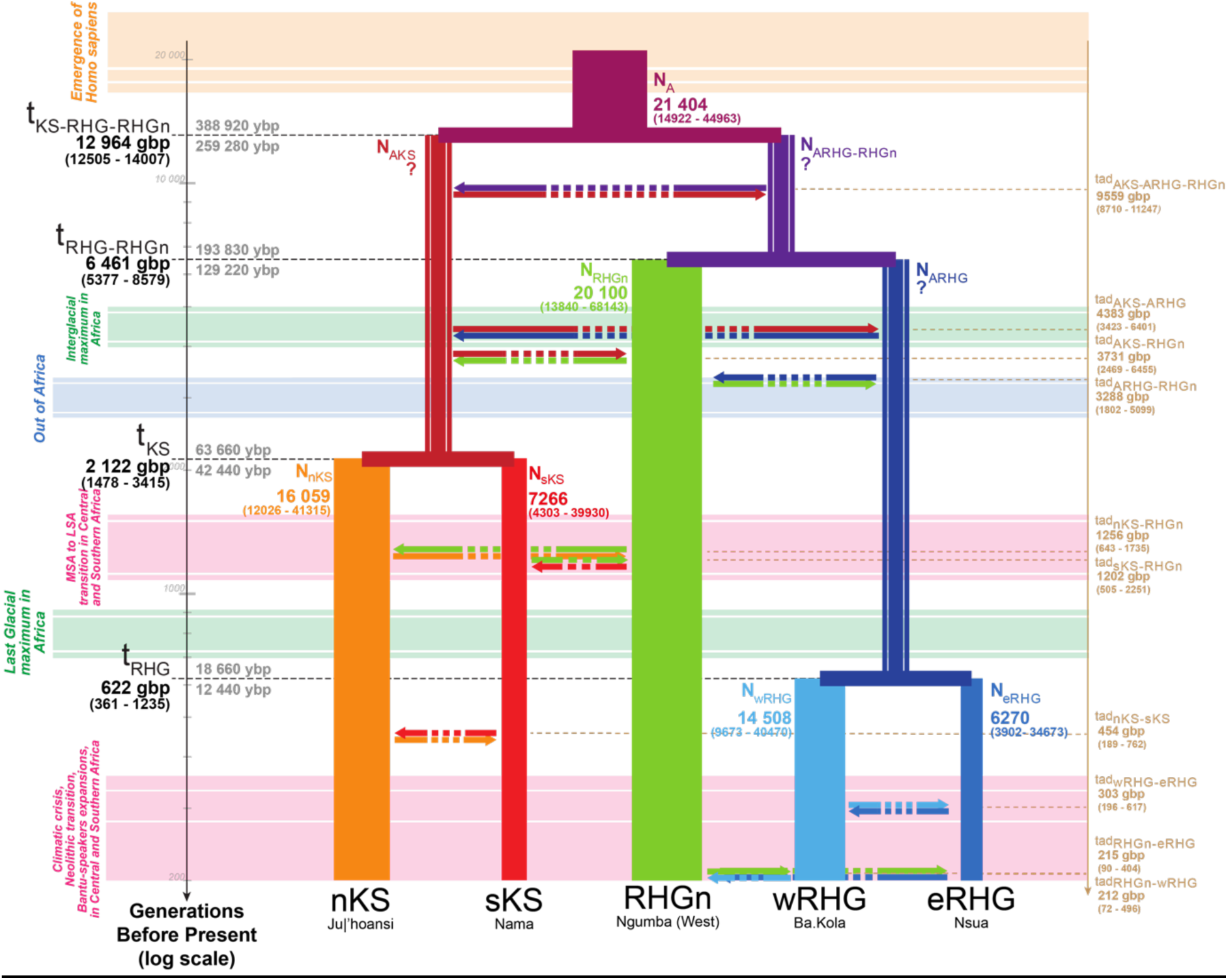
Schematic inferred demographic and migration history of Central and Southern African combination of populations n°14. Schematic representation of Scenario i1-1b (**Figure 4**), and Neural Network ABC posterior parameter mode estimates summarizing results obtained for the 14^th^ combination of sampled populations (**Table S13, Table S14**). Gene-flow arrows are indicated forward in time. For the time of each divergence and gene-flow event, mode point estimates are provided in generations before present (gbp) in bold, and 90% Credibility Intervals are provided between parentheses (**Table S14**). We provide two estimates of the divergence times estimates in years before present (ybp), one (upper) corresponding to 30 years per generation and the other (lower) to 20 years per generation ^66,67^. Mode point estimates of effective population sizes *Ne* are provided in numbers of diploid effective individuals and width of lineages are proportional to the estimated *Ne* (**Table S14**, **Figure S21**). NN-ABC posterior distributions for the effective population sizes of the ancestral Khoe-San lineage (AKS), for the ancestral RHG lineage (ARHG), and for the lineage ancestral to RHGn and ARHG, were all three poorly distinguished from their respective prior distributions and with high cross-validation posterior parameter estimation errors (**Figure S21**, **Table S16, Table S17**), as indicated by the question marks. NN-ABC posterior distributions for instantaneous asymmetric gene-flow rates were overall poorly departing from their priors and with high cross-validation posterior parameter estimation errors (**Figure S22**, **Table S14, Table S16**), as indicated by the doted arrows. All posterior distributions are shown graphically in **Figure 6**, **Figure 7, Figure S21, Figure S22**, and detailed in **Table S14**, with cross-validation posterior errors for each parameter in **Table S16**.

**Figure S37:**
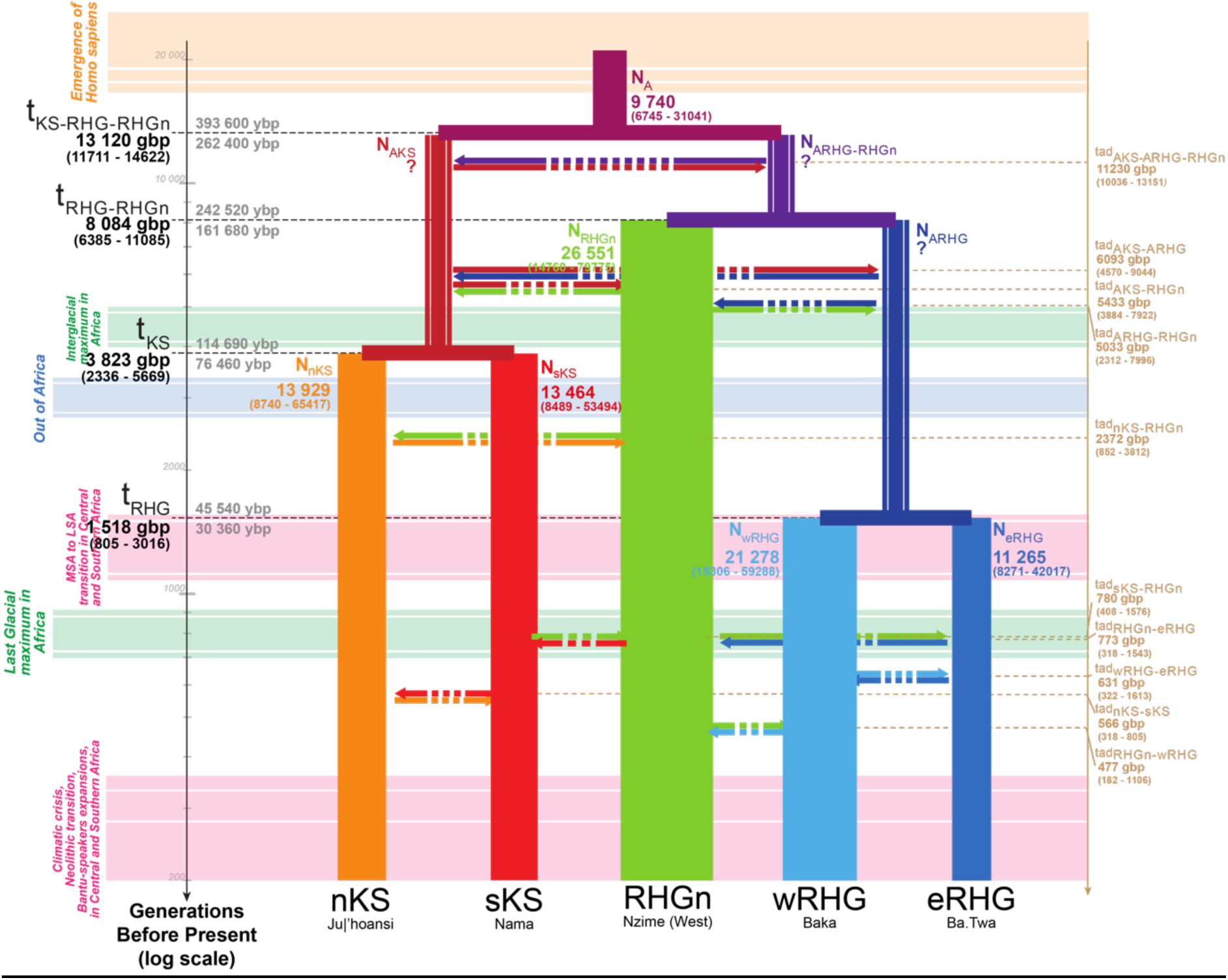
Schematic inferred demographic and migration history of Central and Southern African combination of populations n°15. Schematic representation of Scenario i1-1b (**Figure 4**), and Neural Network ABC posterior parameter mode estimates summarizing results obtained for the 15^th^ combination of sampled populations (**Table S13, Table S14**). Gene-flow arrows are indicated forward in time. For the time of each divergence and gene-flow event, mode point estimates are provided in generations before present (gbp) in bold, and 90% Credibility Intervals are provided between parentheses (**Table S14**). We provide two estimates of the divergence times estimates in years before present (ybp), one (upper) corresponding to 30 years per generation and the other (lower) to 20 years per generation ^66,67^. Mode point estimates of effective population sizes *Ne* are provided in numbers of diploid effective individuals and width of lineages are proportional to the estimated *Ne* (**Table S14**, **Figure S21**). NN-ABC posterior distributions for the effective population sizes of the ancestral Khoe-San lineage (AKS), for the ancestral RHG lineage (ARHG), and for the lineage ancestral to RHGn and ARHG, were all three poorly distinguished from their respective prior distributions and with high cross-validation posterior parameter estimation errors (**Figure S21**, **Table S16, Table S17**), as indicated by the question marks. NN-ABC posterior distributions for instantaneous asymmetric gene-flow rates were overall poorly departing from their priors and with high cross-validation posterior parameter estimation errors (**Figure S22**, **Table S14, Table S16**), as indicated by the doted arrows. All posterior distributions are shown graphically in **Figure 6**, **Figure 7, Figure S21, Figure S22**, and detailed in **Table S14**, with cross-validation posterior errors for each parameter in **Table S16**.

**Figure S38:**
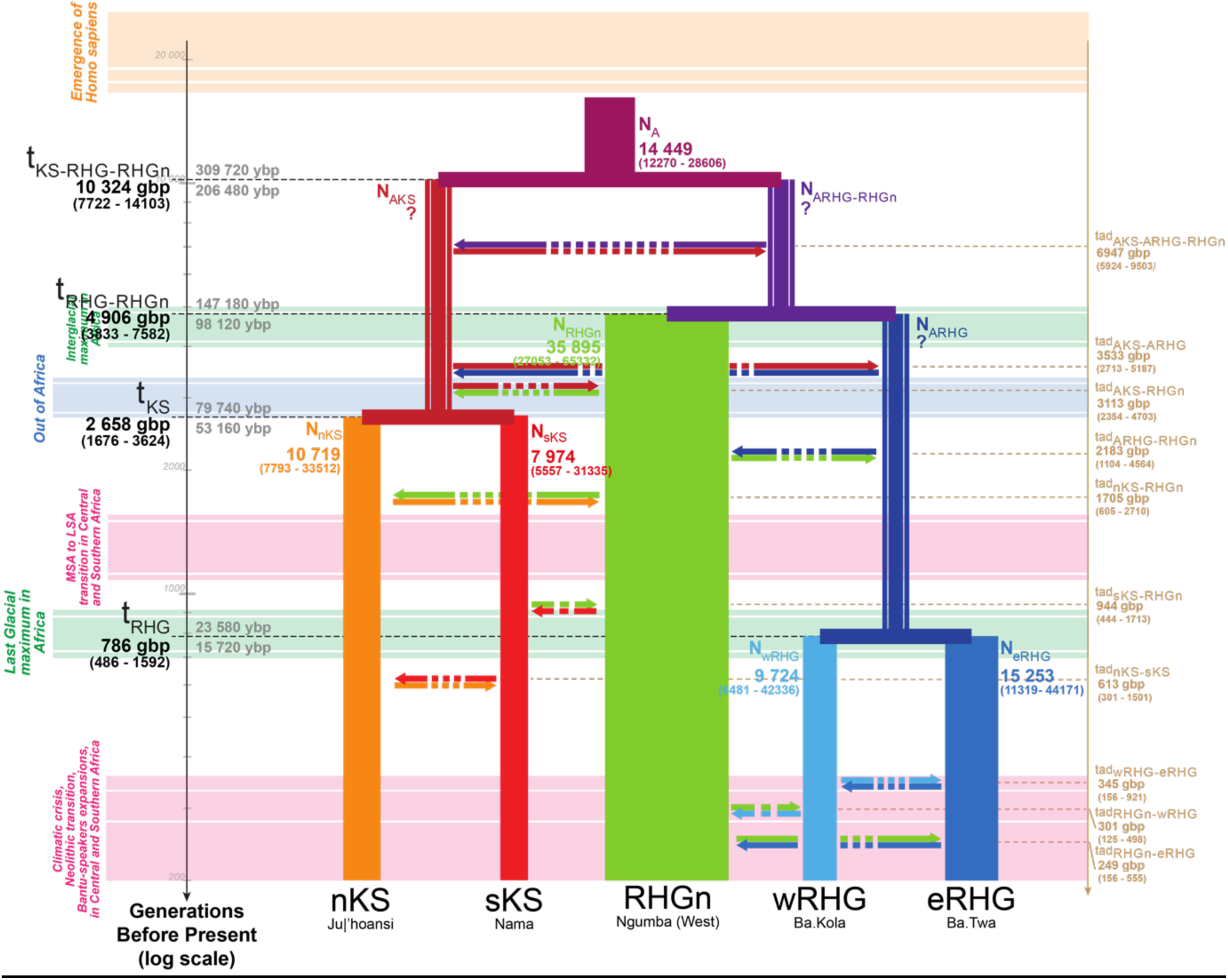
Schematic inferred demographic and migration history of Central and Southern African combination of populations n°16. Schematic representation of Scenario i1-1b (**Figure 4**), and Neural Network ABC posterior parameter mode estimates summarizing results obtained for the 16^th^ combination of sampled populations (**Table S13, Table S14**). Gene-flow arrows are indicated forward in time. For the time of each divergence and gene-flow event, mode point estimates are provided in generations before present (gbp) in bold, and 90% Credibility Intervals are provided between parentheses (**Table S14**). We provide two estimates of the divergence times estimates in years before present (ybp), one (upper) corresponding to 30 years per generation and the other (lower) to 20 years per generation ^66,67^. Mode point estimates of effective population sizes *Ne* are provided in numbers of diploid effective individuals and width of lineages are proportional to the estimated *Ne* (**Table S14**, **Figure S21**). NN-ABC posterior distributions for the effective population sizes of the ancestral Khoe-San lineage (AKS), for the ancestral RHG lineage (ARHG), and for the lineage ancestral to RHGn and ARHG, were all three poorly distinguished from their respective prior distributions and with high cross-validation posterior parameter estimation errors (**Figure S21**, **Table S16, Table S17**), as indicated by the question marks. NN-ABC posterior distributions for instantaneous asymmetric gene-flow rates were overall poorly departing from their priors and with high cross-validation posterior parameter estimation errors (**Figure S22**, **Table S14, Table S16**), as indicated by the doted arrows. All posterior distributions are shown graphically in **Figure 6**, **Figure 7, Figure S21, Figure S22**, and detailed in **Table S14**, with cross-validation posterior errors for each parameter in **Table S16**.

**Figure S39:**
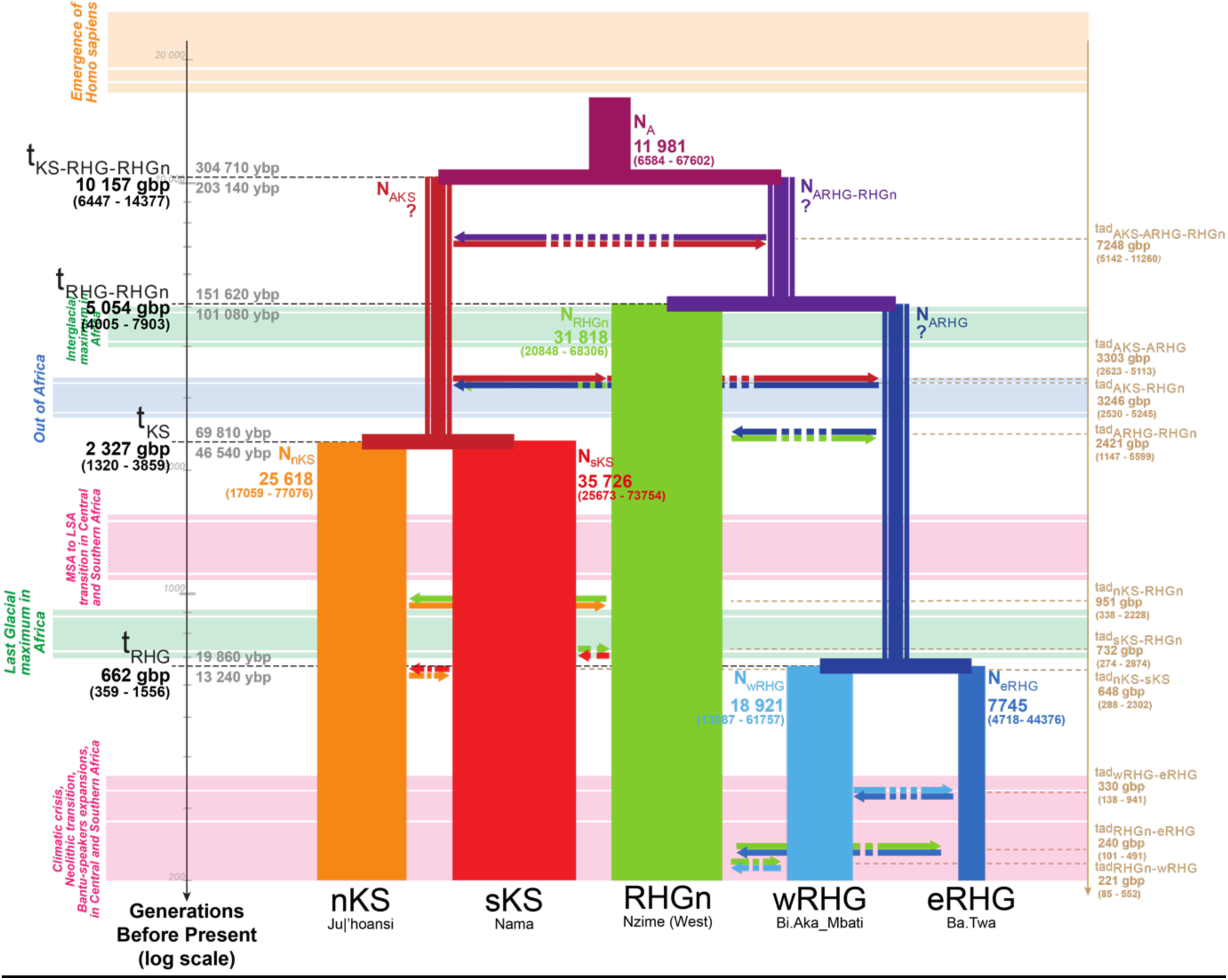
Schematic inferred demographic and migration history of Central and Southern African combination of populations n°17. Schematic representation of Scenario i1-1b (**Figure 4**), and Neural Network ABC posterior parameter mode estimates summarizing results obtained for the 17^th^ combination of sampled populations (**Table S13, Table S14**). Gene-flow arrows are indicated forward in time. For the time of each divergence and gene-flow event, mode point estimates are provided in generations before present (gbp) in bold, and 90% Credibility Intervals are provided between parentheses (**Table S14**). We provide two estimates of the divergence times estimates in years before present (ybp), one (upper) corresponding to 30 years per generation and the other (lower) to 20 years per generation ^66,67^. Mode point estimates of effective population sizes *Ne* are provided in numbers of diploid effective individuals and width of lineages are proportional to the estimated *Ne* (**Table S14**, **Figure S21**). NN-ABC posterior distributions for the effective population sizes of the ancestral Khoe-San lineage (AKS), for the ancestral RHG lineage (ARHG), and for the lineage ancestral to RHGn and ARHG, were all three poorly distinguished from their respective prior distributions and with high cross-validation posterior parameter estimation errors (**Figure S21**, **Table S16, Table S17**), as indicated by the question marks. NN-ABC posterior distributions for instantaneous asymmetric gene-flow rates were overall poorly departing from their priors and with high cross-validation posterior parameter estimation errors (**Figure S22**, **Table S14, Table S16**), as indicated by the doted arrows. All posterior distributions are shown graphically in **Figure 6**, **Figure 7, Figure S21, Figure S22**, and detailed in **Table S14**, with cross-validation posterior errors for each parameter in **Table S16**.

**Figure S40:**
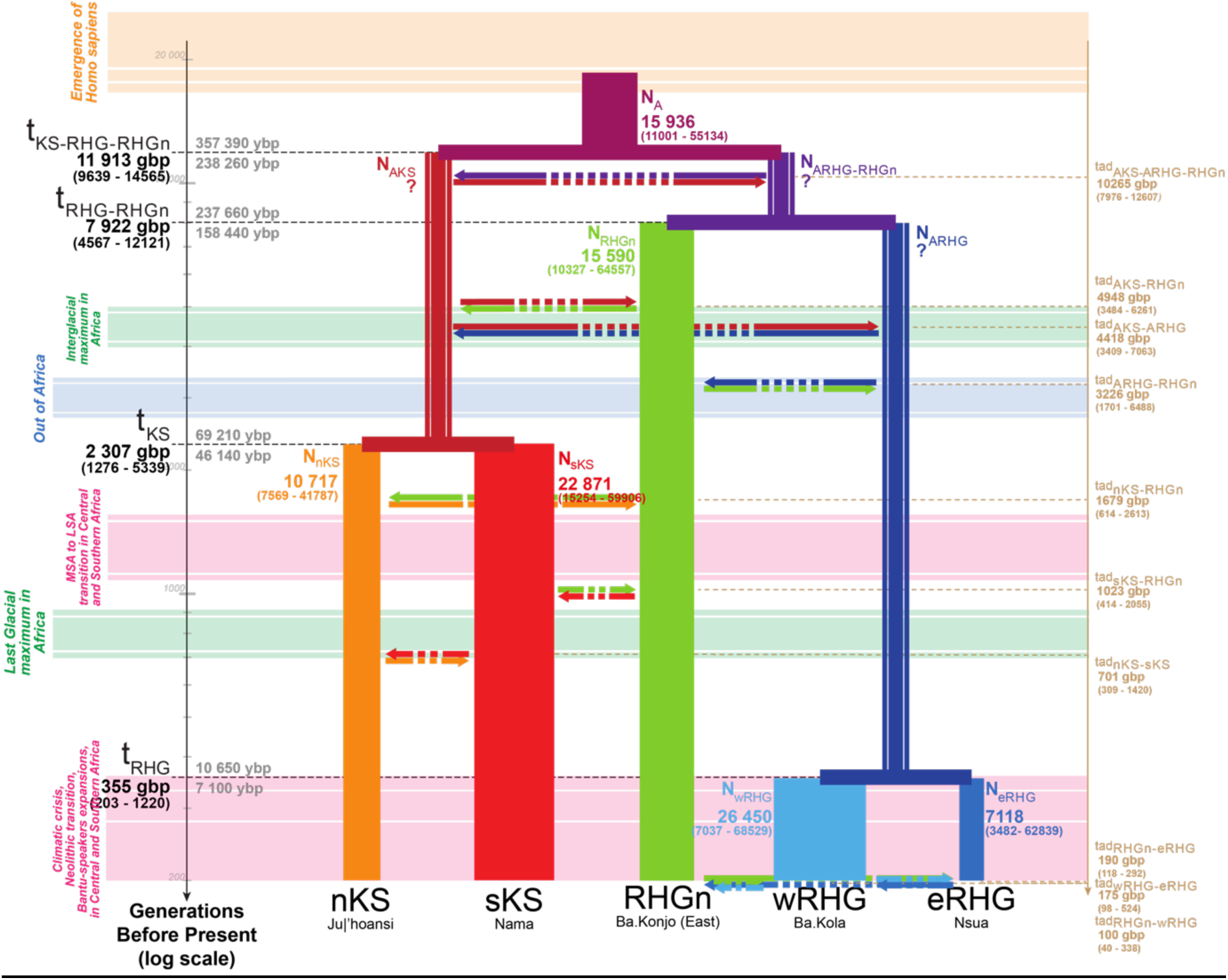
Schematic inferred demographic and migration history of Central and Southern African combination of populations n°18. Schematic representation of Scenario i1-1b (**Figure 4**), and Neural Network ABC posterior parameter mode estimates summarizing results obtained for the 18^th^ combination of sampled populations (**Table S13, Table S14**). Gene-flow arrows are indicated forward in time. For the time of each divergence and gene-flow event, mode point estimates are provided in generations before present (gbp) in bold, and 90% Credibility Intervals are provided between parentheses (**Table S14**). We provide two estimates of the divergence times estimates in years before present (ybp), one (upper) corresponding to 30 years per generation and the other (lower) to 20 years per generation ^66,67^. Mode point estimates of effective population sizes *Ne* are provided in numbers of diploid effective individuals and width of lineages are proportional to the estimated *Ne* (**Table S14**, **Figure S21**). NN-ABC posterior distributions for the effective population sizes of the ancestral Khoe-San lineage (AKS), for the ancestral RHG lineage (ARHG), and for the lineage ancestral to RHGn and ARHG, were all three poorly distinguished from their respective prior distributions and with high cross-validation posterior parameter estimation errors (**Figure S21**, **Table S16, Table S17**), as indicated by the question marks. NN-ABC posterior distributions for instantaneous asymmetric gene-flow rates were overall poorly departing from their priors and with high cross-validation posterior parameter estimation errors (**Figure S22**, **Table S14, Table S16**), as indicated by the doted arrows. All posterior distributions are shown graphically in **Figure 6**, **Figure 7, Figure S21, Figure S22**, and detailed in **Table S14**, with cross-validation posterior errors for each parameter in **Table S16**.

**Figure S41:**
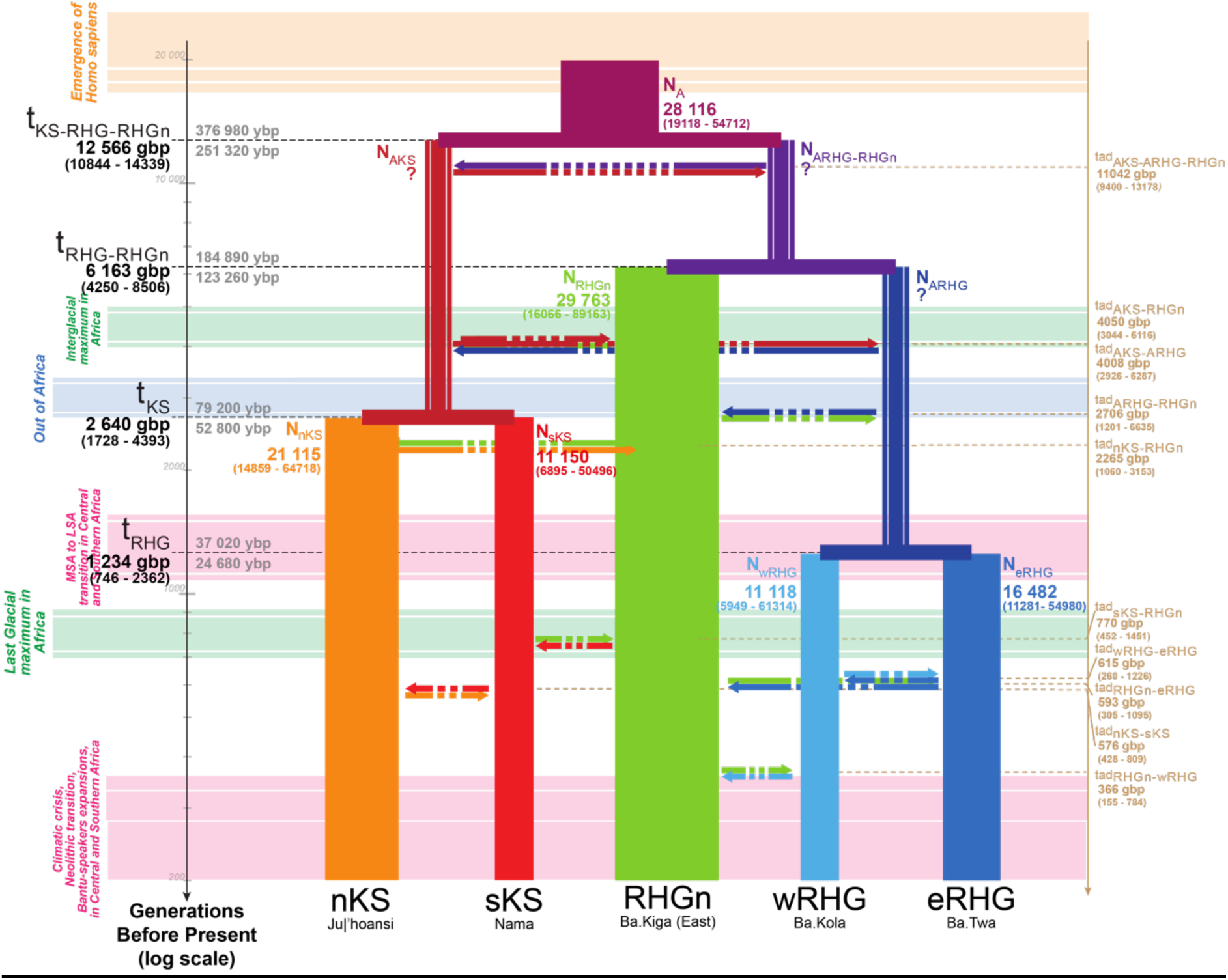
Schematic inferred demographic and migration history of Central and Southern African combination of populations n°19. Schematic representation of Scenario i1-1b (**Figure 4**), and Neural Network ABC posterior parameter mode estimates summarizing results obtained for the 19^th^ combination of sampled populations (**Table S13, Table S14**). Gene-flow arrows are indicated forward in time. For the time of each divergence and gene-flow event, mode point estimates are provided in generations before present (gbp) in bold, and 90% Credibility Intervals are provided between parentheses (**Table S14**). We provide two estimates of the divergence times estimates in years before present (ybp), one (upper) corresponding to 30 years per generation and the other (lower) to 20 years per generation ^66,67^. Mode point estimates of effective population sizes *Ne* are provided in numbers of diploid effective individuals and width of lineages are proportional to the estimated *Ne* (**Table S14**, **Figure S21**). NN-ABC posterior distributions for the effective population sizes of the ancestral Khoe-San lineage (AKS), for the ancestral RHG lineage (ARHG), and for the lineage ancestral to RHGn and ARHG, were all three poorly distinguished from their respective prior distributions and with high cross-validation posterior parameter estimation errors (**Figure S21**, **Table S16, Table S17**), as indicated by the question marks. NN-ABC posterior distributions for instantaneous asymmetric gene-flow rates were overall poorly departing from their priors and with high cross-validation posterior parameter estimation errors (**Figure S22**, **Table S14, Table S16**), as indicated by the doted arrows. All posterior distributions are shown graphically in **Figure 6**, **Figure 7, Figure S21, Figure S22**, and detailed in **Table S14**, with cross-validation posterior errors for each parameter in **Table S16**.

**Figure S42:**
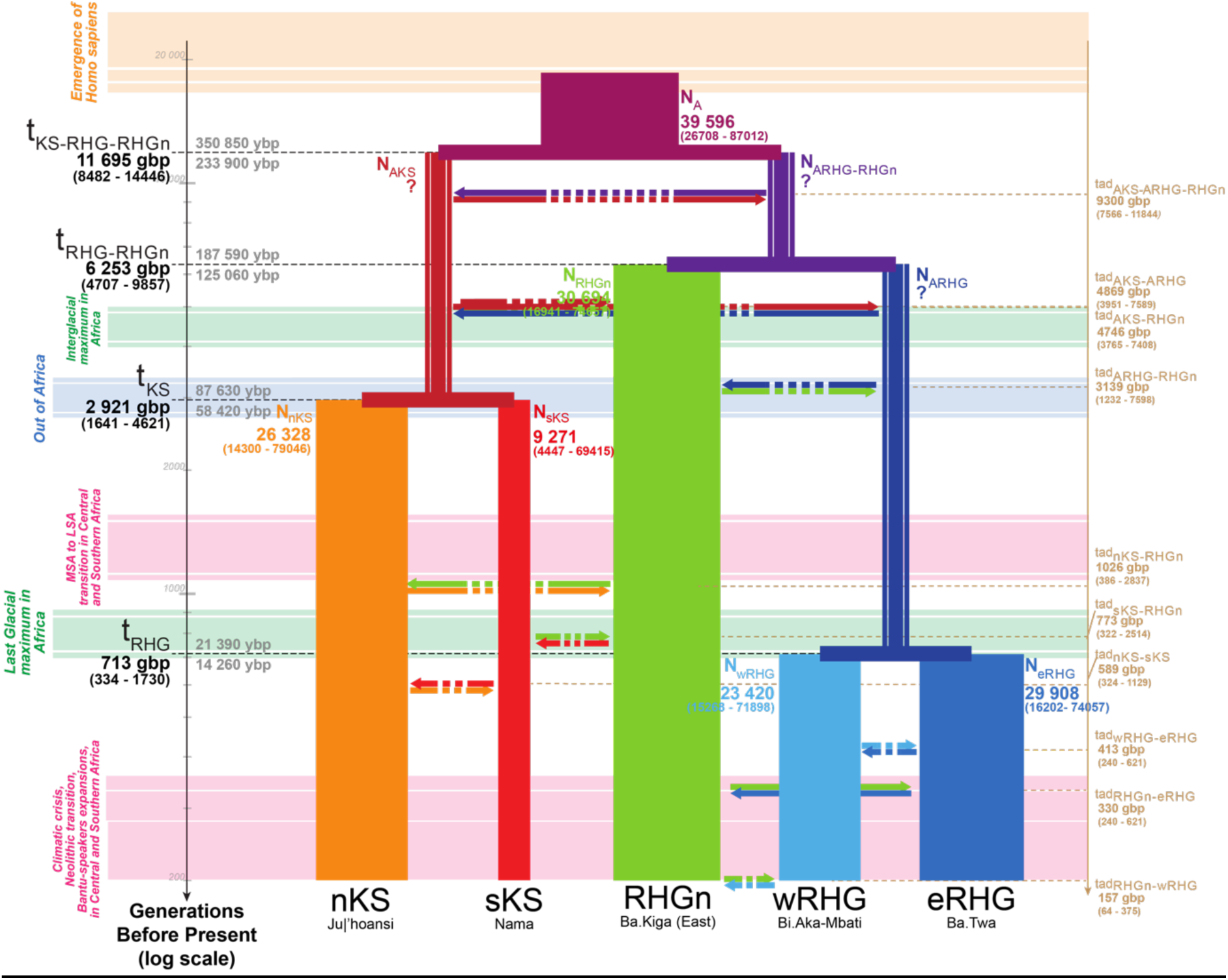
Schematic inferred demographic and migration history of Central and Southern African combination of populations n°20. Schematic representation of Scenario i1-1b (**Figure 4**), and Neural Network ABC posterior parameter mode estimates summarizing results obtained for the 20^th^ combination of sampled populations (**Table S13, Table S14**). Gene-flow arrows are indicated forward in time. For the time of each divergence and gene-flow event, mode point estimates are provided in generations before present (gbp) in bold, and 90% Credibility Intervals are provided between parentheses (**Table S14**). We provide two estimates of the divergence times estimates in years before present (ybp), one (upper) corresponding to 30 years per generation and the other (lower) to 20 years per generation ^66,67^. Mode point estimates of effective population sizes *Ne* are provided in numbers of diploid effective individuals and width of lineages are proportional to the estimated *Ne* (**Table S14**, **Figure S21**). NN-ABC posterior distributions for the effective population sizes of the ancestral Khoe-San lineage (AKS), for the ancestral RHG lineage (ARHG), and for the lineage ancestral to RHGn and ARHG, were all three poorly distinguished from their respective prior distributions and with high cross-validation posterior parameter estimation errors (**Figure S21**, **Table S16, Table S17**), as indicated by the question marks. NN-ABC posterior distributions for instantaneous asymmetric gene-flow rates were overall poorly departing from their priors and with high cross-validation posterior parameter estimation errors (**Figure S22**, **Table S14, Table S16**), as indicated by the doted arrows. All posterior distributions are shown graphically in **Figure 6**, **Figure 7, Figure S21, Figure S22**, and detailed in **Table S14**, with cross-validation posterior errors for each parameter in **Table S16**.

**Figure S43:**
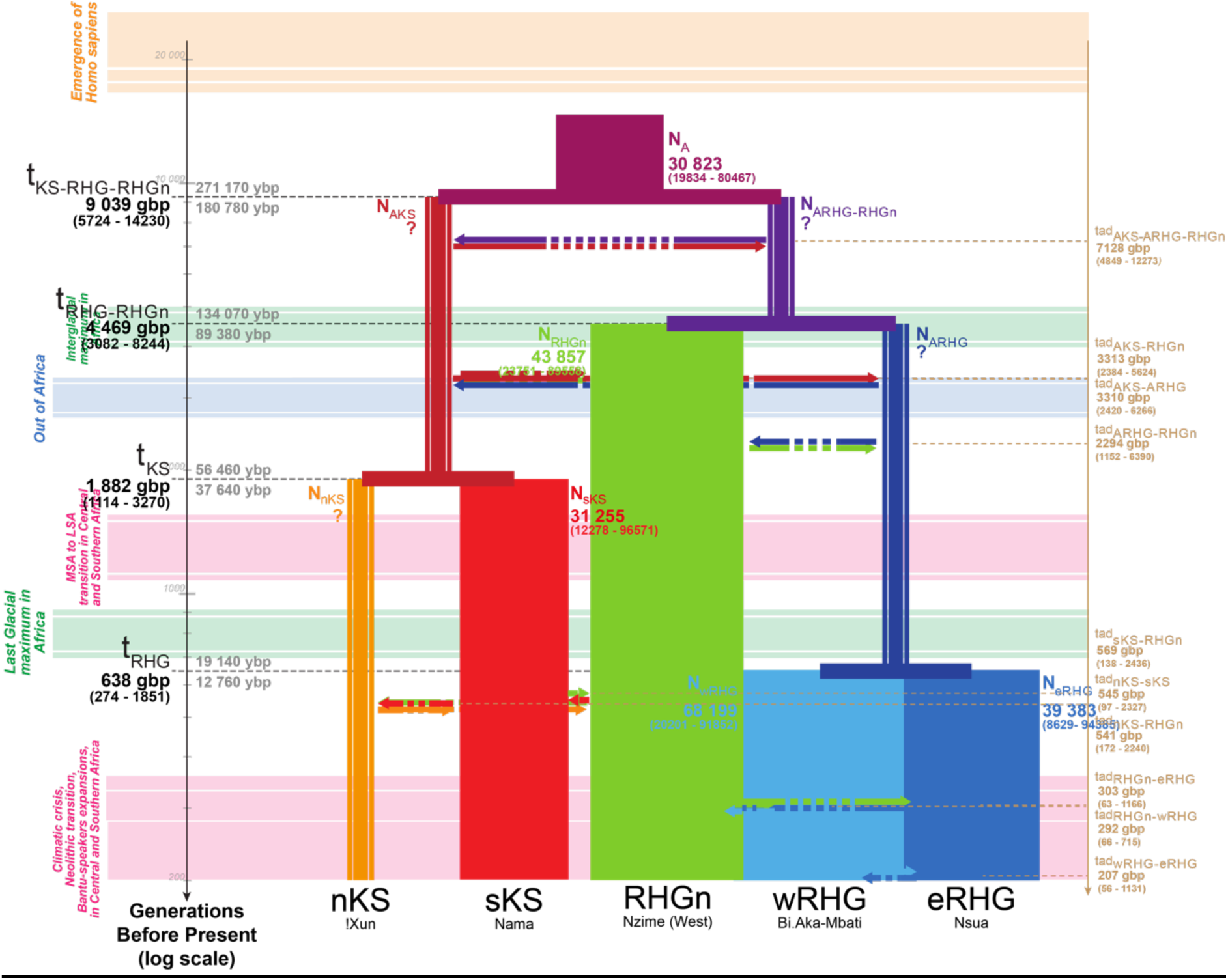
Schematic inferred demographic and migration history of Central and Southern African combination of populations n°21. Schematic representation of Scenario i1-1b (**Figure 4**), and Neural Network ABC posterior parameter mode estimates summarizing results obtained for the 21^th^ combination of sampled populations (**Table S13, Table S14**). Gene-flow arrows are indicated forward in time. For the time of each divergence and gene-flow event, mode point estimates are provided in generations before present (gbp) in bold, and 90% Credibility Intervals are provided between parentheses (**Table S14**). We provide two estimates of the divergence times estimates in years before present (ybp), one (upper) corresponding to 30 years per generation and the other (lower) to 20 years per generation ^66,67^. Mode point estimates of effective population sizes *Ne* are provided in numbers of diploid effective individuals and width of lineages are proportional to the estimated *Ne* (**Table S14**, **Figure S21**). NN-ABC posterior distributions for the effective population sizes of the ancestral Khoe-San lineage (AKS), for the ancestral RHG lineage (ARHG), and for the lineage ancestral to RHGn and ARHG, were all three poorly distinguished from their respective prior distributions and with high cross-validation posterior parameter estimation errors (**Figure S21**, **Table S16, Table S17**), as indicated by the question marks. NN-ABC posterior distributions for instantaneous asymmetric gene-flow rates were overall poorly departing from their priors and with high cross-validation posterior parameter estimation errors (**Figure S22**, **Table S14, Table S16**), as indicated by the doted arrows. All posterior distributions are shown graphically in **Figure 6**, **Figure 7, Figure S21, Figure S22**, and detailed in **Table S14**, with cross-validation posterior errors for each parameter in **Table S16**.

**Figure S44:**
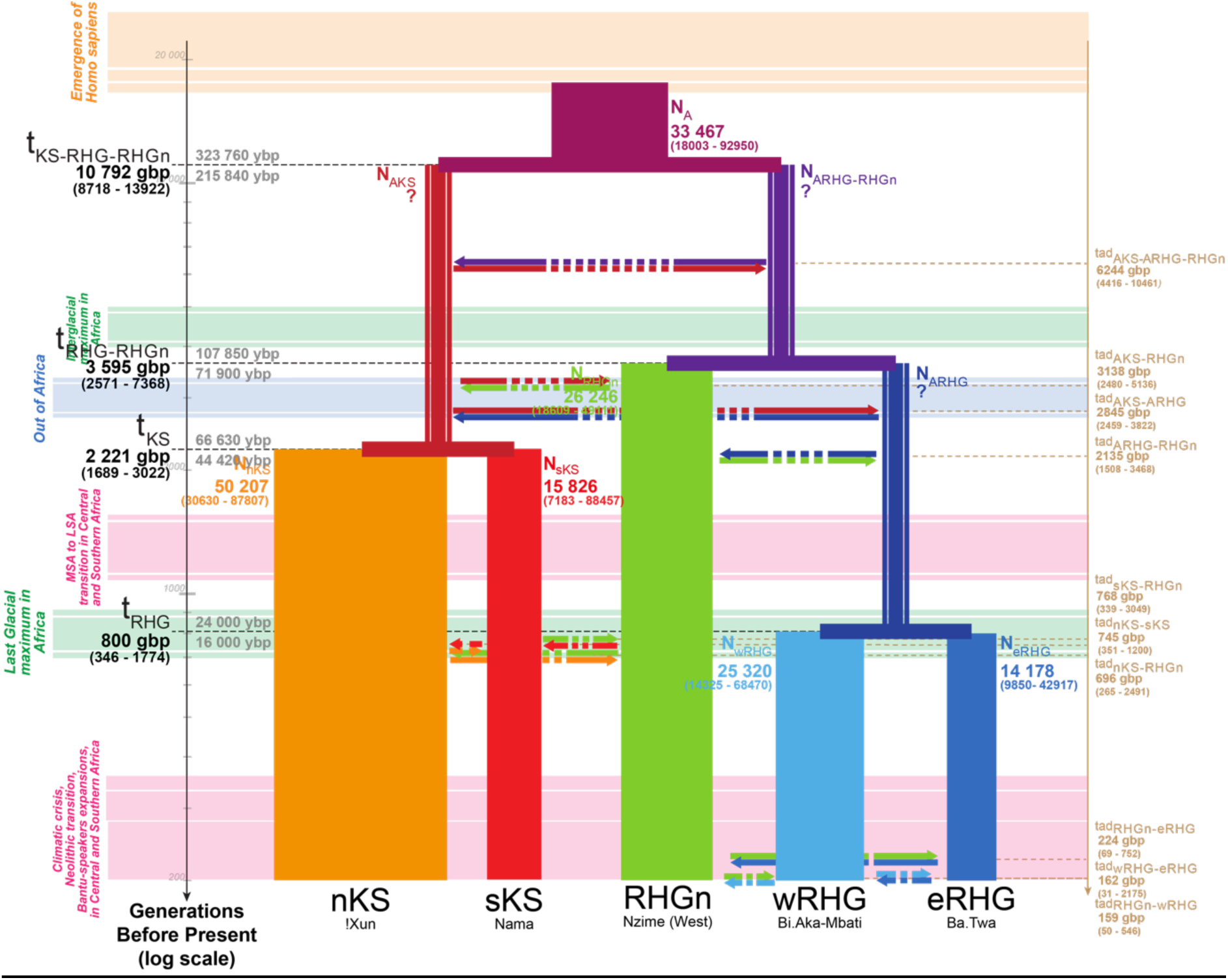
Schematic inferred demographic and migration history of Central and Southern African combination of populations n°22. Schematic representation of Scenario i1-1b (**Figure 4**), and Neural Network ABC posterior parameter mode estimates summarizing results obtained for the 22^th^ combination of sampled populations (**Table S13, Table S14**). Gene-flow arrows are indicated forward in time. For the time of each divergence and gene-flow event, mode point estimates are provided in generations before present (gbp) in bold, and 90% Credibility Intervals are provided between parentheses (**Table S14**). We provide two estimates of the divergence times estimates in years before present (ybp), one (upper) corresponding to 30 years per generation and the other (lower) to 20 years per generation ^66,67^. Mode point estimates of effective population sizes *Ne* are provided in numbers of diploid effective individuals and width of lineages are proportional to the estimated *Ne* (**Table S14**, **Figure S21**). NN-ABC posterior distributions for the effective population sizes of the ancestral Khoe-San lineage (AKS), for the ancestral RHG lineage (ARHG), and for the lineage ancestral to RHGn and ARHG, were all three poorly distinguished from their respective prior distributions and with high cross-validation posterior parameter estimation errors (**Figure S21**, **Table S16, Table S17**), as indicated by the question marks. NN-ABC posterior distributions for instantaneous asymmetric gene-flow rates were overall poorly departing from their priors and with high cross-validation posterior parameter estimation errors (**Figure S22**, **Table S14, Table S16**), as indicated by the doted arrows. All posterior distributions are shown graphically in **Figure 6**, **Figure 7, Figure S21, Figure S22**, and detailed in **Table S14**, with cross-validation posterior errors for each parameter in **Table S16**.

**Figure S45:**
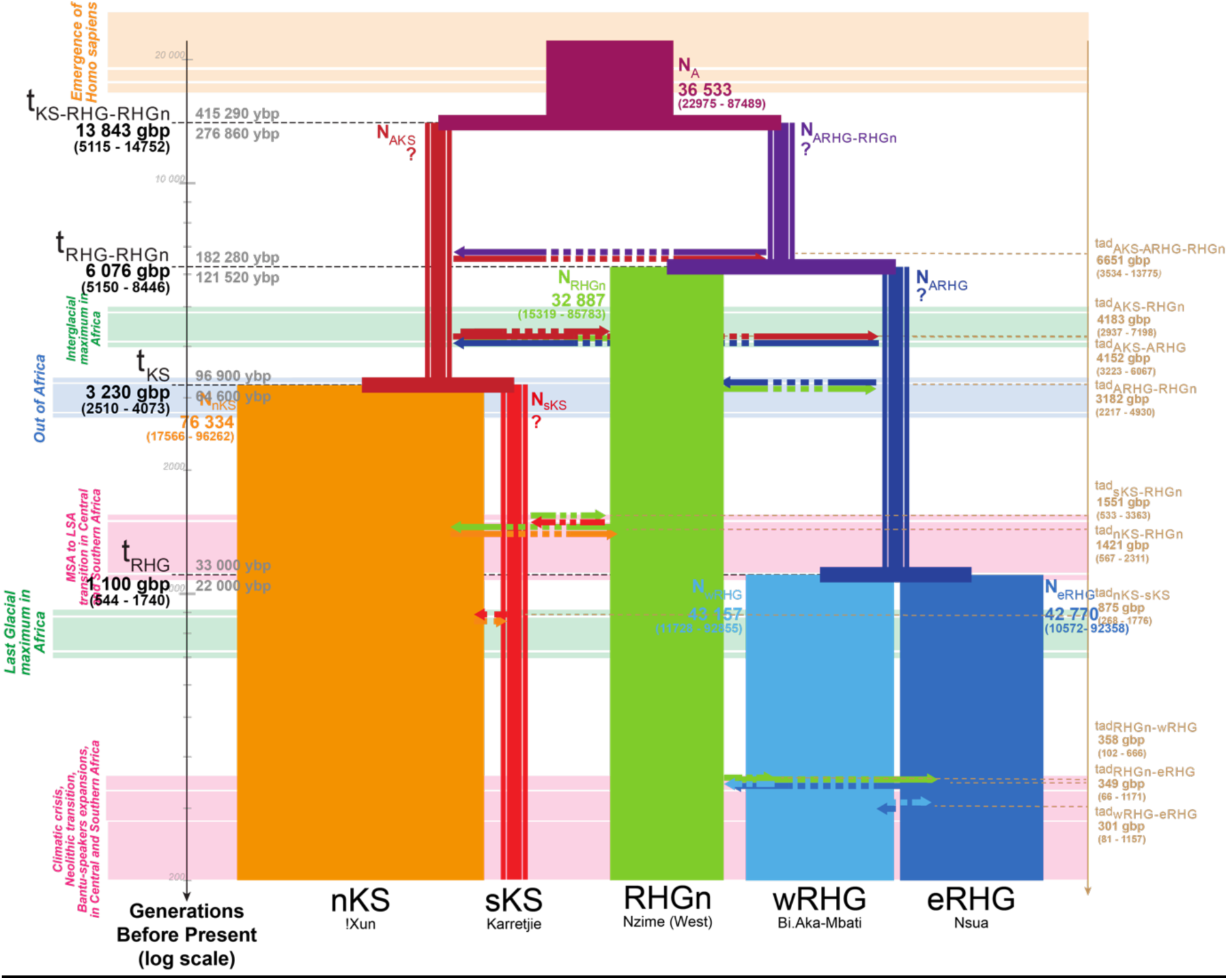
Schematic inferred demographic and migration history of Central and Southern African combination of populations n°23. Schematic representation of Scenario i1-1b (**Figure 4**), and Neural Network ABC posterior parameter mode estimates summarizing results obtained for the 23^th^ combination of sampled populations (**Table S13, Table S14**). Gene-flow arrows are indicated forward in time. For the time of each divergence and gene-flow event, mode point estimates are provided in generations before present (gbp) in bold, and 90% Credibility Intervals are provided between parentheses (**Table S14**). We provide two estimates of the divergence times estimates in years before present (ybp), one (upper) corresponding to 30 years per generation and the other (lower) to 20 years per generation ^66,67^. Mode point estimates of effective population sizes *Ne* are provided in numbers of diploid effective individuals and width of lineages are proportional to the estimated *Ne* (**Table S14**, **Figure S21**). NN-ABC posterior distributions for the effective population sizes of the ancestral Khoe-San lineage (AKS), for the ancestral RHG lineage (ARHG), and for the lineage ancestral to RHGn and ARHG, were all three poorly distinguished from their respective prior distributions and with high cross-validation posterior parameter estimation errors (**Figure S21**, **Table S16, Table S17**), as indicated by the question marks. NN-ABC posterior distributions for instantaneous asymmetric gene-flow rates were overall poorly departing from their priors and with high cross-validation posterior parameter estimation errors (**Figure S22**, **Table S14, Table S16**), as indicated by the doted arrows. All posterior distributions are shown graphically in **Figure 6**, **Figure 7, Figure S21, Figure S22**, and detailed in **Table S14**, with cross-validation posterior errors for each parameter in **Table S16**.

**Figure S46:**
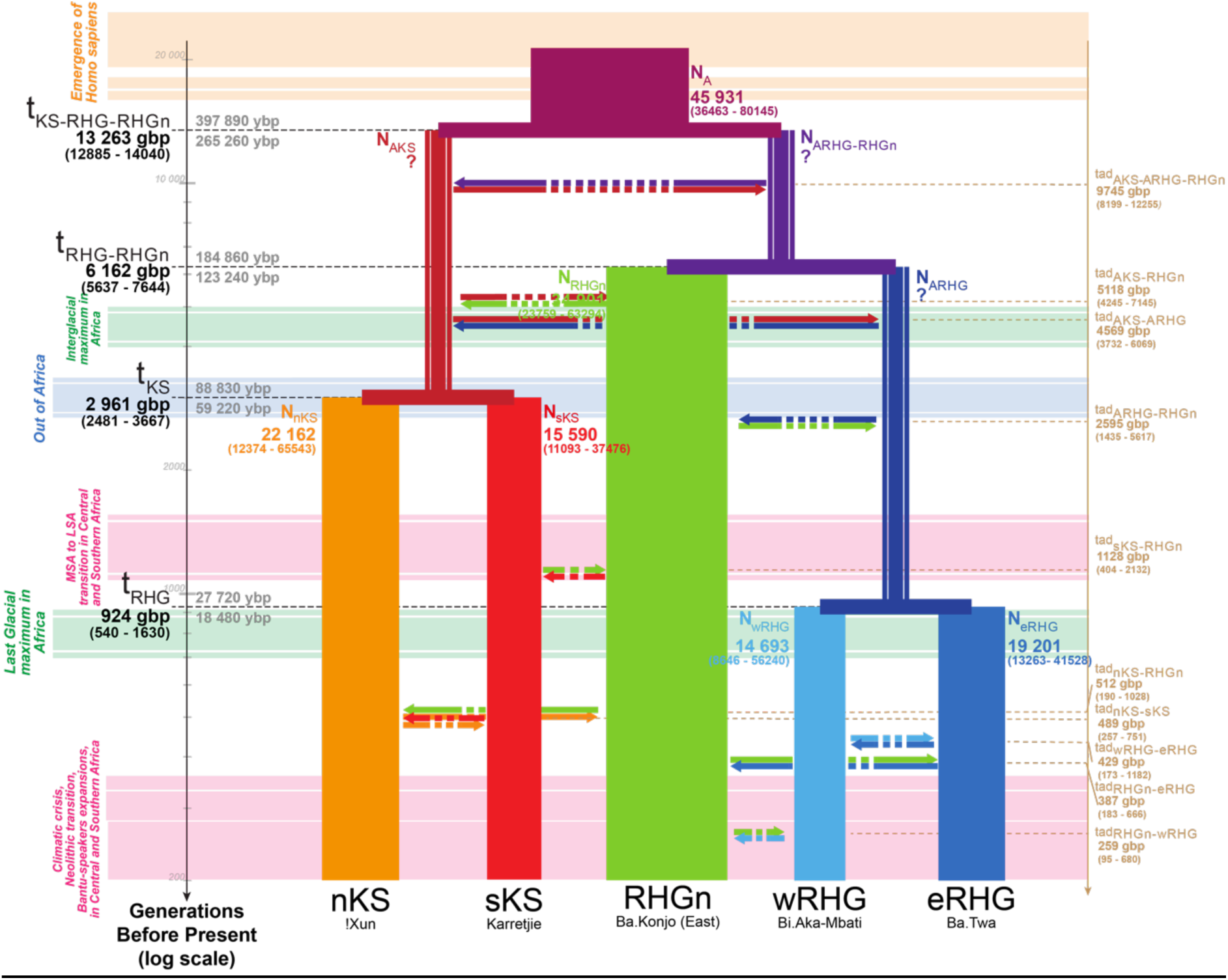
Schematic inferred demographic and migration history of Central and Southern African combination of populations n°24. Schematic representation of Scenario i1-1b (**Figure 4**), and Neural Network ABC posterior parameter mode estimates summarizing results obtained for the 24^th^ combination of sampled populations (**Table S13, Table S14**). Gene-flow arrows are indicated forward in time. For the time of each divergence and gene-flow event, mode point estimates are provided in generations before present (gbp) in bold, and 90% Credibility Intervals are provided between parentheses (**Table S14**). We provide two estimates of the divergence times estimates in years before present (ybp), one (upper) corresponding to 30 years per generation and the other (lower) to 20 years per generation ^66,67^. Mode point estimates of effective population sizes *Ne* are provided in numbers of diploid effective individuals and width of lineages are proportional to the estimated *Ne* (**Table S14**, **Figure S21**). NN-ABC posterior distributions for the effective population sizes of the ancestral Khoe-San lineage (AKS), for the ancestral RHG lineage (ARHG), and for the lineage ancestral to RHGn and ARHG, were all three poorly distinguished from their respective prior distributions and with high cross-validation posterior parameter estimation errors (**Figure S21**, **Table S16, Table S17**), as indicated by the question marks. NN-ABC posterior distributions for instantaneous asymmetric gene-flow rates were overall poorly departing from their priors and with high cross-validation posterior parameter estimation errors (**Figure S22**, **Table S14, Table S16**), as indicated by the doted arrows. All posterior distributions are shown graphically in **Figure 6**, **Figure 7, Figure S21, Figure S22**, and detailed in **Table S14**, with cross-validation posterior errors for each parameter in **Table S16**.

**Figure S47:**
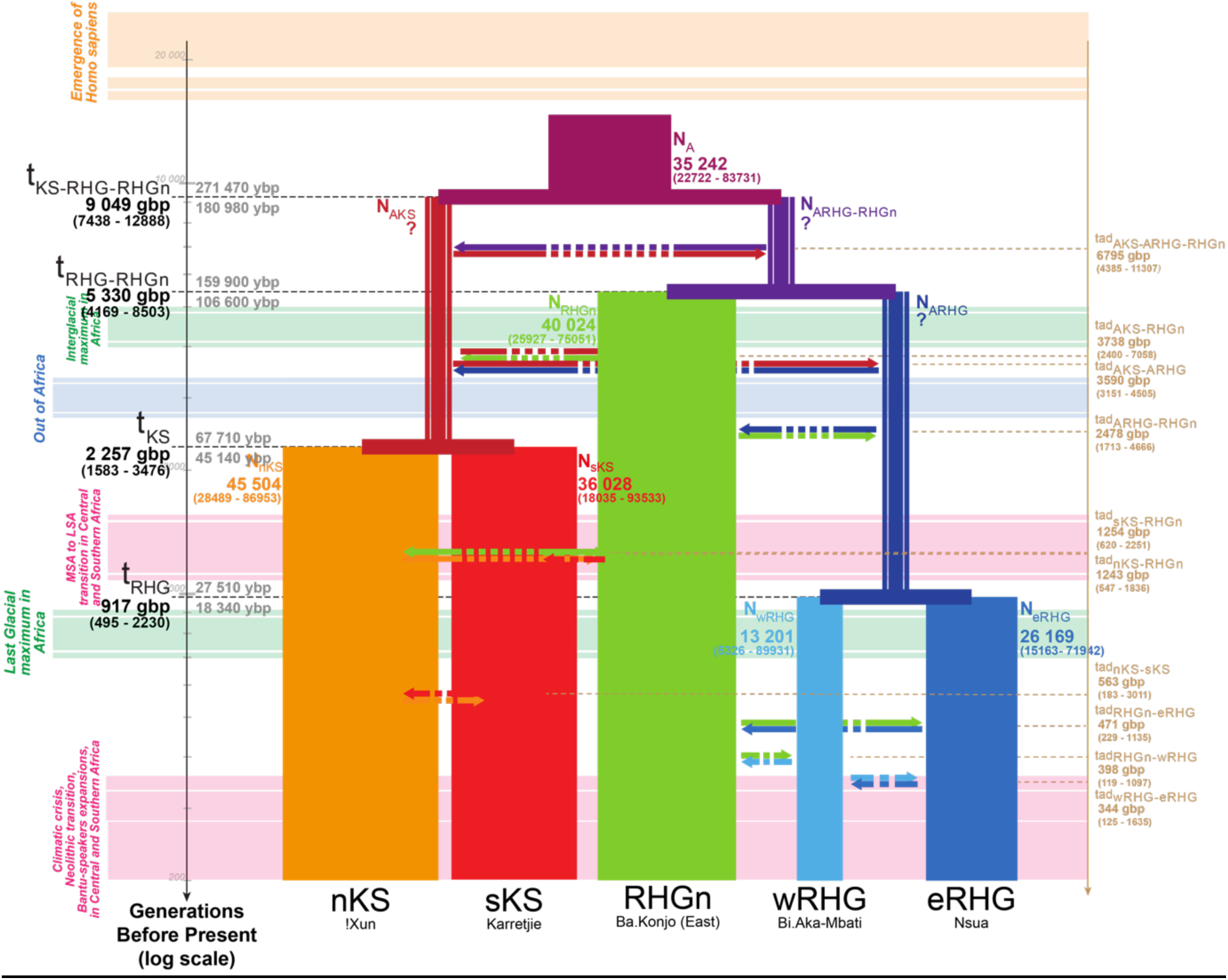
Schematic inferred demographic and migration history of Central and Southern African combination of populations n°25. Schematic representation of Scenario i1-1b (**Figure 4**), and Neural Network ABC posterior parameter mode estimates summarizing results obtained for the 25^th^ combination of sampled populations (**Table S13, Table S14**). Gene-flow arrows are indicated forward in time. For the time of each divergence and gene-flow event, mode point estimates are provided in generations before present (gbp) in bold, and 90% Credibility Intervals are provided between parentheses (**Table S14**). We provide two estimates of the divergence times estimates in years before present (ybp), one (upper) corresponding to 30 years per generation and the other (lower) to 20 years per generation ^66,67^. Mode point estimates of effective population sizes *Ne* are provided in numbers of diploid effective individuals and width of lineages are proportional to the estimated *Ne* (**Table S14**, **Figure S21**). NN-ABC posterior distributions for the effective population sizes of the ancestral Khoe-San lineage (AKS), for the ancestral RHG lineage (ARHG), and for the lineage ancestral to RHGn and ARHG, were all three poorly distinguished from their respective prior distributions and with high cross-validation posterior parameter estimation errors (**Figure S21**, **Table S16, Table S17**), as indicated by the question marks. NN-ABC posterior distributions for instantaneous asymmetric gene-flow rates were overall poorly departing from their priors and with high cross-validation posterior parameter estimation errors (**Figure S22**, **Table S14, Table S16**), as indicated by the doted arrows. All posterior distributions are shown graphically in **Figure 6**, **Figure 7, Figure S21, Figure S22**, and detailed in **Table S14**, with cross-validation posterior errors for each parameter in **Table S16**.

**Figure S48:**
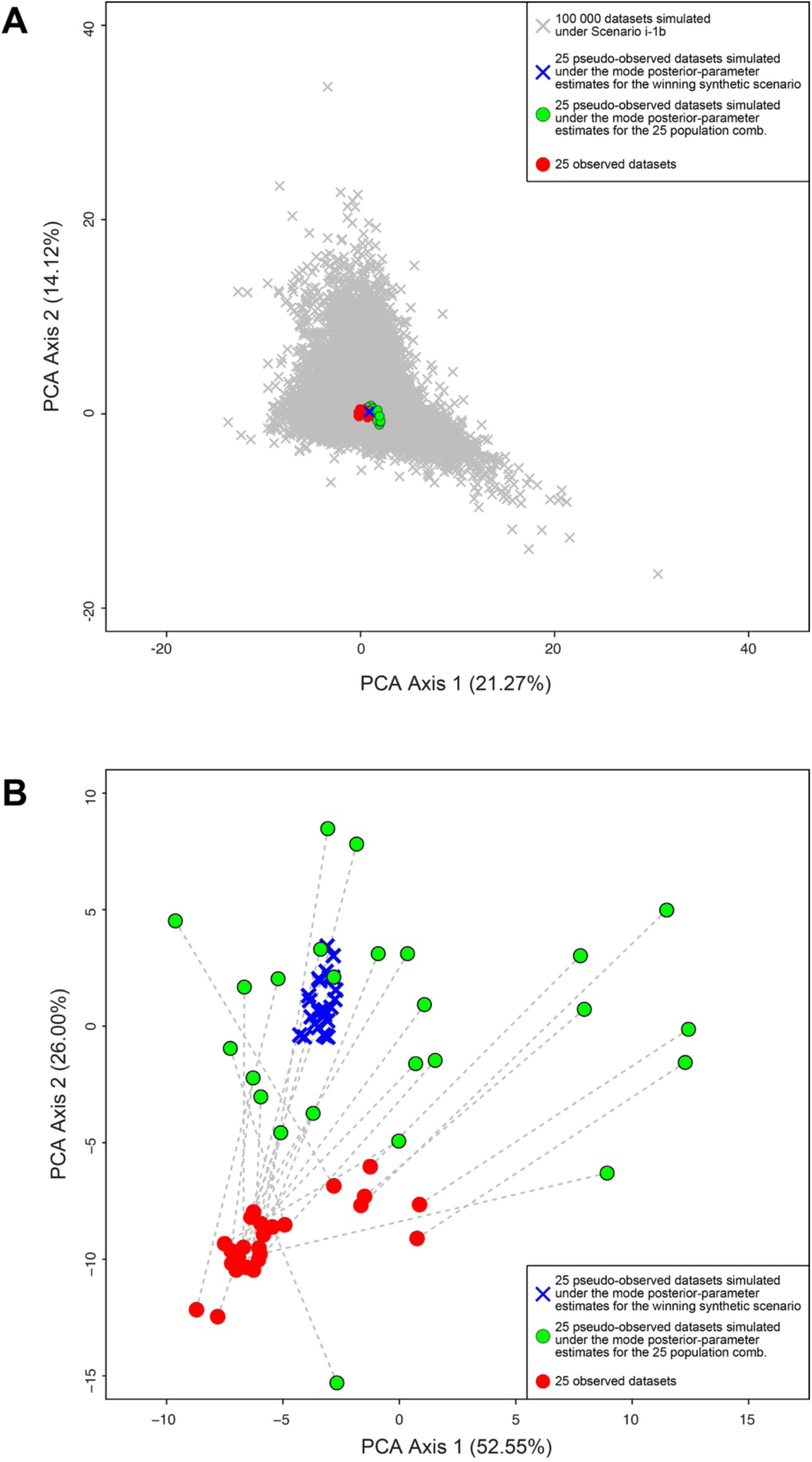
PCA posterior-check for the schematic NN-ABC inferred demographic and migration history of Central and Southern African populations proposed in Figure 8 and Table S14 and Table S17. First two axes of a PCA computed on 202 summary-statistics used to conduct Neural Network ABC posterior-parameter estimations under the best Scenario i1-1b. Observed summary-statistics for the 25 combinations of five Central and Southern African populations each (see **Material and Methods**), are indicated with red dots. Pseudo-observed summary-statistics for 25 separate simulated datasets of five populations each conducted using mode point-estimate values for each model-parameter provided in **Table S17** and schematized in **Figure 8** (see **Material and Methods**), are indicated with blue crosses. Pseudo-observed summary-statistics for 25 separate simulated datasets conducted using mode point-estimate values for each model-parameter provided in **Table S14** for each 25 combinations of five population, separately (see **Material and Methods**), are indicated with green dots. **Panel A** shows the PCA results computed also considering the sets of 202 summary-statistics computed for each 100,000 simulated datasets under Scenario i1-1b used as the reference for all NN-ABC posterior-parameter inferences, and indicated with grey crosses. **Panel B** shows the restricted PCA results where the grey dotted lines join each one of the sets of summary-statistics obtained for the 25 observed combinations of populations with their corresponding sets of summary-statistics from pseudo-observed simulations conducted using mode values of posterior parameter inferences provided in **Table S14.** Pseudo-observed simulations performed under the single synthetic model or under each best model for the 25 combinations of five populations provide summary-statistics reasonably close to that calculated on the 25 observed datasets. In **Panel A**, note that, as expected, most variation is explained among the 100,000 simulated datasets obtained by drawing random parameter-values in prior distributions under Scenario i1-1b and used as the reference table for NN-ABC posterior parameter inferences. In **Panel B**, note that, as expected, most variation in this PCA 2D plot is explained by differences among the 25 observed combinations of five populations each and their corresponding 25 pseudo-observed simulations. See **Figure S49** for visualizing the difference between observed statistics in the 25 observed combinations of sampled populations and the corresponding averaged summary-statistics values across the 25 pseudo-observed simulations conducted under our synthetic scenario in **Figure 8** and **Table S17**.

**Figure S49:**
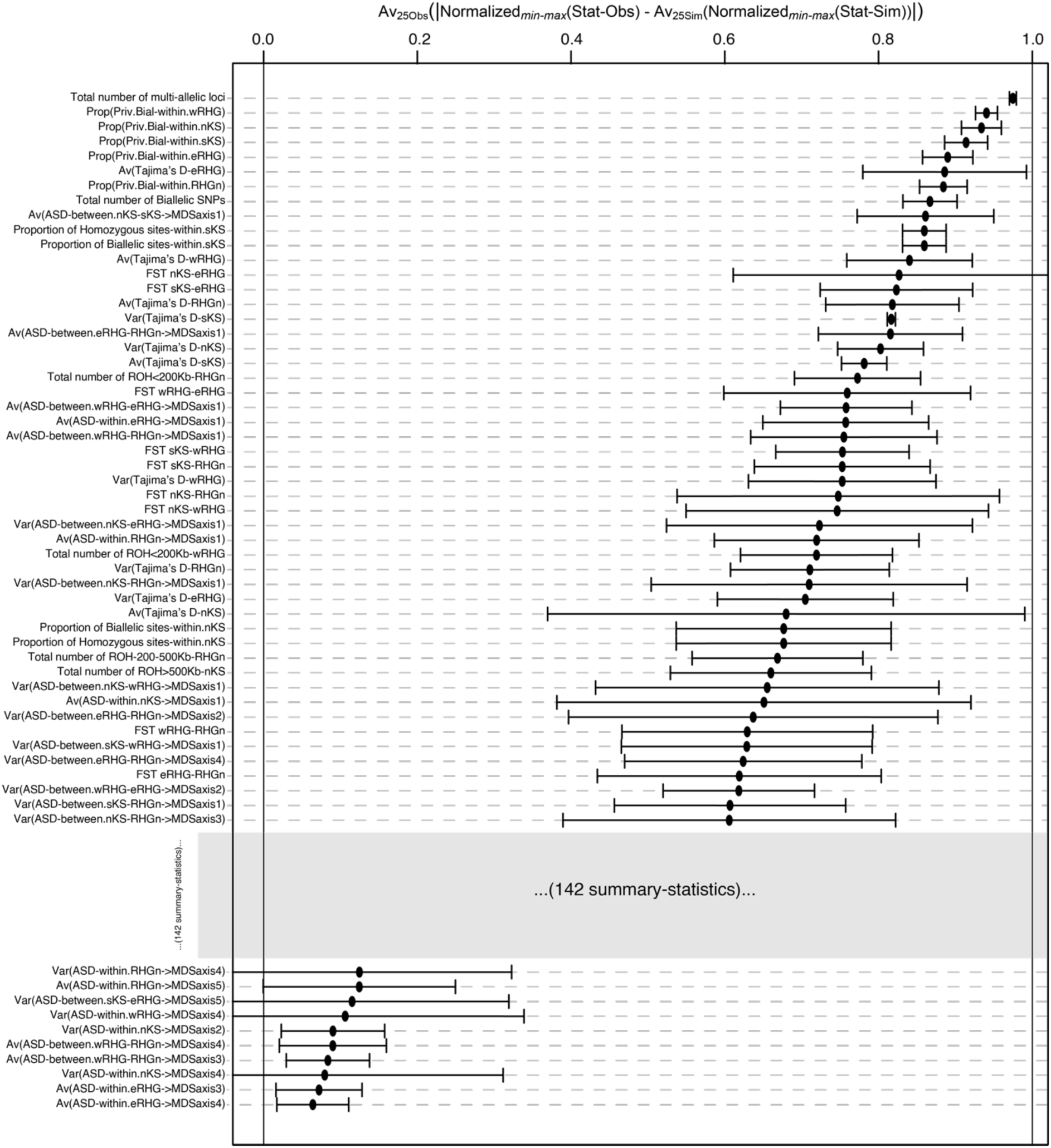
Differences between summary-statistics observed across the 25 combinations of five sampled populations and their average across the 25 pseudo-observed simulations under the synthetic demographic scenario in Figure 8. As mentioned in **Material and Methods**, we normalized (min-max) each summary-statistics among the 202 calculated separately for each one of the 25 combinations of five sampled population (the red dots in **Figure S48**), together with the corresponding statistics computed on the 25 pseudo-observed simulations performed under the synthetic scenario proposed in **Figure 8** and **Table S17** (the blue crosses in **Figure S48**). For each summary-statistics and each 25 observed combinations of population, we then calculated the difference between the normalized observed value and the average of the normalized value among the 25 pseudo-observed simulations. We finally averaged the absolute value of the obtained difference across the 25 observed combination of populations and provide the 50 most differentiated and 10 least differentiated summary-statistics in the figure indicated with black dots, +/- one standard deviation indicated with black tick-marks. The 10 summary-statistics that are most different between the 25 observed combinations of sampled populations and the average of the 25 pseudo-observed simulations under the synthetic demographic scenario in **Figure 8** and **Table S17** are: 1) the total number of multi-allelic loci, 2) the proportion of private biallelic loci within the wRHG, 3) within the nKS, 4) within the sKS, 5) within the eRHG, 6) the average Tajima’s D in the eRHG, 7) the proportion of private biallelic loci within the RHGn, 8) the total number of biallelic SNPs, 9) the average of the ASD-MDS projection on the first dimension between nKS and sKS individuals, and 10) the proportion of homozygous sites within the sKS.

**Figure S50:**
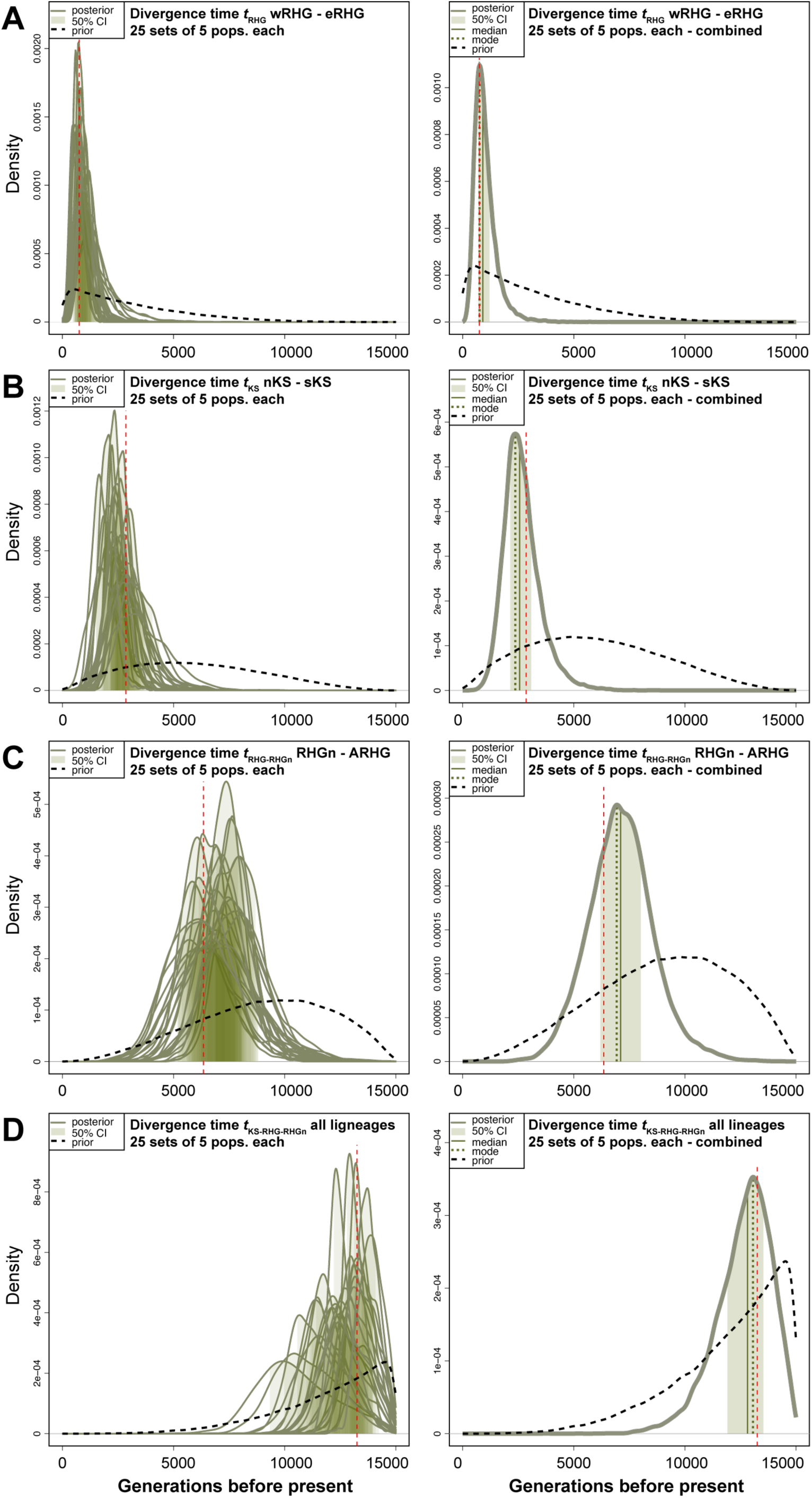
NN-ABC parameter inference posterior check: posterior distribution of divergence time parameters. **Figure** analog to **Figure 6** obtained using 25 pseudo-observed simulations under the synthetic scenario proposed in **Figure 8** and **Table S17** instead of the 25 observed datasets (see **Material and Methods**). Vertical red-doted lines indicate, for each scenario-parameter, the mode point-estimate values from **Table S17** used to perform the 25 simulations under Scenario i1-1b.

**Figure S51:**
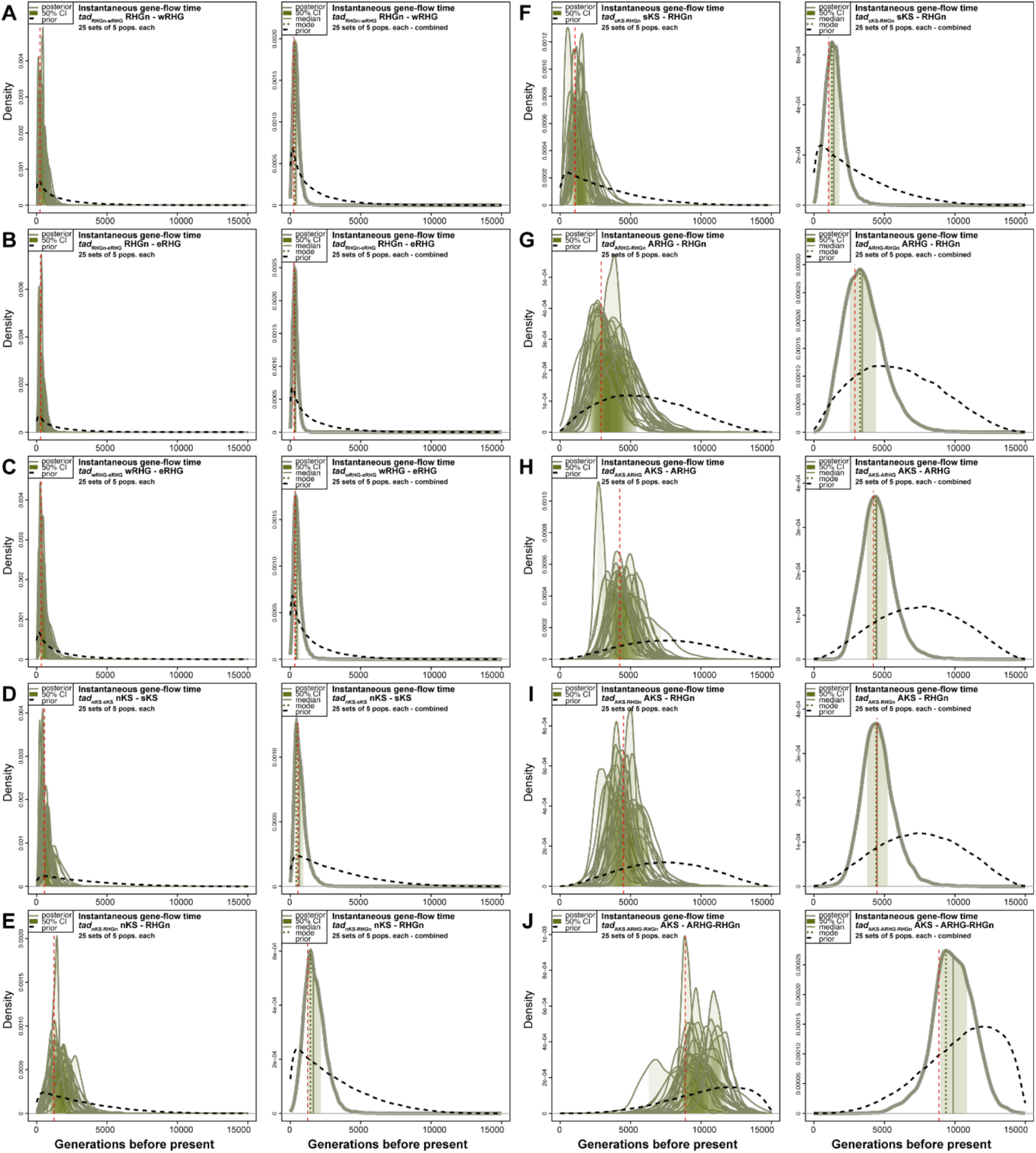
NN-ABC parameter inference posterior check: posterior distribution of potentially asymmetric instantaneous gene-flow time parameters. **Figure** analog to **Figure 7** obtained using 25 pseudo-observed simulations under the synthetic scenario proposed in **Figure 8** and **Table S17** instead of the 25 observed datasets (see **Material and Methods**). Vertical red-doted lines indicate, for each scenario-parameter, the mode point-estimate values from **Table S17** used to perform the 25 simulations under Scenario i1-1b.

**Figure S52:**
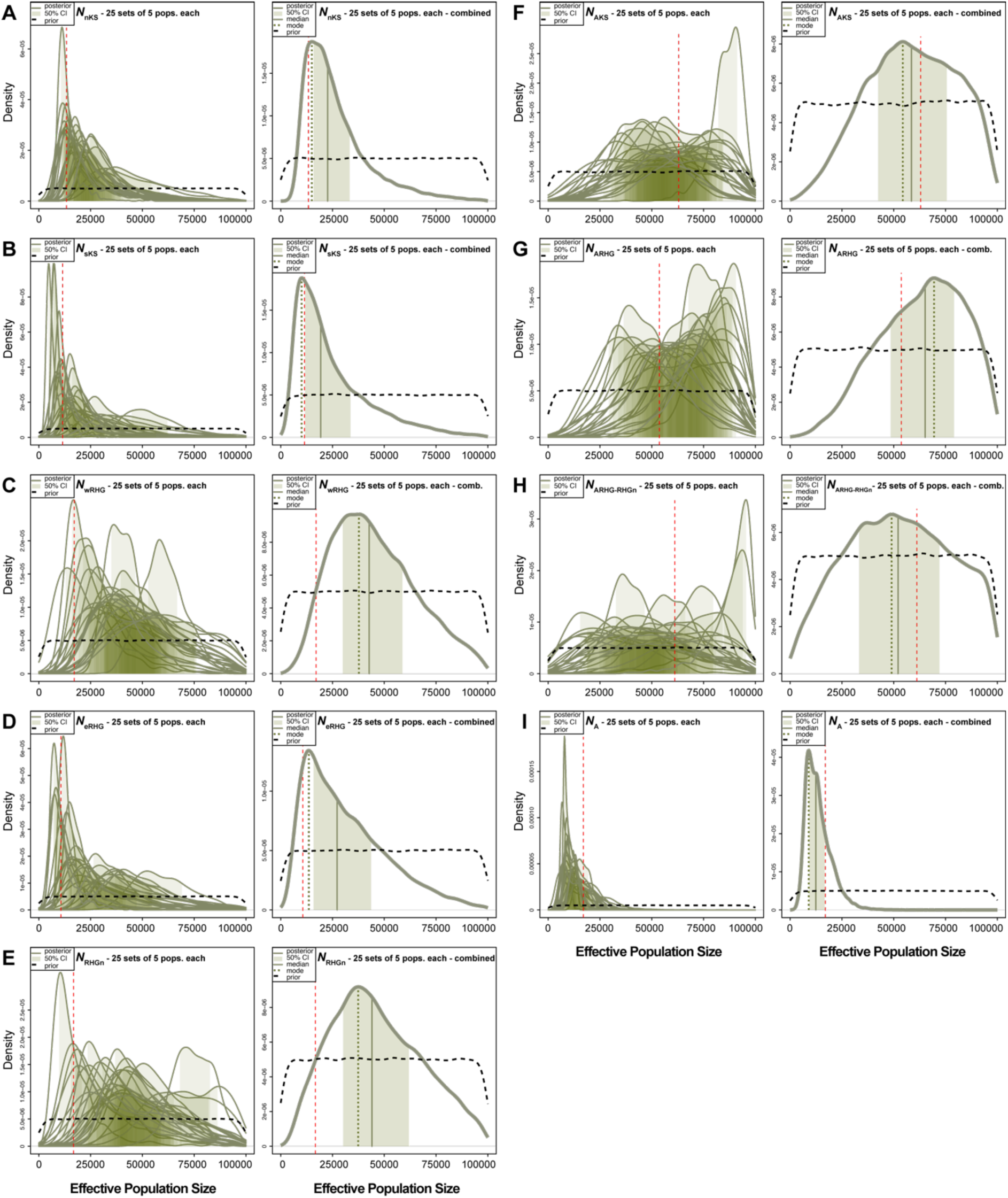
NN-ABC parameter inference posterior check: posterior distribution of effective population size parameters. **Figure** analog to **Figure S21** obtained using 25 pseudo-observed simulations under the synthetic scenario proposed in **Figure 8** and **Table S17** instead of the 25 observed datasets (see **Material and Methods**). Vertical red-doted lines indicate, for each scenario-parameter, the mode point-estimate values from TableS17 used to perform the 25 simulations under Scenario i1-1b.

**Figure S53:**
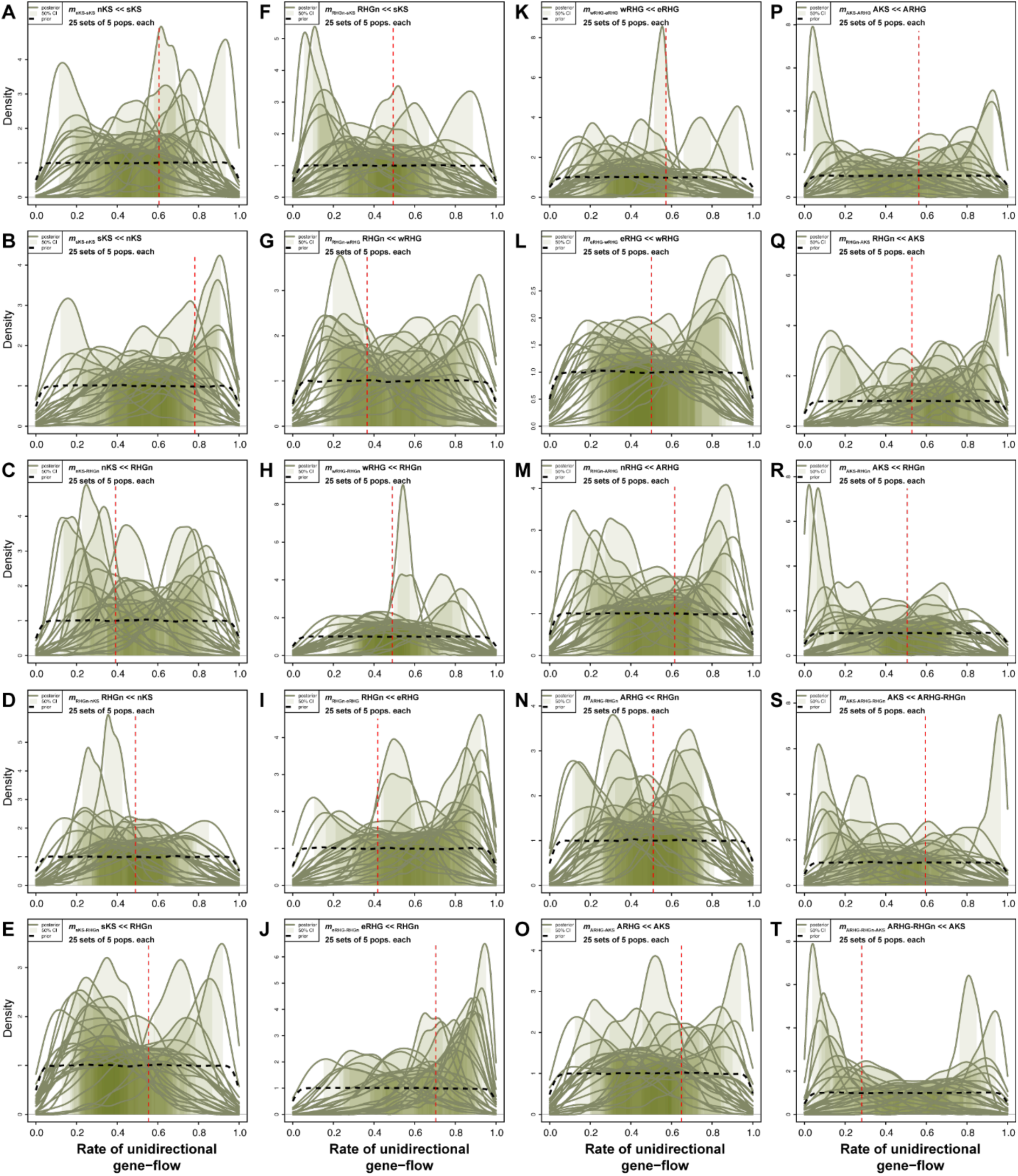
NN-ABC parameter inference posterior check: posterior distribution of potentially asymmetric instantaneous gene-flow intensity parameters. **Figure** analog to **Figure S22** obtained using 25 pseudo-observed simulations under the synthetic scenario proposed in **Figure 8** and **Table S17** instead of the 25 observed datasets (see **Material and Methods**). Vertical red-doted lines indicate, for each scenario-parameter, the mode point-estimate values from **Table S17** used to perform the 25 simulations under Scenario i1-1b.

**Figure S54:**
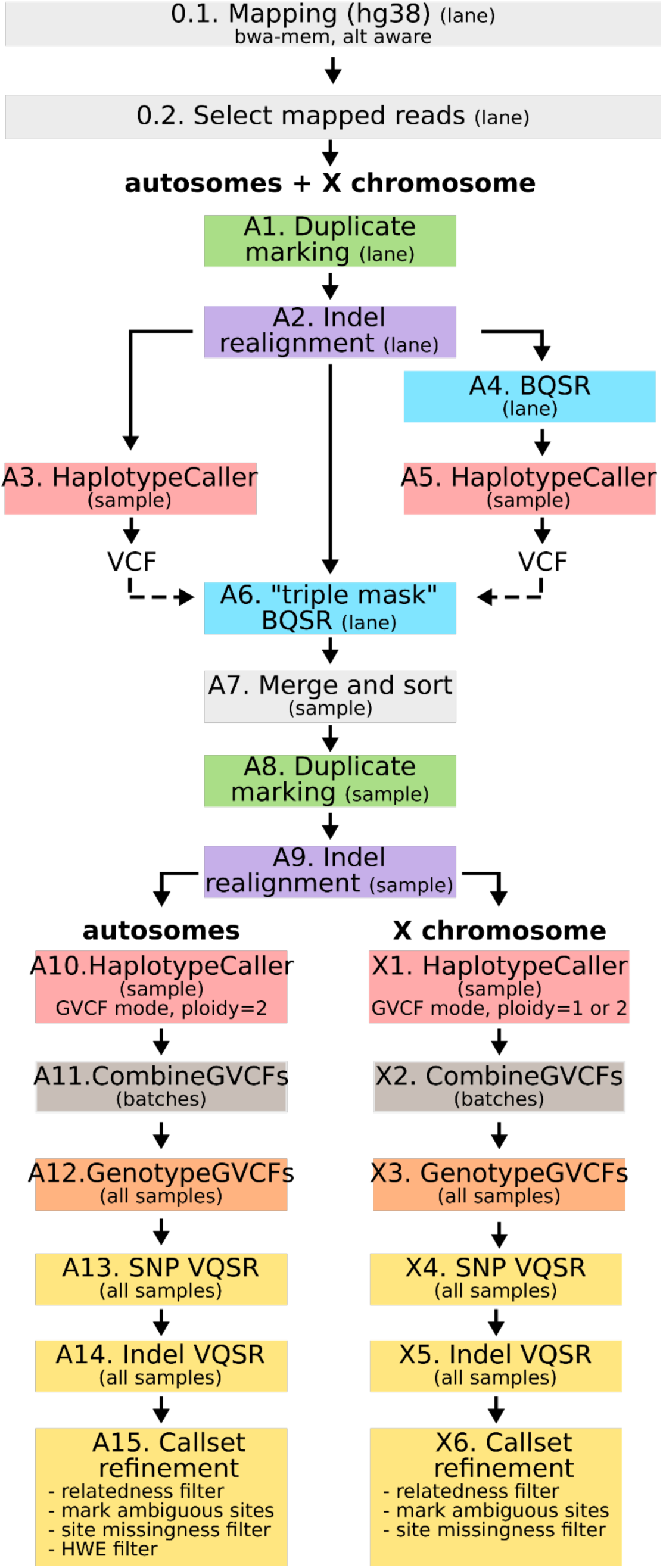
Schematic overview of the sequencing data processing pipeline. All template codes with accompanying detailed explanations for all steps are provided in **Material and Methods** and in the corresponding GitHub repository (https://github.com/Gwennid/africa-wgs-descriptive).

**Table S1:** Complementary population information for genetic-geographic comparisons, and for linguistic and life-style categories for ANOVA analyses. Geographical coordinates correspond to locations in Table 1 and **Figure 1**. “Traditional food-producing strategies” were obtained via ethno-anthropological investigations on the field for all original population-data, and from previously published material for comparison reference-datasets from the SGDP, HGDP, and KGP (see Table 1 and **Main Text** for references). “*nd*” stands for no explicit data concerning the genetic dataset specifically. “AGR” stands for agriculturalist and/or agropastoralist populations. “HG” stands for hunting, gathering, fishing, and horticulturalist populations. “Herder” stands for more strictly pastoral populations.

| Population Name | N | Dataset | GeoCoord:X | GeoCoord:Y | Linguistic Family | Sub-Linguistic Family | Traditional Food-Producing Strategy |
| --- | --- | --- | --- | --- | --- | --- | --- |
| Baka | 7 | this study | 13.86667 | 3.11667 | Niger-Congo | Niger-Congo, Adamawa-Ubangian | HG |
| Nzime | 5 | this study | 14.10000 | 2.98333 | Niger-Congo | Niger-Congo, Narrow-Bantu | AGR |
| Ba.Kola | 5 | this study | 10.48010 | 3.12090 | Niger-Congo | Niger-Congo, Narrow-Bantu | HG |
| Ngumba | 5 | this study | 10.72630 | 3.23490 | Niger-Congo | Niger-Congo, Narrow-Bantu | AGR |
| Bi.Aka Mbat | 5 | this study | 17.83462 | 3.76069 | Niger-Congo | Niger-Congo, Narrow-Bantu | HG |
| Ba.Kiga | 5 | this study | 29.61434 | -0.98659 | Niger-Congo | Niger-Congo, Narrow-Bantu | AGR |
| Ba.Twa | 6 | this study | 29.61434 | -0.98659 | Niger-Congo | Niger-Congo, Narrow-Bantu | HG |
| Nsua | 5 | this study | 30.25151 | 0.95051 | Nilo-Saharan | Nilo-Saharan, Eastern-Sudanic | HG |
| Ba.Konjo | 5 | this study | 30.25151 | 0.91938 | Niger-Congo | Niger-Congo, Narrow-Bantu | AGR |
| Karretjie People | 5 | this study | 25.10101 | -30.71264 | Khoisan | Khoisan | Herder |
| Khutse-San | 5 | this study | 24.66980 | -23.65090 | Khoisan | Khoisan | HG |
| Ju'hoansi | 5 | this study | 20.49500 | -19.59740 | Khoisan | Khoisan | HG |
| Nama | 5 | this study | 17.07275 | -22.55856 | Khoisan | Khoisan | Herder |
| 'Xun | 5 | this study | 17.66602 | -14.62894 | Khoisan | Khoisan | Herder |
| Bantu Herero | 2 | SGDP | 19.00000 | -22.00000 | Niger-Congo | Niger-Congo, Narrow-Bantu | Herder |
| Bantu Kenya | 2 | SGDP | 37.00000 | -3.00000 | Niger-Congo | Niger-Congo, Narrow-Bantu | AGR |
| Bantu Tswana | 2 | SGDP | 24.00000 | -28.00000 | Niger-Congo | Niger-Congo, Narrow-Bantu | AGR |
| Biaka | 2 | SGDP | 17.00000 | 4.00000 | Niger-Congo | Niger-Congo, Narrow-Bantu | HG |
| Coloured | 8 | SAHGP | 18.41740 | -33.92899 | Indo-European | Indo-European, Afrikaans | *nd* |
| Dinka | 4 | SGDP, HGDP | 27.40000 | 8.78333 | Nilo-Saharan | Nilo-Saharan, Eastern-Sudanic | AGR |
| Esan | 3 | SGDP, KGP | 6.00000 | 6.50000 | Niger-Congo | Niger-Congo, Volta-Niger | AGR |
| Gambian | 2 | SGDP | -16.70000 | 13.40000 | Niger-Congo | Niger-Congo, Mande | AGR |
| Igbo | 2 | SGDP | 7.00000 | 6.00000 | Niger-Congo | Niger-Congo, Volta-Niger | AGR |
| Ju'hoansi comp | 4 | SGDP, HGDP | 21.50000 | -18.90000 | Khoisan | Khoisan | HG |
| #Khomani | 2 | SGDP | 20.80000 | -27.00000 | Khoisan | Khoisan | Herder |
| Kongo | 1 | SGDP | 11.80000 | 2.50000 | Niger-Congo | Niger-Congo, Narrow-Bantu | AGR |
| Lemande | 2 | SGDP | 11.10000 | 4.50000 | Niger-Congo | Niger-Congo, Narrow-Bantu | AGR |
| Luhya | 3 | SGDP, KGP | 36.80000 | 1.30000 | Niger-Congo | Niger-Congo, Narrow-Bantu | AGR |
| Luo | 2 | SGDP | 34.30000 | -0.10000 | Nilo-Saharan | Nilo-Saharan, Eastern-Sudanic | AGR |
| Maasai | 2 | SGDP | 35.20000 | -1.50000 | Nilo-Saharan | Nilo-Saharan, Eastern-Sudanic | Herder |
| Mandenka | 4 | SGDP, HGDP | -12.00000 | 12.00000 | Niger-Congo | Niger-Congo, Mande | AGR |
| Mandinka | 1 | KGP | -16.70000 | 13.40000 | Niger-Congo | Niger-Congo, Mande | AGR |
| Mbuti | 5 | SGDP, HGDP | 29.00000 | 1.00000 | Niger-Congo | Niger-Congo, Narrow-Bantu | HG |
| Mende | 3 | SGDP | -13.20000 | 8.20000 | Niger-Congo | Niger-Congo, Mande | AGR |
| Mozabite | 2 | SGDP | 3.00000 | 32.00000 | Afro-Asiatic | Afro-Asiatic, Berber | AGR |
| Saharawi | 2 | SGDP | -8.90000 | 27.30000 | Afro-Asiatic | Afro-Asiatic, Hassaniya | *nd* |
| Somali | 1 | SGDP | 48.30000 | 5.60000 | Afro-Asiatic | Afro-Asiatic, Cushitic | *nd* |
| Sotho | 7 | SAHGP | 27.13588 | -28.08954 | Niger-Congo | Niger-Congo, Narrow-Bantu | *nd* |
| Xhosa | 8 | SAHGP | 25.62075 | -33.96171 | Niger-Congo | Niger-Congo, Narrow-Bantu | *nd* |
| Yoruba | 4 | SGDP, HGDP | 3.90000 | 7.40000 | Niger-Congo | Niger-Congo, Volta-Niger | AGR |
| Zulu | 1 | SAHGP | 28.04972 | -26.20500 | Niger-Congo | Niger-Congo, Narrow-Bantu | *nd* |

**Table S2:** Whole genome total variant counts among 177 unrelated worldwide individuals. Variant counts compared to the reference sequence for the human genome GRCh38 and previously reported variants in dbSNP 156. “SD” stands for “standard deviation”. See **Material and Methods** for details about the quality control, family relatedness filtering, and variant count procedures.

|  | <u>Total variant count</u> |  | <u>Per individual variant count among 177 worldwide unrelated individuals</u> |  |  |  |
| --- | --- | --- | --- | --- | --- | --- |
|  | Original dataset:<br>73 Central and<br>Southern African<br>unrelated individuals | Entire dataset:<br>177 worldwide<br>unrelated<br>individuals | Total | SD | Min | Max |
| Number of biallelic SNPs | 26,780,319 | 36,272,545 | 4,245,000 | 375,379 | 3,093,785 | 4,642,544 |
| Number of biallelic SNPs not<br>in dbSNP 156 | 854,114 | 1,055,245 | 8,052 | 4,563 | 1,152 | 27,406 |
| Number of multiallelic SNPs | 241,428 | 257,906 | 56,330 | 5,162 | 38,057 | 61,716 |
| Number of simple indels | 2,454,965 | 3,159,306 | 385,391 | 35,522 | 278,237 | 421,976 |
| Number of novel simple<br>indels not in dbSNP 156 | 114,362 | 189,124 | 1,651 | 514 | 771 | 627 |
| Number of complex indels | 969,025 | 977,542 | 414,037 | 71,029 | 289,993 | 566,951 |
| Number of variants appearing<br>in a single sample | 8,404,499 | 12,638,316 | 71,403 | 15,094 | 22,444 | 104,211 |
| Number of filtered SNPs | 2,940,629 | 3,296,775 | 247,172 | 48,579 | 66,597 | 350,377 |

**Table S3:** Whole genome biallelic SNPs counts in 46 worldwide populations. Mean number of bi-allelic SNPs across unrelated individuals from 46 worldwide populations, and associated standard deviations (when applicable), compared to the reference sequence for the human genome GRCh38 and compared to previously reported variants in dbSNP 156. See **Material and Methods** for details about the quality control, family relatedness filtering, and variant count procedures. Note that, for the samples original to this study, to reach the mean human coverage reported here required greater or lesser sequencing efforts depending on the biological origin of each sample (saliva or blood), and its quality. Populations are ordered in increasing mean number of SNPs per pop. The information original to this study is indicated in bold. “SD” stands for standard deviation. “*na*” stands for not applicable. Additional population information and geographical location of samples can be found in **Figure 1, Table 1**, and **Table S1**.

| Population Name<sup>1</sup> | Sampling location | Dataset<sup>2</sup> | N<sup>3</sup> | Mean coverage X (SD) | Mean number of biallelic SNPs (SD) | Mean number of biallelic SNPs not in dbSNP 156 (SD) |
| --- | --- | --- | --- | --- | --- | --- |
| Karitiana | Brazil | SGDP, HGDP | 5 | 31.1 (7.8) | 3,214,086 (69,241) | 1,030 (547) |
| Papuan | Papua New Guinea | SGDP, HGDP | 6 | 35.8 (7.8) | 3,414,518 (74,893) | 841 (290) |
| French | France | SGDP, HGDP | 4 | 33.8 (8.7) | 3,455,358 (77,954) | 652 (225) |
| Dai | China | SGDP, KGP | 6 | 37.7 (10.5) | 3,480,744 (67,644) | 695 (407) |
| CEU | United States of America | KGP | 2 | 67.3 (0.2) | 3,551,792 (7,819) | 1,539 (57) |
| Saharawi | Western Sahara | SGDP | 2 | 43.7 (2.2) | 3,726,018 (27,793) | 590 (21) |
| Mozabite | Algeria | SGDP | 2 | 34.9 (1.6) | 3,736,308 (69,393) | 563 (11) |
| Somali | Kenya | SGDP | 1 | 37.7 (na) | 3,940,793 (na) | 540 (na) |
| Coloured | South Africa | SAHGP | 8 | 44.9 (2.2) | 4,000,396 (97,447) | 10,169 (2,319) |
| Dinka | Sudan | SGDP, HGDP | 4 | 31.3 (5.8) | 4,142,138 (97,001) | 3,146 (4,811) |
| Maasai | Kenya | SGDP | 2 | 38.6 (5.5) | 4,169,732 (5,281) | 654 (27) |
| Mandinka | Gambia | KGP | 1 | 27.2 (na) | 4,193,651 (na) | 1,602 (na) |
| Mandenka | Senegal | SGDP, HGDP | 4 | 30 (7.6) | 4,217,398 (107,847) | 914 (464) |
| Yoruba | Nigeria | SGDP, HGDP | 4 | 31.1 (5) | 4,233,500 (11,5371) | 755 (195) |
| **Ba.Kiga** | **Mukono (Uganda)** | **this study** | **5** | **39.4 (4.6)** | **4,274,466 (25,201)** | **5,391 (191)** |
| Gambian | Gambia | SGDP | 2 | 35.5 (0.8) | 4,277,874 (12,976) | 816 (107) |
| Esan | Nigeria | SGDP, KGP | 3 | 39.7 (9.5) | 4,284,041 (18,359) | 704 (67) |
| Bantu Kenya | Kenya | SGDP | 2 | 38.7 (6.8) | 4,288,533 (10,954) | 611 (88) |
| Luo | Kenya | SGDP | 2 | 34 (0.2) | 4,293,632 (1,003) | 722 (4) |
| Luhya | Kenya | SGDP, KGP | 3 | 40.6 (12.1) | 4,294,198 (28,096) | 704 (37) |
| Igbo | Nigeria | SGDP | 2 | 36.3 (8) | 4,294,870 (1,867) | 1,166 (9) |
| Lemende | Cameroon | SGDP | 2 | 36.3 (0.5) | 4,306,880 (5,284) | 1,530 (37) |
| Bantu Herero | Namibia | SGDP | 2 | 38.4 (2.2) | 4,318,604 (33,525) | 690 (161) |
| **Ba.Konjo** | **Mulimassenge (Uganda)** | **this study** | **5** | **43 (6.4)** | **4,322,177 (6,336)** | **5,349 (773)** |
| **Nzime** | **Messea (Cameroon)** | **this study** | **5** | **35.5 (2.6)** | **4,322,734 (18,671)** | **5,305 (208)** |
| **Ngumba** | **Dispersed between Lolodorf and Kribi (Cameroon)** | **this study** | **5** | **40.2 (2.2)** | **4,332,324 (9,273)** | **4,810 (552)** |
| Kongo | Cameroon | SGDP | 1 | 42.8 (na) | 4,336,731 (na) | 1233 (na) |
| Mende | Sierra Leone | SGDP, KGP | 3 | 39.6 (8.3) | 4,341,512 (13,486) | 866 (180) |
| Zulu | South Africa | SAHGP | 1 | 42.8 (na) | 4,376,691 (na) | 8,146 (na) |
| Xhosa | South Africa | SAHGP | 8 | 42.4 (1.1) | 4,385,032 (56,902) | 10,085 (1,300) |
| **Ba.Twa** | **Kebiremu, Byumba, Kitariro, Mgunu, Nteko (Uganda)** | **this study** | **6** | **53.8 (17.7)** | **4,426,223 (32,552)** | **20,251 (1,022)** |
| Bantu Tswana | South Africa | SGDP | 2 | 36.7 (3.1) | 4,430,870 (55,537) | 720 (37) |
| Mbuti | Democratic Republic of Congo | SGDP, HGDP | 5 | 32.1 (7.4) | 4,434,425 (110,658) | 1,227 (488) |
| Sotho | South Africa | SAHGP | 7 | 42.9 (6.4) | 4,446,796 (23,172) | 12,040 (1,150) |
| **Ba.Kola** | **Dispersed between Lolodorf and Kribi (Cameroon)** | **this study** | **5** | **39.2 (1.9)** | **4,475,922 (50,261)** | **6,995 (1,948)** |
| **Nsua** | **Bundimassoli (Uganda)** | **this study** | **5** | **37.6 (4.9)** | **4,486,217 (22,774)** | **16,443 (984)** |
| **Baka** | **Bosquet (Cameroon)** | **this study** | **7** | **40.7 (3.5)** | **4,503,977 (23,946)** | **3,927 (144)** |
| **Bi.Aka_Mbati** | **Bombeketi section of Bagandou (Central African Republic)** | **this study** | **5** | **43.4 (13.6)** | **4,514,318 (17,376)** | **4,587 (198)** |
| Biaka | Central African Republic | SGDP | 2 | 39.1 (0.3) | 4,521,140 (8,155) | 726 (52) |
| **Nama** | **Windhoek (Namibia)** | **this study** | **5** | **48.9 (4.3)** | **4,530,168 (18,807)** | **27,902 (462)** |
| Ju\|'hoansi comp | Namibia | SGDP, HGDP | 4 | 33.9 (7) | 4,554,593 (103,099) | 1,382 (273) |
| #Khomani | South Africa | SGDP | 2 | 44.7 (2.8) | 4,559,846 (79,851) | 1,364 (48) |
| **Khutse_San** | **Kutse Game reserve (Botswana)** | **this study** | **5** | **46.3 (6.7)** | **4,592,852 (27,794)** | **27,989 (1,353)** |
| **Karretjie** | **Colesberg (South Africa)** | **this study** | **5** | **46.6 (5.4)** | **4,601,724 (26,951)** | **27,720 (1,968)** |
| **!Xun** | **Omega camp (Namibia) and Schmidtsdrift (South Africa)<sup>4</sup>** | **this study** | **5** | **45.6 (1.9)** | **4,614,835 (10,047)** | **28,923 (1,371)** |
| Ju\|'hoansi | Tsumkwe (Namibia) | this study | 5 | 49 (4.6) | 4,628,957 (8,139) | 26,665 (6,491) |

**Table S4:** Correlations between genetic pairwise differentiation and geography. Standardized average Allele Sharing Dissimilarities across pairs of individuals from different populations are compared, for all population pairs, with the logarithm of geographic distances using Spearman and Pearson correlations, and tested using 1000 Mantel permutations. See **Figure 1, Table 1**, and **Table S1** for population geographic location and information.

| Geographic Region | Population Samples | Nb pops. | Nb indivs. | Spearman $\rho$ | Mantel Permutation p | Pearson $r$ | Mantel Permutation p |
| --- | --- | --- | --- | --- | --- | --- | --- |
| Africa | All African population samples in **Figure1, Table1, and TableS1** | 41 | 154 | 0.03573 | 0.32268 | 0.05562 | 0.16384 |
| Sub-Saharan Africa | All African population samples in **Figure1, Table1, and TableS1**, except Northern African Sarahwi and Mozabites | 38 | 150 | 0.07321 | 0.16883 | 0.09707 | 0.05294 |
| West Western Africa | Gambian, Mandenka, Mandinka, Mende | 4 | 10 | 0.50022 | 0.09291 | 0.38444 | 0.33467 |
| Wider Central Africa | Igbo, Esan, Yoruba, Ba.Kola (wRHG), Ngumba (wRHGn), Baka (wRHG), Nzime(wRHGn), Kongo, Lemande, Bi.Aka_Mbati (wRHG), Bi.Aka (wRHG), Ba.Twa (eRHG), Ba.Kiga (eRHGn), Nsua (eRHG), Ba.Konjo (eRHGn), Mbuti (eRHG), Dinka | 17 | 71 | 0.15587 | 0.07393 | 0.06298 | 0.21378 |
| Central Africa - Congo Basin | Ba.Kola (wRHG), Ngumba (wRHGn), Baka (wRHG), Nzime(wRHGn), Kongo, Lemande, Bi.Aka_Mbati (wRHG), Bi.Aka (wRHG), Ba.Twa (eRHG), Ba.Kiga (eRHGn), Nsua (eRHG), Ba.Konjo (eRHGn), Mbuti (eRHG) | 13 | 58 | 0.19952 | 0.03996 | 0.13451 | 0.05594 |
| Eastern Africa | Bantu_Kenya, Luhya, Luo, Maasai, Somali | 5 | 10 | 0.13939 | 0.25574 | -0.04705 | 0.36863 |
| Southern Africa | !Xun (nKS), Ju\|'hoansi (nKS), Ju\|'hoansi_comp (nKS), Bantu_Herero, Nama (sKS), Khutse San (Central KS), #Khomani, Zulu, Sotho, Bantu_Tswana, Karretjie_People (sKS), Coloured, Xhosa | 13 | 59 | 0.07123 | 0.20979 | 0.10440 | 0.15485 |

**Table S5:** Permutation ANOVA of average population pairwise ASD explained by population linguistic families at different geographical scale in Africa. See **Figure 1, Table 1** and **Table S1** for population geographical location and linguistic family categorization criteria. Significant ANOVA results are indicated in bold and non-significant results in italics. “*na*” indicates that the ANOVA test is not applicable due to insufficient numbers of levels in the linguistic family category.

| Geographic Region | Population Samples | Nb pops | Nb Linguistic families | ANOVA partial $r^2$: % Var(Av. ASD among pairs of populations) explained | | |
| --- | --- | --- | --- | --- | --- | --- |
|  |  |  |  | among linguistic families | within linguistic families | 1000 ANOVA permutation p |
| Africa | All African population samples in **Figure1**, **Table1**, and **TableS1** | 41 | 5 | **15.064%** | **84.936%** | **<0.001** |
| Sub-Sahara Africa | All African population samples in **Figure1**, **Table1**, and **TableS1**, except Northern African Sarahwi and Mozabites | 39 | 5 | **14.811%** | **85.189%** | **<0.001** |
| West Western Africa | Gambian, Mandenka, Mandinka, Mende | 4 | 1 | *Na* | *Na* | *na* |
| Wider Central Africa | Igbo, Esan, Yoruba, Ba.Kola (wRHG), Ngumba (wRHGn), Baka (wRHG), Nzime(wRHGn), Kongo, Lemande, Bi.Aka_Mbati (wRHG), Bi.Aka (wRHG), Ba.Twa (eRHG), Ba.Kiga (eRHGn), Nsua (eRHG), Ba.Konjo (eRHGn), Mbuti (eRHG), Dinka | 17 | 2 | *6.389%* | *93.611%* | *0.1698* |
| Central Africa - Congo Basin | Ba.Kola (wRHG), Ngumba (wRHGn), Baka (wRHG), Nzime(wRHGn), Kongo, Lemande, Bi.Aka_Mbati (wRHG), Bi.Aka (wRHG), Ba.Twa (eRHG), Ba.Kiga (eRHGn), Nsua (eRHG), Ba.Konjo (eRHGn), Mbuti (eRHG) | 13 | 2 | *9.172%* | *90.828%* | *0.1489* |
| Eastern Africa | Bantu_Kenya, Luhya, Luo, Maasai, Somali | 5 | 3 | *50.311%* | *49.689%* | *0.297* |
| Southern Africa | !Xun (nKS), Ju\|'hoansi (nKS), Ju\|'hoansi_comp (nKS), Bantu_Herero, Nama (sKS), Khutse San (Central KS), #Khomani, Zulu, Sotho, Bantu_Tswana, Karretjie_People (sKS), Coloured, Xhosa | 13 | 3 | **21.544%** | **78.456%** | **<0.001** |

**Table S6:** Permutation ANOVA of average population pairwise ASD explained by population sub-linguistic families at different geographical scale in Africa. See **Figure 1, Table 1** and **Table S1** for population geographical location and sub-linguistic family categorization criteria. Significant ANOVA results are indicated in bold and non-significant results in italics. “*na*” indicates that the ANOVA test is not applicable due to insufficient numbers of levels in the linguistic family category.

| Geographic Region | Population Samples | Nb pops | Nb Sub-linguistic families | ANOVA partial $r^2$: % Var(Av. ASD among pairs of populations) explained | | |
| --- | --- | --- | --- | --- | --- | --- |
|  |  |  |  | among Sub-linguistic families | within Sub-linguistic families | 1000 ANOVA permutation p |
| Africa | All African population samples in **Figure1**, **Table1**, and **TableS1** | 41 | 10 | **26.109%** | **73.891%** | **<0.001** |
| Sub-Sahara Africa | All African population samples in **Figure1**, **Table1**, and **TableS1**, except Northern African Sarahwi and Mozabites | 39 | 8 | **22.759%** | **77.241%** | **<0.001** |
| West Western Africa | Gambian, Mandenka, Mandinka, Mende | 4 | 1 | *na* | *na* | *na* |
| Wider Central Africa | Igbo, Esan, Yoruba, Ba.Kola (wRHG), Ngumba (wRHGn), Baka (wRHG), Nzime(wRHGn), Kongo, Lemande, Bi.Aka_Mbati (wRHG), Bi.Aka (wRHG), Ba.Twa (eRHG), Ba.Kiga (eRHGn), Nsua (eRHG), Ba.Konjo (eRHGn), Mbuti (eRHG), Dinka | 17 | 4 | *19.441%* | *80.559%* | *0.1728* |
| Central Africa - Congo Basin | Ba.Kola (wRHG), Ngumba (wRHGn), Baka (wRHG), Nzime(wRHGn), Kongo, Lemande, Bi.Aka_Mbati (wRHG), Bi.Aka (wRHG), Ba.Twa (eRHG), Ba.Kiga (eRHGn), Nsua (eRHG), Ba.Konjo (eRHGn), Mbuti (eRHG) | 13 | 3 | *17.832%* | *82.168%* | *0.1538* |
| Eastern Africa | Bantu_Kenya, Luhya, Luo, Maasai, Somali | 5 | 3 | *50.311%* | *49.689%* | *0.297* |
| Southern Africa | !Xun (nKS), Ju\|'hoansi (nKS), Ju\|'hoansi_comp (nKS), Bantu_Herero, Nama (sKS), Khutse San (Central KS), #Khomani, Zulu, Sotho, Bantu_Tswana, Karretjie_People (sKS), Coloured, Xhosa | 13 | 3 | **21.544%** | **78.456%** | **<0.001** |

**Table S7:** Permutation ANOVA of average population pairwise ASD explained by population traditional food-production strategies at different geographical scale in Africa. See **Figure 1, Table 1** and **Table S1** for population geographical location and traditional food-production strategy categorization criteria. Significant ANOVA results are indicated in bold and non-significant results in italics. “*na*” indicates that the ANOVA test is not applicable due to insufficient numbers of levels in the linguistic family category.

| Geographic Region | Population Samples | Nb pops | Nb Traditional food-production strategies | ANOVA partial $r^2$: % Var(Av. ASD among pairs of populations) explained | | |
| --- | --- | --- | --- | --- | --- | --- |
|  |  |  |  | among strategies | within strategies | 1000 ANOVA permutation p |
| Africa | All African population samples in Figure1 and Table1, except Saharawi, Coloured, Somali, Sotho, Xhosa, and Zulu populations (TableS1) | 35 | 3 | **9.79%** | **90.21%** | **<0.001** |
| Sub-Sahara Africa | All African population samples in Figure1 and Table1, except Northern African Saharawi and Mozabite, and Southern African Coloured, Somali, Sotho, Xhosa, and Zulu populations (TableS1) | 34 | 3 | **10.01%** | **89.99%** | **<0.001** |
| West Western Africa | Gambian, Mandenka, Mandinka, Mende | 4 | 1 | *na* | *Na* | *na* |
| Wider Central Africa | Igbo, Esan, Yoruba, Ba.Kola (wRHG), Ngumba (wRHGn), Baka (wRHG), Nzime(wRHGn), Kongo, Lemande, Bi.Aka_Mbati (wRHG), Bi.Aka (wRHG), Ba.Twa (eRHG), Ba.Kiga (eRHGn), Nsua (eRHG), Ba.Konjo (eRHGn), Mbuti (eRHG), Dinka | 17 | 2 | **9.538%** | **90.462%** | **<0.001** |
| Central Africa - Congo Basin | Ba.Kola (wRHG), Ngumba (wRHGn), Baka (wRHG), Nzime(wRHGn), Kongo, Lemande, Bi.Aka_Mbati (wRHG), Bi.Aka (wRHG), Ba.Twa (eRHG), Ba.Kiga (eRHGn), Nsua (eRHG), Ba.Konjo (eRHGn), Mbuti (eRHG) | 13 | 2 | **11.153%** | **88.847%** | **0.003** |
| Eastern Africa | Bantu_Kenya, Luhya, Luo, Maasai | 4 | 2 | *34.198%* | *65.802%* | *0.25* |
| Southern Africa | !Xun (nKS), Ju\|'hoansi (nKS), Ju\|'hoansi_comp (nKS), Bantu_Herero, Nama (sKS), Khutse San (Central KS), #Khomani, Bantu_Tswana, Karretjie_People (sKS) | 9 | 3 | *25.979%* | *74.201%* | *0.1728* |

**Table S8:**
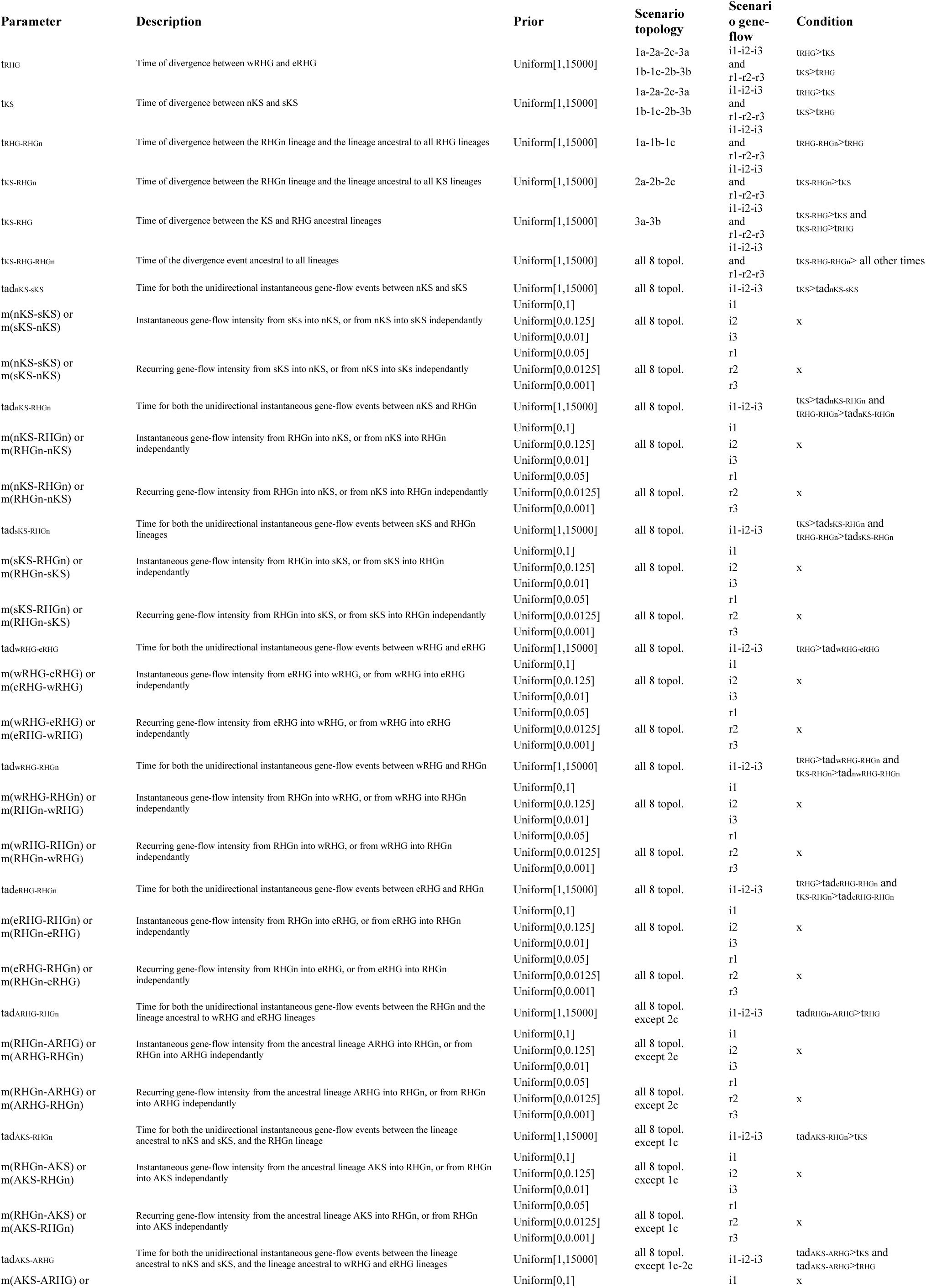
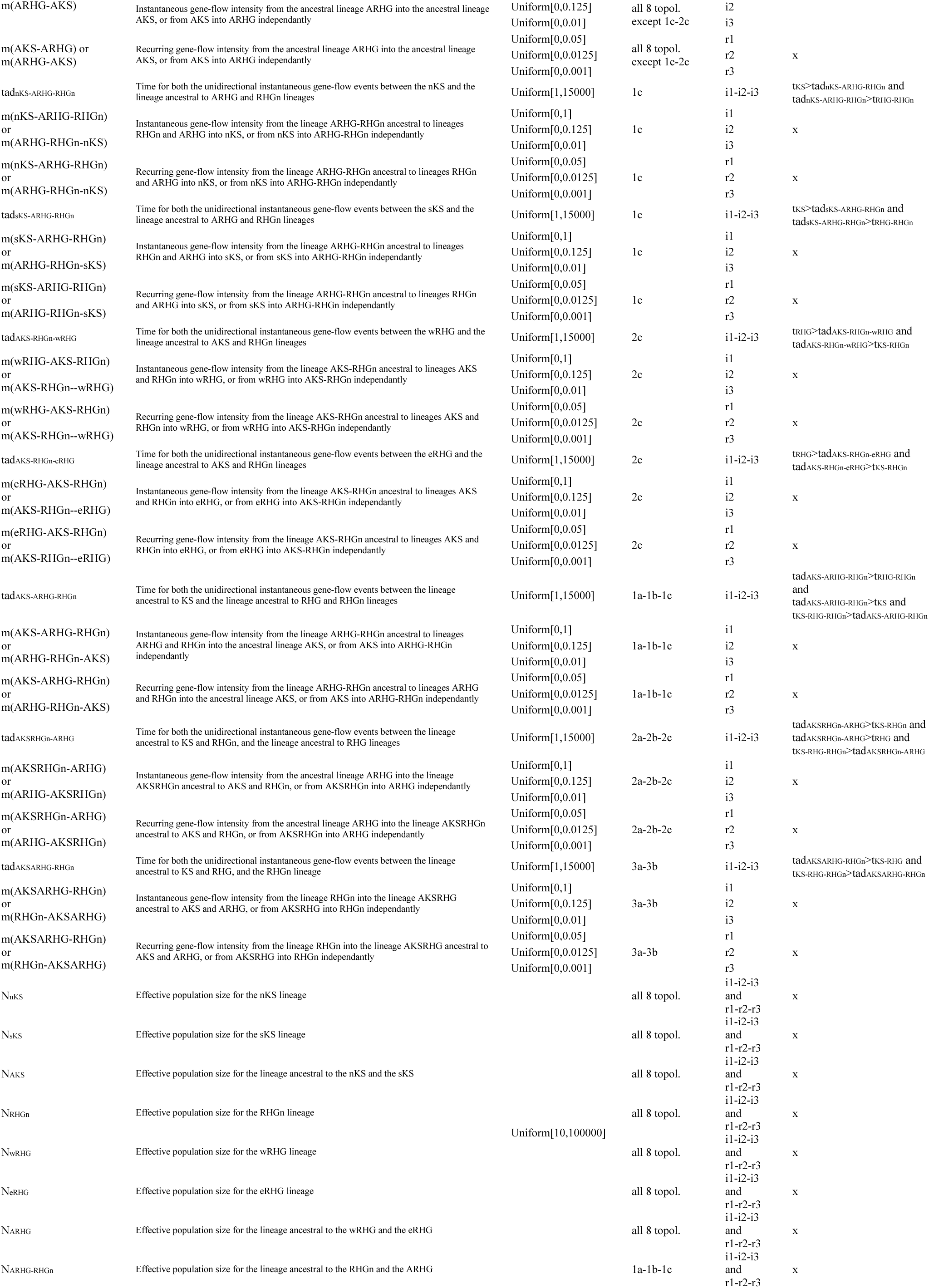
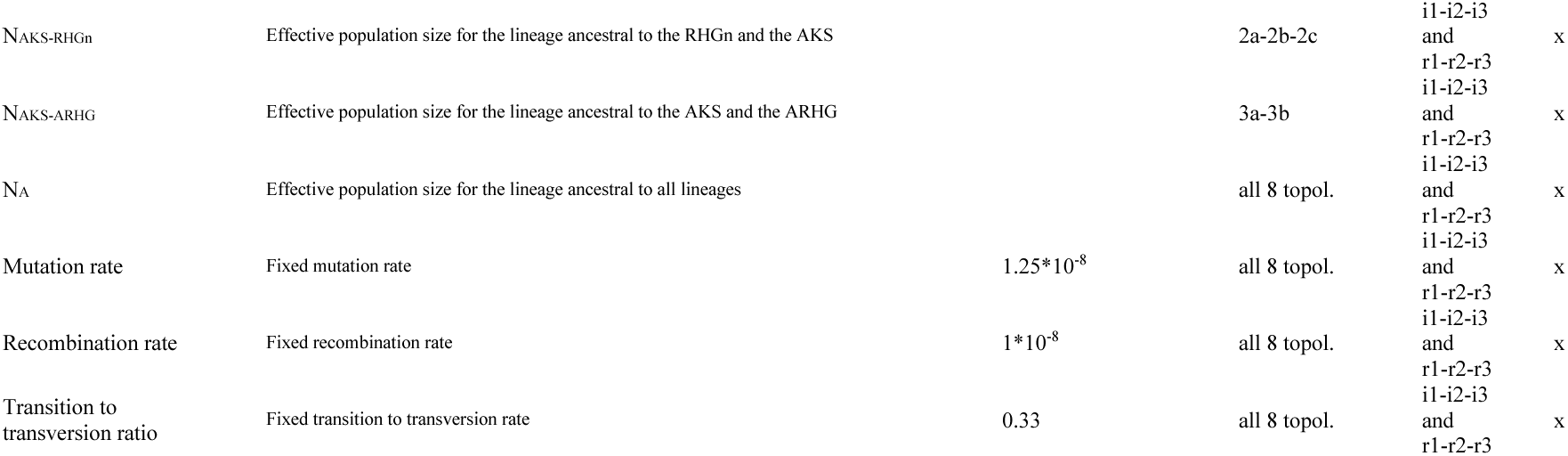
48 competing scenarios parameters’ description and prior distributions. Codes for scenarios topologies and gene-flow processes and all scenario parameters are detailed in **Figure 4** and in **Material and Methods**.

**Table S9:** Summary-statistics used in Random-Forest ABC scenario-choice and Neural-Network ABC posterior-parameter estimation. .

| Name of statistic (or group of statistics) | What was computed | Number of values | RF-ABC scenario-choice | NN-ABC posterior-parameter estimation |
| --- | --- | --- | --- | --- |
| Total number of biallelic sites among the five populations | sum | 1 | yes | yes |
| Total number of multiallelic sites among the five populations | sum | 1 | yes | yes |
| ASD distance within population for each five population separately | average and variance | 10 | yes | no |
| First five dimensions of the projection of the ASD matrix within population for each five population separately | average and variance | 50 | yes | yes |
| Proportion of biallelic sites within each five population separately | sum in a population divided by total number of biallelic sites in the dataset (no missing data) | 5 | yes | yes |
| Proportion of homozygous sites within each five population separately | sum in a population divided by total number of biallelic sites in the dataset (no missing data) | 5 | yes | yes |
| Proportion of homozygous ancestral sites within each five population separately | sum in a population divided by total number of homozygous sites in the population (no missing data) | 5 | yes | no |
| Proportion of private biallelic sites within each five population separately | sum in a population divided by total number of biallelic sites in the dataset (no missing data) | 5 | yes | yes |
| Frequency of the minor allele of the private biallelic sites within each five population separately | average and variance | 10 | yes | no |
| Expected heterozygosity at a segregating site within each five population separately | average and variance | 10 | yes | no |
| Tajima's D within each five population separately | average and variance for chromosomes with at least one variant | 10 | yes | yes |
| 1D Unfolded standardized site frequency spectrum (SFS) within each five population separately | proportion for each of 9 classes | 45 | yes | no |
| Watterson's theta within each five population separately | sum of biallelic sites divided by the harmonic number of the sample size (n=10) | 5 | yes | no |
| Total number of runs of homozygosity (ROH) within each five population separately | sum | 5 | yes | no |
| Total number of ROH by length class (<200 kbp, 200-500 kbp, >500 kbp) within each five population separately | sum | 15 | yes | yes |
| ROH length for each length class within each five population separately | average | 15 | yes | no |
| Total ROH length within each five population separately | average and variance | 10 | yes | no |
| ASD distance between each pair of five populations | average and variance | 20 | yes | no |
| First five dimensions of the projection of the ASD matrix between each pair of five populations | average and variance | 100 | yes | yes |
| Pairwise FST between each pair of five population | Pairwise FST (Weir and Cockerham 1984) | 10 | yes | yes |
| TOTAL number of statistics used in ABC inference |  |  | 337 | 202 |

**Table S10:** Correlations between standardized ASD matrices for each one of the 54 combinations of five population samples computed using 100 separate high-quality 1 Mb-long genomic windows, and corresponding individual-subsets of the standardized ASD matrix computed using the full genomic SNP set and used for Figure 2.

| Combinations of five populations used for ABC inferences |  |  |  |  | Spearman_ρ | 1000<br>Mantel<br>prem. p | Pearson_r | 1000<br>Mantel<br>prem. p |
| --- | --- | --- | --- | --- | --- | --- | --- | --- |
| nKS | sKS | RHGn | wRHG | eRHG |  |  |  |  |
| Ju'hoansi | Karretjie People | Ba.Konjo | Ba.Kola | Mbuti | 0.893 | <0.002 | 0.962 | <0.002 |
| Ju'hoansi | Karretjie People | Ba.Konjo | Ba.Kola | Nsua | 0.896 | <0.002 | 0.963 | <0.002 |
| Ju'hoansi | Karretjie People | Ba.Konjo | Baka | Mbuti | 0.898 | <0.002 | 0.964 | <0.002 |
| Ju'hoansi | Karretjie People | Ba.Konjo | Baka | Nsua | 0.903 | <0.002 | 0.967 | <0.002 |
| Ju'hoansi | Karretjie People | Ba.Konjo | Bi.Aka Mbat | Mbuti | 0.894 | <0.002 | 0.963 | <0.002 |
| Ju'hoansi | Karretjie People | Ba.Konjo | Bi.Aka Mbat | Nsua | 0.895 | <0.002 | 0.965 | <0.002 |
| Ju'hoansi | Karretjie People | Ba.Kiga | Ba.Kola | Ba.Twa | 0.901 | <0.002 | 0.961 | <0.002 |
| Ju'hoansi | Karretjie People | Ba.Kiga | Baka | Ba.Twa | 0.908 | <0.002 | 0.965 | <0.002 |
| Ju'hoansi | Karretjie People | Ba.Kiga | Bi.Aka Mbat | Ba.Twa | 0.887 | <0.002 | 0.961 | <0.002 |
| Ju'hoansi | Karretjie People | Ngumba | Ba.Kola | Ba.Twa | 0.885 | <0.002 | 0.960 | <0.002 |
| Ju'hoansi | Karretjie People | Ngumba | Ba.Kola | Mbuti | 0.907 | <0.002 | 0.963 | <0.002 |
| Ju'hoansi | Karretjie People | Ngumba | Ba.Kola | Nsua | 0.910 | <0.002 | 0.964 | <0.002 |
| Ju'hoansi | Karretjie People | Nzime | Baka | Ba.Twa | 0.914 | <0.002 | 0.968 | <0.002 |
| Ju'hoansi | Karretjie People | Nzime | Baka | Mbuti | 0.929 | <0.002 | 0.971 | <0.002 |
| Ju'hoansi | Karretjie People | Nzime | Baka | Nsua | 0.935 | <0.002 | 0.973 | <0.002 |
| Ju'hoansi | Karretjie People | Nzime | Bi.Aka Mbat | Ba.Twa | 0.896 | <0.002 | 0.965 | <0.002 |
| Ju'hoansi | Karretjie People | Nzime | Bi.Aka Mbat | Mbuti | 0.925 | <0.002 | 0.970 | <0.002 |
| Ju'hoansi | Karretjie People | Nzime | Bi.Aka Mbat | Nsua | 0.927 | <0.002 | 0.970 | <0.002 |
| Ju'hoansi | Nama | Ba.Konjo | Ba.Kola | Nsua | 0.932 | <0.002 | 0.966 | <0.002 |
| Ju'hoansi | Nama | Ba.Konjo | Baka | Nsua | 0.948 | <0.002 | 0.969 | <0.002 |
| Ju'hoansi | Nama | Ba.Konjo | Bi.Aka Mbat | Nsua | 0.944 | <0.002 | 0.967 | <0.002 |
| Ju'hoansi | Nama | Ba.Kiga | Ba.Kola | Ba.Twa | 0.948 | <0.002 | 0.964 | <0.002 |
| Ju'hoansi | Nama | Ba.Kiga | Baka | Ba.Twa | 0.958 | <0.002 | 0.966 | <0.002 |
| Ju'hoansi | Nama | Ba.Kiga | Bi.Aka Mbat | Ba.Twa | 0.945 | <0.002 | 0.962 | <0.002 |
| Ju'hoansi | Nama | Ngumba | Ba.Kola | Ba.Twa | 0.935 | <0.002 | 0.962 | <0.002 |
| Ju'hoansi | Nama | Ngumba | Ba.Kola | Nsua | 0.945 | <0.002 | 0.965 | <0.002 |
| Ju'hoansi | Nama | Nzime | Baka | Ba.Twa | 0.957 | <0.002 | 0.970 | <0.002 |
| Ju'hoansi | Nama | Nzime | Baka | Nsua | 0.963 | <0.002 | 0.974 | <0.002 |
| Ju'hoansi | Nama | Nzime | Bi.Aka Mbat | Ba.Twa | 0.945 | <0.002 | 0.965 | <0.002 |
| Ju'hoansi | Nama | Nzime | Bi.Aka Mbat | Nsua | 0.945 | <0.002 | 0.971 | <0.002 |
| !Xun | Karretjie People | Ba.Konjo | Ba.Kola | Nsua | 0.927 | <0.002 | 0.952 | <0.002 |
| !Xun | Karretjie People | Ba.Konjo | Baka | Nsua | 0.936 | <0.002 | 0.957 | <0.002 |
| !Xun | Karretjie People | Ba.Konjo | Bi.Aka Mbat | Nsua | 0.934 | <0.002 | 0.956 | <0.002 |
| !Xun | Karretjie People | Ba.Kiga | Ba.Kola | Ba.Twa | 0.923 | <0.002 | 0.949 | <0.002 |
| !Xun | Karretjie People | Ba.Kiga | Baka | Ba.Twa | 0.936 | <0.002 | 0.954 | <0.002 |
| !Xun | Karretjie People | Ba.Kiga | Bi.Aka Mbat | Ba.Twa | 0.922 | <0.002 | 0.950 | <0.002 |
| !Xun | Karretjie People | Ngumba | Ba.Kola | Ba.Twa | 0.913 | <0.002 | 0.948 | <0.002 |
| !Xun | Karretjie People | Ngumba | Ba.Kola | Nsua | 0.930 | <0.002 | 0.952 | <0.002 |
| !Xun | Karretjie People | Nzime | Baka | Ba.Twa | 0.945 | <0.002 | 0.959 | <0.002 |
| !Xun | Karretjie People | Nzime | Baka | Nsua | 0.956 | <0.002 | 0.963 | <0.002 |
| !Xun | Karretjie People | Nzime | Bi.Aka Mbat | Ba.Twa | 0.931 | <0.002 | 0.955 | <0.002 |
| !Xun | Karretjie People | Nzime | Bi.Aka Mbat | Nsua | 0.935 | <0.002 | 0.962 | <0.002 |
| !Xun | Nama | Ba.Konjo | Ba.Kola | Nsua | 0.932 | <0.002 | 0.954 | <0.002 |
| !Xun | Nama | Ba.Konjo | Baka | Nsua | 0.945 | <0.002 | 0.957 | <0.002 |
| !Xun | Nama | Ba.Konjo | Bi.Aka Mbat | Nsua | 0.937 | <0.002 | 0.956 | <0.002 |
| !Xun | Nama | Ba.Kiga | Ba.Kola | Ba.Twa | 0.933 | <0.002 | 0.948 | <0.002 |
| !Xun | Nama | Ba.Kiga | Baka | Ba.Twa | 0.942 | <0.002 | 0.951 | <0.002 |
| !Xun | Nama | Ba.Kiga | Bi.Aka Mbat | Ba.Twa | 0.920 | <0.002 | 0.947 | <0.002 |
| !Xun | Nama | Ngumba | Ba.Kola | Ba.Twa | 0.910 | <0.002 | 0.945 | <0.002 |
| !Xun | Nama | Ngumba | Ba.Kola | Nsua | 0.930 | <0.002 | 0.950 | <0.002 |
| !Xun | Nama | Nzime | Baka | Ba.Twa | 0.940 | <0.002 | 0.956 | <0.002 |
| !Xun | Nama | Nzime | Baka | Nsua | 0.954 | <0.002 | 0.962 | <0.002 |
| !Xun | Nama | Nzime | Bi.Aka Mbat | Ba.Twa | 0.915 | <0.002 | 0.951 | <0.002 |
| !Xun | Nama | Nzime | Bi.Aka Mbat | Nsua | 0.919 | <0.002 | 0.959 | <0.002 |

**Table S11:** Correlations between 332 summary-statistics computed on 100 genomic windows of 1Mb each used for ABC inferences and average values for 724 other such windows genome wide, for each one of the 54 combinations of five sampled-populations. We performed separately Spearman correlation tests and Wilcoxon two-sided tests for each one of the 54 combinations of sampled populations considered here between 332 summary-statistics used for ABC inferences computed on 100 genomic windows of 1Mb each and the same summary statistics computed for 724 other such windows genome-wide. Note that we excluded 5 variance related summary-statistics from the comparisons due to scaling irrelevances across sets of windows. Values in italic were not-significant. Values in bold were significant.

| Combinations of five populations used for ABC inferences |  |  |  |  | Spearman_ρ | p-value | Wilcoxon two-sided | p-value |
| --- | --- | --- | --- | --- | --- | --- | --- | --- |
| nKS | sKS | RHGn | wRHG | eRHG |  |  |  |  |
| Ju'hoansi | Karretjie People | Ba.Konjo | Ba.Kola | Mbuti | **0.90025** | <2.2x10-16 | 55505 | 0.8738 |
| Ju'hoansi | Karretjie People | Ba.Konjo | Ba.Kola | Nsua | **0.94243** | <2.2x10-16 | 55699 | 0.8124 |
| Ju'hoansi | Karretjie People | Ba.Konjo | Baka | Mbuti | **0.88935** | <2.2x10-16 | 54958 | 0.9505 |
| Ju'hoansi | Karretjie People | Ba.Konjo | Baka | Nsua | **0.97795** | <2.2x10-16 | 55561 | 0.8560 |
| Ju'hoansi | Karretjie People | Ba.Konjo | Bi.Aka Mbat | Mbuti | **0.88791** | <2.2x10-16 | 54975 | 0.9560 |
| Ju'hoansi | Karretjie People | Ba.Konjo | Bi.Aka Mbat | Nsua | **0.98917** | <2.2x10-16 | 55071 | 0.9869 |
| Ju'hoansi | Karretjie People | Ba.Kiga | Ba.Kola | Ba.Twa | **0.99203** | <2.2x10-16 | 55755 | 0.7949 |
| Ju'hoansi | Karretjie People | Ba.Kiga | Baka | Ba.Twa | **0.88887** | <2.2x10-16 | 55249 | 0.9560 |
| Ju'hoansi | Karretjie People | Ba.Kiga | Bi.Aka Mbat | Ba.Twa | **0.86963** | <2.2x10-16 | 54818 | 0.9055 |
| Ju'hoansi | Karretjie People | Ngumba | Ba.Kola | Ba.Twa | **0.93363** | <2.2x10-16 | 56089 | 0.6928 |
| Ju'hoansi | Karretjie People | Ngumba | Ba.Kola | Mbuti | **0.89903** | <2.2x10-16 | 55667 | 0.8225 |
| Ju'hoansi | Karretjie People | Ngumba | Ba.Kola | Nsua | **0.89407** | <2.2x10-16 | 55683 | 0.8174 |
| Ju'hoansi | Karretjie People | Nzime | Baka | Ba.Twa | **0.93279** | <2.2x10-16 | 55110 | 0.9995 |
| Ju'hoansi | Karretjie People | Nzime | Baka | Mbuti | **0.93047** | <2.2x10-16 | 54903 | 0.9328 |
| Ju'hoansi | Karretjie People | Nzime | Baka | Nsua | **0.97638** | <2.2x10-16 | 55518 | 0.8697 |
| Ju'hoansi | Karretjie People | Nzime | Bi.Aka Mbat | Ba.Twa | **0.91203** | <2.2x10-16 | 55193 | 0.9740 |
| Ju'hoansi | Karretjie People | Nzime | Bi.Aka Mbat | Mbuti | **0.88534** | <2.2x10-16 | 54846 | 0.9145 |
| Ju'hoansi | Karretjie People | Nzime | Bi.Aka Mbat | Nsua | **0.92615** | <2.2x10-16 | 55363 | 0.9193 |
| Ju'hoansi | Nama | Ba.Konjo | Ba.Kola | Nsua | **0.98709** | <2.2x10-16 | 55784 | 0.7859 |
| Ju'hoansi | Nama | Ba.Konjo | Baka | Nsua | **0.95181** | <2.2x10-16 | 55751 | 0.7961 |
| Ju'hoansi | Nama | Ba.Konjo | Bi.Aka Mbat | Nsua | **0.99637** | <2.2x10-16 | 55416 | 0.9023 |
| Ju'hoansi | Nama | Ba.Kiga | Ba.Kola | Ba.Twa | **0.94583** | <2.2x10-16 | 55820 | 0.7747 |
| Ju'hoansi | Nama | Ba.Kiga | Baka | Ba.Twa | **0.97595** | <2.2x10-16 | 55829 | 0.7719 |
| Ju'hoansi | Nama | Ba.Kiga | Bi.Aka Mbat | Ba.Twa | **0.94859** | <2.2x10-16 | 55672 | 0.8209 |
| Ju'hoansi | Nama | Ngumba | Ba.Kola | Ba.Twa | **0.89772** | <2.2x10-16 | 56235 | 0.6497 |
| Ju'hoansi | Nama | Ngumba | Ba.Kola | Nsua | **0.97607** | <2.2x10-16 | 56112 | 0.6859 |
| Ju'hoansi | Nama | Nzime | Baka | Ba.Twa | **0.89324** | <2.2x10-16 | 55925 | 0.7423 |
| Ju'hoansi | Nama | Nzime | Baka | Nsua | **0.89274** | <2.2x10-16 | 55425 | 0.8994 |
| Ju'hoansi | Nama | Nzime | Bi.Aka Mbat | Ba.Twa | **0.94542** | <2.2x10-16 | 55174 | 0.9801 |
| Ju'hoansi | Nama | Nzime | Bi.Aka Mbat | Nsua | **0.91175** | <2.2x10-16 | 55064 | 0.9847 |
| !Xun | Karretjie People | Ba.Konjo | Ba.Kola | Nsua | **0.93807** | <2.2x10-16 | 55773 | 0.7893 |
| !Xun | Karretjie People | Ba.Konjo | Baka | Nsua | **0.92892** | <2.2x10-16 | 55492 | 0.8780 |
| !Xun | Karretjie People | Ba.Konjo | Bi.Aka Mbat | Nsua | **0.99421** | <2.2x10-16 | 55025 | 0.9721 |
| !Xun | Karretjie People | Ba.Kiga | Ba.Kola | Ba.Twa | **0.94753** | <2.2x10-16 | 55946 | 0.7359 |
| !Xun | Karretjie People | Ba.Kiga | Baka | Ba.Twa | **0.94283** | <2.2x10-16 | 55320 | 0.9331 |
| !Xun | Karretjie People | Ba.Kiga | Bi.Aka Mbat | Ba.Twa | **0.88960** | <2.2x10-16 | 55579 | 0.8503 |
| !Xun | Karretjie People | Ngumba | Ba.Kola | Ba.Twa | **0.85660** | <2.2x10-16 | 55508 | 0.8729 |
| !Xun | Karretjie People | Ngumba | Ba.Kola | Nsua | **0.94814** | <2.2x10-16 | 55514 | 0.8709 |
| !Xun | Karretjie People | Nzime | Baka | Ba.Twa | **0.97947** | <2.2x10-16 | 55807 | 0.7787 |
| !Xun | Karretjie People | Nzime | Baka | Nsua | **0.89921** | <2.2x10-16 | 55432 | 0.8971 |
| !Xun | Karretjie People | Nzime | Bi.Aka Mbat | Ba.Twa | **0.94073** | <2.2x10-16 | 55083 | 0.9908 |
| !Xun | Karretjie People | Nzime | Bi.Aka Mbat | Nsua | **0.90798** | <2.2x10-16 | 54884 | 0.9267 |
| !Xun | Nama | Ba.Konjo | Ba.Kola | Nsua | **0.92443** | <2.2x10-16 | 55854 | 0.7642 |
| !Xun | Nama | Ba.Konjo | Baka | Nsua | **0.88787** | <2.2x10-16 | 56213 | 0.6561 |
| !Xun | Nama | Ba.Konjo | Bi.Aka Mbat | Nsua | **0.96611** | <2.2x10-16 | 55590 | 0.8468 |
| !Xun | Nama | Ba.Kiga | Ba.Kola | Ba.Twa | **0.94429** | <2.2x10-16 | 55924 | 0.7427 |
| !Xun | Nama | Ba.Kiga | Baka | Ba.Twa | **0.97282** | <2.2x10-16 | 55637 | 0.8319 |
| !Xun | Nama | Ba.Kiga | Bi.Aka Mbat | Ba.Twa | **0.94463** | <2.2x10-16 | 55365 | 0.9186 |
| !Xun | Nama | Ngumba | Ba.Kola | Ba.Twa | **0.85774** | <2.2x10-16 | 55709 | 0.8093 |
| !Xun | Nama | Ngumba | Ba.Kola | Nsua | **0.93875** | <2.2x10-16 | 56097 | 0.6904 |
| !Xun | Nama | Nzime | Baka | Ba.Twa | **0.94204** | <2.2x10-16 | 55685 | 0.8168 |
| !Xun | Nama | Nzime | Baka | Nsua | **0.95260** | <2.2x10-16 | 55684 | 0.8171 |
| !Xun | Nama | Nzime | Bi.Aka Mbat | Ba.Twa | **0.88873** | <2.2x10-16 | 55332 | 0.9292 |
| !Xun | Nama | Nzime | Bi.Aka Mbat | Nsua | **0.89805** | <2.2x10-16 | 55200 | 0.9718 |

**Table S12:** RF-ABC scenario-choice posterior predictions for each 54 combinations of five sampled populations separately. RF-ABC votes are provided for each competing scenario or group of scenarios, respectively, among the 1000 trees built in the corresponding random forest (see **Material and Methods**). Population combinations are provided in column 2 to 6. Each one of the table sheets correspond to the results provided in box-plot formats in **Figure 5**, **Figure S8**, **Figure S9**, and **Figure S10**, respectively. See “**Table S12_54popResABC_new_rev2.xlsx**” file as the table is too large to reasonably fit in a A4 page format.

**Table S13:** 25 combinations of five population samples out of 54 combinations tested for which Scenario i1-1b is best explaining observed data with Random-Forest ABC among the 48 competing scenarios. Values in the table correspond to posterior densities plotted in the left panels of **Figure 6**, **Figure 7**, **Figure S21**, and **Figure S22**. Parameter definitions and priors are provided in **Table S8** and represented graphically in the corresponding scenario panel of **Figure 4**.

|  | nKS | sKS | RHGn | wRHG | eRHG |
| --- | --- | --- | --- | --- | --- |
| *Combination 1* | Ju\|'hoansi | Karretjie People | Nzime | Baka | Nsua |
| *Combination 2* | Ju\|'hoansi | Karretjie People | Ngumba | Ba.Kola | Nsua |
| *Combination 3* | Ju\|'hoansi | Karretjie People | Ngumba | Ba.Kola | Ba.Twa |
| *Combination 4* | Ju\|'hoansi | Karretjie People | Nzime | Bi.Aka_Mbati | Nsua |
| *Combination 5* | Ju\|'hoansi | Karretjie People | Nzime | Bi.Aka_Mbati | Ba.Twa |
| *Combination 6* | Ju\|'hoansi | Karretjie People | Ngumba | Ba.Kola | Mbuti |
| *Combination 7* | Ju\|'hoansi | Karretjie People | Nzime | Bi.Aka_Mbati | Mbuti |
| *Combination 8* | Ju\|'hoansi | Karretjie People | Ba.Konjo | Baka | Mbuti |
| *Combination 9* | Ju\|'hoansi | Karretjie People | Ba.Konjo | Ba.Kola | Mbuti |
| *Combination 10* | Ju\|'hoansi | Karretjie People | Ba.Konjo | Bi.Aka_Mbati | Mbuti |
| *Combination 11* | Ju\|'hoansi | Nama | Ba.Konjo | Baka | Nsua |
| *Combination 12* | !Xun | Nama | Ba.Konjo | Baka | Nsua |
| *Combination 13* | Ju\|'hoansi | Nama | Nzime | Baka | Nsua |
| *Combination 14* | Ju\|'hoansi | Nama | Ngumba | Ba.Kola | Nsua |
| *Combination 15* | Ju\|'hoansi | Nama | Nzime | Baka | Ba.Twa |
| *Combination 16* | Ju\|'hoansi | Nama | Ngumba | Ba.Kola | Ba.Twa |
| *Combination 17* | Ju\|'hoansi | Nama | Nzime | Bi.Aka_Mbati | Ba.Twa |
| *Combination 18* | Ju\|'hoansi | Nama | Ba.Konjo | Ba.Kola | Nsua |
| *Combination 19* | Ju\|'hoansi | Nama | Ba.Kiga | Ba.Kola | Ba.Twa |
| *Combination 20* | Ju\|'hoansi | Nama | Ba.Kiga | Bi.Aka_Mbati | Ba.Twa |
| *Combination 21* | !Xun | Nama | Nzime | Bi.Aka_Mbati | Nsua |
| *Combination 22* | !Xun | Nama | Nzime | Bi.Aka_Mbati | Ba.Twa |
| *Combination 23* | !Xun | Karretjie People | Nzime | Bi.Aka_Mbati | Nsua |
| *Combination 24* | !Xun | Karretjie People | Nzime | Bi.Aka_Mbati | Ba.Twa |
| *Combination 25* | !Xun | Karretjie People | Ba.Konjo | Bi.Aka_Mbati | Nsua |

**Table S14:** NN-ABC posterior parameter estimation of all parameters in Scenario i1-1b for results from each 25 sets of five populations each, separately. Table summarizing posterior parameter distributions obtained with Neural Network ABC for each 25 combinations of sampled-populations separately. Full posterior distributions are provided in plot formats in main **Figure 6** (divergence times t), **Figure 7** (admixture times tad), supplementary **Figure S21** (Ne), and supplementary **Figure S22** (asymmetric admixture rates). Column-headers indicate the categorization of sampled populations into Northern Khoisan (nKS), Southern Khoisan (sKS), Rainforest Hunter-Gatherer neighbors (RHGn), Western Rainforest Hunter-Gatherer (wRHG), or Eastern Rainsforest Hunter-Gatherer (eRHG) groups, as per scenario topology and simulation design explicated in **Material and Methods** and in **Figure 4**. Note that each sampled population considered in the RF-ABC scenario choice analyses are represented twice or more in the 25 combinations. “CI” stands for ABC Credibility Intervals. See main **Figure 4**, supplementary **Table S8** and **Material and Methods** for detailed scenario-parameter descriptions. **Table S15** provides Pearson correlations between each pair of NN-ABC posterior-parameter estimates (median or mode values) for the 25 combinations of five populations. See “**Table S14_new_rev2.xlsx**” file as the table is too large to reasonably fit in a A4 page format.

**Table S15:** Pearson correlations between each pair of NN-ABC posterior-parameter estimates (median or mode values) for the 25 combinations of five populations provided in Table S14. Each parameter inferred in Scenario i1-1b using NN-ABC for each one of the 25 combinations of sampled populations (**Table S14**), is indicated in the first column and the first row of the matrix. Pearson rho values are indicated in the lower triangle, and corresponding Pearson correlation test p-values are indicated in the upper triangle of the matrix. The first spreadsheet corresponds to tests computed on the median point estimate of **Table S14**; the second spreadsheet correspond to the same testing considering mode point-estimate instead. Significant correlation results after Bonferroni correction for multiple testing (separately for the median and mode results) are indicated in bold with “**” for p-values < 1% and “*” for p-values between 1% and 5%. Note that only 16 pairs of parameters (on 903 tested pairs), were significant (11 with p-value< 1%, and five with p-value between 1% and 5%), when considering the median point posterior-estimates, and 12, (9 with p-value< 1%, and three with p-value between 1% and 5%), when considering the mode instead. See “**Table S15_corparam_new_rev2.xls**” file as the table is too large to reasonably fit in a A4 page format.

**Table S16:** NN-ABC posterior parameter estimation cross-validation errors of all parameters in Scenario i1-1b for results from 25 sets of five populations each. Cross-validation errors are based on 50 (5%) randomly chosen simulation out of the 1000 simulations closest to the observed data for each 25 combinations of sampled populations, respectively, each considered in-turn as pseudo-observed data and for which NN-ABC joint posterior parameter estimations were performed with the same hyper-parameters. For each parameter and each 25 combinations of sampled populations, the first column provides variance-scaled mean-squared errors and the second column provides simple average estimation bias (see **Material and Methods**). Relatively high variance-scaled mean-squared errors (close-to or above 1) are highlighted in grey. See “**Table S16_new2_rev2.xlsx**” file as the table is too large to reasonably fit in a A4 page format.

**Table S17:** NN-ABC posterior parameter estimation of all parameters in Scenario i1-1b for results from 25 sets of five populations each, combined altogether. Entire posterior distributions for each 25 analyses separately (**Table S14**), for which summary statistics are presented in **Table S3**, are combined (concatenated) together before re-computing the Mode, Median, Mean, 50% CI and 90% CI presented in this table. Therefore, values in the table correspond to posterior densities plotted in the right panels of **Figure 6**, **Figure 7**, **Figure S21**, and **Figure S22**. Parameter definitions and priors are provided in **Table S8** and represented graphically in the corresponding scenario panel of **Figure 4**. Values in this table are used for the synthetic schematic representation of the demographic and migration history of Central and Southern African populations presented in **Figure 8**. Values in italic are not satisfactorily departing from the priors and are associated with high cross-validation posterior parameter estimation errors in the vicinity of the observed data (**Table S16**).

| Parameter | Mode | Median | Mean | 50% CI | 90% CI |
| --- | --- | --- | --- | --- | --- |
| N<sub>nKS</sub> | 13387 | 27960 | 33648 | 16400-46042 | 8648-78425 |
| N<sub>sKS</sub> | 11496 | 22584 | 29465 | 12698-40535 | 6158-76677 |
| N<sub>wRHG</sub> | 17083 | 23544 | 29128 | 15201-37922 | 7606-71133 |
| N<sub>eRHG</sub> | 10702 | 17589 | 24112 | 10882-31019 | 5769-67086 |
| N<sub>RHGn</sub> | 16730 | 31250 | 34896 | 18914-47062 | 9111-74177 |
| N<sub>AKS</sub> | 62991 | 57534 | 55788 | 39887-72762 | 16051-90666 |
| N<sub>ARHG</sub> | 53641 | 51921 | 51674 | 33228-69906 | 12751-90806 |
| N<sub>ARHG-RHGn</sub> | 61127 | 55333 | 53839 | 35566-73204 | 11601-91048 |
| N<sub>A</sub> | 17005 | 26518 | 31283 | 17496-40248 | 9890-69775 |
| t<sub>KS</sub> | 2857 | 2881 | 2969 | 2304-3509 | 1624-4662 |
| t<sub>RHG</sub> | 751 | 970 | 1089 | 670-1374 | 370-2222 |
| t<sub>RHG-RHGn</sub> | 6347 | 6478 | 6617 | 5462-7566 | 4098-9734 |
| t<sub>KS-RHG-RHGn</sub> | 13254 | 12412 | 11909 | 10715-13411 | 8044-14397 |
| tad<sub>nKS-sKS</sub> | 550 | 611 | 718 | 434-847 | 227-1580 |
| tad<sub>nKS-RHGn</sub> | 1241 | 1406 | 1524 | 969-1956 | 447-3003 |
| tad<sub>sKS-RHGn</sub> | 1045 | 1243 | 1354 | 829-1739 | 406-2678 |
| tad<sub>wRHG-RHGn</sub> | 239 | 310 | 373 | 197-478 | 86-874 |
| tad<sub>eRHG-RHGn</sub> | 277 | 446 | 528 | 278-694 | 135-1190 |
| tad<sub>wRHG-eRHG</sub> | 332 | 449 | 547 | 290-701 | 130-1263 |
| tad<sub>RHGn-ARHG</sub> | 2905 | 3161 | 3369 | 2370-4114 | 1435-6007 |
| tad<sub>AKS-RHGn</sub> | 4492 | 4483 | 4582 | 3710-5303 | 2728-6825 |
| tad<sub>AKS-ARHG</sub> | 4228 | 4427 | 4579 | 3691-5261 | 2814-6954 |
| tad<sub>AKS-ARHG-RHGn</sub> | 8881 | 9332 | 9307 | 7970-10762 | 5923-12505 |
| m(nKS-sKS) | 0.6058 | 0.6129 | 0.5888 | 0.4309-0.7707 | 0.1613-0.9241 |
| m(sKS-nKS) | 0.7823 | 0.6187 | 0.5925 | 0.4298-0.7824 | 0.1544-0.9309 |
| m(nKS-RHGn) | 0.3921 | 0.4800 | 0.4918 | 0.3218-0.6569 | 0.1356-0.8794 |
| m(RHGn-nKS) | 0.4897 | 0.5207 | 0.5203 | 0.3162-0.7340 | 0.0859-0.9382 |
| m(sKS-RHGn) | 0.5539 | 0.5241 | 0.5215 | 0.3463-0.6994 | 0.1343-0.8974 |
| m(RHGn-sKS) | 0.4941 | 0.4824 | 0.4816 | 0.2931-0.6633 | 0.0872-0.8868 |
| m(RHGn-wRHG) | 0.3666 | 0.5076 | 0.5055 | 0.2935-0.7189 | 0.0839-0.9216 |
| m(wRHG-RHGn) | 0.4899 | 0.5039 | 0.5038 | 0.3374-0.6693 | 0.1311-0.8837 |
| m(RHGn-eRHG) | 0.4183 | 0.4678 | 0.4749 | 0.2694-0.6774 | 0.0795-0.8886 |
| m(eRHG-RHGn) | 0.7038 | 0.6242 | 0.5957 | 0.4363-0.7746 | 0.1733-0.9270 |
| m(wRHG-eRHG) | 0.5720 | 0.5104 | 0.5087 | 0.3200-0.6976 | 0.1213-0.8929 |
| m(eRHG-wRHG) | 0.5011 | 0.4922 | 0.4936 | 0.3066-0.6788 | 0.0969-0.8911 |
| m(RHGn-ARHG) | 0.6155 | 0.5087 | 0.5011 | 0.3188-0.6820 | 0.1078-0.8798 |
| m(ARHG-RHGn) | 0.5104 | 0.4846 | 0.4893 | 0.2734-0.6991 | 0.0869-0.9104 |
| m(ARHG-AKS) | 0.6493 | 0.5414 | 0.5299 | 0.3316-0.7375 | 0.0935-0.9294 |
| m(AKS-ARHG) | 0.5627 | 0.4814 | 0.4807 | 0.2950-0.6624 | 0.0897-0.8766 |
| m(RHGn-AKS) | 0.5286 | 0.4940 | 0.4913 | 0.3026-0.6776 | 0.0998-0.8863 |
| m(AKS-RHGn) | 0.5057 | 0.4919 | 0.4888 | 0.3026-0.6751 | 0.0912-0.8781 |
| m(AKS-ARHG-RHGn) | 0.5948 | 0.5145 | 0.5096 | 0.3263-0.6970 | 0.1123-0.8887 |
| m(ARHG-RHGn-AKS) | 0.2813 | 0.4507 | 0.4642 | 0.2482-0.6707 | 0.0761-0.8966 |

**Table S18:**
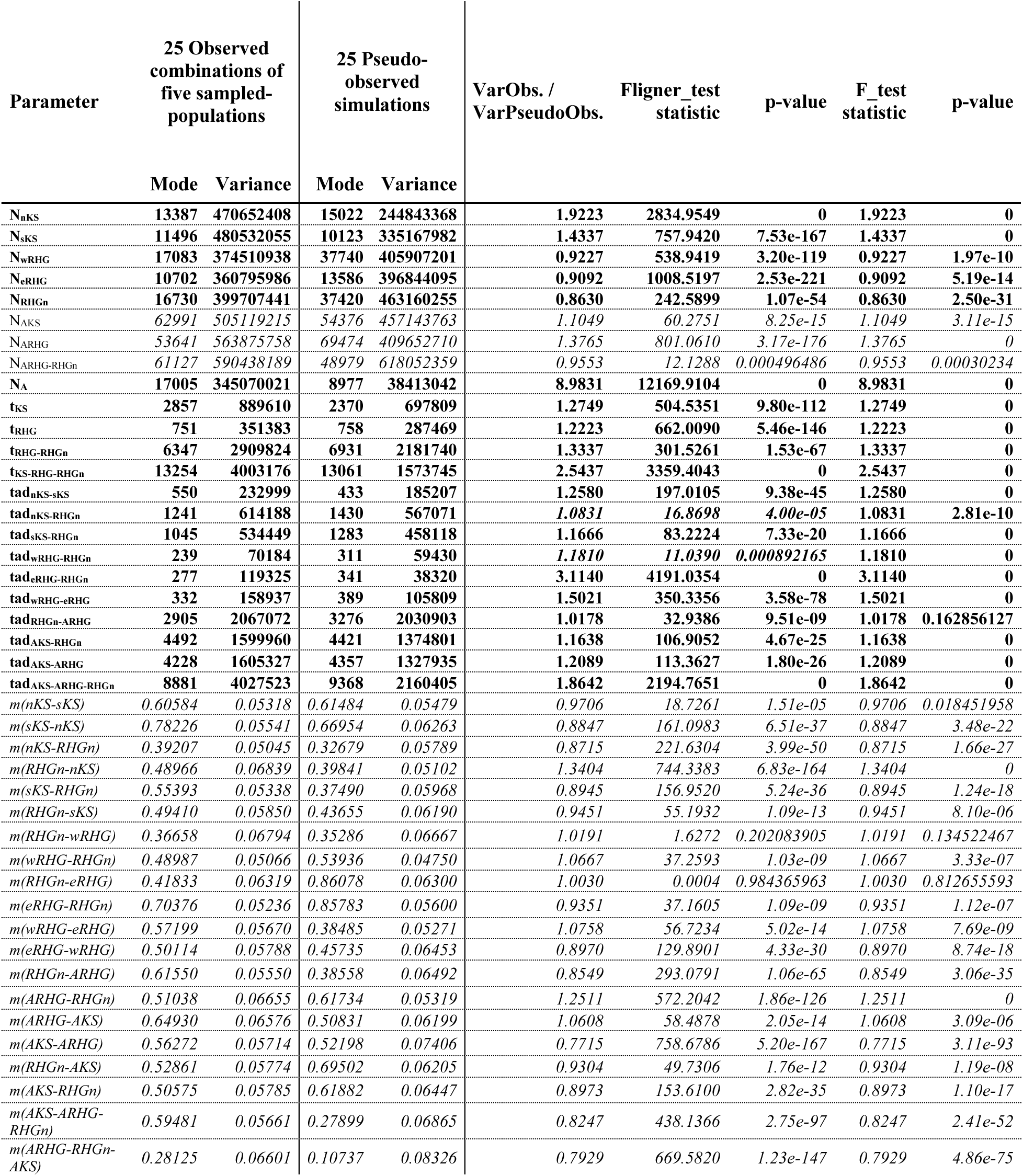
NN-ABC parameter inference posterior check: Fligner and F tests of homogeneity of variance between the NN-ABC posterior-parameter combined (concatenated) distributions obtained for 25 different observed combinations of five sampled-populations and 25 pseudo-observed simulated datasets under the Scenario i1-1b proposed in Figure 8 and Table S17. Each line corresponds to a single scenario-parameter under Scenario i1-1b (**Figure 4** and **Table S8**). For each parameter in Scenario i1-1b, the first two columns provide the Mode and Variance of posterior distributions plotted in the right panels of **Figure 6**, **Figure 7**, **Figure S21**, and **Figure S22**, and corresponding **Table S17** and synthetic **Figure 8**. The third and fourth column provide the mode and Variance of NN-ABC posterior-parameter estimated distributions obtained as for the 25 observed datasets but using instead 25 pseudo-observed simulated datasets, each obtained using the same point-value of scenario-parameters provided in the first column (see **Material and Methods**). Therefore, values in the third and fourth column correspond to the Mode and Variance of the combined (concatenated) densities of **Figure S50**, **Figure S51**, **Figure S52**, and in **Figure S53**. Fifth column corresponds to scaled difference in observed variance between NN-ABC results obtained from observed data and NN-ABC results obtained from pseudo-observed simulations. Sixth and Seventh columns present results from Fligner tests of homogeneity of variance performed between NN-ABC posterior parameter combined distributions obtained between observed data and pseudo-observed simulations. The eight and ninth column present results from simple F-tests of homogeneity of variance between the same distributions. Values in italic correspond to scenario-parameters NN-ABC posterior estimates that are not satisfactorily departing from the priors and are associated with large posterior CI and high cross-validation posterior parameter estimation errors in the vicinity of the observed data (**Table S16**).

